# NanoFAST: Structure-based design of a small fluorogen-activating protein with only 98 amino acids

**DOI:** 10.1101/2020.12.29.424647

**Authors:** Konstantin S. Mineev, Sergey A. Goncharuk, Marina V. Goncharuk, Natalia V. Povarova, Nadezhda S. Baleeva, Alexander Yu. Smirnov, Ivan N Myasnyanko, Dmitry A. Ruchkin, Sergey Bukhdruker, Alina Remeeva, Alexey Mishin, Valentin Borshchevskiy, Valentin Gordeliy, Alexander S. Arseniev, Dmitriy A. Gorbachev, Alexey S. Gavrikov, Alexander S. Mishin, Mikhail S. Baranov

## Abstract

One of the essential characteristics of any tag used in bioscience and medical applications is its size. The larger the label, the more it may affect the studied object, and the more it may distort its behavior. In this paper, using NMR spectroscopy and X-ray crystallography, we have studied the structure of fluorogen-activating protein FAST both in the apo form and in complex with the fluorogen. We shown that significant change in the protein occurs upon interaction with the ligand. While the protein is completely ordered in the complex, its apo form is characterized by higher mobility and disordering of its N-terminus. We used structural information to design the shortened FAST (which we named nanoFAST) by truncating 26 N-terminal residues. Thus, we created the shortest genetically encoded tag among all known fluorescent and fluorogen-activating proteins, which is composed of only 98 amino acids.

## Introduction

Various fluorescent tags have long and widely been used in modern biomolecular research.[1] Over the past decades, many such tags have been developed, mostly fluorescent proteins (FP),[2] which are genetically encoded tags formed solely from the internal components of a biological object. Alternatively, completely external chemical tags can also be used.[3] However, the greatest interest in recent years has been acquired by the combined tags, with one of the components being internally genetically encoded, while the second, small-molecule component is supplied from the outside. The best known labels of this kind are two-component Halo-,[4] and SNAP-tags[5] or three-component system based on mutant forms of lipoic acid ligase.[6] Nevertheless, all these tags have a number of drawbacks. Fluorescent proteins are quite large and require a considerable time and the presence of oxygen for maturation[7], while the use of chemical fluorescent dyes in any role often leads to off-target labeling.[8]

In this regard, the approaches that employ the so-called fluorogens-substances with very weak fluorescence in the free state, which become bright only when they bind to the tag’s secondary component look more promising. The internal component of such labels can be nucleic acid[9] or fluorogen-activating protein (FAP).[10] Such tags do not require oxygen and can be used under anaerobic conditions. Their maturation time is small and corresponds to the time of protein folding, while the fluorescent signal can be induced or removed on demand by simple addition or washing out of fluorogen.[11] The size of the tag is also an important characteristic. The larger it is, the more it affects the natural dynamics of the tagged protein.[12] Apart from their other advantages, FAPs are almost two times smaller than FP. Nevertheless, the size of such proteins is still about 120-150 amino acids while the shorter FAPs are too dim to be used for imaging.[13]

One of the most promising protein among various FAPs is the so-called FAST protein (“Fluorescence-Activating and absorption-Shifting Tag”)[14], an engineered variant of the photoreceptor from *Halorhodospira halophila* - Photoactive Yellow Protein (PYP).[15] This photoactive protein covalently binds a hydroxycinnamic acid. However, the replacement of several amino acids (including the key cysteine responsible for covalent binding) allowed using it as an FAP with a group of fluorogens. Over the past five years, a series of multi-colored variants [16] and split constructs [17] have been created based on this FAP.

Nevertheless, the structure of this protein and its complexes was unknown. It was only evident that the interaction with fluorogens resembled the binding of the native ligand of the PYP, since in the FAST:fluorogen complex, the phenolic fragment of the fluorogen was deprotonated, probably due to interaction with amino acids E46 and Y42.[18]

In this work, we have used NMR spectroscopy to study the structure of both the FAST apo form and its complex with the previously proposed fluorogen **N871b** (Fig. 4).[19] We have shown that significant change in the protein occurs upon interaction with the ligand. Using the data, we found that the N-terminally truncated variant of FAST protein (which we named nanoFAST) can also be used as an FAP.

## Results and Discussion

Through the whole of our study, we crystallized the FAST protein several times, either in the presence of various ligands or without them. Surprisingly, we found that the protein adopts a form of the ligand-free domain-swapped dimer in crystals under a wide variety of conditions. In this dimer, the first three strands of the core β sheet (A30-L33, I39-N43, and T50-R52) are exchanged with the corresponding elements from the symmetrical molecule, forming together one large twelve-stranded β sheet (Figure 1 SI Part 3).

**Figure 1.**
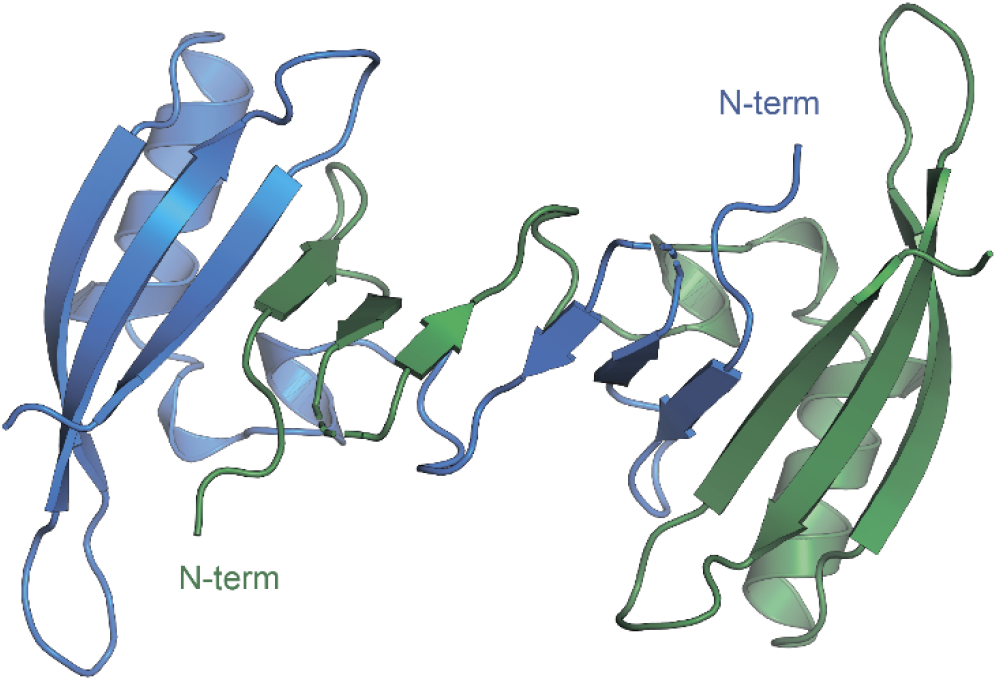
Crystal structure of FAST in the domain-swapped dimer form.

To further investigate the FAST ligand binding, we produced the ^13^C/^15^N isotope-labeled FAST and solved the spatial structure of the protein in complex with a promising ligand **N871b** previously proposed by us[19] and in the apo state using NMR spectroscopy (SI Part 4). Contrary to X-ray data and in good correlation with reported previously,[14] both forms of protein are present in solution exclusively in the monomeric form (the determined hydrodynamic radius was 2.1±0.1 nm, which corresponds to the 15 kDa).

Initial characterization of FAST-apo revealed the poor quality of NMR spectra due to the enhanced slow conformational transitions (Figures S4.1, S4.2, S4.3). In contrast, ligand binding stabilized the protein substantially, providing a perfect NMR spectrum. Thus, we first investigated the structure of FAST in complex with **N871b**. The high quality of NMR data allowed obtaining 97% of possible chemical shift assignment and determining the structure in a semi-automated manner, with the intermolecular distances being observed directly via the isotope-filtered experiments (Figure S4.4).[20] In complex with **N871b**, FAST reveals the typical architecture of a PAS domain,[21] similar to the fold of its parent PYP protein (Figure 2). [22] According to the PDBeFOLD server, the backbone atoms of the complex may be superimposed with the coordinates of PYP (PDBID 1F98 [22]) with the RMSD of 1.16 Å.

**Figure 2.**
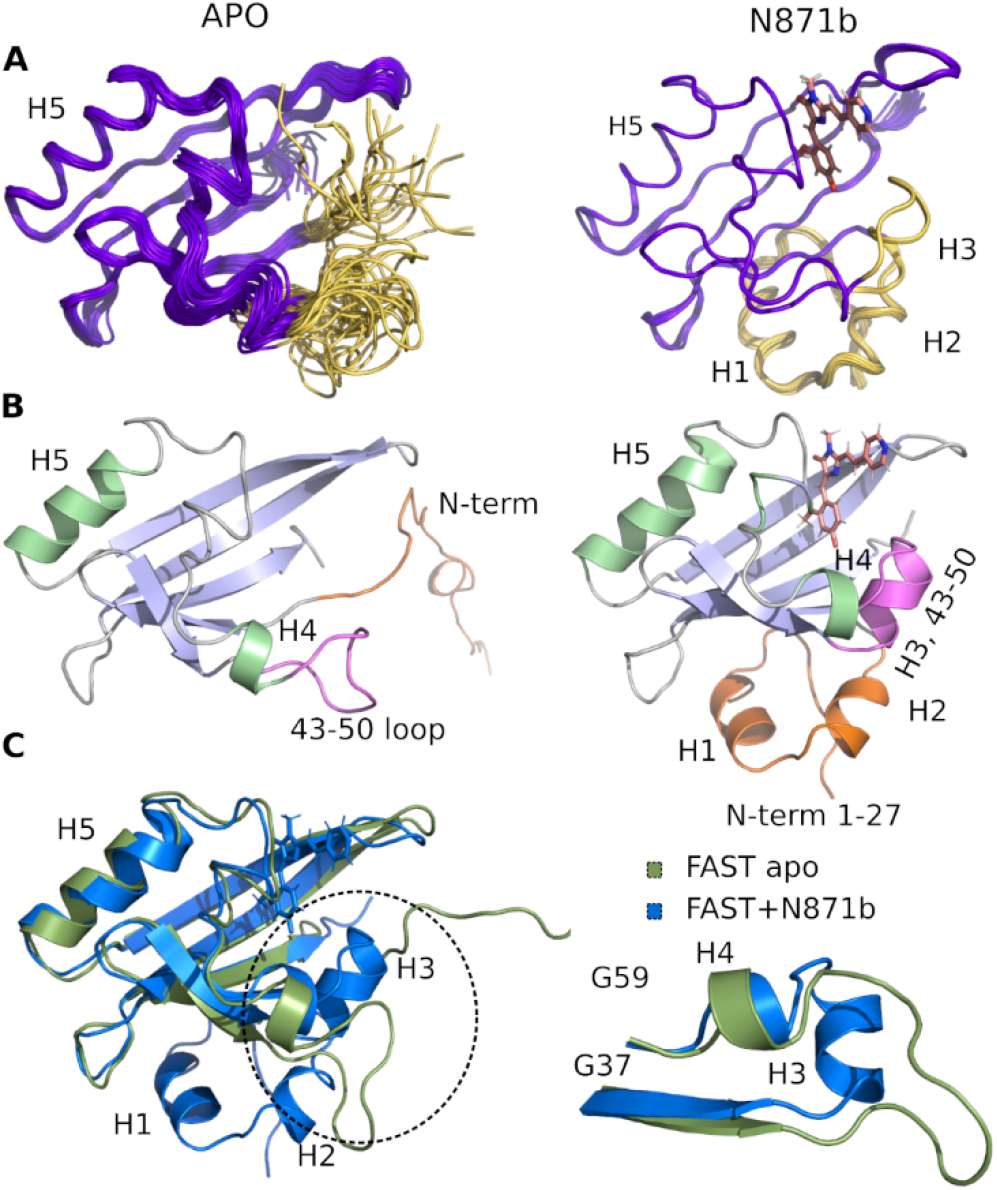
3D Structure of FAST-apo (**A**, **B** left, superposition on **C**) and FAST:**N871b** complex (**A**, **B** right, superposition on **C**). **A** - 20 best NMR structures, superimposed over the backbone atoms of the secondary structure elements. N-terminus and region 43-50 that change the structure upon ligand binding are shown in ivory. **B** - representative structures. α-helices are shown in green and β-strands are shown in blue. N-terminus and region 43-50 are highlighted by orange and magenta, respectively. **C** – structures of FAST-apo (green) and FAST:**N871b** (blue), superimposed over the backbone atoms of 5 β-strands and helix H5. Unstructured C-terminal His6 tag is not shown.

FAST chain forms a 5-stranded β-sheet and 5 α-helices, and the ligand is placed inside a hydrophobic cavity, stabilized by three hydrogen bonds (Figure 3). Like 4-hydroxycinnamic acid residue in PYP, the oxygen of **N871b** phenol ring forms an H-bond with the protonated sidechain of E46 (Hε2) and a more distant polar contact with phenol group of Y42 (Hη), which is supported by the presence of two broad low-field peaks in ^1^H NMR spectra of these labile protons (Figure 3). In addition, the carbonyl group of a 5-member ring is engaged in an H-bond with the ε1-imino-group of W94, which explains the necessity of this residue identified during the creation of the FAST protein.[14] Besides, the ligand binding is favored by the interactions with hydrophobic sidechains of I31, T50, V66, A67, P68, T70, I96, P97, V107 and V122 and π-stacking with the rings of F62 and F75. The ligand molecule is larger than the cavity, and it exits the protein near the C-terminus of helix 3, with the direct contact of the solvent-exposed pyridine group with the positively charged R52 sidechain.

**Figure 3.**
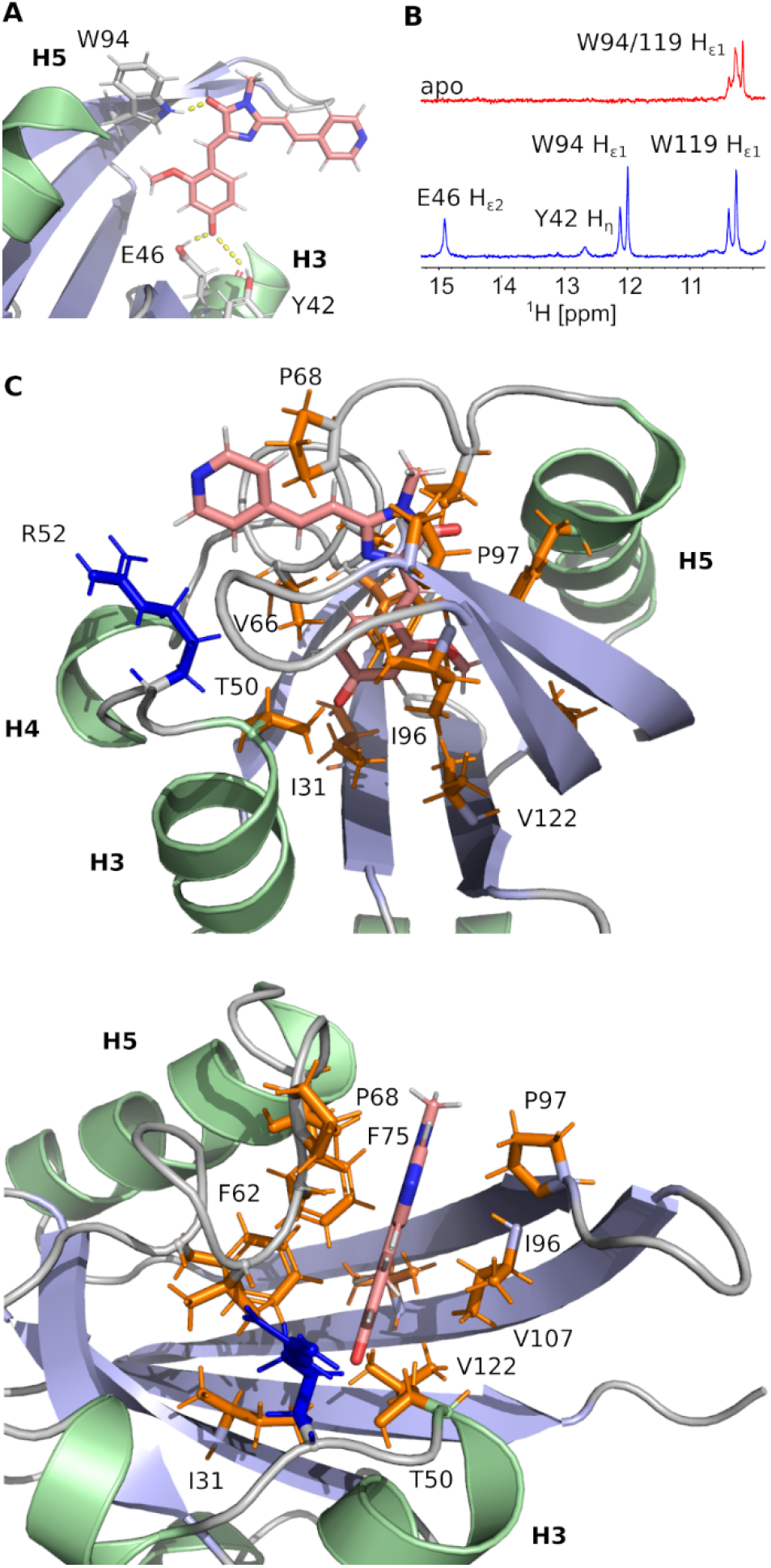
Details of the **N871b** binding by FAST. **A** – hydrogen bonds involved in **N871b** binding. **B** – a low-field region of 1H NMR spectra of FAST in the apo state (red) and in complex with **N871b**(blue). Both spectra were recorded at pH 7.0, 25oC. The assignment of protons is indicated. **C** – non-bonding interactions, detected within the FAST:**N871b** complex. Sidechains of hydrophobic residues are painted in orange, R52 positively charged sidechain is shown in blue.

To investigate the structure of ligand-free FAST, we had to heat the protein to 45oC and use a lower-field NMR magnet. Together, these two actions allowed reducing the effect of slow conformational transitions and solving the spatial structure (Table S4.1). The initial analysis revealed a drastic difference between the apo and bound states of FAST (Figures 2 and S4.5). The whole N-terminal part, which includes helices H1 and H2 appeared completely disordered in the apo state. Region of helix H3, containing three residues, engaged in the ligand binding in PYP and FAST:**N871b** complex, became unstructured. However, the coordinates of the remaining elements of secondary structure - a 5-stranded β-sheet and helix H5 remain unchanged with respect to the FAST:**N871b** complex and could be superimposed with the RMSD as small as 0.9 Å. Thus, the ligand binding to FAST induces the formation of helix H3 and stabilizes the N-terminal residues. This behavior can possibly lead to dimer formation upon protein crystallization and correlates well with the proposed PYP action mechanism - the apo-state of FAST is highly similar to the light-induced state of PYP.[23]

Since the ligand binding should begin with the initial apo form before the protein rearrangement, we hypothesized that the binding can occur in the absence of N-terminus and the shortened FAST can retain fluorogen-activating properties.

Thus, next, we created an N-terminally shortened variant of FAST protein (truncated up to the 26th residue - nanoFAST). We found that it is inactive against such known fluorogens as **HMBR**, **HBR-DOM**, **N871b**, and others (SI, Part 5). Since the protein pocket is slightly enlarged in the apo form, the pocket of nanoFAST also should be bigger than in the original FAST. Thus, at the next stage, we created a library of compounds with enlarged benzylidene moiety (Figure 4, SI Part 9).

**Figure 4.**
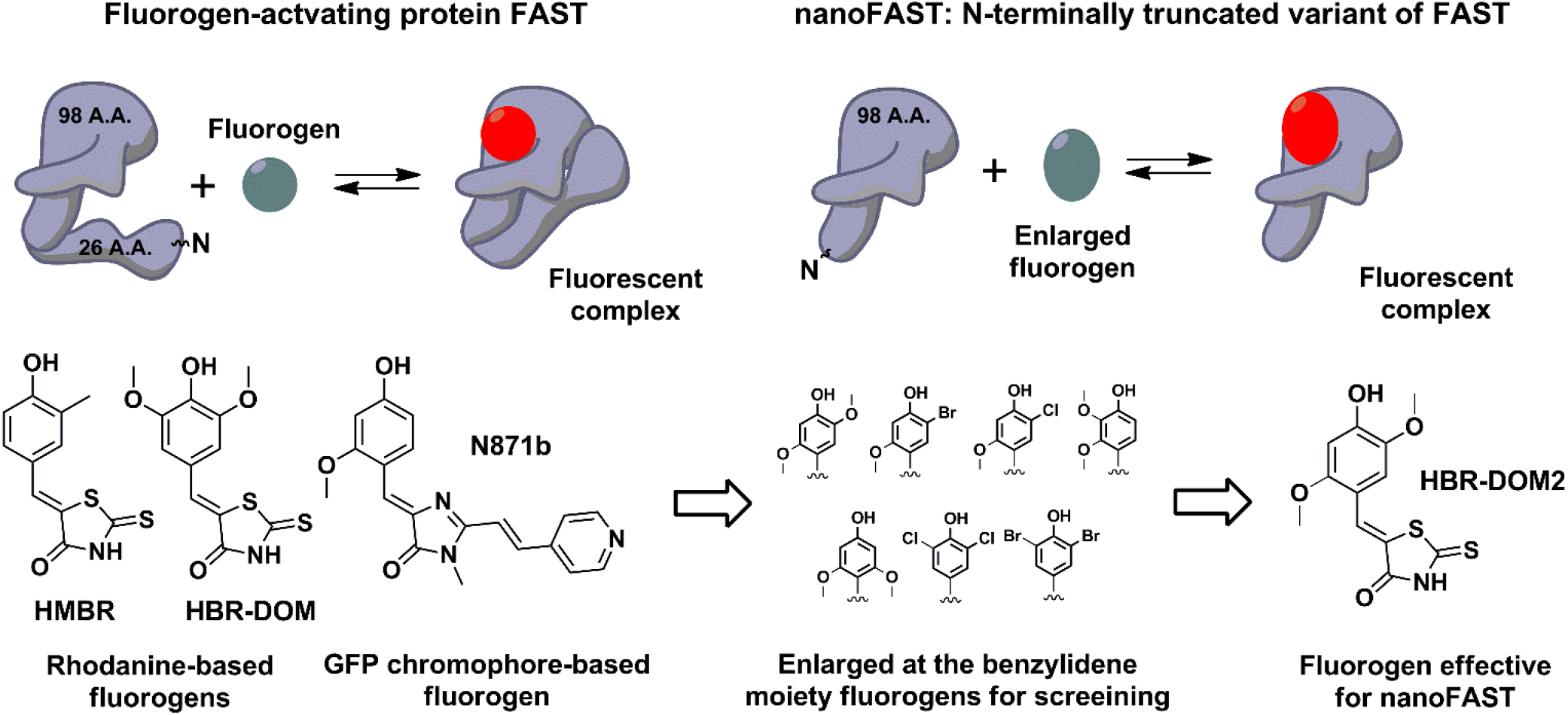
Principal scheme of FAST protein action revealed by NMR analysis, proposed nanoFAST protein and their fluorogens.

The screening of this library (SI Part 5) showed that the introduction of additional bulky substituents allows recovering the ligand binding with the protein. It turned out to be most effective in the case of 2.5 disubstituted substances, and especially 2.5 dimethoxy rhodanine - compound **HBR-DOM2**. This substance’s fluorescence intensity increased by more than a hundred times upon interaction with the nanoFAST protein. The fluorescence quantum yield of the complex reached 55%, while the binding constant turned out to be close to 1 μM, which is similar to the characteristic of previously obtained pairs with FAST (Table 1, SI Parts 6 and 7).

**Table 1.** Optical properties of chromophores and their complexes with proteins

| Complex | Absorbance maxima position, nm<sup>a</sup> | | | Emission maxima position, nm<sup>a</sup> | | | $\epsilon$, M<sup>-1</sup>·cm<sup>-1</sup><sub>b</sub> | Fluorescence quantum yield (FQY), %<sup>b</sup> | Brightness (=ε·FQY) | K<sub>d</sub>, μM |
| --- | --- | --- | --- | --- | --- | --- | --- | --- | --- | --- |
|  | neutral | anionic | bound | neutral | anionic | bound |  |  |  |  |
| **FAST:HMBR**<sup>[14,19]</sup> | 400 | 461 | 483 | 493 | 560 | 540 | 45000 | 33 | 14850 | 0.13 |
| **FAST:HBR-DOM**<sup>[14,19]</sup> | 406 | 486 | 520 | 530 | 600 | 600 | 39000 | 31 | 12000 | 0.97 |
| **FAST:N871b**<sup>[19]</sup> | 455 | 528 | 562 | 570 | 630 | 606 | 23000 | 25 | 5750 | 0.25 |
| **FAST:HBR-DOM2** | ~420 | 492 | 510 | ~500 | 572 | 566 | 30500 | 54±3 | 16500 | 0.021±0.001 |
| **nanoFAST:HBR-DOM2** |  |  | 502 |  |  | 563 | 25500 | 56±3 | 14300 | 0.85±0.01 |

a – characteristics of neutral forms were investigated in 1mM phosphate buffer pH 5.5, anionic forms – in 1mM phosphate buffer pH 9.5; b – for chromophore:protein complexes.

The absorption and emission spectra of the obtained complex lay between the spectra of complexes of FAST protein with fluorogens **N871b** and **HMBR** (Figure 5, Table 1). The fluorogen **HBR-DOM2** also efficiently binds to the original FAST protein with a more than an order of magnitude lower constant and a similar orange fluorescence color. The changes in the spectra occurring upon the **HBR-DOM2** binding to FAST and nanoFAST also reveal the deprotonation of its phenolic moiety. In a protein-free form, this fluorogen is already partially deprotonated at neutral pH; however, the low quantum yield of fluorescence of a free form allows avoiding an off-target signal.

**Figure 5.**
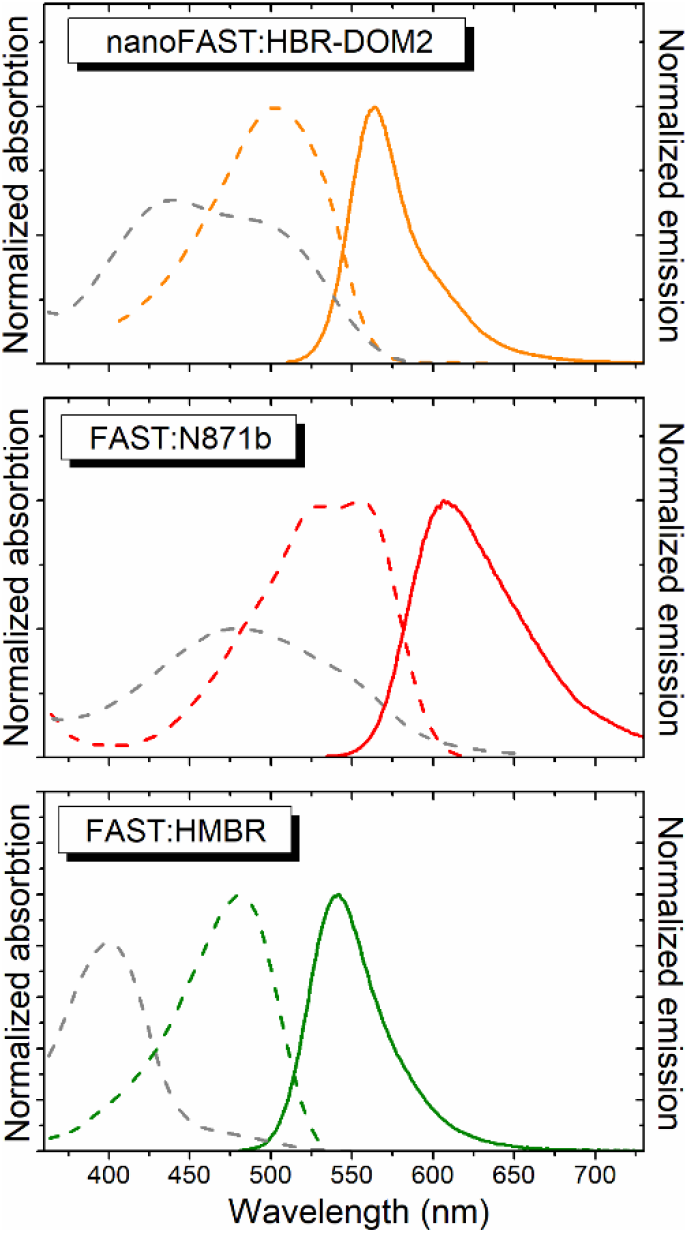
The absorption (dashed lines) and emission (solid line) spectra of free fluorogens (grey) and protein:fluorogen complexes (colored). The ratio of the absorption intensity of the complex and free fluorogene corresponds to the real change in the extinction coefficient and shape of the spectra in PBS.

We demonstrated the efficiency and utility of the proposed pair on a series of cells transiently transfected with various nanoFAST fusions (Figure 6A-D, SI Part 8).

**Figure 6.**
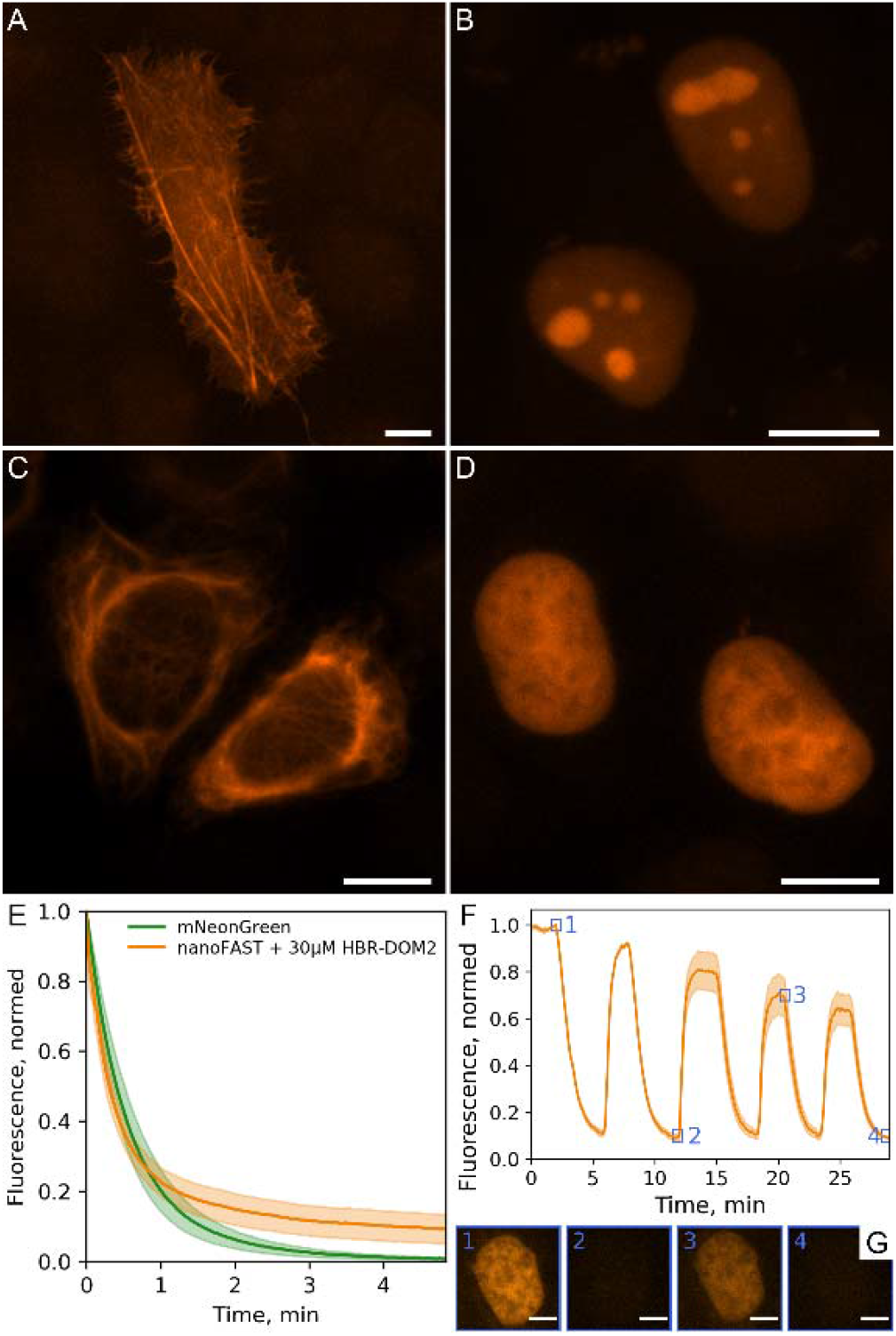
Performance of fluorogenic complex of nanoFAST with **HBR-DOM2** in fluorescence microscopy. Fluorescence imaging of live U2OS cells transiently transfected with (A) lifeact-nanoFAST and live HeLa cells transiently transfected (B) 3xNLS-nanoFAST, (C) vimentin-nanoFAST and (D) H2B-nanoFAST constructs in the presence of 5μM **HBR-DOM2**; Scale bars are 10 μm. (E) Photobleaching of nanoFAST in the presence of 30 μM **HBR-DOM2** in comparison with mNeonGreen under confocal microscopy conditions. (F) Sequential staining and washout of live HeLa cells transiently transfected with H2B-nanoFAST construct; 5 μM concentration of **HBR-DOM2** was used for staining; individual frames of single labeled nuclei from timings indicated by numbered blue squares presented on panel (G); Scale bars are 5 μm.

The fluorescent signal of the resulting complex turned out to be sufficiently photostable. We compared it with the mNeonGreen fluorescent protein, [24] which has a similar absorption spectrum and showed that their photobleaching curves behave in a similar way (Figure 6E). We have also confirmed that the binding of our fluorogen with nanoFAST protein is non-covalent and, if necessary, it can be easily washed out from cell media (Figure 6F, SI Video).

## Conclusion

In this paper, we report the first structures of FAST fluorogen-activating protein in the apo state and in complex with the ligand. Our data reveal that the structural organization of FAST is highly similar to that of parent photoactive yellow protein, with the FAST apo-state corresponding to the light-induced form of PYP and ligand-bound state corresponding to the ground conformation of the protein. The main feature of protein behavior is a significant rearrangement and organization of its N-terminus upon binding. Based on this, we assumed that removing this part of the protein can be carried out without the loss of fluorogen-activating properties. We shortened the FAST protein by 26 amino acid residues, found a suitable ligand, and thus created the shortest genetically encoded fluorogen-activating tag – nanoFAST - containing only 98 amino acids and suitable for fluorescent microscopy.

## Supporting information

Supporting material All

## Acknowledgements

Crystallization, X-ray data collection and treatment were supported by RFBR (№ 20-34-70034). The other work were supported by Russian Science Foundation, grant No. 18-73-10105. We sincerely thank Dr. Anastasia Mamontova and Dr. Alexey Bogdanov (IBCH RAS, Biophotonics department) for their kind help with the large-scale protein isolation, Egor Marin for the assistance with X-Ray data acquisition, Kirill Kovalev for the help with crystal harvesting and data collection. We acknowledge the ESRF Structural Biology Group (Grenoble, France), the EMBL unit at DESY (Hamburg, Germany) and SLS PSI (Villigen, Switzerland).

## Entry for the Table of Contents

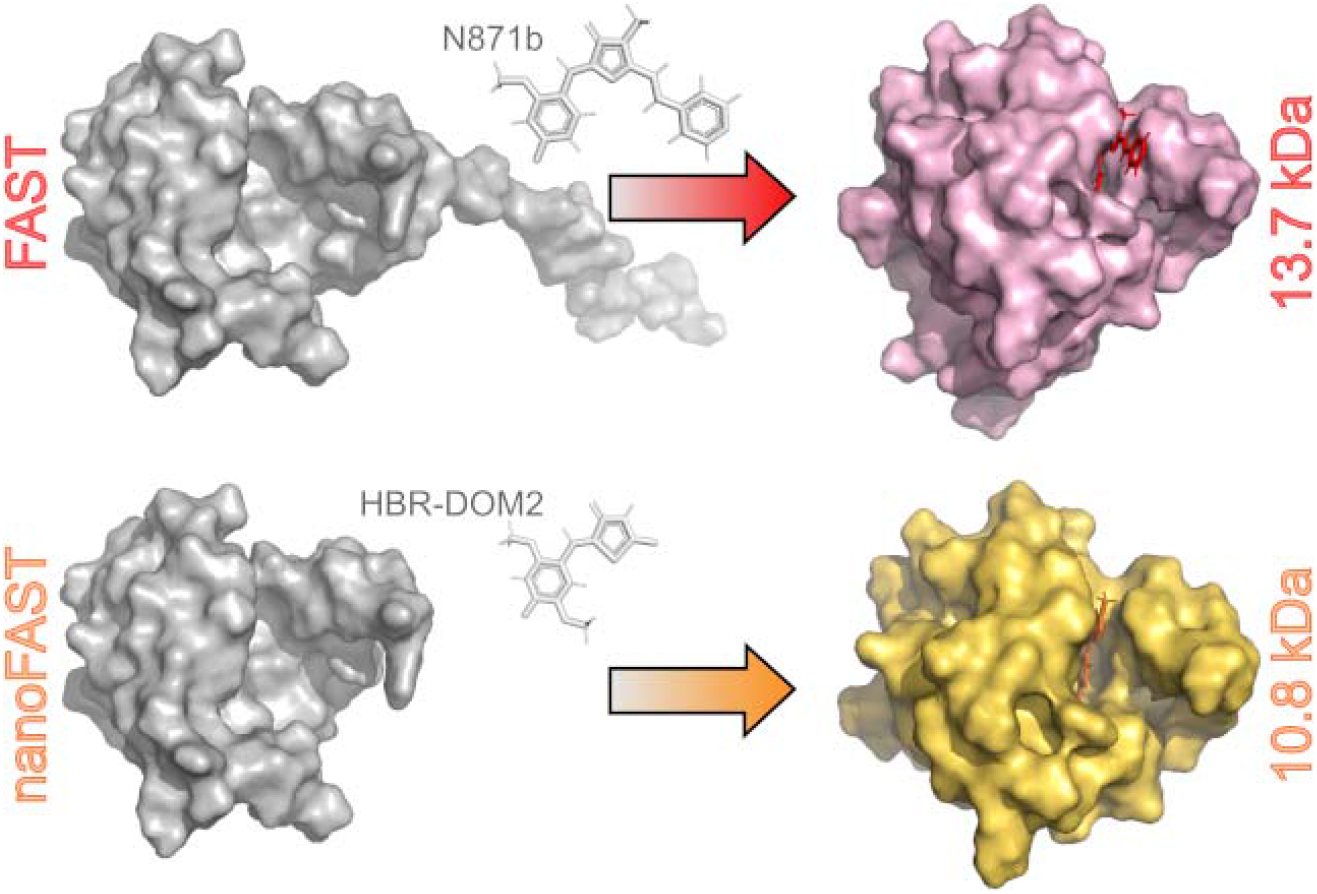
Novel tiny genetically encoded fluorescent tag nanoFAST for live cell microscopy is presented. The tag is based on the well known fluorogen-activating protein FAST. The structure of this protein was studied by NMR and its truncated variant was proposed. NanoFAST is just 98 amino acids long, the shortest genetically encoded tag among all known for today.

