## Supporting material All for "NanoFAST: Structure-based design of a small fluorogen-activating protein with only 98 amino acids"

### Contents

**1. General** **S2**

**2. Genes expression and protein purification** **S4**

**3. Crystallization, X-ray data collection and structure determination** **S6**

**4. NMR spectroscopy** **S13**

**5. Screening *in vitro*** **S21**

**6. Fluorescence quantum yields determination** **S26**

**7. Determination of affinity constants** **S27**

**8. Fluorescence microscopy, photobleaching, and sequential imaging** **S28**

**9. Synthetic procedures** **S29**

**10. $^{1}$H and $^{13}$C NMR spectra** **S30**

### 1. General

***Chromophores stock solutions***

All solid chromophores were dissolved in DMSO (Sigma Aldrich, “for molecular biology” grade, #cat D8418) in 10 mM concentration, and stored in dark place at -20°C for no more than 3 months.

***Fluorescence and absorption spectra***

UV-VIS spectra were recorded on a Varian Cary 100 spectrophotometer. Fluorescence excitation and emission spectra were recorded on Agilent Cary Eclipse fluorescence spectrophotometer.

***Molecular cloning***

Constructs for mammalian expression encoding fuses of nanoFAST with vimentin, histone H2B, lifeact, 3xNLS, and plasmid encoding fuse of mNeonGreen with histone H2B were assembled using Golden Gate cloning following MoClo standard.[[1]] Each transcriptional unit for mammalian expression consisted of the CMV promoter, coding sequence for the fusion protein, and the SV40 terminator. Cloning reactions were performed in the T4 ligase buffer (SibEnzyme, Russia) supplied with 10 U of T4 ligase, 20 U of either BsaI or BpiI (ThermoFisher, USA), and approximately 100 ng of DNA of each DNA fragments. Golden Gate reactions were performed with the following cycling conditions: 25 cycles between 37°C and 16°C (90 sec at 37°C, 180 sec at 16°C).

***FAST amino acid sequence***

(M)EHVAFGSEDIENTLAKMDDGQLDGLAFGAIQLDGDGNILQYNAAEGDITGRD PKQVIGKNFFKDVAPGTDSPEFYGKFKEGVASGNLNTMFEWMIPTSRGPTKVKVHMKK ALSGDSYWVFVKRV

***nanoFAST amino acid sequence***

(M)FGAIQLDGDGNILQYNAAEGDITGRDPKQVIGKNFFKDVAPGTDSPEFYGKF KEGVASGNLNTMFEWMIPTSRGPTKVKVHMKKALSGDSYWVFVKRV

[1] C. Lu, J. Browse, J. Wallis. *cDNA Libraries. Methods in Molecular Biology (Methods and Protocols),* **2011**. *729*. Humana Press. (C. Engler, S. Marillonnet Generation of Families of Construct Variants Using Golden Gate Shuffling.)

##### *Cell culture and transient transfection*

HeLa cells were grown in Dulbecco’s modification of Eagle’s medium (DMEM) (PanEco) supplied with 50 U/ml penicillin and 50 µg/ml streptomycin (PanEco), 2 mM L-glutamine (PanEco) and 10% fetal bovine serum (HyClone, Thermo Scientific) at 37 °C and 5% $CO_2$. For transient transfection, FuGENE 6 reagent (Promega) was used. Immediately before imaging DMEM was replaced with Hanks Buffer (PanEco) supplemented with 20 mM HEPES (Sigma).

##### *Synthesis*

Commercially available reagents were used without additional purification. Merck Kieselgel 60 was used for column chromatography. Thin layer chromatography (TLC) was performed on silica gel 60 $F_{254}$ glass-backed plates (MERCK). Visualization was effected by UV light (254 or 312nm) and staining with $KMnO_4$.

NMR spectra were recorded on a 700 MHz Bruker Avance III NMR at 303 K, 800 MHz Bruker Avance III NMR at 333 K and Bruker Fourier 300. Chemical shifts are reported relative to residue peaks of $CDCl_3$ (7.27 ppm for $^1H$ and 77.0 ppm for $^{13}C$) or DMSO-$d_6$ (2.51 ppm for $^1H$ and 39.5 ppm for $^{13}C$). Melting points were measured on a SMP 30 apparatus. High-resolution mass spectra (HRMS) spectra were recorded on a Bruker micrOTOF II instrument using electrospray ionization (ESI). The measurements were done in a positive ion mode (interface capillary voltage – 4500 V) or in a negative ion mode (3200 V); mass range from m/z 50 to m/z 3000; external or internal calibration was done with ESI Tuning Mix, Agilent. A syringe injection was used for solutions in acetonitrile, methanol, or water (flow rate 3 mL/min). Nitrogen was applied as a dry gas; interface temperature was set at 180 °C.

#### 2. Genes expression and proteins purification

Production protocols were the same for both variants (FAST and nanoFAST). Both coding sequences of FAST and nanoFAST were cloned into pet24b backbone with C-terminal his-tag. Proteins were expressed in BL21(DE3) *E. coli* strain in M9 medium. To obtain the $^{15}N$-labeled or $^{15}N$-$^{13}C$-labeled FAST, we used $^{15}NH_4Cl$ or $^{15}NH_4Cl$ and [U-$^{13}C$]-glucose, respectively. The cells were grown at 37 °C for several hours in a shaking incubator (New Brunswick Innova 44R) at 250 rpm. Protein expression was induced at OD600 ~ 0.6 by isopropyl β-d-1-thiogalactopyranoside (IPTG) at final concentration of 0.25 mM. Expression was carried out for 4-5 hours and cells were harvested by centrifugation at 7000×g for 10 min at 4 °C and stored at -20 °C.

Proteins were purified using metal affinity and size-exclusion chromatography. The cells from 600 ml of M9 medium were resuspended in 30 ml of lysis buffer (20 mM Tris, pH 8.0, 500 mM NaCl, 20 mM Imidazole) with 200 µM PMSF and disrupted on ice by 15 circles of ultrasonication (Bandelin Sonoplus, GM 2200, titanium tapered tip KE76) in pulse mode with an active interval of 15 sec at 60% of power followed by cooling for 3 min. The lysate was clarified by centrifugation at 14000×g for 60 min at 4 °C. The supernatant was filtered using a Millipore filter unit with 0.22 µm pore size and loaded to the column equipped with circa 5 ml $Ni^{2+}$ Sepharose HP resin (GE) pre-equilibrated with a lysis buffer. The column was washed with 5 column volumes (CV) of IMAC-buffer (20 mM Tris, pH 8.0, 200 mM NaCl) with 20 mM Imidazole followed by 5 CV of IMAC-buffer with 60 mM Imidazole. Target protein was eluted by IMAC-buffer with 500 mM Imidazole and dialyzed against the 1xPBS buffer with 1 мM EDTA overnight at 4 °C. Protein was concentrated up to 15 mg/ml by ultrafiltration (10 kDa MWCO, Amicon Ultra), clarified at 25000×g for 1 h and loaded to a Superdex 75 Tricorn 10/300 (GE) gel filtration column equilibrated in 1xPBS buffer. After SDS-PAGE analysis, fractions containing the pure target protein were combined. For NMR applications the target protein was dialyzed against the NMR-buffer (20 mM NaPi, pH 7.0, 20 mM NaCl) overnight at 4 °C and concentrated up to 15 mg/ml.

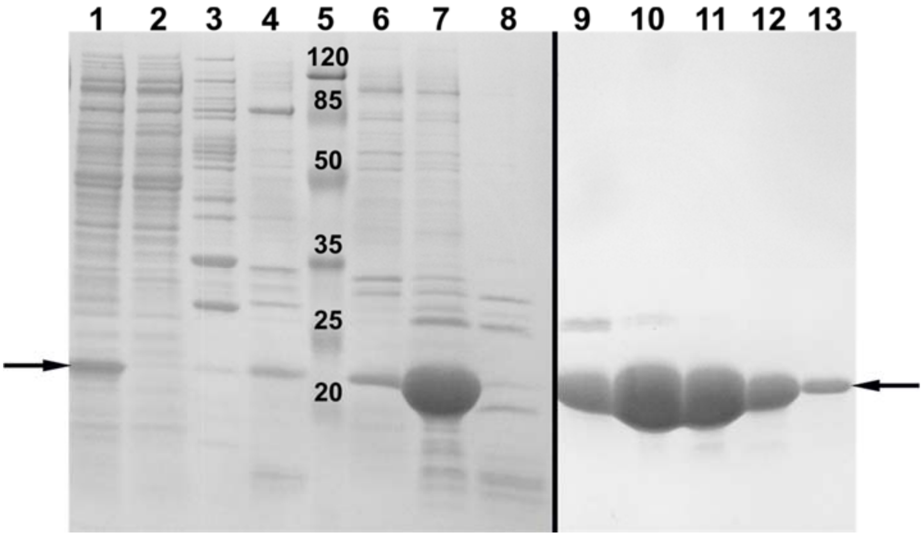


**Figure S2.1.** Purification summary for the FAST. Lines 1-8 correspond to immobilized metal affinity chromatography and lines 9-13 correspond to size-exclusion chromatography. FAST is indicated by arrows. 1 - clarified and filtered lysate, 2 - unbound proteins after washing with 20 mM Imidazole, 3 and 4 - unbound proteins after washing with 40 mM and 80 mM Imidazole, respectively, 4 - molecular weight markers, 6-8 elution with 500 mM Imidazole, 9-13 fractions of FAST after size-exclusion chromatography.

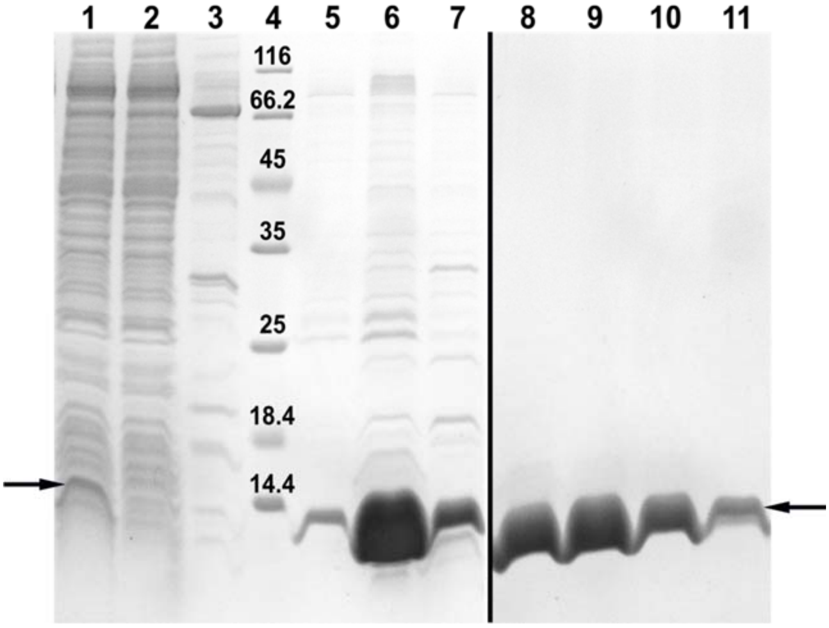


**Figure S2.2.** Purification summary for the nanoFAST. Lines 1-7 correspond to immobilized metal affinity chromatography and lines 8-11 correspond to size-exclusion chromatography. nanoFAST is indicated by arrows. 1 - clarified and filtered lysate, 2 - unbound proteins after washing with 20 mM Imidazole, 3 - unbound proteins after washing with 60 mM Imidazole, 4 - molecular weight markers, 5-7 elution with 500 mM Imidazole, 8-11 fractions of purified nanoFAST after size-exclusion chromatography.

#### 3. Crystallization, X-ray data collection and structure determination

***Crystallization of FAST***

Initially, we crystallized FAST with **N871b** and without the ligand. Later on we co-crystalized the protein with the ligand for nanoFAST (**HBR-DOM2**). All crystallizations gave crystals of the same morphology, space group and protein structure, albeit of different crystallographic resolution. In all crystallizations, purified protein was concentrated to 13-22 mg/mL. Ligands were dissolved in DMSO and added to the protein solution at final concentrations of 150 nM and 1.5 mM in the case of **HBR-DOM2** and **N871b**, correspondingly. Protein crystals were obtained by sitting drop vapor diffusion approach in 96-well crystallization plates (SPT Labtech, UK). Crystallization trials were set up by the NT8 robotic system (Formulatrix, USA). The drops contained 200 nL concentrated protein solution and 100 nL reservoir solution. Crystallization Screens JCSG-plus and Morpheus (Molecular Dimensions, UK) were used for screening of crystallization conditions. Crystals were grown at 20 °C. The crystallization buffer composition was the following: 0.2 M Magnesium chloride hexahydrate, 0.1 M Bis-Tris, pH 5.5, 25 % w/v PEG 3350 for ligand-free; in 0.2 M Sodium chloride, 0.1 M Bis-Tris, pH 5.5, 25 % w/v PEG 3350 for N871b; 0.2 M magnesium formate, 20 % w/v PEG 3350 for HBR-DOM2 (Fig. 1c). Upon these conditions, crystals were grown to the final size of ~100 µm (Fig. S3.1a), ~50 µm (Fig. S3.1b) and ~300 µm (Fig. S3.1c), correspondingly.

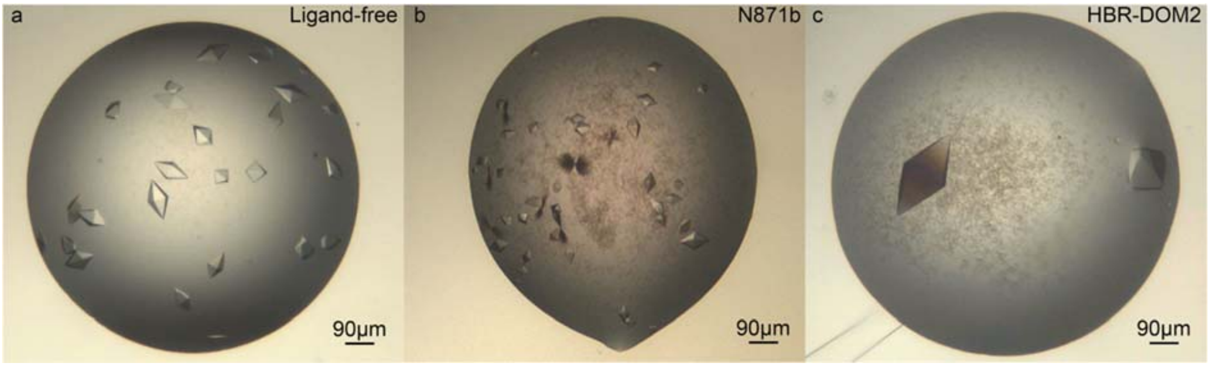


**Figure S3.1.** Crystals of FAST used for data collection. **a**, crystals grown in the absence of ligand. **b**, crystals grown in the presence of **N871b** **c**, crystals grown in the presence of **HBR-DOM2.**

##### *X-ray diffraction data collection*

X-ray diffraction data were collected at beamline P13 (DESY, Hamburg, Germany), X06SA (SLS, Paul Scherrer Institute, Villigen, Switzerland) and ID23-1 (ESRF, Grenoble, France) for Ligand-free, **N871b** and **HBR-DOM2**, correspondingly. Crystals were fished in MiTeGen micromounts and immediately flash-frozen in liquid nitrogen. All data were collected at 100 K. The data collection details were the following: 0.1°/0.1°/0.05° oscillation and 360°/200°/360° total range for ligand-free/**N871b**/**HBR-DOM2**, correspondingly. Total absorbed dose did not exceed 1.5 MGy for each dataset as assessed by *RADDOSE-3D*.[[2]] Diffraction data were integrated and scaled in the *XDS* program suite.[[3]] Data collection statistics can be found in Table S3.1.

The resolution of datasets was 2.5 Å, 3.1 Å and 1.5 Å, for Ligand-free, **N871b** and **HBR-DOM2**, correspondingly. The crystallographic models were built for all datasets and showed no difference. Model building for the best dataset (for the crystals grown in the presence of **HBR-DOM2**) is described below. Phase problem was solved by molecular replacement with the *MoRDa* pipeline[[4]] from the *CCP4* web service[[5]] with PDB ID 1ODV[[6]]) automatically selected as a starting model. The solution was subsequently rebuilt in the *ARP/wARP* web service[[7]]. Coordinate and B-factor refinement with 9 TLS groups was performed in *phenix.Refine*.[[8,9]] *Coot*[[10]] was utilized for manual model building between refinement cycles. Final resolution cutoff was determined by paired refinement.[[11]] Structure was validated in *phenix.MolProbity*.[[12]] Refinement statistics can be found in Table S3.2. The secondary structure has been assigned with DSSP.[[13]] All model figures were rendered in *PyMOL* (Schrödinger LLC, New York, NY, USA).

---

*Fo-Fo* q-weighted maps[14] were built between **HBR-DOM2/N871b** and ligand-free with the phases from the obtained model and showed no signs of ligands.

The diffraction images and processed data for the **HBR-DOM2** dataset were deposited to Integrated Resource for Reproducibility in Macromolecular Crystallography[15] (http://proteindiffraction.org/) under accession number 7AV6. The corresponding structure for FAST in the domain-swapped dimer form was deposited in the Protein Data Bank under accession number 7AV6.

[14] D. Bourgeois, *Acta Crystallogr. Sect. D Biol. Crystallogr.* **1999**, 55, 1733–1741.

[15] M. Grabowski, K. M. Langner, M. Cymborowski, P. J. Porebski, P. Sroka, H. Zheng, D. R. Cooper, M. D. Zimmerman, M. A. Elsliger, S. K. Burley, W. Minor, *Acta Crystallogr. Sect. D Struct. Biol.* **2016**, *72*, 1181–1193.

**Table S3.1**. Data collection statistics

| Dataset | Ligand-free | **N871b** | **HBR-DOM2** |
|---|---|---|---|
| Beamline | PETRA III P13 | SLS X06SA | ESRF ID23-1 |
| Detector | PILATUS 6M | EIGER 16M | PILATUS 6M-F |
| Wavelength (Å) | 0.976 | 1.000 | 0.977 |
| Space group | $P4_12_12$ | $P4_12_12$ | $P4_12_12$ |
| a, b, c (Å) | 44.76, 44.76, 107.56 | 44.65, 44.65, 107.28 | 45.92, 45.92, 105.04 |
| α, β, γ (°) | 90, 90, 90 | 90, 90, 90 | 90, 90, 90 |
| Resolution (Å)* | 41.33 - 2.50 (2.59 - 2.50) | 41.23 - 3.10 (3.21 - 3.10) | 42.09 - 1.50 (1.54 - 1.50) |
| No. of observations* | 101249 (8479) | 20432 (1808) | 471330 (30388) |
| No. of unique reflections* | 4159 (389) | 2156 (212) | 18549 (1274) |
| Rpim* | 0.013 (1.110) | 0.015 (0.433) | 0.007 (0.726) |
| I/σI* | 21.5 (0.4) | 20.9 (1.3) | 32.8 (0.7) |
| CC1/2* | 100 (77.0) | 100 (83.7) | 100 (32.5) |
| Completeness (%)* | 99.9 (100) | 96.1 (96.4) | 98.5 (96.6) |
| Redundancy* | 24.3 (21.8) | 9.5 (8.5) | 25.4 (23.9) |
| Wilson B-factor ($Å^2$) | 84.6 | 118.8 | 34.2 |

*Values in parentheses are for highest-resolution shell.

**Table S3.2**. Refinement statistics

| | |
|---|---|
| Resolution (Å)* | 42.08 - 1.50 (1.55 - 1.50) |
| No. of reflections (work/free)* | 18507/925 (1766/87) |
| Rwork/Rfree* | 0.215/0.239 (0.367/0.392) |
| No. of atoms | |
| Protein | 764 |
| Solvent | 29 |
| B-factors ($Å^2$) | |
| Protein | 50.4 |
| Solvent | 54.7 |
| R.m.s.d | |
| Bond lengths (Å) | 0.005 |
| Bond angles (°) | 0.73 |
| Ramachandran statistics | |
| Residues in favored regions (%) | 97.89 |
| Residues in allowed regions (%) | 100 |

*Values in parentheses are for highest-resolution shell.

***Crystallographic structure of FAST***

FAST crystallized in a form of a domain-swapped dimer that contains no chromophore molecule (Fig. S3.2a). The three β-strands: β1 (A30-L33), β2 (I39-N43), and a newly formed β2′ (T50-R52) - are swapped with the corresponding elements from the symmetrical molecule, forming together one twelve-stranded β sheet protein.

Compared to the NMR ligand-bound structure, significant differences in the secondary and tertiary structures are observed (Fig. S3.2b). First, helix H3 (A44-G51) and the adjacent R52 transform into the β2′ strand and the β2-β2′ turn (A44-I49). Second, the N-term, which includes helices H1 (D10-K17) and H2 (D20-G25), becomes disordered in the domain-swapped dimer. The C-term linker and His-tag (L127-H138) were not resolved in our crystal structure, correlating with their high flexibility in the NMR complex.

Despite these tremendous rearrangements, the dimer partially conserves binding site, with the key β3-β5 strands (N89-R124) and H4H5-loop (I58-S72) not deviating significantly compared to the NMR ligand-bound model (with Cα-Cα RMSD not exceeding 1.8 Å and 1.5 Å, respectively). In particular, W94 from β3, which forms H-bond with the carbonyl group of the imidazolone ring in the FAST:**N871b** complex, preserves its position in the swapped dimer. In contrast, E46 at helix H3, which H-bonded the phenol ring of the fluorogen, is located in the swapped part of the crystallographic model ~7 Å away from its position in FAST:**N871b** complex. The loss of the stabilizing H-bond with E46 presumably ties with the chromophore uncoupling, destabilization of protein tertiary structure and finally with the domain swapping.

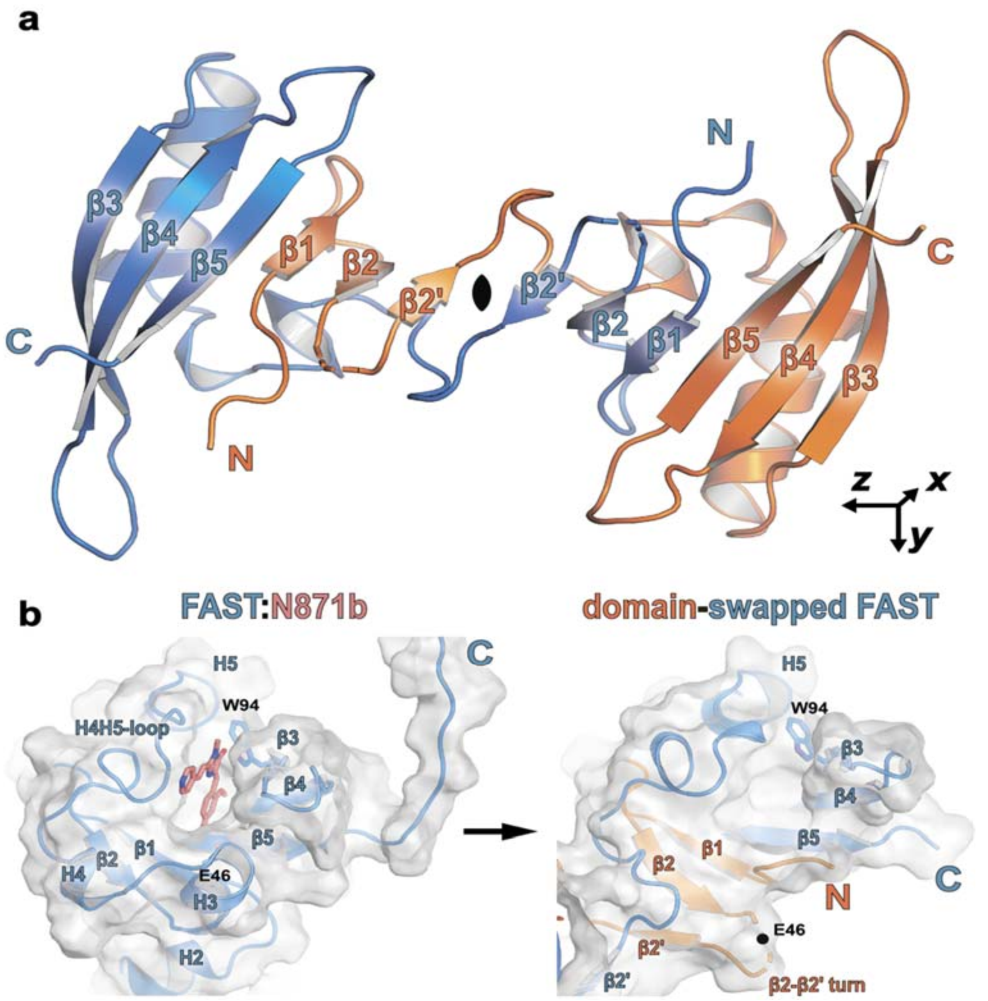


**Figure S3.2**. **a**, Crystal structure of FAST in the domain-swapped dimer form. The asymmetric unit consists of one protein molecule (colored blue), while the second (colored orange) is obtained by applying the twofold rotation about the x-axis. **b**, Structural rearrangements in the chromophore binding pocket. On the left - FAST:**N871b** NMR complex (protein colored blue, ligand colored pink). On the right - domain-swapped dimer with the swapped N-term displayed in orange. Approximate position of E46 Cα from the disordered part of the β2-β2’ turn is shown by the dot in the latter case.

### 4. NMR spectroscopy

All NMR spectra were acquired at pH 7.0, 20 mM NaPi, 20 mM NaCl. Concentration of protein was 0.85 mM, concentration of **N871b** was taken equal 0.89 mM. FAST apo state was investigated at 45 °C using the Bruker Avance III 600 MHz spectrometer, FAST/**N871b** complex was studied at 25 °C on the Bruker Avance III 800 MHz spectrometer, both devices were equipped with helium cryogenic triple-resonance probes. NMR chemical shift assignment was made manually using the full set of triple resonance spectra (HNCO, HNCA, HNCACAB, HN(CA)CO, HN(CO)CA and CBCA(CO)NH), HC(C)H-TCOSY, (HB)CB(CGCC)Har [16] and TROSY-based aromatic (H)CCH-COSY.[17] All triple resonance experiments were run in BEST-TROSY versions.[18] Non-uniform sampling in indirect dimensions was applied and spectra were processed in qMDD software[19] using the IST algorithm and virtual echo. $^{3}J_{CCo}$ and $^{3}J_{NCo}$ couplings were measured from the spin-echo difference constant-time HSQC spectra.[20] $^{3}J_{NH\beta}$ couplings were derived from the cross-peak intensities in a J-quantitative 3D HNHB experiment.[21] Due to the limited lifetime of FAST apo samples (5 days), HNHB was not recorded. Spatial structure of the protein was calculated based on the distance restraints derived from the 3D 1H,15N-NOESY-HSQC and 1H,13C-NOESY-HSQC (two spectra with 13C offsets set at 48 and 120 ppm, recorded in $D_2O$) experiments acquired with 80 ms mixing time and dihedral restraints ($\varphi$, $\chi_1$) obtained from the analysis of J-couplings and chemical shifts in TALOS-N software.[22] Structure calculation was performed in the automated mode as implemented in CYANA 3.98 software.[23] Intermolecular distance restraints were determined directly using the 13C-filtered-NOESY-HSQC spectrum.[24] Ligand chemical shifts were assigned by the analysis of superimposed 2D-NOESY 120 ms spectrum, recorded with and without the $^{13}C/^{15}N$ decoupling. In addition, the HMBC-derived assignment of **N871b** in DMSO solution was utilized (Figure S4.4). NMR structures, peak lists and chemical shifts were deposited to PDB data based under the accession codes 7AVB and 7AVA for the FAST apo state and **N871b** complex, respectively. MOLMOL[25] and PyMOL software (Schrödinger LLC) was used for 3D visualization.

---

16 F. Löhr, R. Hänsel, V. V. Rogov, and V. Dötsch, *J. Biomol. NMR*, **2007**, *37*, 3, 205–224.

17 K. Pervushin, R. Riek, G. Wider, and K. Wüthrich, *J. Am. Chem. Soc.*, **1998**, *120*, 25, 6394–6400.

18 A. Favier and B. Brutscher, *J. Biomol. NMR*, **2011**, *49*, 1, 9–15.

19 M. Mayzel, K. Kazimierczuk, and V. Y. Orekhov, *Chem. Commun. (Camb.)*, **2014**, *50*, 64, 8947–8950.

20 (a) S. Grzesiek, G. W. Vuister, and A. Bax, *Journal of biomolecular NMR*, **1993**, *3*, 4, 487–493. (b) G. W. Vuister, A. C. Wang, and A. Bax, *Journal of the American Chemical Society*, **1993**, *115*, 12, 5334–5335.

21 P. Düx, B. Whitehead, R. Boelens, R. Kaptein, and G. W. Vuister, *J. Biomol. NMR*, **1997**, *10*, 3, 301–306.

22 Y. Shen and A. Bax, *Methods Mol. Biol.*, **2015**, *1260*, 17–32.

23 P. Güntert and L. Buchner, "*J Biomol NMR*, **2015**, *62*, 4, 453–471.

24 C. Zwahlen, P. Legault, S. J. F. Vincent, J. Greenblatt, R. Konrat, and L. E. Kay, *J. Am. Chem. Soc.*, **1997**, *119*, 29, 6711–6721.

25 R. Koradi, M. Billeter, and K. Wüthrich, *J Mol Graph*, **1996**, *14*, 1, 51–55.

Hydrodynamic radii were measured using the $^1H$ diffusion NMR spectroscopy at 30 °C and PGSTE-WATERGATE pulse program,[26] region of methyl groups (1.0-0.5 ppm) was used for the analysis. The obtained diffusion coefficients were then converted to the hydrodynamic radius according to the Stokes-Einstein relationship, and to the molecular mass of the protein, according to the well-established correlation between the diffusion coefficients and weights of globular proteins.[27]

[26] G. Zheng, T. Stait-Gardner, P.G. A. Kumar, A. M. Torres, W. S. Price, *Journal of Magnetic Resonance*, **2008**, *191*, 1, 159-163.
[27] M.T. Tyn, T.W. Gusek. *Biotechnol Bioeng* **1990**. *35*. 327–338.

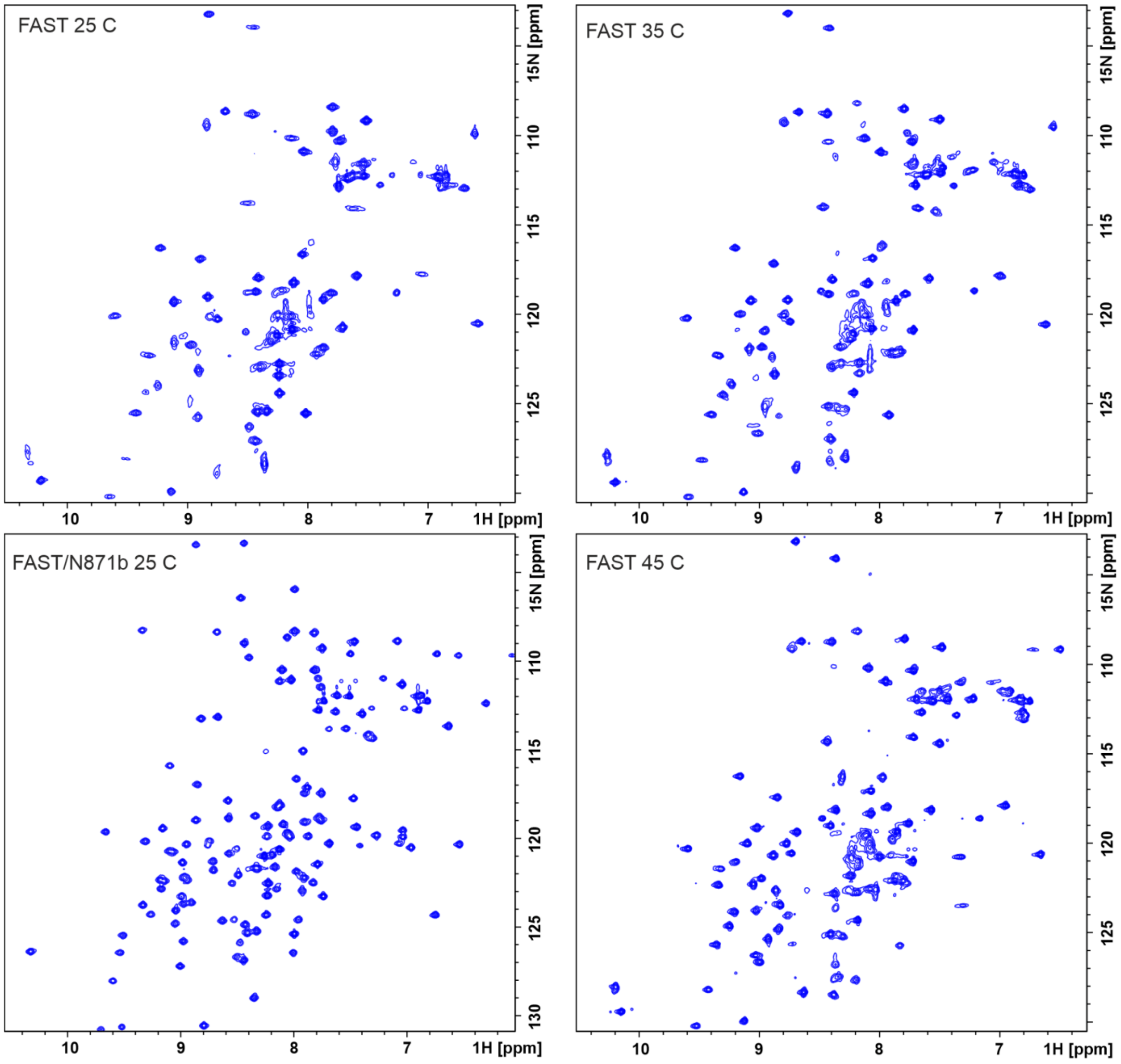


**Figure S4.1.** NMR spectra of FAST. Shown are the $^{1}H,^{15}N$-HSQC spectra of FAST apo at 25, 35 and 45 °C and spectrum of FAST:**N871b** complex at 25 °C. Spectra were recorded at pH 7.0, 800 MHz, concentration of protein was 0.2 mM, protein/ligand ratio equaled 0.95.

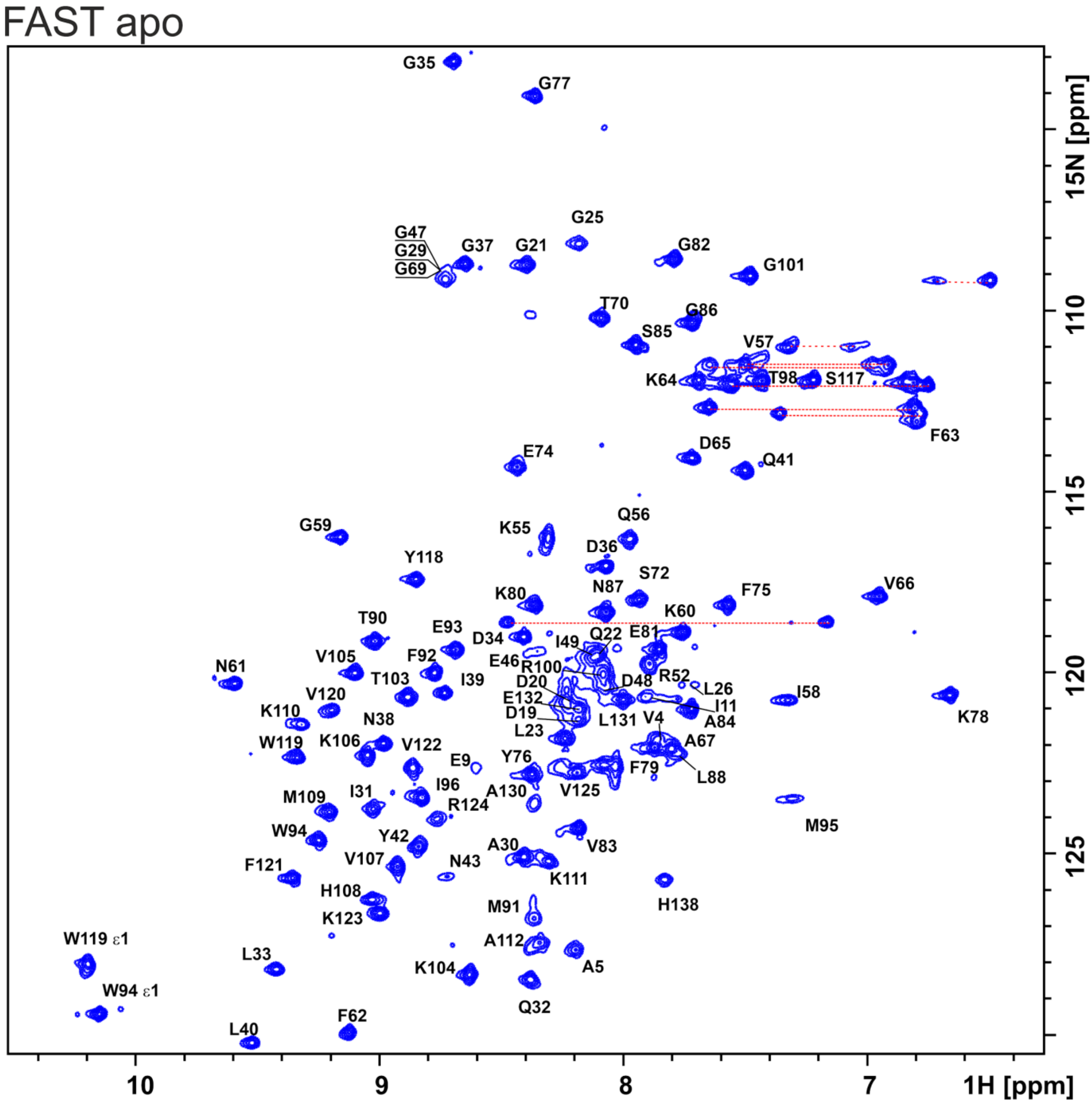


**Figure S4.2.** $^{1}$H,$^{15}$N-HSQC spectrum of FAST apo. Spectrum was recorded at 45 ºC, pH 7.0, 800 MHz. Concentration of the protein was 0.2 mM. Assignment of signals is indicated. $NH_2$ groups of Asn and Gln sidechains are shown by red dashed lines.

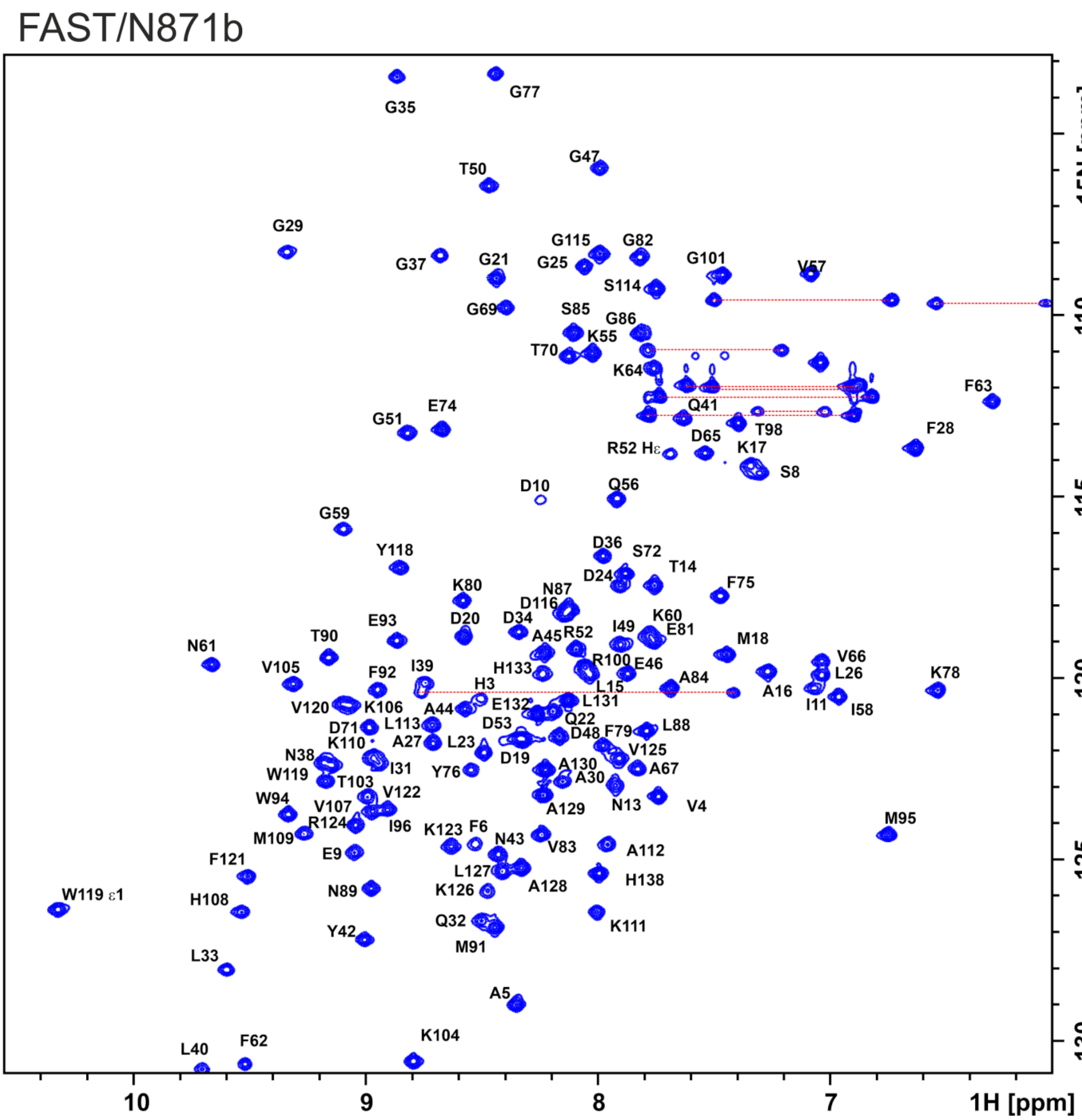


**Figure S4.3.** $^{1}H,^{15}N$-HSQC spectrum of FAST:**N871b**. Spectrum was recorded at 25 °C, pH 7.0, 800 MHz. Concentration of the protein was 0.2 mM, of N871b – 0.22 mM. Assignment of signals is indicated. $NH_2$ groups of Asn and Gln sidechains are shown by red dashed lines.

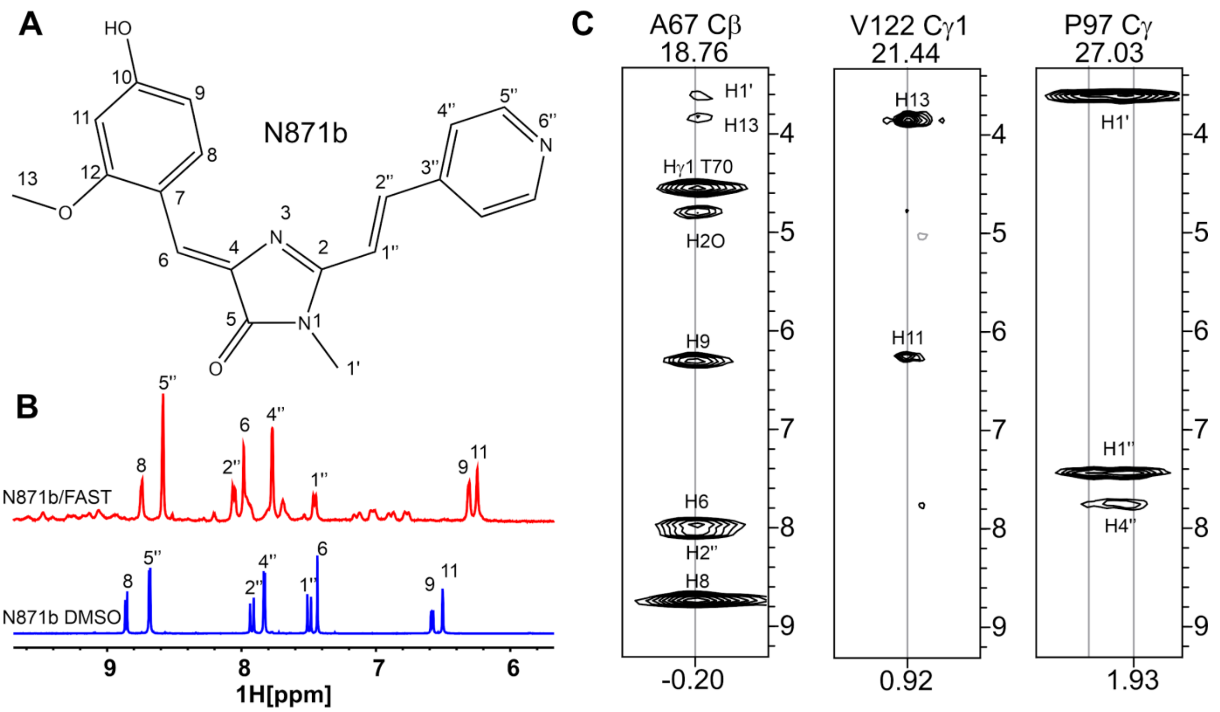


**Figure S4.4.** Ligand-protein interactions in FAST:**N871b** complex. A – the structure of **N871b** with the atom numbering indicated. **B** - fragment of $^{1}$H NMR spectra of **N871b** in DMSO (blue) is superimposed with the corresponding region of a $^{13}$C-filtered $^{1}$H spectrum of FAST:**N871b** in D2O (red). Smaller signals in the spectrum correspond to the amide groups protons that have not been completely exchanged to deuterium. Assignment of peaks is shown. **C** – representative 2D strips extracted from the 3D $^{13}$C,$^{15}$N-filtered $^{13}$C-edited NOESY-HSQC spectrum of FAST:**N871b** complex (0.85/0.89 mM, pH 7.0, 25 $^{o}$C, 120 ms mixing time). Assignment of cross-peaks is shown.

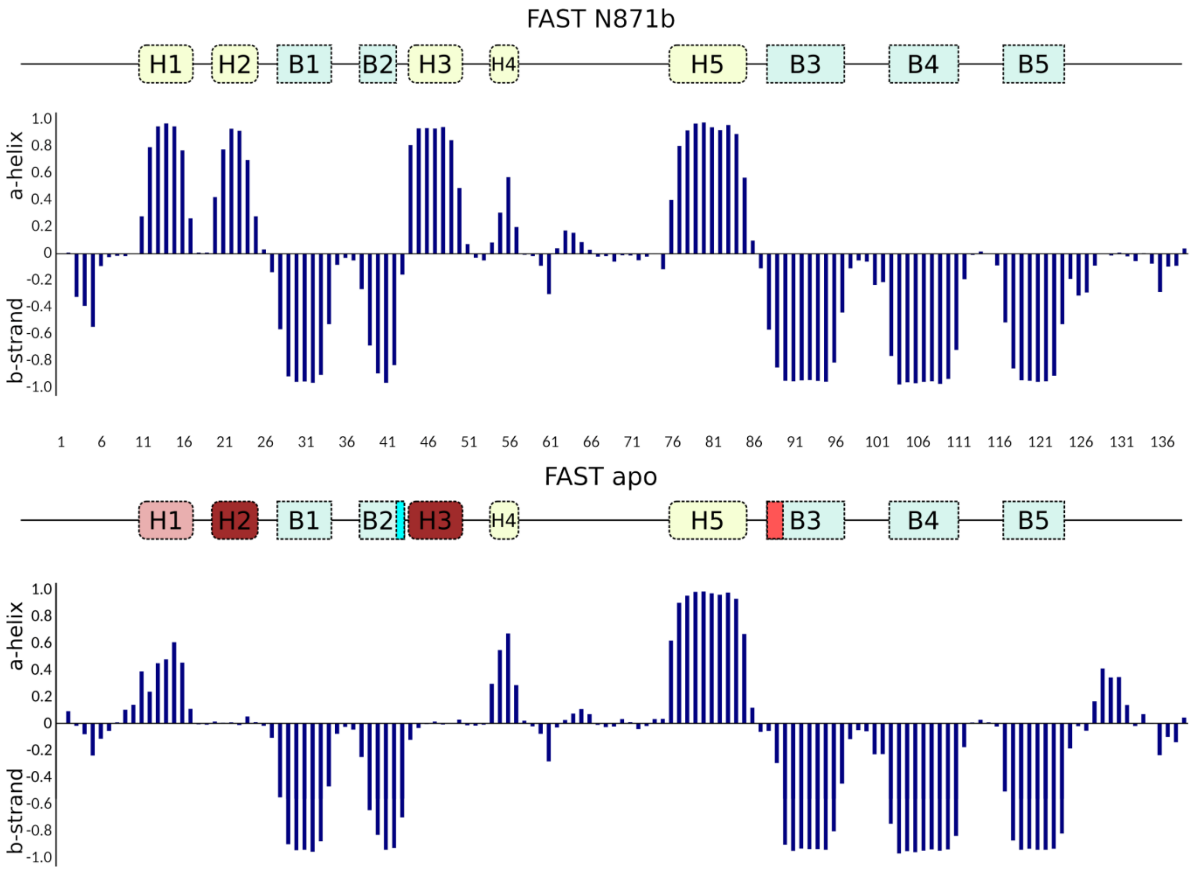


**Figure S4.5.** Secondary structure of FAST apo state and in complex with **N871b**. Shown is the secondary structure propensity predicted by the TALOS-N software [ref] based on the NMR chemical shift. Negative values correspond to the extended conformation, positive – to the α-helices. Secondary structure of FAST:**N871b** is schematically shown on top. Elements that are lost completely or partially in the apo state are colored by red. Helices H2 and H3 are lost completely, helix H1 is lost partially. Strand B3 becomes 2 residues shorter, while strand B2 becomes one residue longer.

**Table S4.1.** Input data and statistics of 20 best NMR structures calculated out of 100 random starting points for FAST apo and FAST:**N871b** complex.

| **Distance and Angle restraints** | | |
|---|---|---|
| | FAST apo | FAST/N871b |
| Total NOE contacts | 1162 | 2397 |
| intraresidual | 292 | 492 |
| sequential (\|i-j\|=1) | 317 | 587 |
| medium-range (1<\|i-j\|<4) | 171 | 518 |
| long-range (\|i-j\|>4) | 382 | 762 |
| intermolecular | 0 | 38 |
| H-bond restraints | 13 | 42 |
| TALOS-N angles (φ/ϕ) | 175 | 231 |
| Total J-couplings | 86 | 165 |
| $^3J_{NH\beta}$ | 0 | 59 |
| $^3J_{C\gamma C'}$ | 43 | 53 |
| $^3J_{C\gamma N}$ | 43 | 53 |
| Total restraints/per residue (1-125): | 1501/12.0 | 3045/24.3 |
| **Statistics of the obtained structures** | | |
| Structures calculated/selected | 100/20 | 100/20 |
| CYANA target function (Å$^2$) | 1.52±0.07 | 2.16±0.05 |
| *Violations of restraints* | | |
| Distance (>0.2 Å) | 0 | 0 |
| Dihedral angles (>5 °) | 2 | 4 |
| RMSD (Å), regular structure | **(29-43,54-67,75-124)** | **(11-57,76-124)** |
| Backbone | 0.84±0.22 | 0.22±0.05 |
| All heavy atoms | 1.44±0.17 | 0.79±0.08 |
| RMSD (Å) full PAS domain | **(1-125)** | **(1-125)** |
| Backbone | 12.6±4.8 | 0.31±0.07 |
| All heavy atoms | 12.9±4.7 | 0.90±0.07 |
| *Ramachandran analysis* | | |
| Residues in favored regions (%) | 81.9 | 77.6 |
| Residues in allowed regions (%) | 100 | 100 |

### 5. Screening *in vitro*

Optical properties of chromophores were investigated using 10 μM solutions in water (pH 5). Chromophore-protein binding was tested using solutions containing 10 μM and 1 μM of nanoFAST and chromophore respectively in PBS buffer (pH 7.4, #cat E404-200TABS, Amresco).

Fluorescence intensity enhancement was defined as the ratio of fluorescence intensity of chromophore+protein solution to the fluorescence intensity of free chromophore solution recorded on Tecan Infinite 200 Pro M Nano dual mode plate reader (1 μM chromophore and 10 μM protein solutions). Both solutions were excited at similar wavelengths listed in Table S5.1. Due to the concentrations used, there can be non-saturated conditions. Correction to the absorbance intensity was not made.

**Table S5.1.** Optical properties of fluorogenes and the results of interaction with nanoFAST

| Cmpd | Structure | Abs[a] | Em[a] | Enhancement | | | | | |
|---|---|---|---|---|---|---|---|---|---|
| | | | | 380 | 430 | 480 | 530 | 580 | 630 |
| **HMBR** | 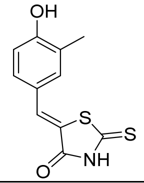 | 400 | 493 | 1.3 | 2.0 | 3.9 | 1.2 | 0.9 | 1.1 |
| **HBR-DOM** | 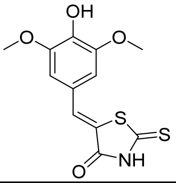 | 406 | 530 | 1.1 | 1.2 | 2.1 | 4.1 | 1.1 | 1.0 |
| **HBR-DOM2** | 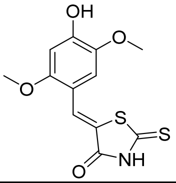 | ~420 | ~500 | 2.6 | 18.0 | 100.0 | 252.9 | 1.4 | 1.0 |
| **N 871b** | 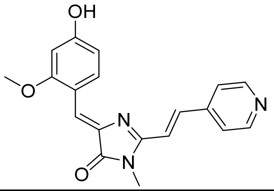 | 455 | 570 | 1.1 | 1.1 | 1.6 | 2.6 | 2.5 | 1.3 |
| **SAI 112** | 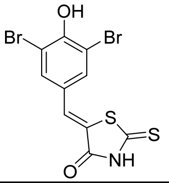 | 384 [b] | 445 [b] | 1.3 | 2.4 | 5.7 | 1.7 | 1.1 | 1.1 |
| **SAI 117** | 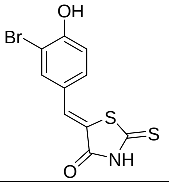 | 387 [b] | 445 [b] | 1.4 | 2.5 | 4.3 | 1.3 | 1.0 | 0.6 |

| | | | | | | | | | |
|---|---|---|---|---|---|---|---|---|---|
| **SAI 122** | 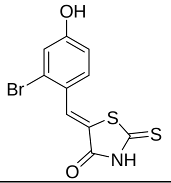 | 386 [b] | 446 [b] | 1.5 | 5.2 | 5.8 | 1.2 | 1.1 | 0.8 |
| **SAI 127** | 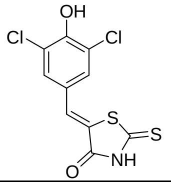 | 383 [b] | 448 [b] | 1.3 | 2.8 | 6.0 | 1.5 | 1.2 | 0.8 |
| **SAI 125** | 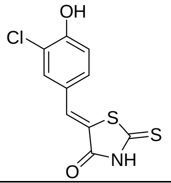 | 386 [b] | 445 [b] | 1.4 | 3.2 | 5.4 | 1.2 | 1.0 | 0.5 |
| **SAI 199** | 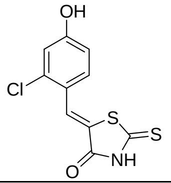 | 387 [b] | 448 [b] | 1.9 | 8.6 | 9.8 | 1.1 | 0.9 | 1.3 |
| **SAI 118** | 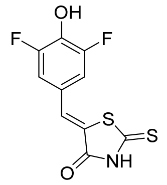 | 382 [b] | 445 [b] | 1.2 | 1.6 | 2.0 | 1.3 | 1.0 | 1.2 |
| **SAI 121** | 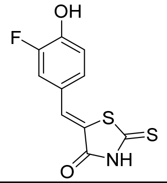 | 386 [b] | 443 [b] | 1.2 | 1.8 | 2.3 | 1.2 | 1.2 | 0.8 |
| **SAI 120** | 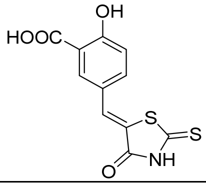 | 390 [b] | 447 [b] | 1.1 | 1.1 | 1.2 | 1.2 | 1.1 | 1.8 |
| **HBR-2,5-DM** | 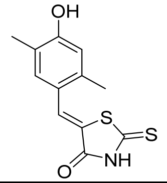 | 397 [b] | ~475 [b] | 1.6 | 7.6 | 42.9 | 8.2 | 1.0 | 0.8 |
| **SAI 366** | 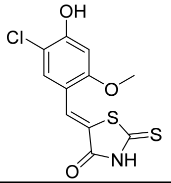 | 403 [b] | ~480 [b] | 2.2 | 17.2 | 31.2 | 3.1 | 0.9 | 1.4 |
| **SAI 365** | 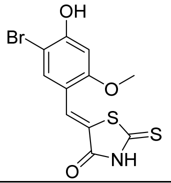 | 403 [b] | ~480 [b] | 1.7 | 12.7 | 37.4 | 3.3 | 1.1 | 1.3 |

| | | | | | | | | | |
|---|---|---|---|---|---|---|---|---|---|
| **SAI 363** | 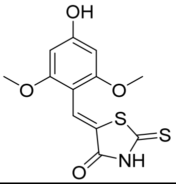 | 410[b] | ~470[b] | 1.4 | 6.8 | 16.2 | 1.3 | 1.0 | 1.2 |
| **SAI 362** | 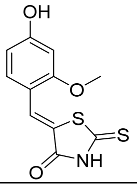 | 460[b] | ~475[b] | 2.2 | 18.7 | 62.8 | 3.0 | 1.1 | 1.0 |
| **SAI 367** | 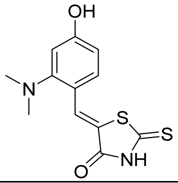 | 402[b] | ~570[b] | 1.2 | 1.6 | 1.5 | 1.2 | 0.9 | 1.0 |
| **ZS 319.2** | 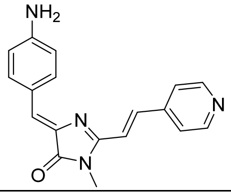 | 413 | 615 | 1.1 | 1.0 | 1.0 | 1.0 | 1.1 | 1.0 |
| **ZS 309** | 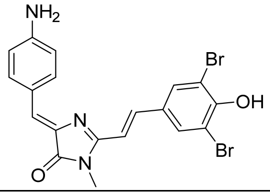 | 453 | 583 | 1.2 | 1.2 | 1.5 | 2.2 | 1.3 | 1.1 |
| **N 976** | 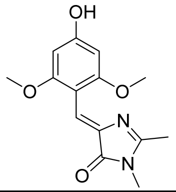 | 393 | 485 | 1.1 | 1.1 | 1.0 | 1.0 | 1.1 | 0.8 |
| **ZS 316** | 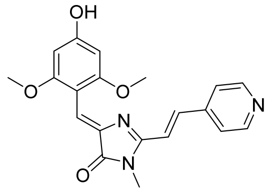 | 446 | 550 | 1.1 | 1.1 | 1.1 | 1.0 | 0.9 | 1.1 |
| **N 973** | 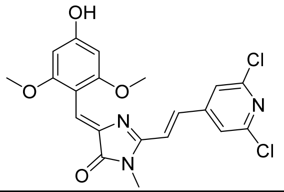 | 453 | 576 | 1.1 | 1.1 | 1.1 | 1.1 | 0.9 | 0.9 |
| **N 971** | 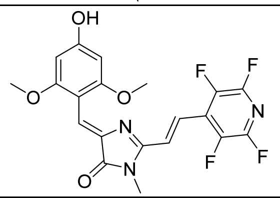 | 453 | 567 | 1.1 | 1.1 | 1.1 | 1.1 | 1.1 | 1.2 |
| **N 979** | 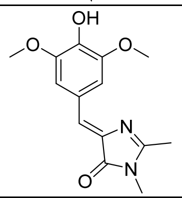 | 383 | 485 | 1.1 | 1.0 | 1.0 | 1.0 | 1.1 | 0.9 |

| | | | | | | | | | |
|---|---|---|---|---|---|---|---|---|---|
| **N 960a** | 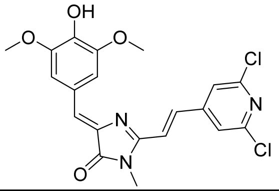 | 437 | -[c] | 1.1 | 1.1 | 1.0 | 1.1 | 1.2 | 1.2 |
| **N 980** | 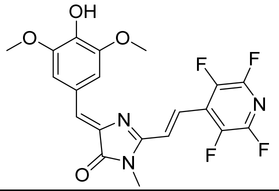 | 468 | -[c] | 1.1 | 1.0 | 1.0 | 1.1 | 1.1 | 1.0 |
| **ZS 325** | 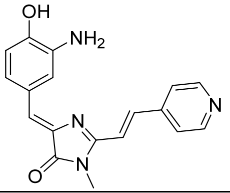 | 393 | -[c] | 1.1 | 1.1 | 1.0 | 1.0 | 1.4 | 1.0 |
| **N 1036** | 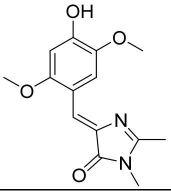 | 410 | 520 | 1.1 | 1.1 | 1.1 | 1.1 | 1.0 | 1.9 |
| **N 1048** | 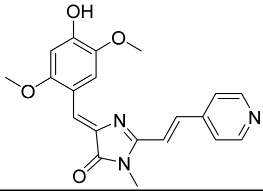 | 485 | 615 | 1.2 | 1.1 | 1.3 | 2.6 | 3.9 | 1.0 |
| **N 1056** | 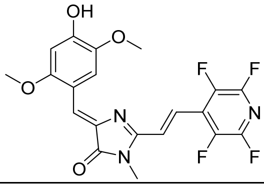 | 478 | 590 | 1.1 | 1.0 | 1.0 | 1.2 | 1.5 | 1.1 |
| **N 1027** | 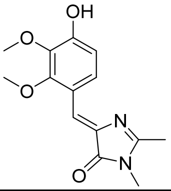 | 382 | 450 | 1.1 | 1.0 | 1.0 | 1.0 | 1.1 | 0.8 |
| **N 1068** | 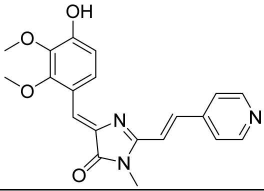 | 453 | 539 | 1.1 | 1.1 | 1.0 | 1.1 | 1.0 | 1.0 |
| **N 1042** | 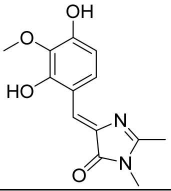 | 381 | 440 | 1.1 | 1.0 | 1.0 | 1.0 | 1.1 | 1.1 |
| **N 1069** | 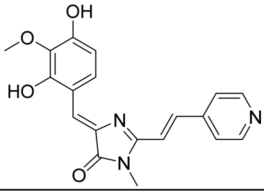 | 469 | 571 | 1.1 | 1.0 | 1.1 | 1.2 | 1.3 | 0.9 |

| | | | | | | | | | |
|---|---|---|---|---|---|---|---|---|---|
| **M 2767** | 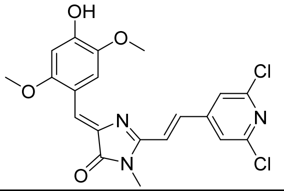 | 488 | 630 | 1.2 | 1.2 | 1.6 | 2.4 | 3.2 | 2.0 |
| **N 1179** | 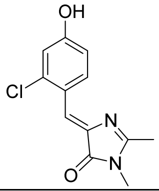 | 368 | ~400 | 1.1 | 1.0 | 1.1 | 1.0 | 1.0 | 0.9 |
| **N 1204** | 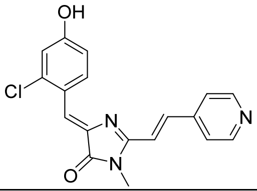 | 438 | 550 | 1.1 | 1.2 | 1.6 | 2.3 | 1.8 | 0.9 |
| **N 1180** | 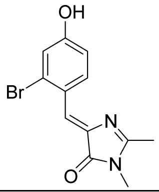 | 367 | -[c] | 1.1 | 1.0 | 1.1 | 1.1 | 1.2 | 1.0 |
| **N 1205** | 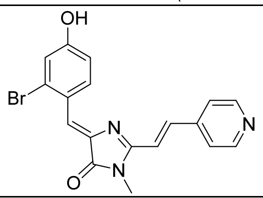 | 438 | 555 | 1.1 | 1.3 | 1.8 | 3.0 | 1.7 | 1.0 |
| **M 2876** | 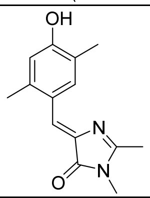 | 380 | ~400 | 1.1 | 1.1 | 1.0 | 1.0 | 1.3 | 0.6 |
| **N 1206** | 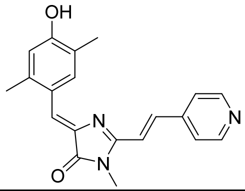 | 470 | -[c] | 1.1 | 1.1 | 1.1 | 1.0 | 1.1 | 0.9 |
| **MID 213** | 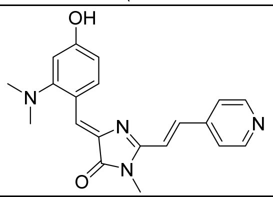 | 460 | 640 | 1.1 | 1.1 | 1.1 | 1.1 | 1.1 | 1.1 |

a – maxima position in nm;
b –in acetonitrile (non soluble enough in water);
c – non fluorescent;
d – in methanol (non soluble enough in water).

#### 6. Fluorescence quantum yields determination

Fluorescence quantum yields for complexes [protein:**HBR-DOM-2**] were calculated according to the procedure described in the literature[28] with the use of Rodamine 6G as standard. The chromophores concentration was 1.67, 0.5, and 0.167 µM for emission and 5 µM for absorption, protein were added in such concentration that lead α > 95%. The quantum yield was calculated by the formula:

$$\Phi_x = \Phi_{st} \times \frac{F_x}{F_{st}} \times \frac{f_{st}}{f_x} \times \frac{n_x^2}{n_{st}^2}$$

where *F* is the area under the emission peak, *f* is the absorption factor (see below), *n* is the refractive index of the solvent, Φ is the quantum yield, the subscript *x* corresponds to the novel compounds, the subscript *st* – for standards.

$$f = 1 - 10^{-A}$$

where *A* is absorbance at the excitation wavelength.

Data represent mean ± sem (n = 3) from various concentration.

[28] Würth C., Grabolle M., Pauli J., Spieles M., Resch-Genger U. *Nature Protocols*, **2013**, 8(8), 1535.

#### 7. Determination of affinity constants

The affinity constants for complexes [protein:**HBR-DOM2**] were determined by spectrofluorometric titration of protein by chromophore solutions with various concentrations on the Tecan Infinite 200 Pro M Nano dual mode plate reader. The protein concentration was 0.10 μM. Data represent mean ± sem (n = 3). Least squares fit (line) gave the dissociation constants $K_D$ presented in Table S7.1.

The titration experiments were performed at 25 °C in pH 7.4 PBS (#cat E404-200TABS, Amresco). Fitting was performed using Origing 8.6 software.

**Table S7.1.** Dissociation constants values of complexes [protein:**HBR-DOM2**]

| **Protein** | **$K_D$, μM** |
|---|---|
| FAST | 0.021±0.001 |
| nanoFAST | 0.85±0.01 |

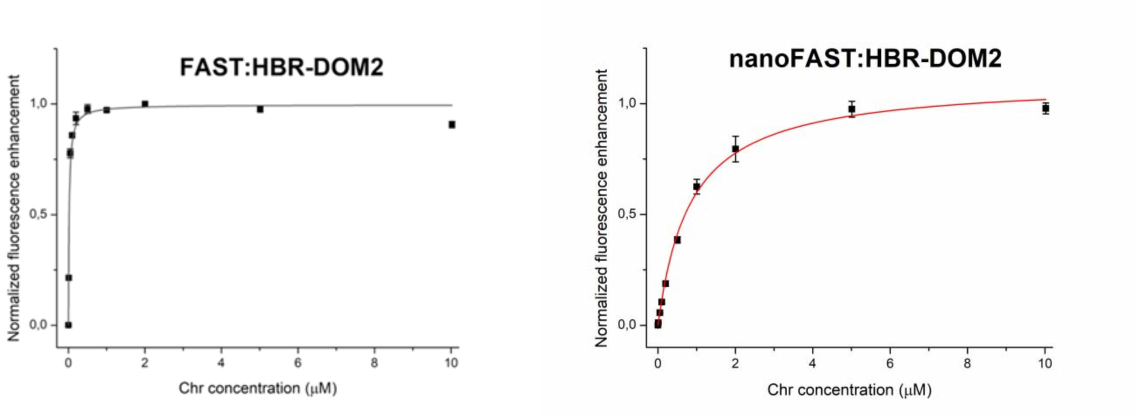


**Figure S7.1.** Titration curves. The graphs show the bound fraction for various **HBR-DOM2** concentrations.

#### 8. Fluorescence microscopy, photobleaching, and sequential imaging

Widefield fluorescence microscopy was performed on Leica 6000 with Leica HCX PL APO 100X/1.40-0.70NA oil immersion objective, Zyla sCMOS camera (Andor), and CoolLED pE-300 as a light source. TRITC cube filter was used. The standard concentration of **HBR-DOM2** for target proteins imaging, labeled with nanoFAST was 5µM

Photobleaching experiments were performed using Leica confocal microscope DMIRE2 TCS SP2 (Leica, Wetzlar, Germany) with HCX PL APO lbd.BL 63.0x1.40 OIL oil immersion objective. Live HeLa cells transiently transfected with H2B-nanoFAST (in the presence of saturating concentration of 30µM **HBR-DOM2**) or H2B-mNeonGreen were illuminated by 488 nm laser (87 µW).

Experiment with sequential staining and washout was performed using µ–Slide I 0.4 Luer ibiTreat. Injection of normal and **HBR-DOM2** (5 µM) containing HHBS (Hanks' Buffer with 20 mM Hepes) into the µ–Slide was controlled with a self-made system with multiple injectors controlled by Raspberry Pi 3 with custom python scripts. Flow speed was ~120 µl/min (that means ~1 exchange of cell media in µ–Slide volume per minute respectively).

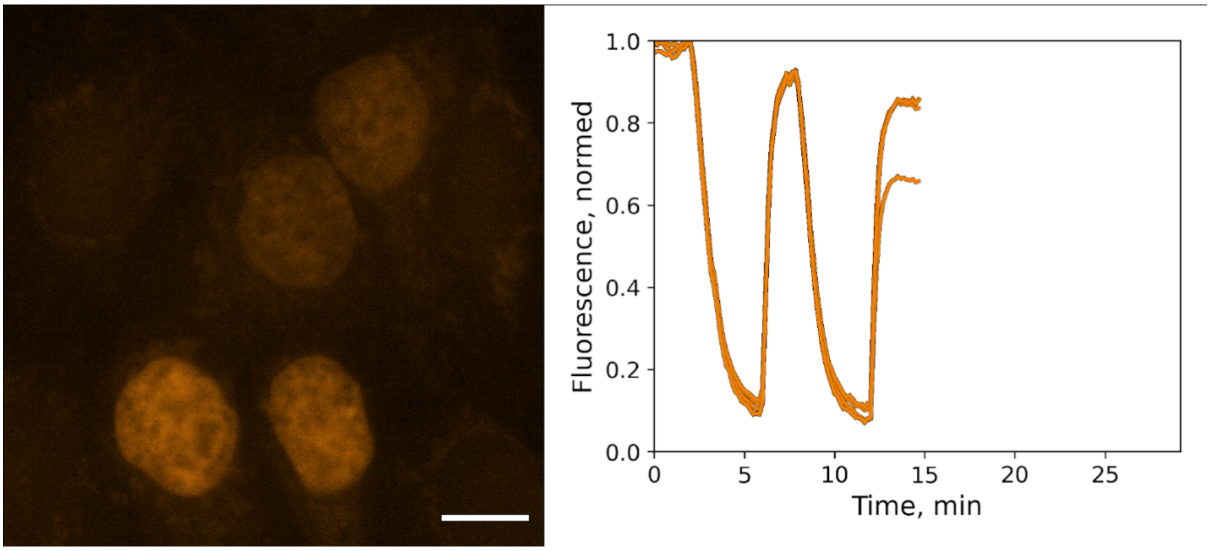


**Supplementary video 8.1**. Sequential staining and washout of live HeLa cells transiently transfected with H2B-nanoFAST construct; 5 µM concentration of **HBR-DOM2** was used for staining; Scale bar is 10 µm. Each line on the graph represents the normed mean value of fluorescence intensity in nuclei. Video plays at 30 fps.

#### 9. Synthetic procedures

**Preparation of (*Z*)-5-(4-hydroxybenzylidene)-2-thioxothiazolidin-4-one**

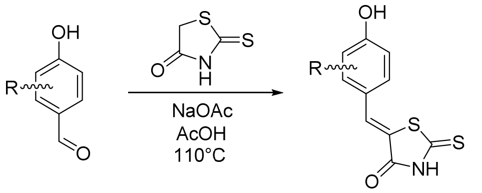


A mixture of corresponding aromatic aldehyde (2 mmol), 2-thioxothiazolidin-4-one (2.4 mmol, 320 mg), anhydrous sodium acetate (6 mmol, 492 mg) and 5 ml of glacial acetic acid were added to a screw cap test tube equipped with a magnetic stirrer and heated in an oil bath at 110°C for 2-5 hours. The progress of the reaction was monitored with TLC ($CHCl_3$/EtOH, v/v 100/5). After completion of the reaction, the reaction mixture was successively cooled to a room temperature, poured into 100 ml of water and acidified to pH=2 with a 5% water solution of hydrochloric acid. A suspension was filtered off and washed twice with 30 ml of water. Precipitate was dissolved in a mixture of $CH_2Cl_2$/MeOH (v/v, 10/1) after what n-hexane was added until the solution turned opaque. The resulted mixture was stored at 5°C for 2 hours. Filtration afforded pure product as yellow to orange powder.

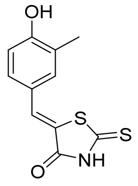

**(*Z*)-5-(4-hydroxy-3-methylbenzylidene)-2-thioxothiazolidin-4-one (HMBR)**

Previously characterized.[29]

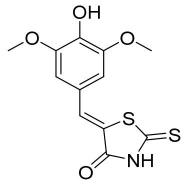

**(*Z*)-5-(4-hydroxy-3,5-dimethoxylbenzylidene)-2-thioxothiazolidin-4-one (HBR-DOM)**

Previously characterized.[30]

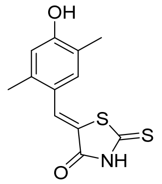

**(*Z*)-5-(4-hydroxy-2,5-dimethylbenzylidene)-2-thioxothiazolidin-4-one (HBR-2,5-DM)**

Previously characterized.[31]

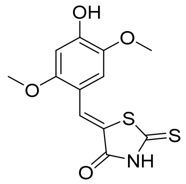

**(*Z*)-5-(4-hydroxy-2,5-dimethoxybenzylidene)-2-thioxothiazolidin-4-one (HBR-DOM2)**

Yield 60 mg (10%), red solid, mp = 278-279 ºC; $^{1}$H NMR (800 MHz, DMSO-$d_6$) δ ppm 3.80 (s, 3H), 3.82 (s, 3H), 6.62 (s, 1H), 6.87 (s, 1H), 7.75 (s, 1H), 10.21 (s, 1H), 13.55 (brs, 1H); $^{13}$C NMR (201 MHz, DMSO-$d_6$) δ ppm 55.9, 56.2, 100.5, 111.7, 113.1, 120.5, 127.4, 140.2, 142.3, 152.2, 154.8, 195.6; HRMS (ESI) m/z: 296.0057 found (calcd for $C_{12}H_{10}NO_4S_2^-$, [M-H]$^-$ 296.0059).

[29] M. A. Plamont, R. Billon-Denis, S. Maurin, C. Gauron, F. M. Pimenta, C. G. Specht, J. Shi, J. Qurard, B. Pan, J. Rossignol, K. Moncoq, N. Morellet, M. Volovitch, E. Lescop, Y. Chen, A. Triller, S. Vriz, T. Saux, L. Jullien, A. Gautier, Proc. Natl. Acad. Sci. USA **2016**, 113, 497.
[30] C. Li, M. A. Plamont, H. L. Sladitschek, V. Rodrigues, I. Aujard, P. Neveu, T. Saux, L. Jullien, A. Gautier, Chem. Sci. **2017**, 8, 5598.
[31] C. Li, M. A. Plamont, H. L. Sladitschek, V. Rodrigues, I. Aujard, P. Neveu, T. Saux, L. Jullien, A. Gautier, Chem. Sci. **2017**, 8, 5598.

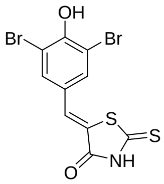

**(*Z*)-5-(3,5-dibromo-4-hydroxybenzylidene)-2-thioxothiazolidin-4-one (SAI 112)**

Yield 139 mg (16%), orange solid, mp ~ 310-312 ºC with decomposition; $^{1}H$ NMR (700 MHz, DMSO-$d_6$) δ ppm 7.54 (s, 1H), 7.75 (s, 2H), 10.95 (brs, 1H), 13.77 (brs, 1H); $^{13}C$ NMR (176 MHz, DMSO-$d_6$) δ ppm 112.4, 124.8, 127.4, 129.0, 134.1, 152.9, 169.1, 194.8; HRMS (ESI) m/z: 393.8041 found (calcd for $C_{10}H_5Br_2NO_2S_2^-$, $[M-H]^-$ 393.8035).

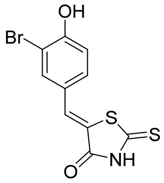

**(*Z*)-5-(3-bromo-4-hydroxybenzylidene)-2-thioxothiazolidin-4-one (SAI 117)**

Yield 417 mg (66%), yellow solid, mp = 263-265 ºC; $^{1}H$ NMR (700 MHz, DMSO-$d_6$) δ ppm 7.10 (d, *J* = 8.4 Hz, 1H), 7.42 (dd, *J* = 8.5, 2.3 Hz, 1H), 7.54 (s, 1H), 7.78 (d, *J* = 2.2 Hz, 1H), 11.22 (s, 1H), 13.73 (brs, 1H); $^{13}C$ NMR (176 MHz, DMSO-$d_6$) δ ppm 110.3, 117.1, 122.7, 125.6, 130.7, 130.9, 136.0, 156.6, 169.3, 195.2; HRMS (ESI) m/z: 313.8957 found (calcd for $C_{10}H_5BrNO_2S_2^-$, $[M-H]^-$ 313.8951).

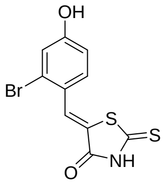

**(*Z*)-5-(2-bromo-4-hydroxybenzylidene)-2-thioxothiazolidin-4-one (SAI 122)**

Yield 120 mg (19%), orange solid, mp = 234-236 ºC; $^{1}H$ NMR (700 MHz, DMSO-$d_6$) δ ppm 6.98 (dd, *J* = 8.7, 2.5 Hz, 1H), 7.19 (d, *J* = 2.6 Hz, 1H), 7.38 (d, *J* = 8.6 Hz, 1H), 7.71 (s, 1H), 10.80 (s, 1H), 13.80 (brs, 1H); $^{13}C$ NMR (176 MHz, DMSO-$d_6$) δ ppm 116.2, 120.4, 122.9, 124.6, 127.5, 129.4, 130.9, 160.6, 169.3, 195.4; HRMS (ESI) m/z: 313.8957 found (calcd for $C_{10}H_5BrNO_2S_2^-$, $[M-H]^-$ 313.8951).

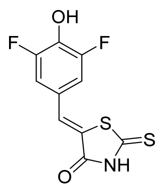

**(*Z*)-5-(3,5-difluoro-4-hydroxybenzylidene)-2-thioxothiazolidin-4-one (SAI 118)**

Yield 390 mg (71%), yellow solid, mp = 272-273 ºC; $^1$H NMR (700 MHz, DMSO-*d*$_6$) δ ppm 7.27 (d, *J* = 8.5 Hz, 2H), 7.51 (s, 1H), 11.23 (brs, 1H), 13.78 (brs, 1H); $^{13}$C NMR (176 MHz, DMSO-*d*$_6$) δ ppm 114.1 (dd, *J* = 17.3, 5.3 Hz), 123.3 (t, *J*=9.2 Hz), 124.6, 129.9, 136.5 (t, *J* = 16.4 Hz), 152.2 (dd, *J* = 243.7, 7.8 Hz), 169.2, 195.0; HRMS (ESI) m/z: 271.9654 found (calcd for $C_{10}H_4F_2NO_2S_2^-$, $[M-H]^-$ 271.9657).

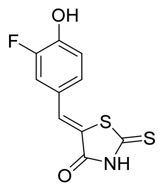

**(*Z*)-5-(3-fluoro-4-hydroxybenzylidene)-2-thioxothiazolidin-4-one (SAI 121)**

Yield 65 mg (13%), orange solid, mp = 261-263 ºC; $^1$H NMR (700 MHz, DMSO-*d*$_6$) δ ppm 7.11 (t, *J* = 8.7 Hz, 1H), 7.27 (dd, *J* = 8.4, 2.2 Hz, 1H), 7.45 (dd, *J* = 12.2, 2.3 Hz, 1H), 7.56 (s, 1H), 10.85 (s, 1H), 13.74 (brs, 1H); $^{13}$C NMR (176 MHz, DMSO-*d*$_6$) δ ppm 118.6, 122.8, 124.6 (d, *J* = 6.6 Hz), 127.6, 131.1, 132.9 (d, *J* = 3.7 Hz), 147.9 (d, *J* = 11.7 Hz), 151.0 (d, *J* = 243.2 Hz), 169.4, 195.3; HRMS (ESI) m/z: 253.9753 found (calcd for $C_{10}H_5FNO_2S_2^-$, $[M-H]^-$ 253.9751).

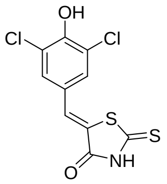


**(*Z*)-5-(3,5-dichloro-4-hydroxybenzylidene)-2-thioxothiazolidin-4-one (SAI 127)**

Yield 202 mg (33%), yellow solid, mp = 268-269 ºC; $^{1}$H NMR (700 MHz, DMSO-$d_6$) δ ppm 7.53 (s, 1H), 7.57 (s, 2H), 11.21 (brs, 1H), 13.69 (brs, 1H); $^{13}$C NMR (176 MHz, DMSO-$d_6$) δ ppm 122.8, 124.7, 125.8, 129.1, 130.3, 151.0, 169.0, 194.7; HRMS (ESI) m/z: 357.9072 found (calcd for $C_{10}H_4Cl_2NO_2S_2^-$, [M-H]$^-$ 303.9066).

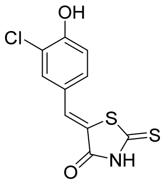


**(*Z*)-5-(3-chloro-4-hydroxybenzylidene)-2-thioxothiazolidin-4-one (SAI 125)**

Yield 158 mg (29%), orange solid, mp = 265-266 ºC; $^{1}$H NMR (700 MHz, DMSO-$d_6$) δ ppm 7.12 (d, *J* = 8.6 Hz, 1H), 7.39 (dd, *J* = 8.6, 2.3 Hz, 1H), 7.55 (s, 1H), 7.65 (d, *J* = 2.3 Hz, 1H), 11.16 (s, 1H), 13.74 (brs, 1H); $^{13}$C NMR (176 MHz, DMSO-$d_6$) δ ppm 117.4, 120.8, 122.8, 125.2, 130.3, 130.8, 132.9, 155.6, 169.3, 195.2; HRMS (ESI) m/z: 269.9455 found (calcd for $C_{10}H_7ClNO_2S_2^-$, [M-H]$^-$ 269.9456).

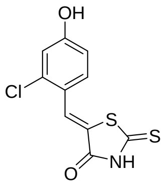


**(*Z*)-5-(2-chloro-4-hydroxybenzylidene)-2-thioxothiazolidin-4-one (SAI 199)**

Yield 433 mg (80%), orange solid, mp = 267-269 ºC; $^{1}$H NMR (700 MHz, DMSO-$d_6$) δ ppm 6.95 (dd, *J* = 8.6, 2.3 Hz, 1H), 7.02 (d, *J* = 2.3 Hz, 1H), 7.40 (d, *J* = 8.6 Hz, 1H), 7.73 (s, 1H), 10.83 (s, 1H), 13.82 (brs, 1H); $^{13}$C NMR (176 MHz, DMSO-$d_6$) δ ppm 115.8, 117.2, 121.3, 124.4, 126.6, 130.9, 136.6, 160.7, 169.3, 195.4; HRMS (ESI) m/z: 269.9455 found (calcd for $C_{10}H_5ClNO_2S_2^-$, [M-H]$^-$ 269.9456).

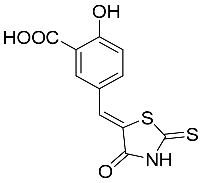

**(*Z*)-2-hydroxy-5-((4-oxo-2-thioxothiazolidin-5-ylidene)methyl)benzoic acid (SAI 120)**

Yield 247 mg (44%), dark yellow solid, mp ~ 278-280°C with decomposition; $^1$H NMR (700 MHz, DMSO-$d_6$) δ ppm 7.00 (d, *J* = 8.7 Hz, 1H), 7.58 (s, 1H), 7.65 (dd, *J* = 8.6, 2.3 Hz, 1H), 7.98 (d, *J* = 2.3 Hz, 1H); $^{13}$C NMR (176 MHz, DMSO-$d_6$) δ ppm 115.9, 118.5, 121.8, 123.0, 131.7, 132.9, 137.0, 164.4, 169.4, 170.8, 195.2; HRMS (ESI) m/z: 279.9746 found (calcd for $C_{11}H_6NO_4S_2^-$, [M-H]$^-$ 279.9744).

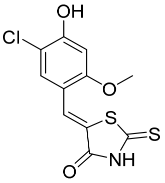

**(*Z*)-5-(5-chloro-4-hydroxy-2-methoxybenzylidene)-2-thioxothiazolidin-4-one (SAI 366)**

Yield 532 mg (59%), yellow solid, mp = 264-266°C; $^1$H NMR (700 MHz, DMSO-$d_6$) δ ppm 3.86 (s, 3H), 6.70 (s, 1H), 7.33 (s, 1H), 7.65 (s, 1H), 11.26 (s, 1H), 13.67 (brs, 1H); $^{13}$C NMR (176 MHz, DMSO-$d_6$) δ ppm 55.9, 100.3, 112.5, 113.9, 122.3, 126.1, 130.8, 157.4, 158.4, 169.4, 195.5; HRMS (ESI) m/z: 301.9705 found (calcd for $C_{11}H_9ClNO_3S_2^+$, [M+H]$^+$ 301.9707).

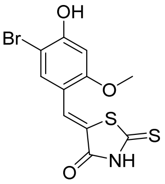

**(*Z*)-5-(5-bromo-4-hydroxy-2-methoxybenzylidene)-2-thioxothiazolidin-4-one (SAI 365)**

Yield 829 mg (80%), yellow solid, mp = 232-234°C; $^1$H NMR (700 MHz, DMSO-$d_6$) δ ppm 3.86 (s, 3H), 6.68 (s, 1H), 7.45 (s, 1H), 7.64 (s, 1H), 11.33 (s, 1H), 13.67 (brs, 1H); $^{13}$C NMR (176 MHz, DMSO-$d_6$) δ ppm 55.8, 100.1, 101.1, 114.5, 122.3, 126.1, 133.8, 158.4, 159.0, 169.4, 195.5; HRMS (ESI) m/z: 345.9195 found (calcd for $C_{11}H_9BrNO_3S_2^+$, [M+H]$^+$ 345.9202).

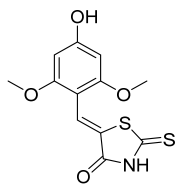

**(*Z*)-5-(4-hydroxy-2,6-dimethoxybenzylidene)-2-thioxothiazolidin-4-one (SAI 363)**

Yield 566 mg (63%), orange solid, mp = 292-294°C with decomposition; a mixture of two isomers was observed; isomer's peaks are highlighted with an asterisk sign (see NMR copy); major isomer: $^{1}$H NMR (700 MHz, DMSO-*d6*) δ ppm 3.83 (s, 6H), 6.14 (s, 2H), 7.84 (s, 1H), 10.57 (s, 1H), 13.31 (brs, 1H); minor isomer: $^{1}$H NMR (700 MHz, DMSO-*d6*) δ ppm 3.69 (s, 6H), 6.07 (s, 2H), 7.11 (s, 1H), 9.99 (s, 1H), 13.21 (brs, 1H); $^{13}$C NMR (176 MHz, DMSO-*d*6) δ 55.5, 92.2, 102.2, 120.9, 126.3, 160.3, 164.0, 169.9, 197.2; HRMS (ESI) m/z: 298.0203 found (calcd for $C_{12}H_{12}NO_4S_2^+$, $[M+H]^+$ 298.0202).

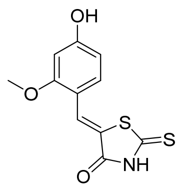

**(*Z*)-5-(4-hydroxy-2-methoxybenzylidene)-2-thioxothiazolidin-4-one (SAI 362)**

Yield 602 mg (75%), orange powder, mp = 297-300°C with decomposition; $^{1}$H NMR (700 MHz, DMSO-*d*6) δ ppm 3.85 (s, 3H), 6.51 (d, *J* = 2.3 Hz, 1H), 6.55 (dd, *J* = 8.6, 2.3 Hz, 1H), 7.24 (d, *J* = 8.6 Hz, 1H), 7.75 (s, 1H), 10.51 (s, 1H), 13.58 (brs, 1H); $^{13}$C NMR (176 MHz, DMSO-*d*6) δ ppm 55.6, 99.4, 109.0, 112.8, 120.3, 127.4, 131.6, 160.4, 162.7, 169.6, 195.8; HRMS (ESI) m/z: 268.0097 found (calcd for $C_{11}H_{10}NO_3S_2^+$, $[M+H]^+$ 268.0097).

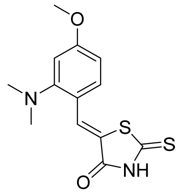

**(*Z*)-5-(2-(dimethylamino)-4-methoxybenzylidene)-2-thioxothiazolidin-4-one (SAI 364)**

Yield 747 mg (85%), orange powder, mp = 175-178°C; $^{1}$H NMR (700 MHz, DMSO-*d6*) δ ppm 2.68 (s, 6H), 3.83 (s, 3H), 6.69 (d, J = 2.5 Hz, 1H), 6.72 (dd, J = 8.6, 2.5 Hz, 1H), 7.42 (d, J = 8.6 Hz, 1H), 7.63 (s, 1H), 13.50 (brs, 1H); $^{13}$C NMR (176 MHz, DMSO-*d*6) δ 44.0, 55.4, 104.9, 108.2, 118.4, 121.6, 130.8, 133.2, 155.3, 162.5, 170.1, 197.2; HRMS (ESI) m/z: 295.0575 found (calcd for $C_{13}H_{15}N_2O_2S_2^+$, $[M+H]^+$ 295.056).

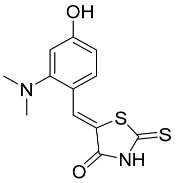


**(*Z*)-5-(2-(dimethylamino)-4-hydroxybenzylidene)-2-thioxothiazolidin-4-one (SAI 367)**

(*Z*)-5-(2-(dimethylamino)-4-methoxybenzylidene)-2-thioxothiazolidin-4-one (294 mg, 1 mmol) was suspended in dry DCM (10 ml), the flask was cooled in an ice-cold water bath. A solution of $BBr_3$ in DCM (1M, 5 ml, 5 mmol) was added dropwise under vigorous stirring. The reaction mixture was stirred at 0°C for an hour, then it was allowed to warm to room temperature and continued to stir for 5 h. After completion, the reaction mixture was diluted with DCM (100 ml), EtOH (10 ml). Organic layer was washed with water (1x100 ml) and dried over $Na_2SO_4$. The solvents were removed under reduced pressure. The crude product was purified by silica gel column chromatography ($CHCl_3$/EtOH, v/v 100/1) to afford the desired compound as a deep red solid. Yield 134 mg (48%), orange solid, mp = 256-258°C with decomposition; $^1$H NMR (700 MHz, DMSO-*d6*) δ ppm 2.64 (s, 6H), 6.54 (dd, J = 8.4, 2.4 Hz, 1H), 6.56 (d, J = 2.3 Hz, 1H), 7.30 (d, J = 8.4 Hz, 1H), 7.60 (s, 1H), 10.22 (s, 1H), 13.45 (brs, 1H); $^{13}$C NMR (176 MHz, DMSO-*d6*) δ ppm 44.1, 105.9, 110.2, 117.0, 120.9, 130.8, 133.3, 155.7, 161.4, 170.8, 197.4; HRMS (ESI) m/z: 281.0413 found (calcd for $C_{12}H_{13}N_2O_2S_2^+$, $[M+H]^+$ 281.0413).

**Preparation of 5-(*Z*)-benzylidene-2-methyl-3-alkyl/aryl-3,5-dihydro-4H-imidazol-4-ones**

General method was reported previously.[32]

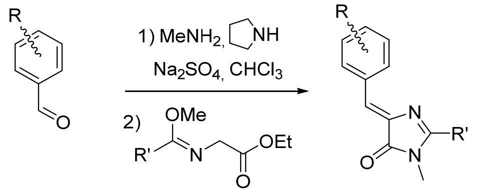


The corresponding aromatic aldehyde (10 mmol) was dissolved in $CHCl_3$ (50 mL) and mixed with methylamine solution (40% aqueous, 2.5 mL), pyrrolidine (7 mg, 0.1 mmol) and anhydrous $Na_2SO_4$ (10 g). The mixture was stirred for 48 h at room temperature, filtered and dried over the additional $Na_2SO_4$. The solvent was evaporated and corresponding ethyl((methoxy)amino)acetate (20 mmol) was added to the residue (5-10 mL of methanol was also added if it wouldn't mixed). The mixture was stirred for 24 h at room temperature, solvents were evaporated and the product was purified by column chromatography ($CHCl_3$-EtOH).

Most of imidazolones we used in this work were previously described and characterized.

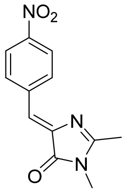


**(*Z*)-2,3-dimethyl-5-(4-nitrobenzylidene)-3,5-dihydro-4H-imidazol-4-one**

Previously characterized.[33]

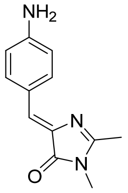


**(*Z*)-5-(4-aminobenzylidene)-2,3-dimethyl-3,5-dihydro-4H-imidazol-4-one**

Previously characterized.[34]

[32] Baldridge A., Kowalik J., Tolbert L. M. *Synthesisю* **2010**, 14, 2424.

[33] Zaitseva S.O., Golodukhina S.V., Baleeva N.S., Levina E.A., Smirnov A.Yu., Zagudaylova M.B., Baranov M.S. Chem. Select, **2018**, 3 (30), 8593-8596.

[34] Baranov, M. S., Solntsev, K. M., Baleeva, N. S., Mishin, A. S., Lukyanov, S. A., Lukyanov, K. A. and Yampolsky, I. V., Chem. Eur. J., **2014**, 20, 13234-13241.

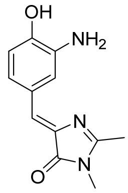

**(*Z*)-5-(3-amino-4-hydroxybenzylidene)-2,3-dimethyl-3,5-dihydro-4H-imidazol-4-one**

Previously characterized.[35]

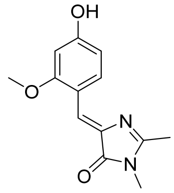

**(Z)-5-(4-hydroxy-2-methoxybenzylidene)-2,3-dimethyl-3,5-dihydro-4H-imidazol-4-one**

Previously characterized.[36]

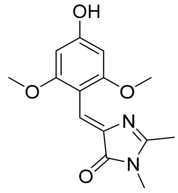

**(*Z*)-5-(4-hydroxy-2,6-dimethoxybenzylidene)-2,3-dimethyl-3,5-dihydro-4H-imidazol-4-one (N 976)**

Orange solid (1.32 g, 48%); mp ~200 ºC with decomposition; $^{1}$H NMR (700 MHz, DMSO-*d*$_6$) δ ppm 9.87 (br. s., 1 H), 6.89 (s, 1 H), 6.10 (s, 2 H), 3.70 (s, 6 H), 3.05 (s, 3 H), 2.22 (s, 3 H); $^{13}$C NMR (75 MHz, DMSO-*d*$_6$) δ ppm 169.5, 161.0, 160.6, 159.6, 138.8, 120.2, 103.2, 92.2, 55.6, 26.2, 15.2; HRMS (ESI) m/z: 277.1184 found (calcd for $C_{14}H_{17}N_2O_4^+$, $[M+H]^+$ 277.1183).

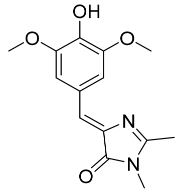

**(*Z*)-5-(4-hydroxy-3,5-dimethoxybenzylidene)-2,3-dimethyl-3,5-dihydro-4H-imidazol-4-one (N 979)**

$^{1}$H NMR (300 MHz, DMSO-*d*$_6$) δ ppm 9.15 (br. s., 1 H), 7.63 (s, 2 H), 6.90 (s, 1 H), 3.79 (s, 6 H), 3.08 (s, 3 H), 2.33 (s, 3 H). Previously characterized.[37]

---

[35] Chen C., Baranov M.S., Zhu L., Tang L, Baleeva N.S., Smirnov A.Yu., Zaitseva S.O., Yampolsky I.V., Solntsev K.M., Fang C., *Chem. Commun.*, **2019**, 55, 2537-2540.

[36] Povarova N.V., Zaitseva S.O., Baleeva N.S., Smirnov A.Yu., Myasnyanko I.N., Zagudaylova M.B., Bozhanova N.G., Gorbachev D.A., Malyshevskaya K.K., Gavrikov A.S., Mishin A.S., Baranov M.S. Chem. Eur. J. **2019**, 25 (41), 9592-9596.

[37] Paige J.S., Wu K.Y., Jaffrey S.R., Science, **2011**, 333 (6042), 642-646.

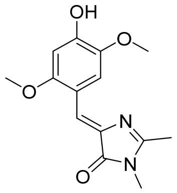


**(*Z*)-5-(4-hydroxy-2,5-dimethoxybenzylidene)-2,3-dimethyl-3,5-dihydro-4H-imidazol-4-one (N 1036)**

Orange solid (1.38 g, 50%); mp = 202-205 ºC; $^1$H NMR (300 MHz, DMSO-$d_6$) δ ppm 9.99 (br. s., 1 H), 8.49 (s, 1 H), 7.24 (s, 1 H), 6.53 (s, 1 H), 3.82-3.72 (m, 6 H), 3.08 (s, 3 H), 2.32 (s, 3 H); $^{13}$C NMR (75 MHz, DMSO-$d_6$) δ ppm 169.8, 161.5, 155.0, 151.2, 141.8, 135.3, 118.6, 115.8, 113.1, 99.8, 56.3, 56.0, 26.2, 15.5; HRMS (ESI) m/z: 277.1179 found (calcd for $C_{14}H_{17}N_2O_4^+$, $[M+H]^+$ 277.1183).

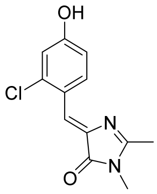


**(*Z*)-5-(2-chloro-4-hyroxybenzylidene)-2,3-dimethyl-3,5-dihydro-4H-imidazol-4-one (N 1179)**

Brown solid (2.05 g, 80%); mp = 152-155 ºC; $^1$H NMR (700 MHz, DMSO-$d_6$) δ ppm 8.78 (d, *J*=8.8 Hz, 1 H), 7.15 (s, 1 H), 6.87 (d, *J*=2.1 Hz, 1 H), 6.82 (dd, *J*=8.7, 2.0 Hz, 1 H), 3.09 (s, 3 H), 2.34 (s, 3 H); $^{13}$C NMR (176 MHz, DMSO-$d_6$) δ ppm 169.7, 163.7, 161.2, 137.3, 136.1, 134.3, 121.7, 119.1, 116.4, 115.6, 26.2, 15.3; HRMS (ESI) m/z: 251.0583 found (calcd for $C_{12}H_{12}ClN_2O_2^+$, $[M+H]^+$ 251.0582).

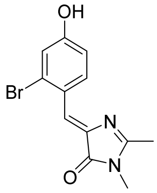


**(*Z*)-5-(2-bromo-4-hyroxybenzylidene)-2,3-dimethyl-3,5-dihydro-4H-imidazol-4-one (N 1180)**

Brown solid (2.23 g, 76%); mp = 174-177 ºC; $^1$H NMR (700 MHz, DMSO-$d_6$) δ ppm 8.78 (d, *J*=8.8 Hz, 1 H), 7.14 (s, 1 H), 7.04 (br. s., 1 H), 6.84 (d, *J*=8.8 Hz, 1 H), 3.09 (s, 3 H), 2.34 (s, 3 H); $^{13}$C NMR (176 MHz, DMSO-$d_6$) δ ppm 169.7, 163.5, 161.6, 137.1, 134.3, 127.4, 122.8, 122.1, 119.8, 116.1, 26.2, 15.3; HRMS (ESI) m/z: 295.0078 found (calcd for $C_{12}H_{12}BrN_2O_2^+$, $[M+H]^+$ 295.0077).

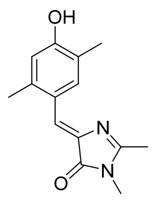

**(*Z*)-5-(4-hyroxy-2,5-dimethylbenzylidene)-2,3-dimethyl-3,5-dihydro-4H-imidazol-4-one (M 2876)**

Yellow solid (1.98 g, 81%); mp = 239-242 ºC; $^{1}$H NMR (700 MHz, DMSO-*d*6) δ ppm 9.87 (br. s., 1 H), 8.54 (s, 1 H), 7.02 (s, 1 H), 6.66 (s, 1 H), 3.09 (s, 3 H), 2.33 (s, 3 H), 2.32 (s, 3 H), 2.10 (s, 3 H); $^{13}$C NMR (176 MHz, DMSO-*d*6) δ ppm 169.9, 162.2, 157.5, 138.7, 135.9, 134.4, 123.4, 122.0, 121.7, 116.5, 26.1, 19.2, 15.7, 15.2; HRMS (ESI) m/z: 245.1280 found (calcd for $C_{14}H_{17}N_2O_2^+$, $[M+H]^+$ 245.1285).

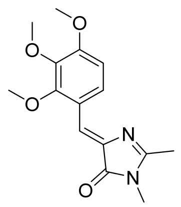

**(*Z*)-5-(2,3,4-trimethoxybenzylidene)-2,3-dimethyl-3,5-dihydro-4H-imidazol-4-one (N 1043)**

Yellow solid (1.86 g, 64%); mp = 192-195 ºC; $^{1}$H NMR (700 MHz, DMSO-*d*6) δ ppm 8.53 (d, *J*=9.0 Hz, 1 H), 7.13 (s, 1 H), 6.95 (d, *J*=9.0 Hz, 1 H), 3.88-3.85 (m, 6 H), 3.77 (s, 3 H), 3.09 (s, 3 H), 2.34 (s, 3 H); $^{13}$C NMR (75 MHz, DMSO-*d*6) δ ppm 169.9, 163.5, 155.4, 153.3, 141.4, 137.6, 127.6, 120.4, 118.0, 108.4, 61.7, 60.5, 56.0, 26.3, 15.4; HRMS (ESI) m/z: 291.1337 found (calcd for $C_{15}H_{19}N_2O_4^+$, $[M+H]^+$ 291.1339).

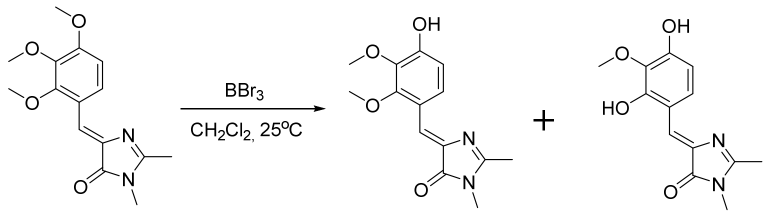


To the solution of (Z)-5-(2,3,4-trimethoxybenzylidene)-2,3-dimethyl-3,5-dihydro-4H-imidazol-4-one (5 mmol) in $CH_2Cl_2$ (50 mL) borone tribromide in $CH_2Cl_2$ (1M, 5 mL, 5 mmol) were added. The mixture was stirred for 2 h at room temperature under inert atmosphere. The mixture was diluted with $CHCl_3$ (100 mL), washed with water (2x50 mL) and brine (2x50 mL) and dried over $Na_2SO_4$. The solvent was evaporated and the mixture of products was separated by column chromatography ($CHCl_3$-EtOH, 49:1).

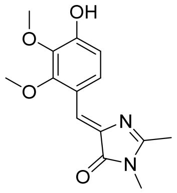

**(*Z*)-5-(4-hydroxy-2,3-dimethoxybenzylidene)-2,3-dimethyl-3,5-dihydro-4H-imidazol-4-one (N 1027)**

Yellow solid (0.41 g, 30%); mp = 193-196 ºC; $^{1}H$ NMR (300 MHz, DMSO-*d*6) δ ppm 12.44 (br. s., 1 H), 7.75 (d, *J*=8.9 Hz, 1 H), 7.15 (s, 1 H), 6.64 (d, *J*=9.0 Hz, 1 H), 3.83 (s, 3 H), 3.67 (s, 3 H), 3.12 (s, 3 H), 2.37 (s, 3 H); $^{13}C$ NMR (75 MHz, DMSO-*d*6)  ppm 168.1, 160.3, 156.2, 151.9, 136.7, 133.0, 130.4, 124.5, 115.1, 104.1, 59.8, 55.9, 26.4, 15.1; HRMS (ESI) m/z: 277.1178 found (calcd for $C_{14}H_{17}N_2O_4^+$, $[M+H]^+$ 277.1183).

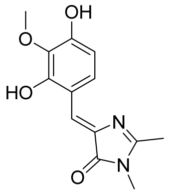

**(*Z*)-5-(2,4-dihydroxy-3-methoxybenzylidene)-2,3-dimethyl-3,5-dihydro-4H-imidazol-4-one (N 1042)**

Orange solid (0.46 g, 35%); mp =231-234 ºC; $^{1}H$ NMR (300 MHz, DMSO-*d*6) δ ppm 12.71 (br. s., 1 H), 8.41 (br. s., 1 H), 7.37 (d, *J*=8.8 Hz, 1 H), 7.12 (s, 1 H), 6.58 (d, *J*=8.8 Hz, 1 H), 3.81 (s, 3 H), 3.12 (s, 3 H), 2.36 (s, 3 H); $^{13}C$ NMR (75 MHz, DMSO-*d*6) δ ppm 167.9, 159.6, 151.3, 146.7, 134.6, 132.3, 125.9, 125.7, 114.4, 104.1, 55.8, 26.4, 15.1; HRMS (ESI) m/z: 263.1024 found (calcd for $C_{13}H_{15}N_2O_4^+$, $[M+H]^+$ 263.1026).

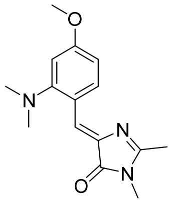

**(*Z*)-5-(2-(dimethylamino)-4-methoxybenzylidene)-2,3-dimethyl-3,5-dihydro-4H-imidazol-4-one (MID 173)**

Yellow solid (1.53 g, 56%); mp =181-184 ºC; $^{1}H$ NMR (700 MHz, DMSO-*d*6) δ ppm 8.58 (d, *J*=8.8 Hz, 1 H), 7.19 (s, 1 H), 6.69 (dd, *J*=8.8, 2.4 Hz, 1 H), 6.62 (d, *J*=2.4 Hz, 1 H), 3.80 (s, 3 H), 3.09 (s, 3 H), 2.72 (s, 6 H), 2.33 (s, 3 H); $^{13}C$ NMR (75 MHz, DMSO-*d*6) δ ppm 170.1, 162.2, 161.5, 156.7, 136.0, 134.3, 121.7, 119.5, 108.2, 104.3, 55.2, 45.0 (2 C), 26.2, 15.3; HRMS (ESI) m/z: 274.1546 found (calcd for $C_{15}H_{20}N_3O_2^+$, $[M+H]^+$ 274.1550).

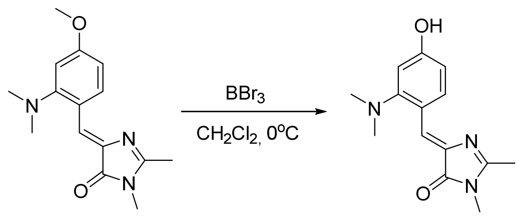


**(*Z*)-5-(2-(dimethylamino)-4-hydroxybenzylidene)-2,3-dimethyl-3,5-dihydro-4H-imidazol-4-one (MID 198)**

To the solution of (Z)-5-(2-(dimethylamino)-4-methoxybenzylidene)-2,3-dimethyl-3,5-dihydro-4H-imidazol-4-one (1.37 g, 5 mmol) in $CH_2Cl_2$ (45 mL) borone tribromide in $CH_2Cl_2$ (1M, 40 mL, 40 mmol) were added at 0 $^oC$. The mixture was stirred for 3 h at room temperature under inert atmosphere. The mixture was diluted with $NaHCO_3$ (100 ml), washed with water (2x50 mL) and brine (2x50 mL) and dried over $Na_2SO_4$. The solvent was evaporated and the product was purified by column chromatography ($CHCl_3$-EtOH, 9:1). Red solid (129 mg, 10 %); mp = 210-213 °C; $^1$H NMR (700 MHz, DMSO-$d_6$) δ ppm 9.95 (s, 1 H), 8.51 (d, *J*=8.4 Hz, 1 H), 7.19 (s, 1 H), 6.46 - 6.55 (m, 2 H), 3.08 (s, 3 H), 2.68 (s, 6 H), 2.31 (s, 3 H); $^{13}$C NMR (75 MHz, DMSO-$d_6$) □ ppm 170.1, 161.5, 160.3, 157.0, 135.2, 134.5, 122.2, 118.1, 110.1, 105.4, 45.0, 26.2, 15.2; HRMS (ESI) m/z: 260.1391 found (calcd for $C_{14}H_{18}N_3O_2^+$, $[M+H]^+$ 260.1394).

**Preparation of 5-(Z)-benzylidene-2-(E)-arylvinyl-3-methyl-3,5-dihydro-4H-imidazol-4-ones**

**Method A**

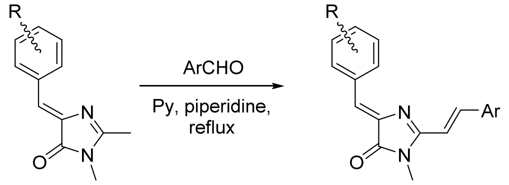


To the solution of 5-(Z)-benzylidene-2,3-dimethyl-3,5-dihydro-4H-imidazol-4-one (1 mmol) in pyridine (5 mL) piperidine (0.03 mL) and corresponding aldehyde (5 mmol) were added. The mixture was refluxed for 8-12 h and the solvent was evaporated. The product was purified by column chromatography ($CHCl_3$-EtOH, typically 19:1).

**Method B**

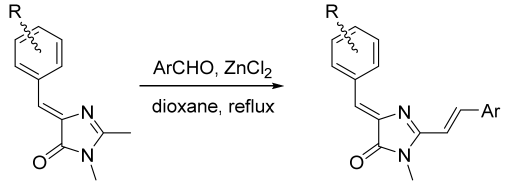


To the solution of 5-(Z)-benzylidene-2,3-dimethyl-3,5-dihydro-4H-imidazol-4-one (1 mmol) in dioxane (5 mL) anhydrous zinc chloride (30 mg, 0.22 mmol) and corresponding aldehyde (1.2 mmol) were added. The mixture was refluxed for 1-6 h and the solvent was evaporated. The mixture was dissolved in EtOAc (50 mL) and washed by EDTA solution (0.5%, 10 mL), water (3x10 mL) and brine (1x10 mL). The mixture was dried over $Na_2SO_4$. The solvent was evaporated and the product was purified by column chromatography ($CHCl_3$-EtOH, typically 19:1).

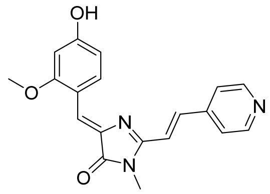


**(Z)-5-(4-hydroxy-2-methoxybenzylidene)-3-methyl-2-((E)-2-(pyridin-4-yl)vinyl)-3,5-dihydro-4H-imidazol-4-one (N871b)**

Synthesized by the **method A**. Previously characterized.[38]

[38] Povarova N.V., Zaitseva S.O., Baleeva N.S., Smirnov A.Yu., Myasnyanko I.N., Zagudaylova M.B., Bozhanova N.G., Gorbachev D.A., Malyshevskaya K.K., Gavrikov A.S., Mishin A.S., Baranov M.S. Chem. Eur. J. **2019**, 25 (41), 9592-9596.

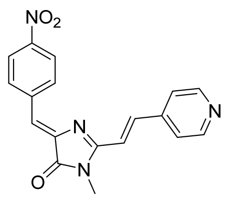


**5-((Z)-4-nitrobenzylidene)-3-methyl-2-((E)-2-(pyrydin-4-yl)vinyl)-3,5-dihydro-4H-imidazol-4-one (ZS 315)**

Synthesized by the **method A**.

Orange solid (157 mg, 47%); mp ~250 ºC with decomposition; $^{1}H$ NMR (700 MHz, DMSO-*d*6) δ ppm 8.70 (d, *J*=5.9 Hz, 2 H), 8.58 (d, *J*=8.8 Hz, 2 H), 8.30 (d, *J*=9.0 Hz, 2 H), 8.10 (d, *J*=15.8 Hz, 1 H), 7.84 (d, *J*=5.9 Hz, 2 H), 7.56 (d, *J*=15.8 Hz, 1 H), 7.21 (s, 1 H), 3.32 (s, 3 H); $^{13}C$ NMR (176 MHz, DMSO-*d*6) δ ppm 169.7, 162.4, 150.2, 147.1, 141.8, 141.7, 140.6, 138.9, 132.7, 123.4, 122.3, 122.0, 118.3, 26.5; HRMS (ESI) m/z: 335.1142 found (calcd for $C_{18}H_{15}N_4O_3^+$, $[M+H]^+$ 335.1139).

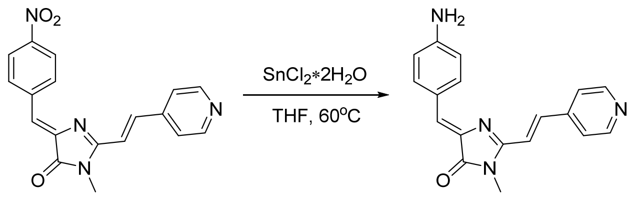


**5-((Z)-4-aminobenzylidene)-3-methyl-2-((E)-2-(pyrydin-4-yl)vinyl)-3,5-dihydro-4H-imidazol-4-one (ZS 319.2)**

5-((Z)-4-nitrobenzylidene)-3-methyl-2-((E)-2-(pyrydin-4-yl)vinyl)-3,5-dihydro-4H-midazol-4-one (0.4 mmol) and tin (II) chloride dihydrate (0.65g, 3 mmol) were suspended in 7 mL of THF. The mixture was refluxed for 3h. Afterwards it was neutralized with 10% $Na_2CO_3$ solution, filtered, extracted with dichloromethane (4x75 mL) and dried over $Na_2SO_4$. The solvent was evaporated and crude product was purified by column chromatography ($CHCl_3$-EtOH 99:1).

Red solid (77 mg, 63%); mp = 165-169 ºC; $^{1}H$ NMR (700 MHz, DMSO-*d*6) δ ppm 8.64 (d, *J*=5.7 Hz, 2 H), 8.04 (br. d., *J*=7.8 Hz, 2 H), 7.85 (d, *J*=15.8 Hz, 1 H), 7.80 (d, *J*=5.9 Hz, 2 H), 7.45 (d, *J*=15.8 Hz, 1 H), 6.96 (s, 1 H), 6.64 (d, *J*=8.8 Hz, 2 H), 6.15 (br. s., 2 H), 3.28 (s, 3 H); $^{13}C$ NMR (201 MHz, DMSO-*d*6) δ ppm 169.5, 155.7, 152.1, 150.1, 142.5, 135.2, 135.0, 134.5, 128.8, 121.9, 121.8, 119.0, 113.7, 26.4; HRMS (ESI) m/z: 305.1399 found (calcd for $C_{18}H_{17}N_4O^+$, $[M+H]^+$ 305.1397).

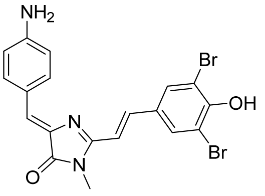

**5-((Z)-4-aminobenzylidene)-2-((E)-3,5-dibromo-4-hyrdoxystyryl)-3-methyl-3,5-dihydro-4H-imidazol-4-one (ZS 309)**

Synthesized by the **method A**.

Red solid (190 mg, 40%); mp ~260 ºC with decomposition; $^{1}$H NMR (700 MHz, DMSO-*d*$_6$) δ ppm 10.35 (br. s., 1 H), 8.12 (s, 2 H), 8.02 (d, *J*=8.2 Hz, 2 H), 7.78 (d, *J*=15.8 Hz, 1 H), 7.17 (d, *J*=15.8 Hz, 1 H), 6.87 (s, 1 H), 6.62 (d, *J*=8.8 Hz, 2 H), 6.07 (br. s., 2 H), 3.25 (s, 3 H); $^{13}$C NMR (75 MHz, DMSO-*d*$_6$) δ ppm 169.8, 156.5, 151.7, 135.5, 134.8, 134.7, 131.9, 130.3, 127.4, 122.1, 114.1, 113.7, 112.2, 26.4; HRMS (ESI) m/z: 477.9589 found (calcd for $C_{19}H_{16}Br_2N_3O_2^+$, $[M+H]^+$ 477.9583).

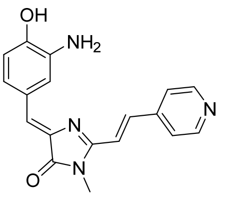

**5-((Z)-3-amino-4-hydroxybenzylidene)-3-methyl-2-((E)-2-(pyridin-4-yl)vinyl)-3,5-dihydro-4H-imidazol-4-one (ZS 325)**

Synthesized by the **method A**. $^{1}$H NMR (700 MHz, DMSO-*d*$_6$) δ ppm 8.67 (br. s., 2H), 7.97 (d, *J*=15.8 Hz, 1H), 7.87 (s, 1H), 7.79 (d, *J*=4.8 Hz, 2H), 7.48 (d, *J*=15.8 Hz, 1H), 7.27 (d, *J*=7.8 Hz, 1H), 6.89 (s, 1H), 6.75 (d, *J*=8.2 Hz, 1H), 3.28 (s, 3H). Previously characterized.[39]

[39] Chen C., Baranov M.S., Zhu L., Tang L, Baleeva N.S., Smirnov A.Yu., Zaitseva S.O., Yampolsky I.V., Solntsev K.M., Fang C., *Chem. Commun.*, **2019**, 55, 2537-2540.

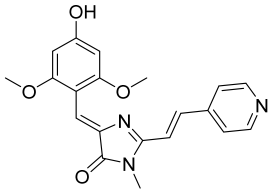


**5-((Z)-4-hydroxy-2,6-dimethoxybenzylidene)-3-methyl-2-((E)-2-(pyridin-4-yl)vinyl)-3,5-dihydro-4H-imidazol-4-one (ZS 316)**

Synthesized by the **method A**.

Red solid (146 mg, 40%); mp = 193-196 ºC; $^{1}$H NMR (700 MHz, DMSO-$d_6$) δ ppm 10.08 (s, 1 H), 8.63 (d, *J*=5.9 Hz, 2 H), 7.70 - 7.75 (m, 3 H), 7.43 (d, *J*=15.8 Hz, 1 H), 7.12 (s, 1 H), 6.15 (s, 2 H), 3.77 (s, 6 H); $^{13}$C NMR (75 MHz, DMSO-$d_6$) δ ppm 169.7, 161.8, 160.2, 156.3, 150.3, 142.3, 138.3, 135.8, 122.1, 121.9, 119.2, 103.9, 92.4, 55.7, 26.5; HRMS (ESI) m/z: 366.1448 found (calcd for $C_{20}H_{20}N_3O_4^+$, $[M+H]^+$ 366.1448).

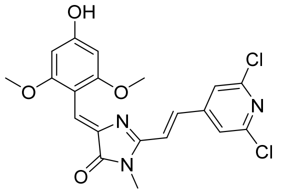


**5-((Z)-4-hydroxy-2,6-dimethoxybenzylidene)-2-((E)-2-(2,6-dichloropyridin-4-yl)vinyl)-3-methyl-3,5-dihydro-4H-imidazol-4-one (N 973)**

Synthesized by the **method B**.

Dark red solid (195 mg, 45%); mp ~260 ºC with decomposition; $^{1}$H NMR (700 MHz, DMSO-$d_6$) δ ppm 10.13 (s, 1 H), 8.03 (s, 2 H), 7.68 (d, *J*=15.8 Hz, 1 H), 7.61 (d, *J*=15.8 Hz, 1 H), 7.17 (s, 1 H), 6.14 (s, 2 H), 3.77 (s, 6 H), 3.25 (s, 3 H); $^{13}$C NMR (201 MHz, DMSO-$d_6$) δ ppm 169.5, 162.0, 160.3, 155.7, 149.9, 149.1, 138.1, 132.7, 122.8, 121.5, 120.8, 104.0, 92.4, 55.7, 26.4; HRMS (ESI) m/z: 434.0678 found (calcd for $C_{20}H_{18}Cl_2N_3O_4^+$, $[M+H]^+$ 434.0669).

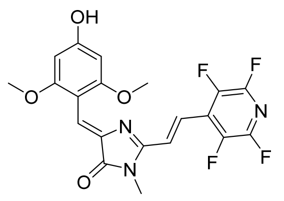

**5-((Z)-4-hydroxy-2,6-dimethoxybenzylidene)-3-methyl-2-((E)-2-(perfluoropyridin-4-yl)vinyl)-3,5-dihydro-4H-imidazol-4-one (N 971)**

Synthesized by the **method B**.

Red solid (131 mg, 30%); mp ~270 ºC with decomposition; $^{1}$H NMR (700 MHz, DMSO-$d_6$) δ ppm 10.24 (s, 1 H), 7.73 (d, *J*=16.0 Hz, 1 H), 7.43 (d, *J*=16.0 Hz, 1 H), 7.27 (s, 1 H), 6.15 (s, 2 H), 3.78 (s, 6 H), 3.22 (s, 3 H); $^{13}$C NMR (201 MHz, DMSO-$d_6$) δ ppm 169.5, 162.6, 160.6, 155.1, 143.0 (dm, *J*=242.1 Hz), 139.7 (dm, *J*=260.8 Hz), 137.2, 127.3, 126.4, 123.5, 120.6, 103.9, 92.4, 55.6, 26.2; HRMS (ESI) m/z: 438.1075 found (calcd for $C_{20}H_{16}F_4N_3O_4^+$, $[M+H]^+$ 438.1071).

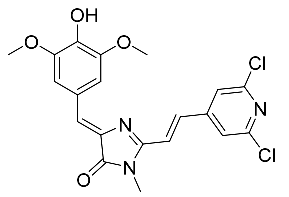

**5-((Z)-4-hydroxy-3,5-dimethoxybenzylidene)-2-((E)-2-(2,6-dichloropyridin-4-yl)vinyl)-3-methyl-3,5-dihydro-4H-imidazol-4-one (N 960a)**

Synthesized by the **method B**.

Red solid (329 mg, 76%); mp = 287-290 ºC; $^{1}$H NMR (700 MHz, DMSO-$d_6$) δ ppm 9.31 (s, 1 H), 8.06 (s, 2 H), 7.82 (d, *J*=15.8 Hz, 1 H), 7.74 (s, 2 H), 7.66 (d, *J*=15.8 Hz, 1 H), 7.09 (s, 1 H), 3.86 (s, 6 H), 3.29 (s, 3 H); $^{13}$C NMR (75 MHz, DMSO-$d_6$) δ ppm 169.6, 157.5, 150.0, 149.0, 147.9, 139.3, 137.0, 133.3, 128.6, 124.7, 121.8, 121.7, 110.5, 56.1, 26.5; HRMS (ESI) m/z: 434.0685 found (calcd for $C_{20}H_{18}Cl_2N_3O_4^+$, $[M+H]^+$ 434.0669).

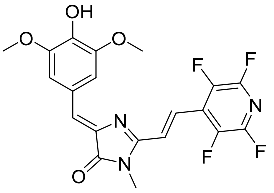


**5-((Z)-4-hydroxy-3,5-dimethoxybenzylidene)-2-((E)-2-(perfluoropyridin-4-yl)vinyl)-3-methyl-3,5-dihydro-4H-imidazol-4-one (N 980)**

Synthesized by the **method B**.

Red solid (135 mg, 31%); mp ~295 ºC with decomposition; $^{1}$H NMR (700 MHz, DMSO-*d*6) δ ppm 9.41 (br. s., 1 H), 7.91 (d, *J*=16.0 Hz, 1 H), 7.80 (br. s., 2 H), 7.47 (d, *J*=16.2 Hz, 1 H), 7.16 (s, 1 H), 3.85 (s, 6 H), 3.26 (s, 3 H); $^{13}$C NMR (201 MHz, DMSO-*d*6) δ ppm 169.2, 157.0, 147.8, 142.9 (dm, *J*=242.8 Hz), 139.8 (dm, *J*=263.7 Hz), 139.6, 138.9, 136.6, 129.1, 127.1, 125.6, 124.6, 121.1, 110.7, 55.6, 26.1; HRMS (ESI) m/z: 438.1076 found (calcd for $C_{20}H_{16}F_4N_3O_4^+$, [M+H]$^+$ 438.1071).

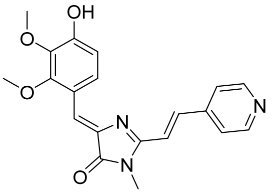


**5-((Z)-4-hydroxy-2,3-dimethoxybenzylidene)-3-methyl-2-((E)-2-(pyridin-4-yl)vinyl)-3,5-dihydro-4H-imidazol-4-one (N 1068)**

Synthesized by the **method B**.

Orange solid (223 mg, 61%); mp = 269-272 ºC; $^{1}$H NMR (700 MHz, DMSO-*d*6) δ ppm 11.66 (br. s., 1 H), 8.67 (d, *J*=5.7 Hz, 2 H), 8.10 (d, *J*=8.4 Hz, 1 H), 7.79 - 7.83 (m, 3 H), 7.51 (d, *J*=16.0 Hz, 1 H), 7.34 (s, 1 H), 6.71 (d, *J*=9.0 Hz, 1 H), 3.87 (s, 3 H), 3.72 (s, 3 H), 3.32 (s, 3 H); $^{13}$C NMR (75 MHz, DMSO-*d*6) δ ppm 168.7, 156.3, 156.3, 152.0, 150.4, 142.0, 136.9, 136.4, 134.6, 130.1, 124.5, 122.1, 118.5, 115.7, 104.6, 60.1, 55.9, 26.7; HRMS (ESI) m/z: 366.1439 found (calcd for $C_{20}H_{20}N_3O_4^+$, [M+H]$^+$ 366.1448).

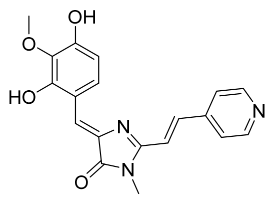

**5-((Z)-2,4-dihydroxy-3-methoxybenzylidene)-3-methyl-2-((E)-2-(pyridin-4-yl)vinyl)-3,5-dihydro-4H-imidazol-4-one (N 1069)**

Synthesized by the **method B**.

Orange solid (158 mg, 45%); mp =295-298 ºC; $^1$H NMR (700 MHz, DMSO-*d*6) δ ppm 8.68 (d, *J*=5.9 Hz, 2 H), 8.54 (br. s., 1 H), 7.76 - 7.81 (m, 3 H), 7.71 (d, J=8.4 Hz, 1 H), 7.52 (d, *J*=16.0 Hz, 1 H), 7.33 (s, 1 H), 6.65 (d, *J*=9.0 Hz, 1 H), 3.85 (s, 3 H), 3.32 (s, 3 H); $^{13}$C NMR (75 MHz, DMSO-*d*6) δ ppm 168.4, 155.6, 151.7, 150.4, 147.2, 142.0, 136.7, 134.3, 133.8, 126.1, 125.7, 122.0, 118.4, 115.3, 104.4, 55.9, 26.7; HRMS (ESI) m/z: 352.1286 found (calcd for $C_{19}H_{18}N_3O_4^+$, $[M+H]^+$ 352.1292).

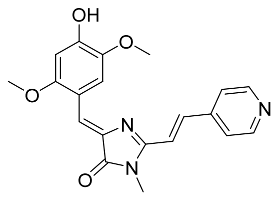

**5-((Z)-4-hydroxy-2,5-dimethoxybenzylidene)-3-methyl-2-((E)-2-(pyridin-4-yl)vinyl)-3,5-dihydro-4H-imidazol-4-one (N 1048)**

Synthesized by the **method B**.

Red solid (234 mg, 64%); mp ~250 ºC with decomposition; $^1$H NMR (700 MHz, DMSO-*d*6) δ ppm 10.11 (s, 1 H), 8.62 - 8.69 (m, 3 H), 7.84 (d, *J*=15.8 Hz, 1 H), 7.76 (d, *J*=5.9 Hz, 2 H), 7.49 (d, *J*=15.8 Hz, 1 H), 7.40 (s, 1 H), 6.56 (s, 1 H), 3.88 (s, 3 H), 3.82 (s, 3 H), 3.29 (s, 3 H); $^{13}$C NMR (75 MHz, DMSO-*d*6) δ ppm 169.8, 157.1, 155.6, 151.9, 150.3, 142.2, 142.2, 135.9, 135.7, 122.0, 120.4, 118.9, 115.1, 113.6, 99.7, 56.0, 55.9, 26.4; HRMS (ESI) m/z: 366.1440 found (calcd for $C_{20}H_{20}N_3O_4^+$, $[M+H]^+$ 366.1448).

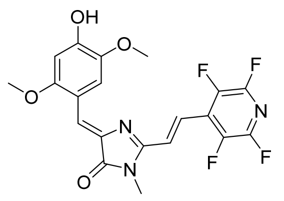

**5-((Z)-4-hydroxy-2,5-dimethoxybenzylidene)-3-methyl-2-((E)-2-(perfluoropyridin-4-yl)vinyl)-3,5-dihydro-4H-imidazol-4-one (N 1056)**

Synthesized by the **method B**.

Dark red solid (66 mg, 15%); mp ~290 ºC with decomposition; $^{1}$H NMR (700 MHz, DMSO-*d*$_6$) δ ppm 10.29 (br. s., 1 H), 8.69 (s, 1 H), 7.87 (d, *J*=16.2 Hz, 1 H), 7.43 - 7.49 (m, 2 H), 6.55 (s, 1 H), 3.85 (s, 3 H), 3.83 (s, 3 H), 3.25 (s, 3 H); $^{13}$C NMR (201 MHz, DMSO-*d*$_6$) δ ppm 169.2, 156.0, 155.9, 152.5, 142.9 (dm, *J*=242.1 Hz), 142.2, 139.5 (dm, *J*=262.3 Hz), 135.2, 127.1, 125.5, 121.8, 120.5, 114.7, 113.5, 99.7, 56.0, 55.2, 26.0; HRMS (ESI) m/z: 438.1069 found (calcd for $C_{20}H_{16}F_4N_3O_4^+$, $[M+H]^+$ 438.1071).

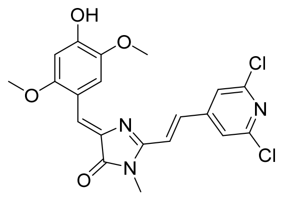

**5-((Z)-4-hydroxy-2,5-dimethoxybenzylidene)-3-methyl-2-((E)-2-(2,6-dichloropyridin-4-yl)vinyl)-3,5-dihydro-4H-imidazol-4-one (M 2767)**

Synthesized by the **method B**.

Dark red solid (229 mg, 53%); mp ~280 ºC with decomposition; $^{1}$H NMR (700 MHz, DMSO-*d*$_6$) δ ppm 10.18 (br. s., 1 H), 8.60 (s, 1 H), 8.04 (s, 2 H), 7.77 (d, *J*=15.8 Hz, 1 H), 7.64 (d, *J*=15.8 Hz, 1 H), 7.43 (s, 1 H), 6.55 (s, 1 H), 3.87 (s, 3 H), 3.82 (s, 3 H), 3.28 (s, 3 H); $^{13}$C NMR (176 MHz, DMSO-*d*$_6$) δ ppm 169.5, 156.4, 155.8, 152.2, 149.9, 149.0, 142.3, 135.6, 132.7, 121.8, 121.5, 121.2, 115.2, 113.5, 99.7, 56.1, 56.0, 26.4; HRMS (ESI) m/z: 432.0530 found (calcd for $C_{20}H_{16}Cl_2N_3O_4^-$, $[M-H]^-$ 432.0523).

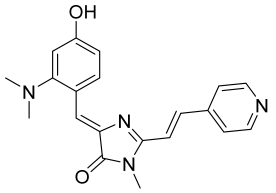

**5-((Z)-2-(dimethylamino)-4-hydroxybenzylidene)-3-methyl-2-((E)-2-(pyridin-4-yl)vinyl)-3,5-dihydro-4H-imidazol-4-one (MID 213)**

Synthesized by the **method B**.

Red solid (129 mg, 37%); mp = 211-214 °C ; $^1$H NMR (700 MHz, DMSO-$d_6$) □ ppm 10.13 (s, 1 H), 8.69 (d, *J*=8.7 Hz, 1 H), 8.65 (d, *J*=6.1 Hz, 2 H), 7.87 (d, *J*=15.8 Hz, 1 H), 7.81 (d, *J*=6.1 Hz, 2 H), 7.48 (d, *J*=15.8 Hz, 1 H), 7.34 (s, 1 H), 6.58 (dd, *J*=8.7, 2.4 Hz, 1 H), 6.52 (d, *J*=2.4 Hz, 1 H), 3.29 (s, 3 H), 2.72 (s, 6 H); $^{13}$C NMR (201 MHz, DMSO-$d_6$) δ ppm 169.9, 160.8, 157.6, 157.1, 149.9, 142.5, 135.8, 135.4, 135.0, 124.1, 121.8, 119.1, 118.3, 110.3, 105.3, 44.9, 26.3; HRMS (ESI) m/z: 349.1657 found (calcd for $C_{20}H_{21}N_4O_2$, $[M+H]^+$ 349.1659).

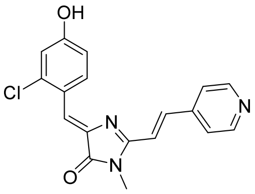

**5-((Z)-2-chloro-4-hydroxybenzylidene)-3-methyl-2-((E)-2-(pyridine-4-yl)vinyl)-3,5-dihydro-4H-imidazol-4-one (N 1204)**

Synthesized by the **method B**.

Orange solid (227 mg, 67%); mp = 284-287 ºC; $^1$H NMR (700 MHz, DMSO-$d_6$) δ ppm 10.70 (br. s., 1 H), 8.99 (d, *J*=8.8 Hz, 1 H), 8.67 (d, *J*=5.7 Hz, 2 H), 7.97 (d, *J*=15.8 Hz, 1 H), 7.82 (d, *J*=5.9 Hz, 2 H), 7.50 (d, *J*=15.8 Hz, 1 H), 7.30 (s, 1 H), 6.96 (d, *J*=2.3 Hz, 1 H), 6.93 (dd, *J*=8.8, 2.5 Hz, 1 H), 3.30 (s, 3 H); $^{13}$C NMR (176 MHz, DMSO-$d_6$) δ ppm 169.8, 160.3, 159.6, 150.2, 142.0, 138.1, 137.4, 136.6, 134.9, 122.7, 122.0, 120.3, 118.6, 116.3, 115.5, 26.5; HRMS (ESI) m/z: 340.0841 found (calcd for $C_{18}H_{15}ClN_3O_2^+$, $[M+H]^+$ 340.0847).

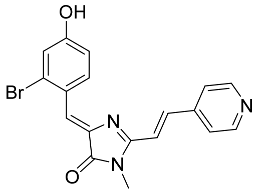


**5-((Z)-2-bromo-4-hydroxybenzylidene)-3-methyl-2-((E)-2-(pyridine-4-yl)vinyl)-3,5-dihydro-4H-imidazol-4-one (N 1205)**

Synthesized by the **method B**.

Red solid (218 mg, 57%); mp = 273-276 ºC; $^{1}$H NMR (700 MHz, DMSO-$d_6$) δ ppm 10.67 (br. s., 1 H), 8.99 (d, *J*=8.8 Hz, 1 H), 8.67 (d, *J*=5.9 Hz, 2 H), 7.97 (d, *J*=15.8 Hz, 1 H), 7.82 (d, *J*=6.1 Hz, 2 H), 7.50 (d, *J*=16.0 Hz, 1 H), 7.28 (s, 1 H), 7.14 (d, *J*=2.3 Hz, 1 H), 6.97 (dd, *J*=8.8, 2.5 Hz, 1 H), 3.30 (s, 3H); $^{13}$C NMR (176 MHz, DMSO-$d_6$) δ 169.8, 160.2, 159.7, 150.3, 142.0, 138.1, 137.4, 134.9, 127.7, 124.3, 123.2, 122.0, 119.7, 118.6, 115.9, 26.5; HRMS (ESI) m/z: 384.0344 found (calcd for $C_{18}H_{15}BrN_3O_2^+$, $[M+H]^+$ 384.0342).

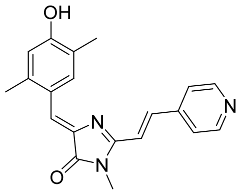


**5-((Z)-4-hydroxy-2,5-dimethylbenzylidene)-3-methyl-2-((E)-2-(pyridine-4-yl)vinyl)-3,5-dihydro-4H-imidazol-4-one (N 1206)**

Synthesized by the **method B**.

Red solid (273 mg, 82%); mp = 278-301 ºC; $^{1}$H NMR (700 MHz, DMSO-$d_6$) δ ppm 10.05 (br. s., 1 H), 8.75 (s, 1 H), 8.66 (d, *J*=5.9 Hz, 2 H), 7.89 (d, *J*=16.0 Hz, 1 H), 7.81 (d, *J*=6.1 Hz, 2 H), 7.49 (d, *J*=15.8 Hz, 1 H), 7.17 (s, 1 H), 6.70 (s, 1 H), 3.30 (s, 3 H), 2.37 (s, 3 H), 2.19 (s, 3 H); $^{13}$C NMR (176 MHz, DMSO-$d_6$) δ ppm 169.9, 158.2, 158.0, 150.2, 142.2, 139.6, 136.3, 136.3, 135.1, 123.8, 123.5, 122.5, 121.9, 118.9, 116.6, 26.5, 19.2, 15.9; HRMS (ESI) m/z: 334.1551 found (calcd for $C_{20}H_{20}N_3O_2^+$, $[M+H]^+$ 334.1550).

#### 10. $^{1}$H and $^{13}$C NMR spectra

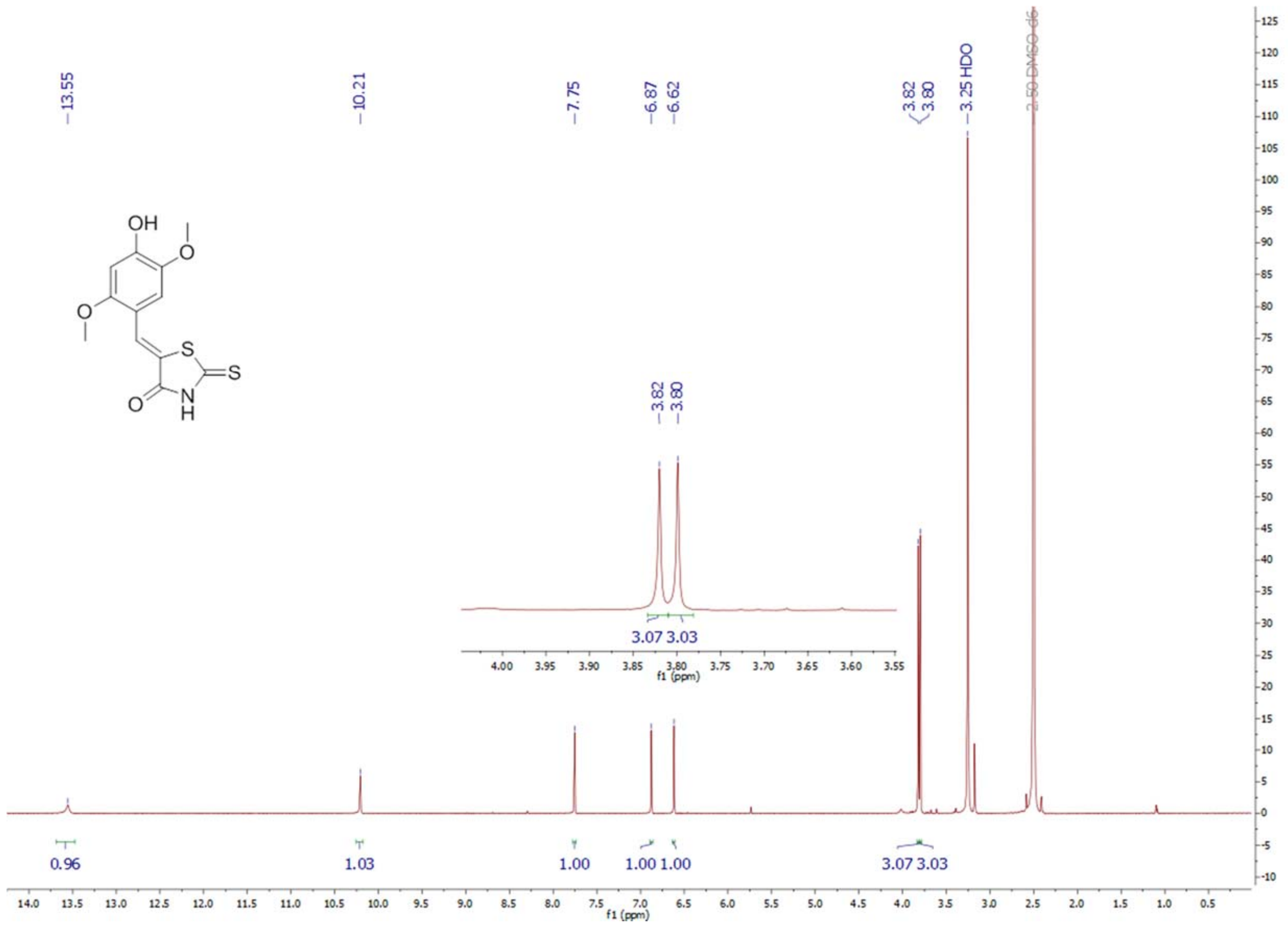

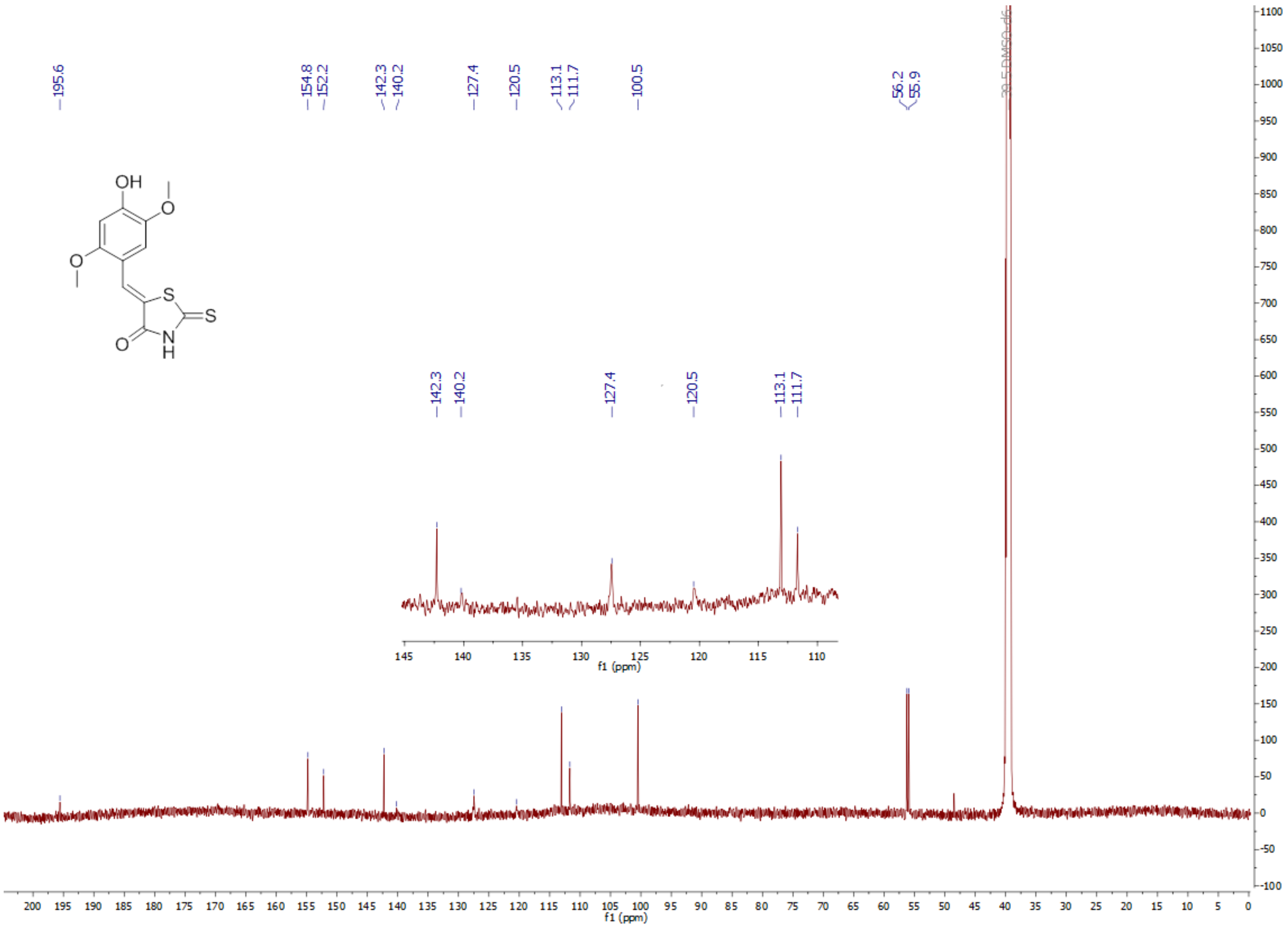

195.6
154.8
152.2
142.3
140.2
127.4
120.5
113.1
111.7
100.5
56.2
55.9
OH
O
O
S
S
N
H
O
f1 (ppm)

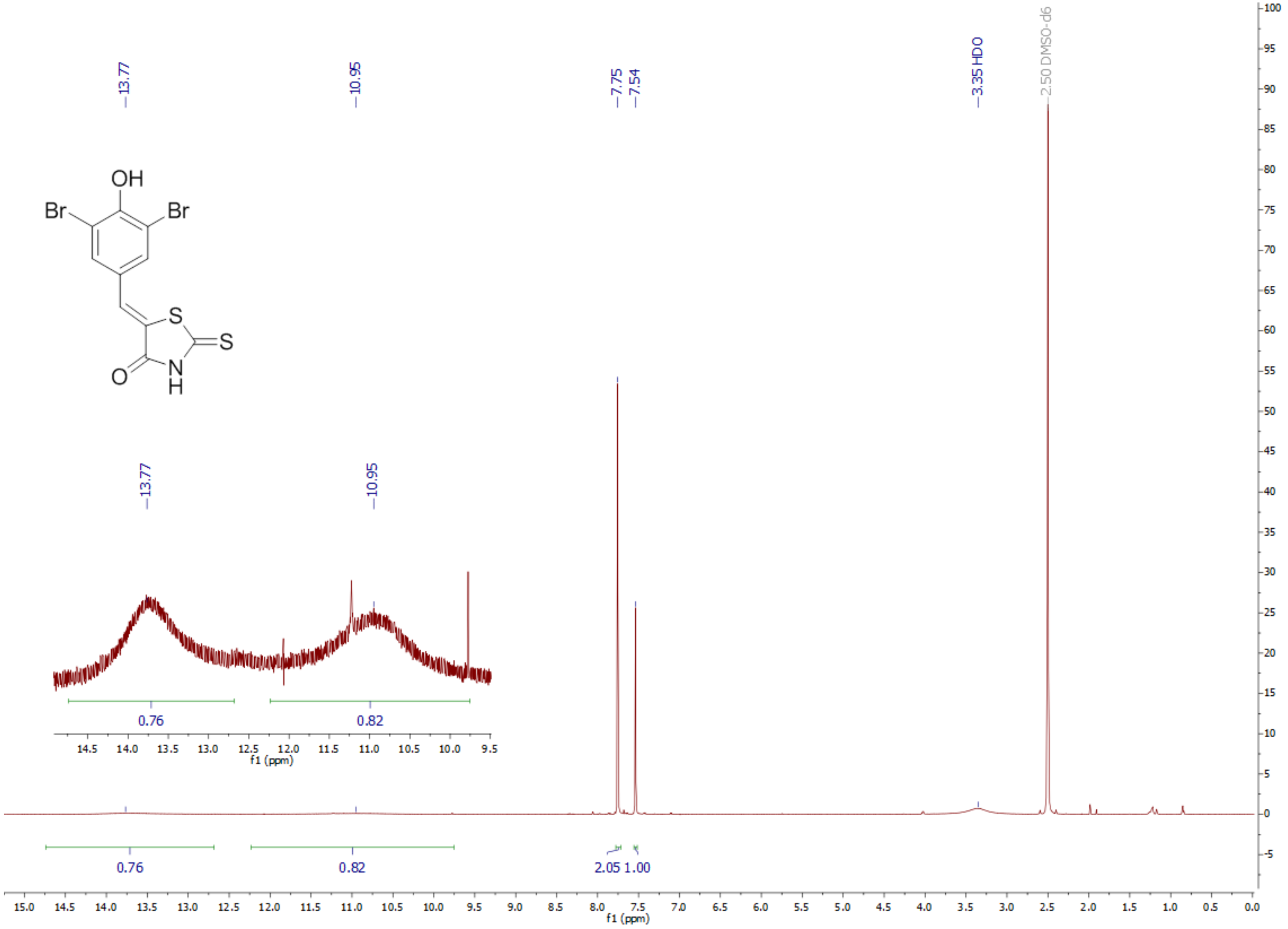

13.77
10.95
7.75
7.54
3.35 HDO
2.50 DMSO-d6
OH
Br
Br
S
S
O
N
H
13.77
10.95
0.76
0.82
f1 (ppm)
0.76
0.82
2.05 1.00
f1 (ppm)

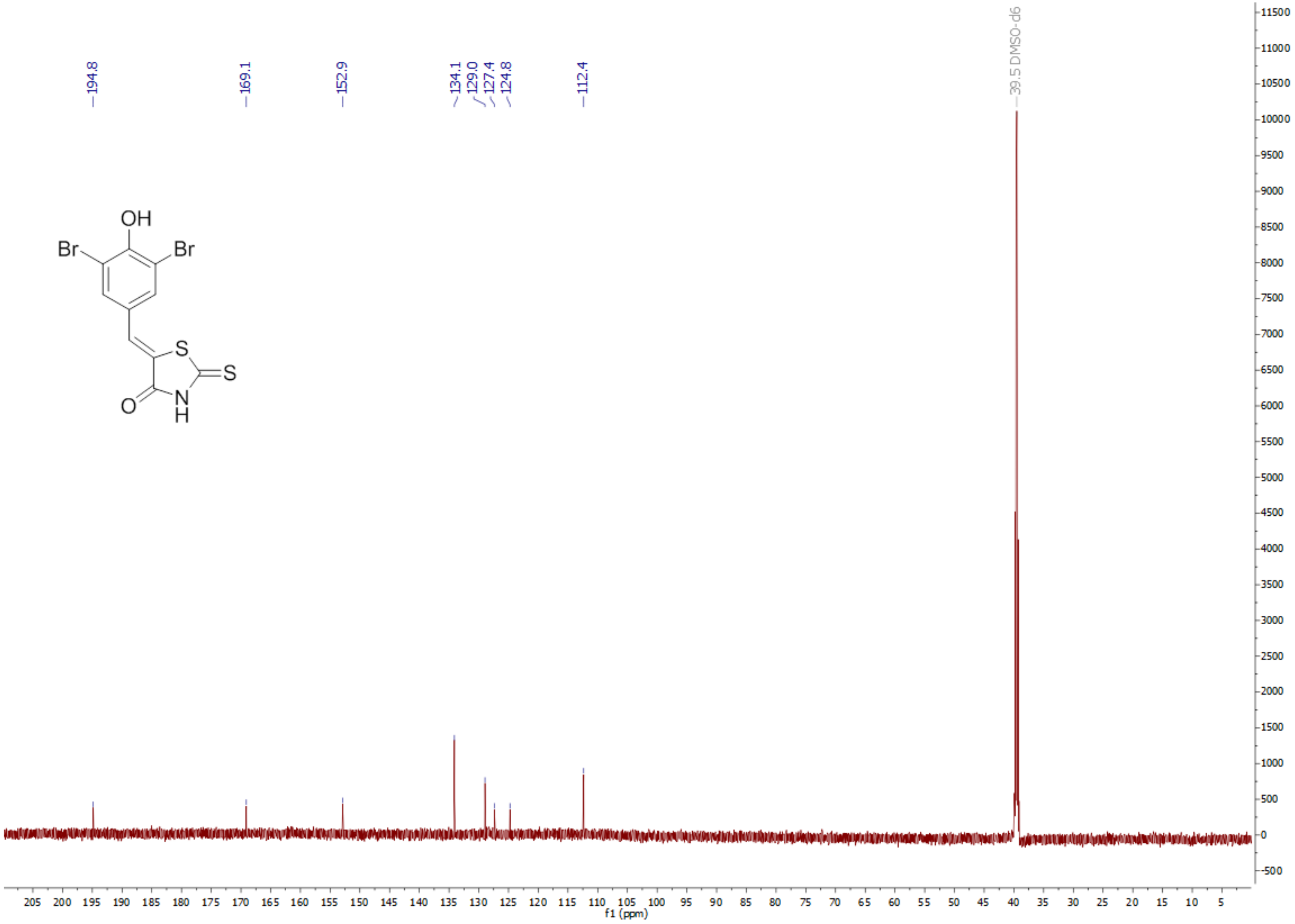
194.8
169.1
152.9
134.1
129.0
127.4
124.8
112.4
39.5 DMSO-d6
OH
Br
Br
S
S
O
N
H
f1 (ppm)

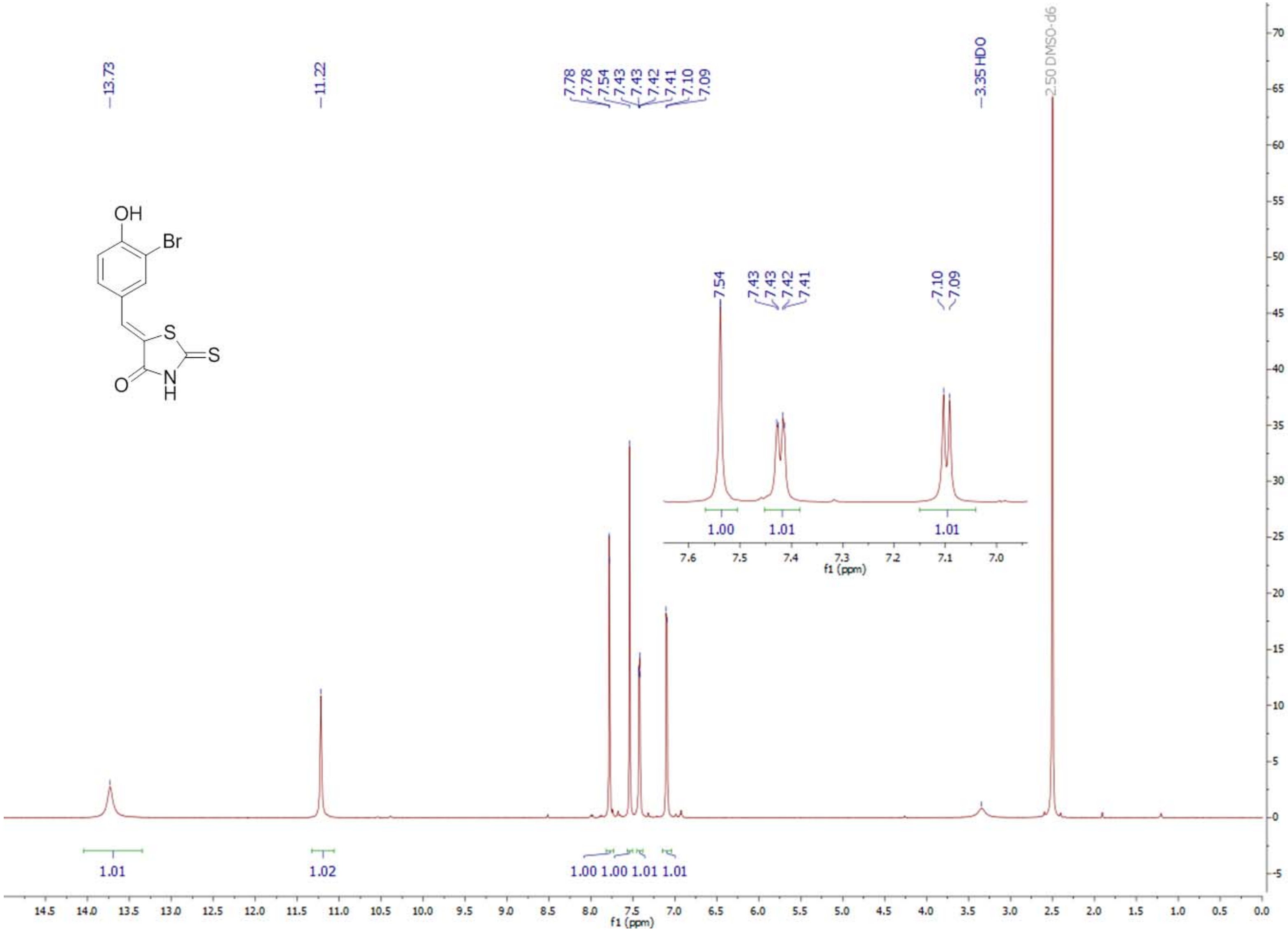

13.73
11.22
7.78
7.78
7.54
7.43
7.43
7.42
7.41
7.10
7.09
3.35 HDO
2.50 DMSO-d6
OH
Br
S
S
O
N
H
7.54
7.43
7.43
7.42
7.41
7.10
7.09
1.00
1.01
1.01
f1 (ppm)
1.01
1.02
1.00 1.00 1.01 1.01
f1 (ppm)

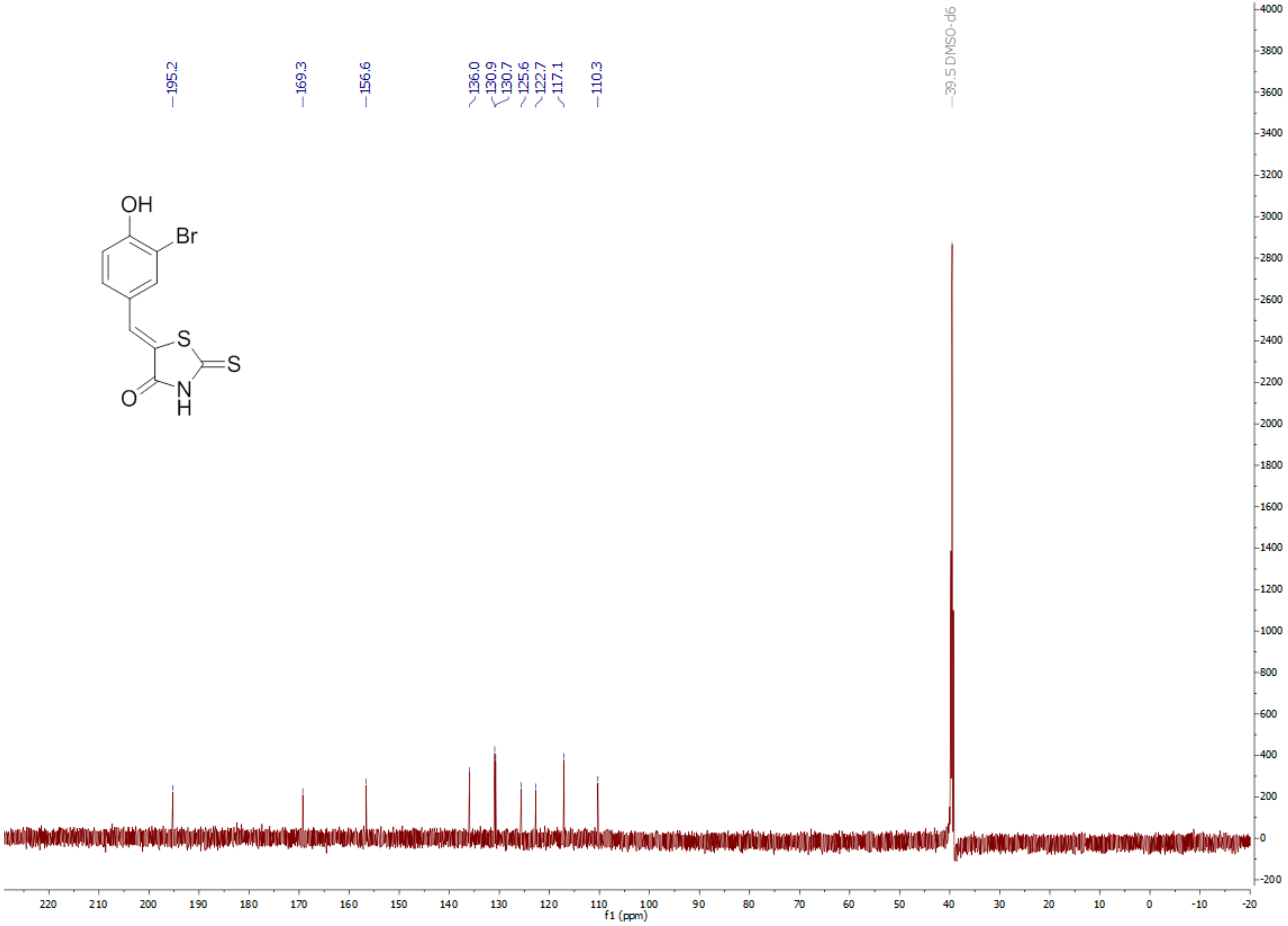
195.2
169.3
156.6
136.0
130.9
130.7
125.6
122.7
117.1
110.3
39.5 DMSO-d6
OH
Br
S
S
O
N
H
f1 (ppm)

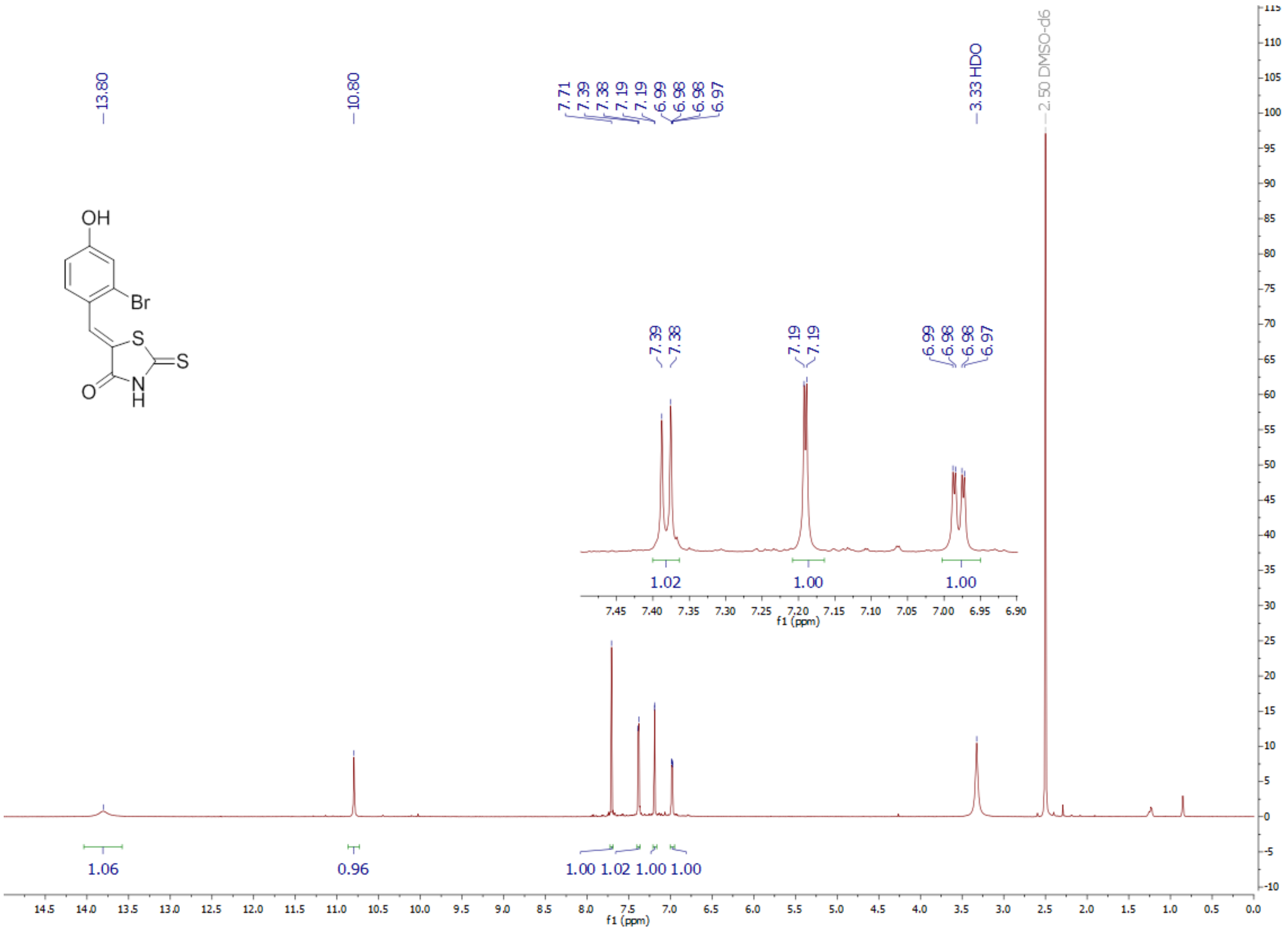

OH
Br
S
S
O
N
H
13.80
10.80
7.71
7.39
7.38
7.19
7.19
6.99
6.98
6.98
6.97
3.33 HDO
2.50 DMSO-d6
1.06
0.96
1.00 1.02 1.00 1.00
1.02
1.00
1.00
f1 (ppm)

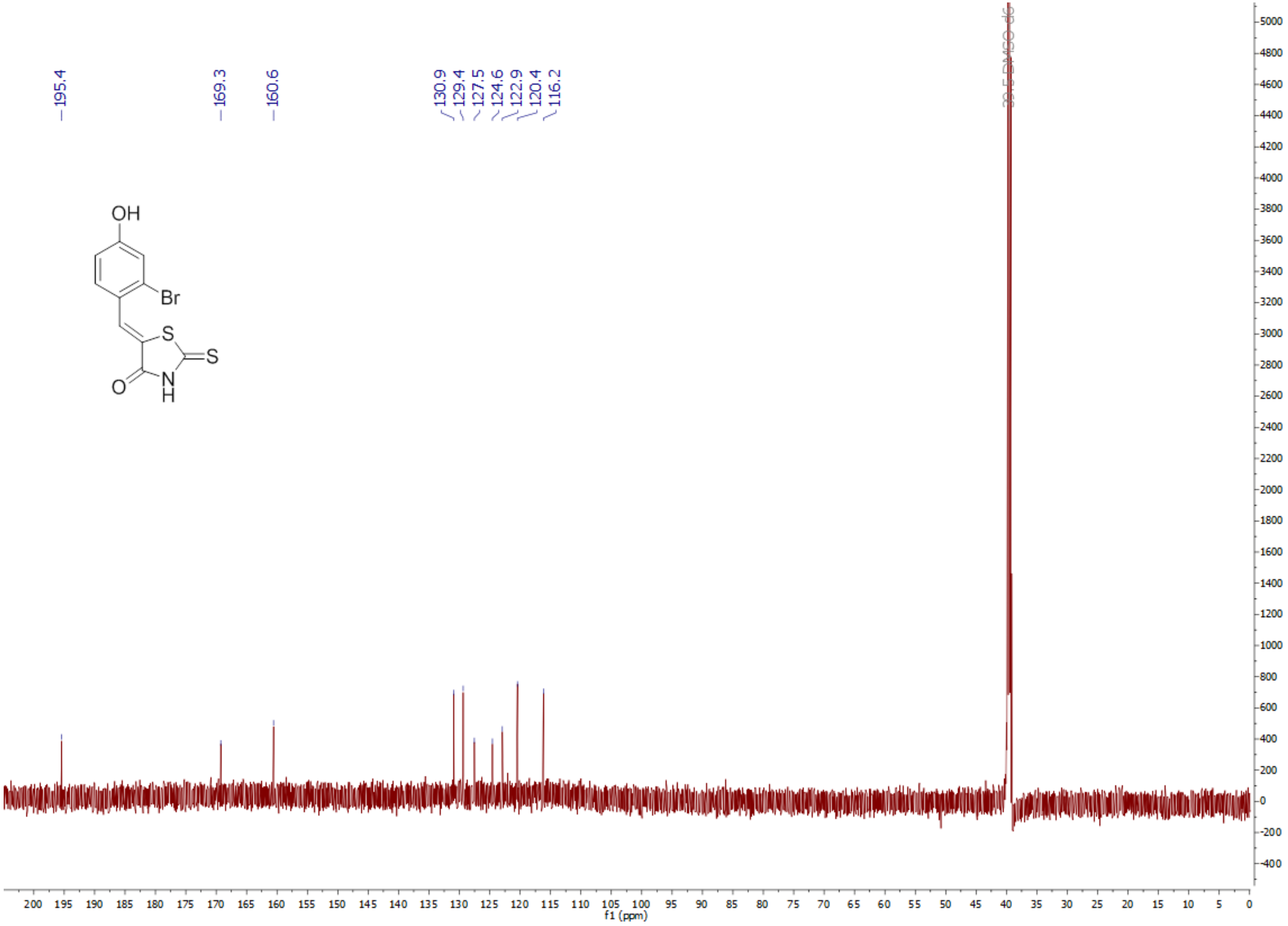

195.4
169.3
160.6
130.9
129.4
127.5
124.6
122.9
120.4
116.2
39.5 DMSO-d6
OH
Br
S
S
N
H
O
f1 (ppm)

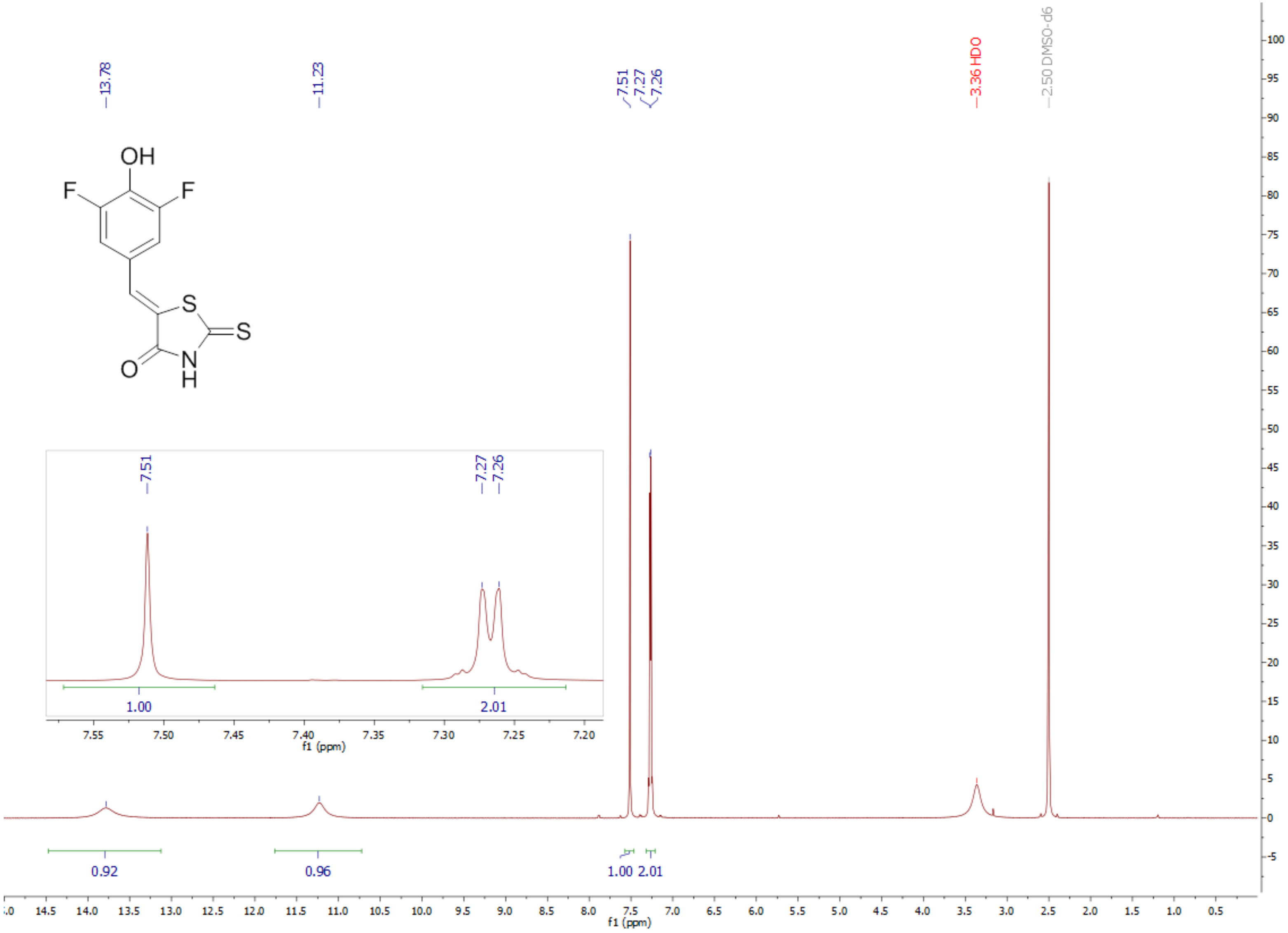

13.78
11.23
7.51
7.27
7.26
3.36 HDO
2.50 DMSO-d6
OH
F
F
S
S
O
N
H
1.00
2.01
f1 (ppm)
0.92
0.96
1.00 2.01
f1 (ppm)

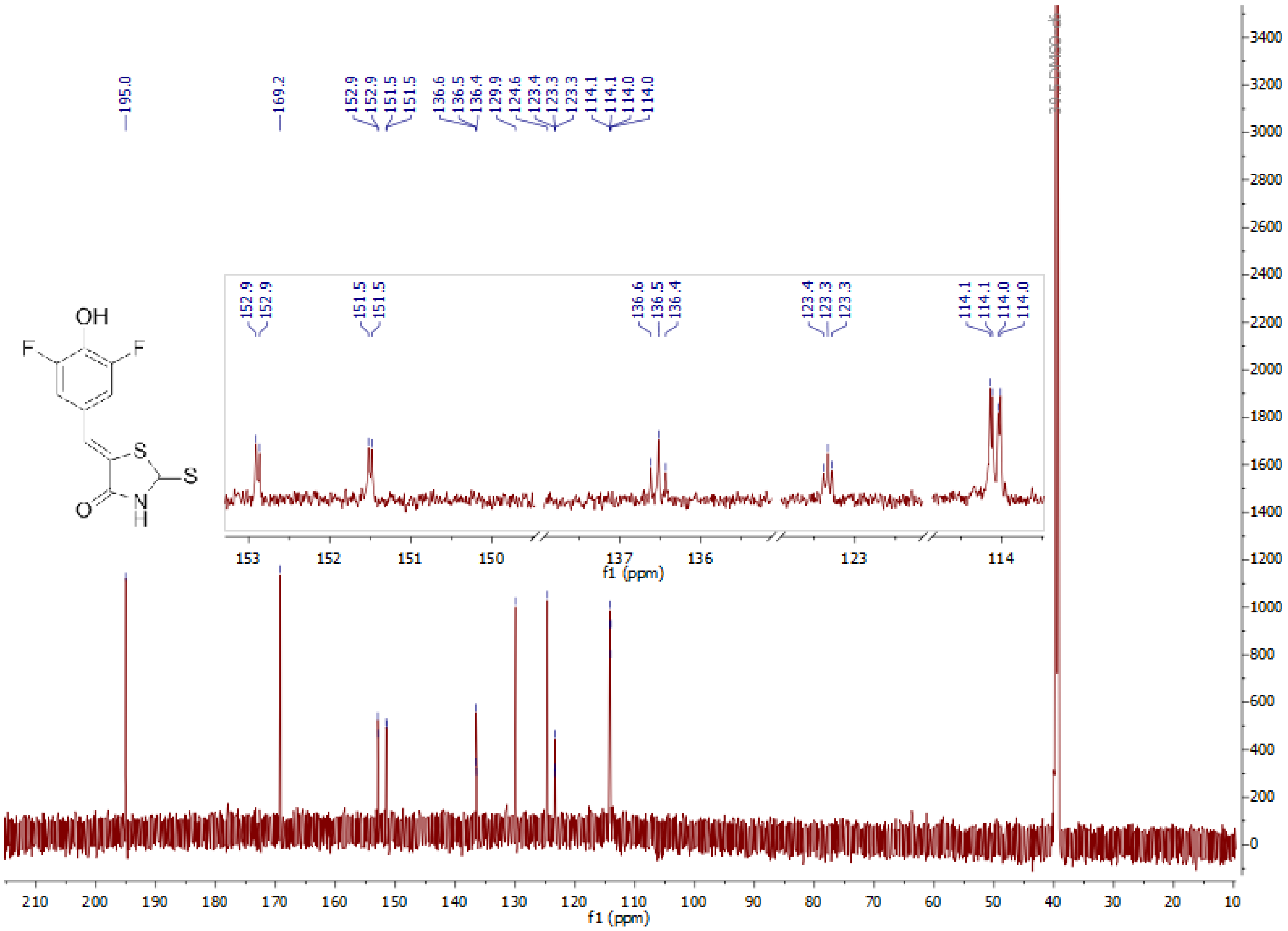

195.0
169.2
152.9
152.9
151.5
151.5
136.6
136.5
136.4
129.9
124.6
123.4
123.3
123.3
114.1
114.1
114.0
114.0
39.5 DMSO-d6
OH
F
F
S
S
O
N
H
f1 (ppm)

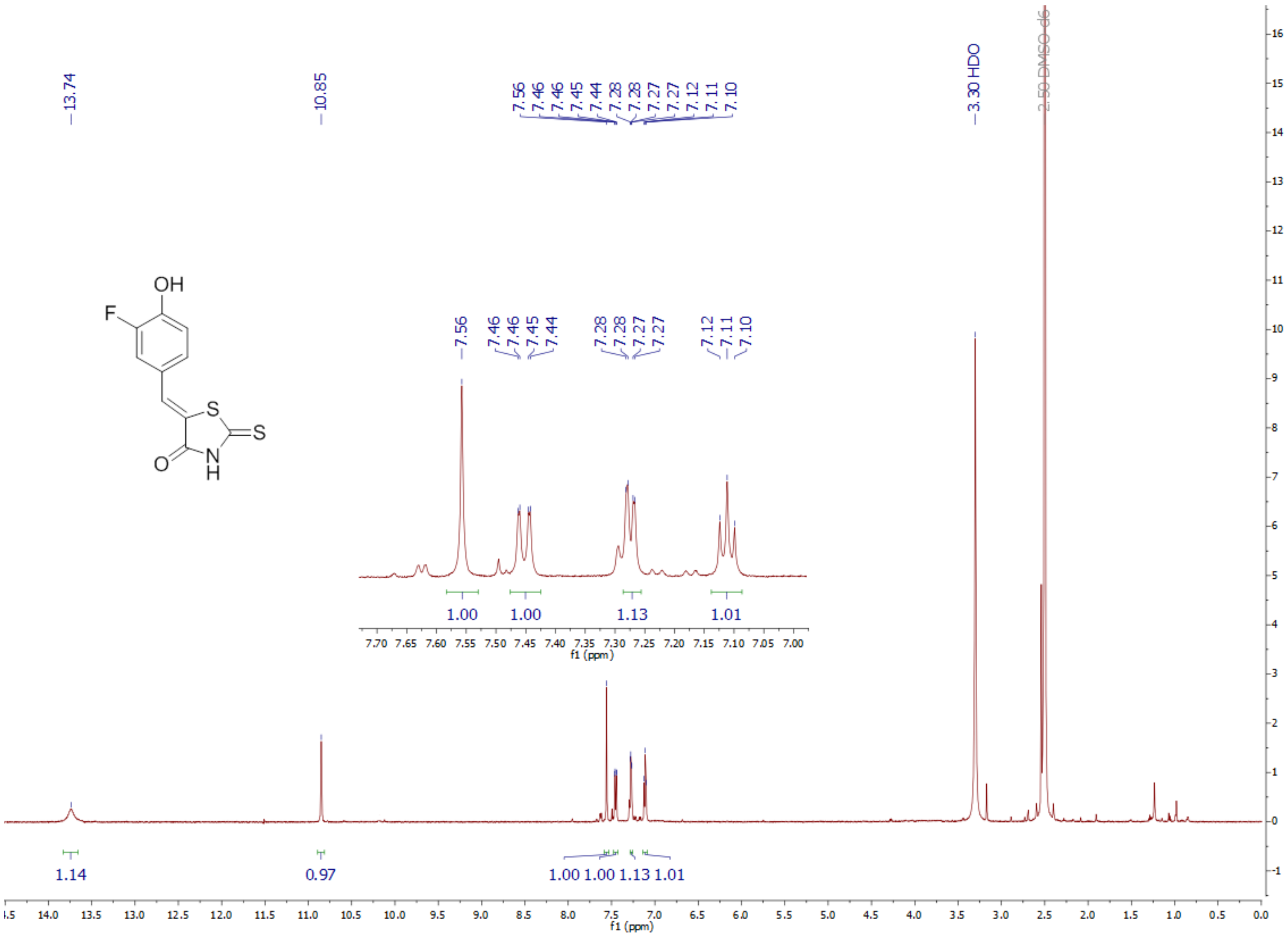

13.74
10.85
7.56
7.46
7.46
7.45
7.44
7.28
7.28
7.27
7.27
7.12
7.11
7.10
3.30 HDO
2.50 DMSO-d6
OH
F
S
S
O
N
H
1.00
1.00
1.13
1.01
f1 (ppm)
1.14
0.97
1.00 1.00 1.13 1.01
f1 (ppm)

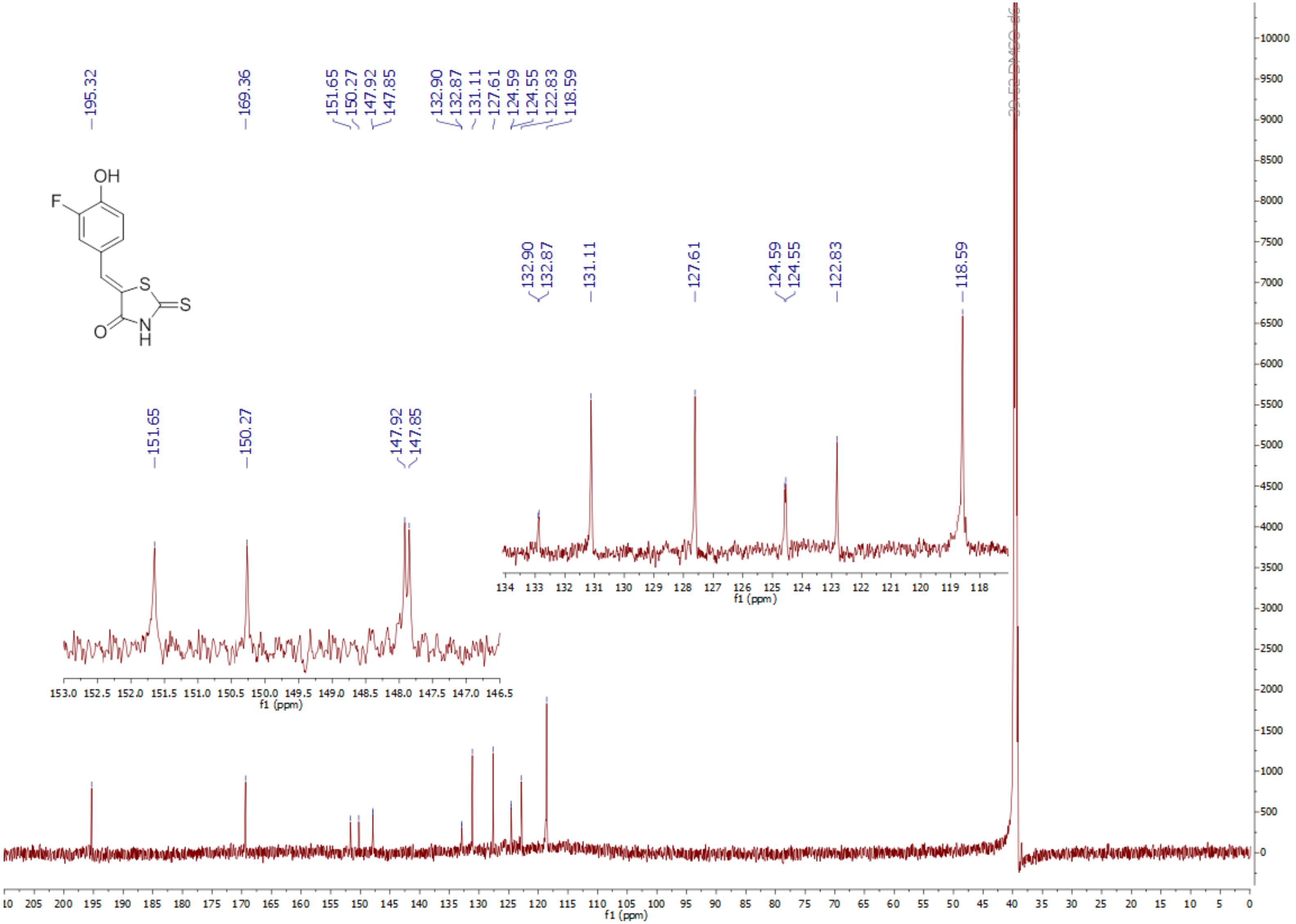
195.32
169.36
151.65
150.27
147.92
147.85
132.90
132.87
131.11
127.61
124.59
124.55
122.83
118.59
OH
F
S
S
N
H
O
f1 (ppm)
f1 (ppm)
f1 (ppm)

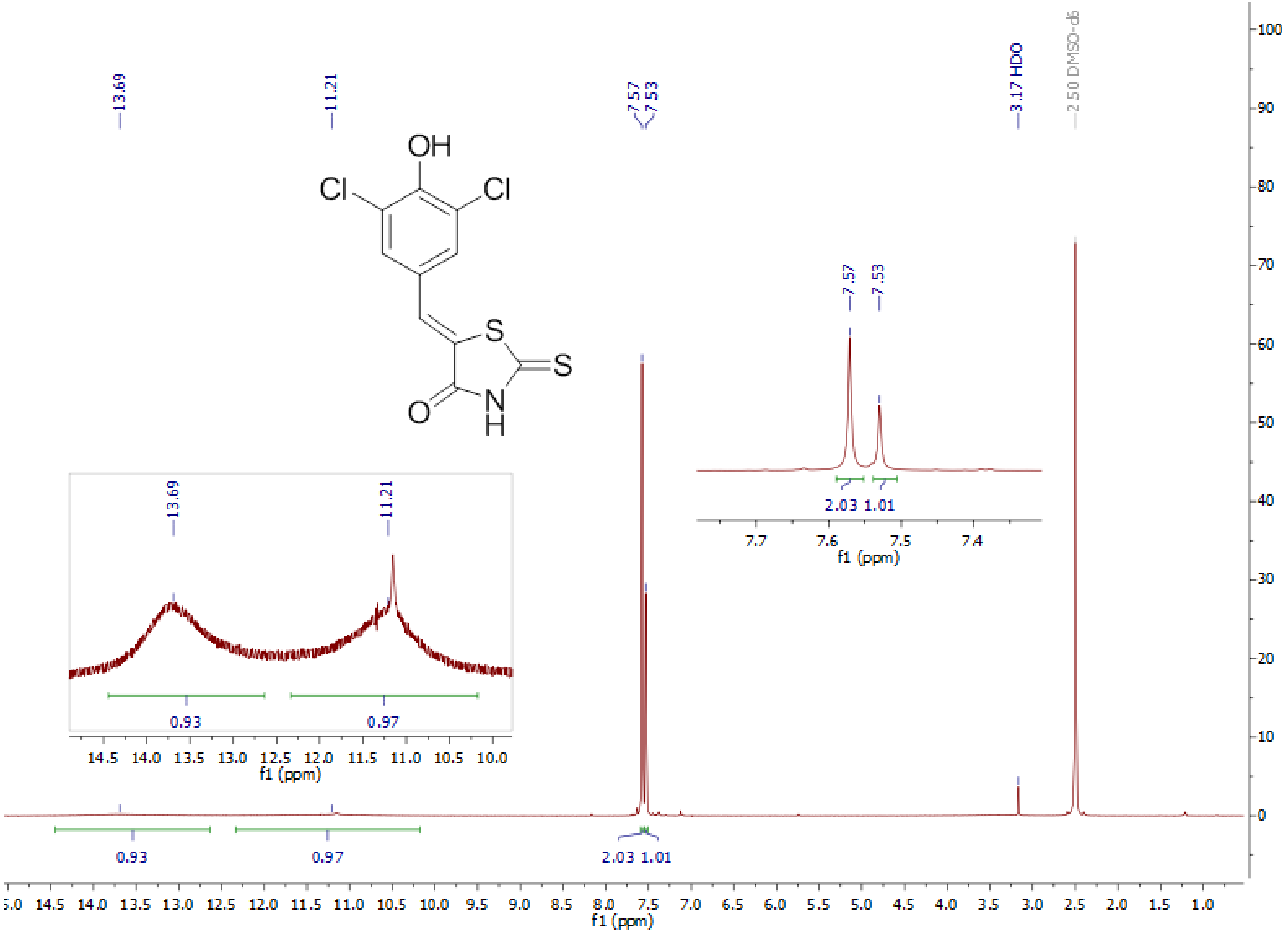

13.69
11.21
7.57
7.53
3.17 HDO
2.50 DMSO-d6
OH
Cl
Cl
S
S
N
H
O
0.93
0.97
2.03
1.01
f1 (ppm)

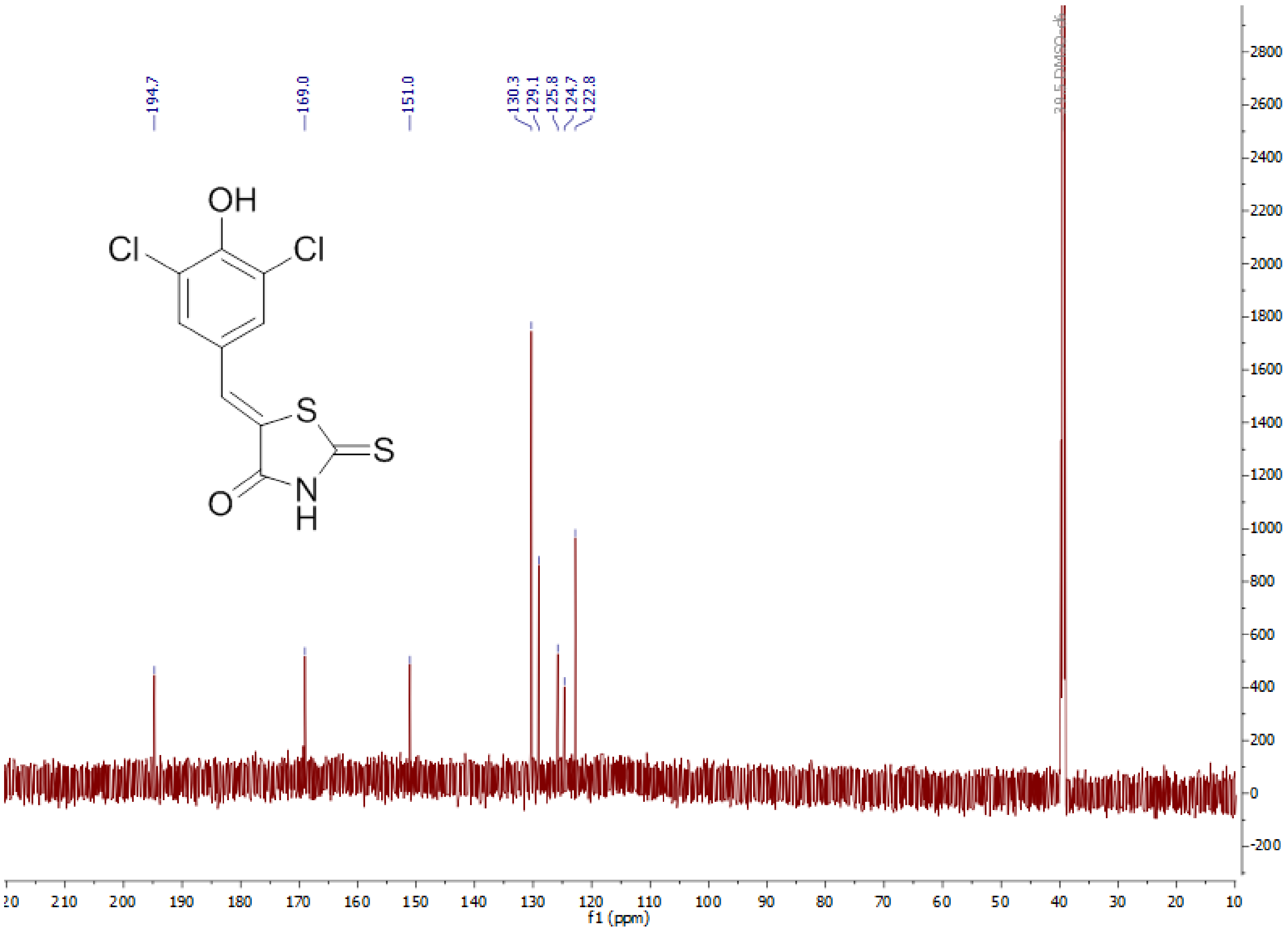
194.7
169.0
151.0
130.3
129.1
125.8
124.7
122.8
39.5 DMSO-d6
OH
Cl
Cl
S
S
O
N
H
f1 (ppm)

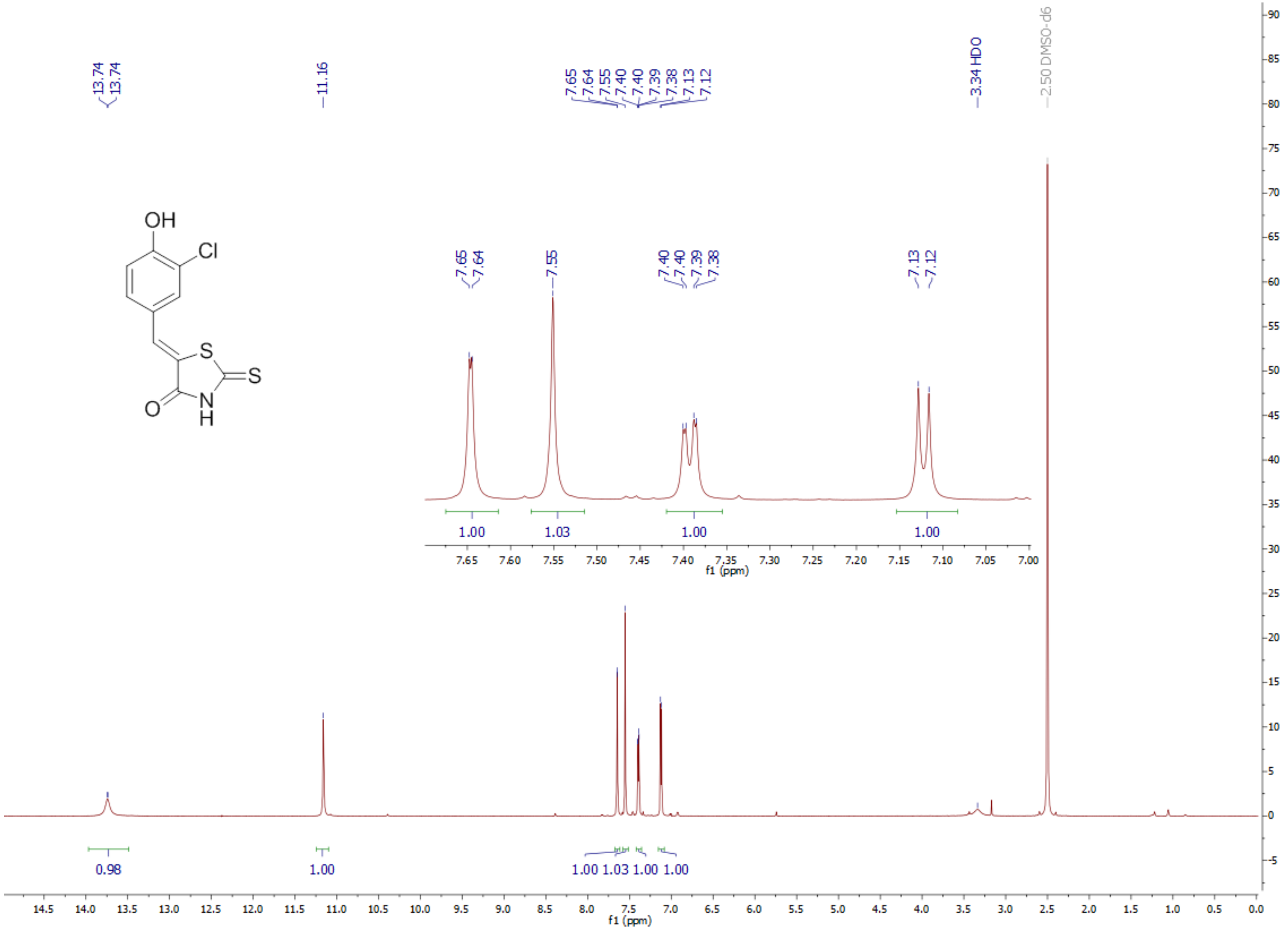
13.74
13.74
11.16
7.65
7.64
7.55
7.40
7.40
7.39
7.38
7.13
7.12
3.34 HDO
2.50 DMSO-d6
OH
Cl
S
S
N
H
O
0.98
1.00
1.00 1.03 1.00 1.00
f1 (ppm)

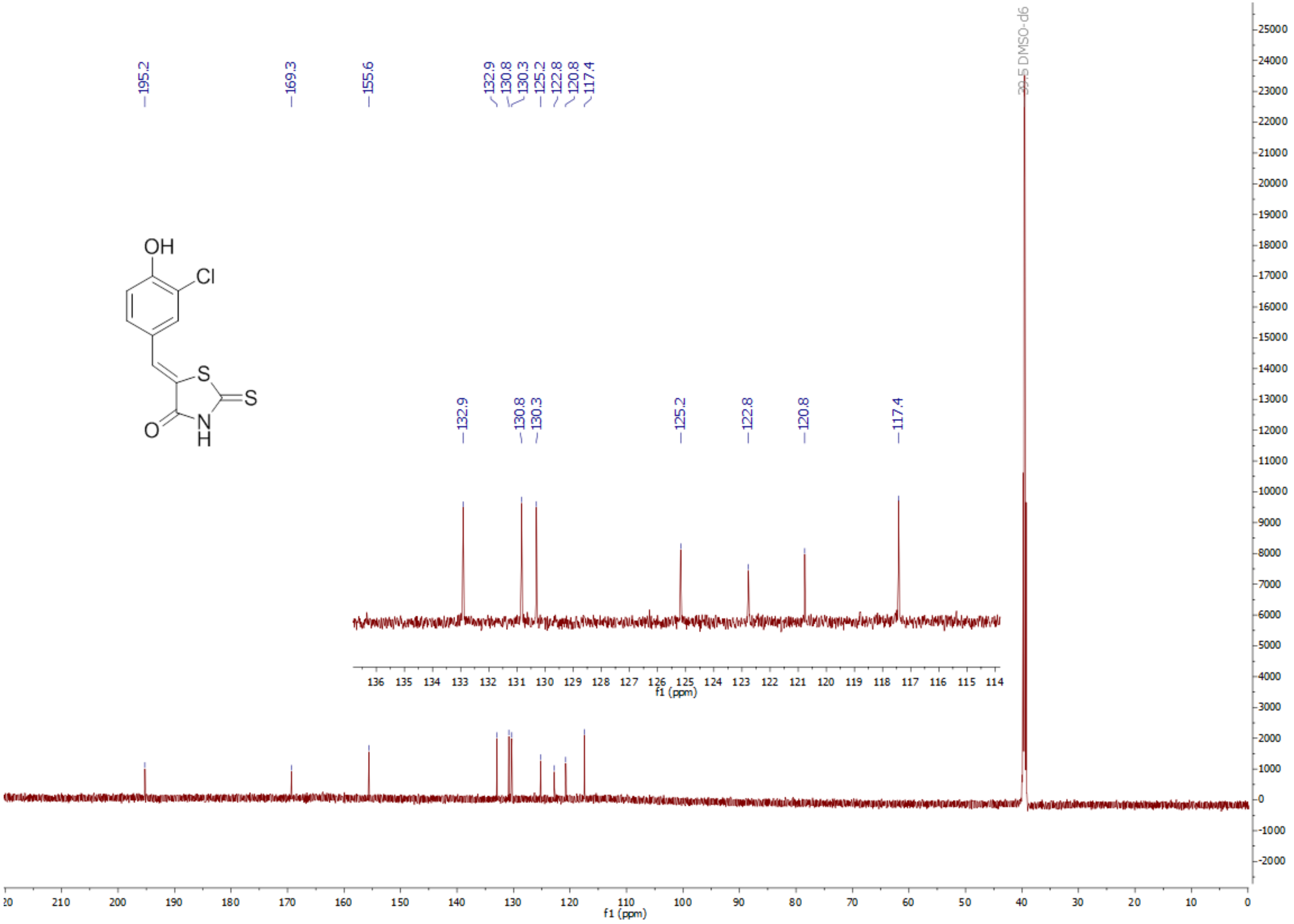
195.2
169.3
155.6
132.9
130.8
130.3
125.2
122.8
120.8
117.4
39.5 DMSO-d6
OH
Cl
S
S
O
N
H
f1 (ppm)

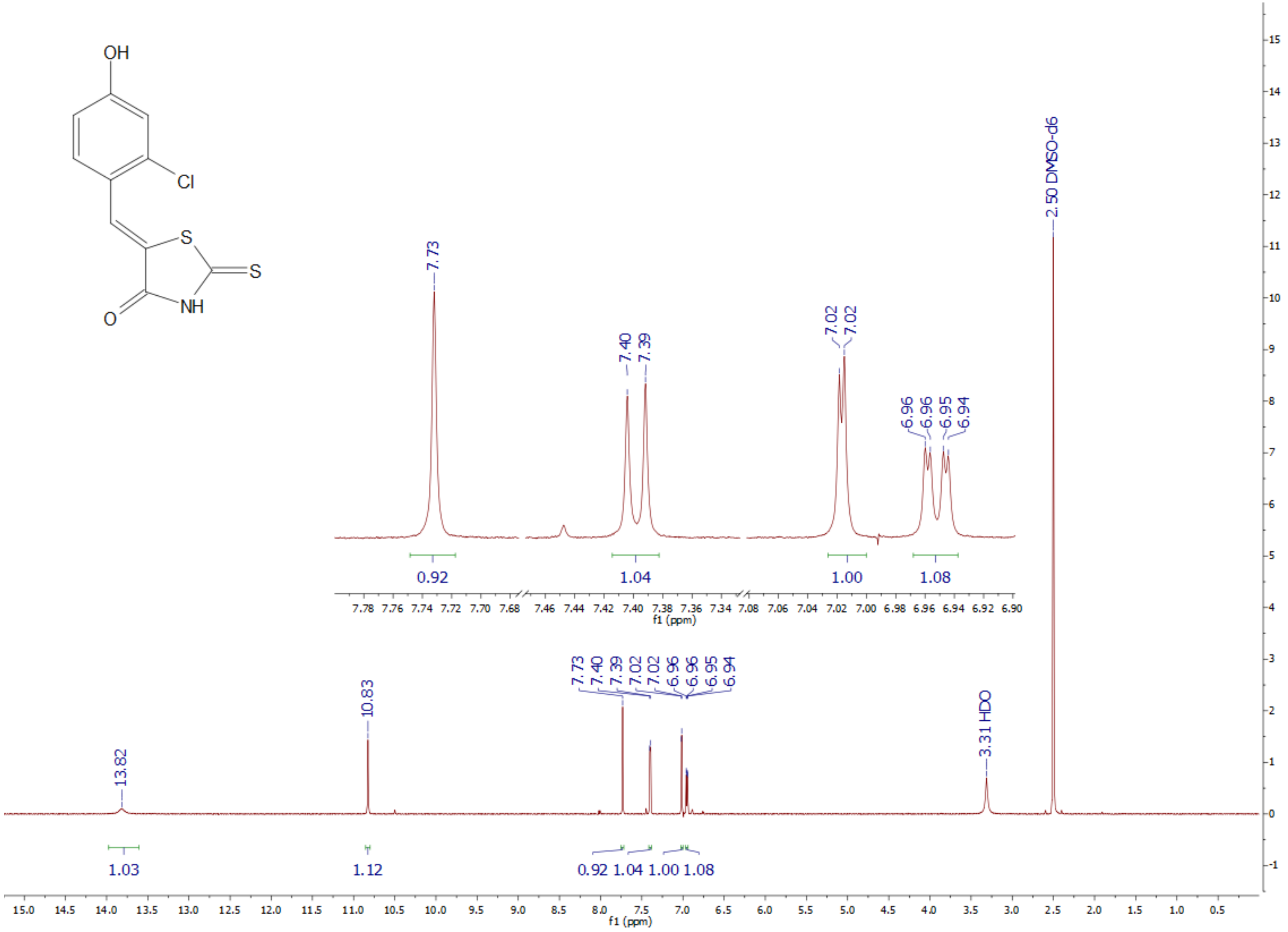
OH
Cl
S
S
NH
O
7.73
7.40
7.39
7.02
7.02
6.96
6.96
6.95
6.94
0.92
1.04
1.00
1.08
f1 (ppm)
2.50 DMSO-d6
13.82
10.83
3.31 HDO
1.03
1.12
0.92 1.04 1.00 1.08
f1 (ppm)

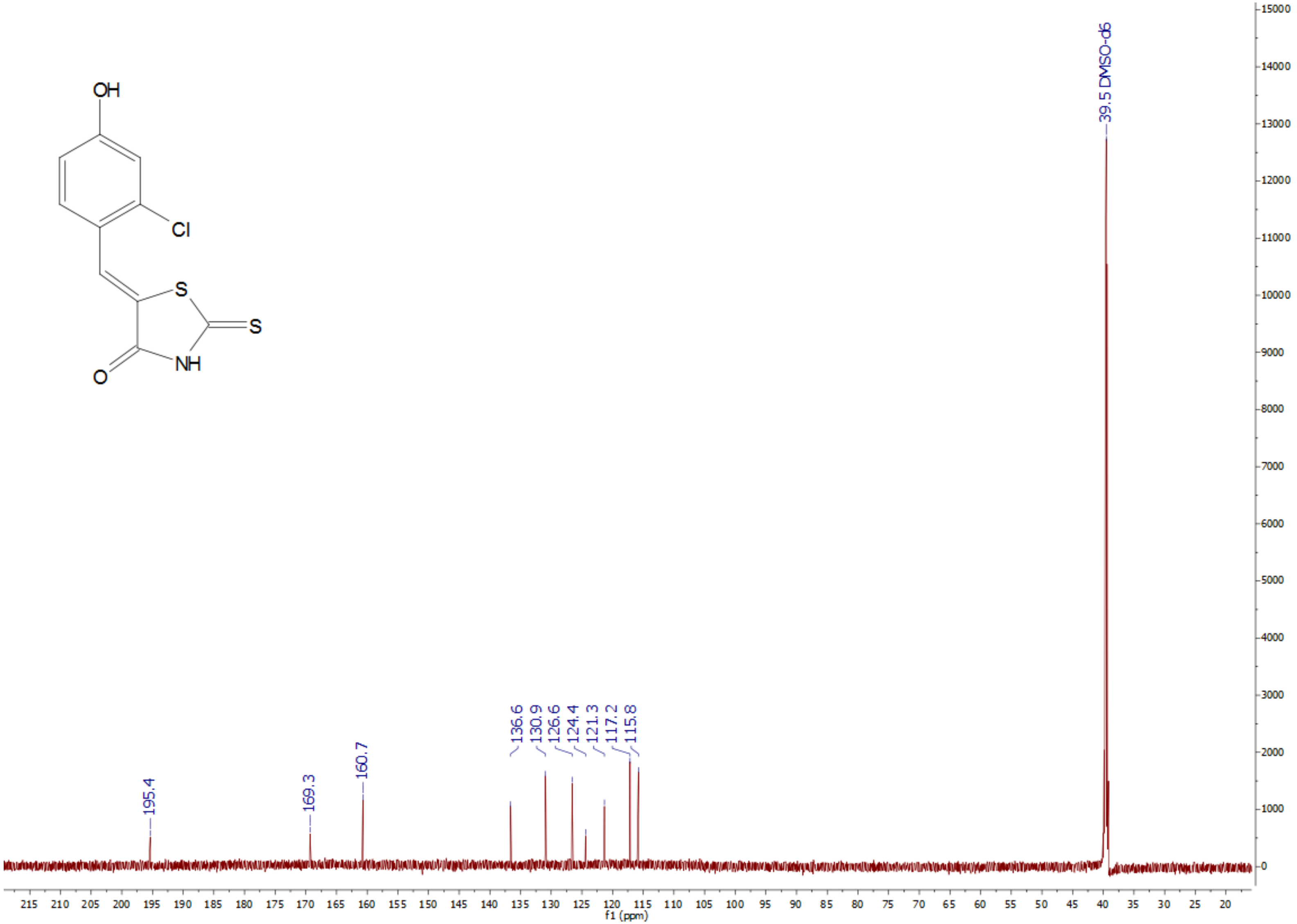
OH
Cl
S
S
NH
O
39.5 DMSO-d6
195.4
169.3
160.7
136.6
130.9
126.6
124.4
121.3
117.2
115.8
f1 (ppm)

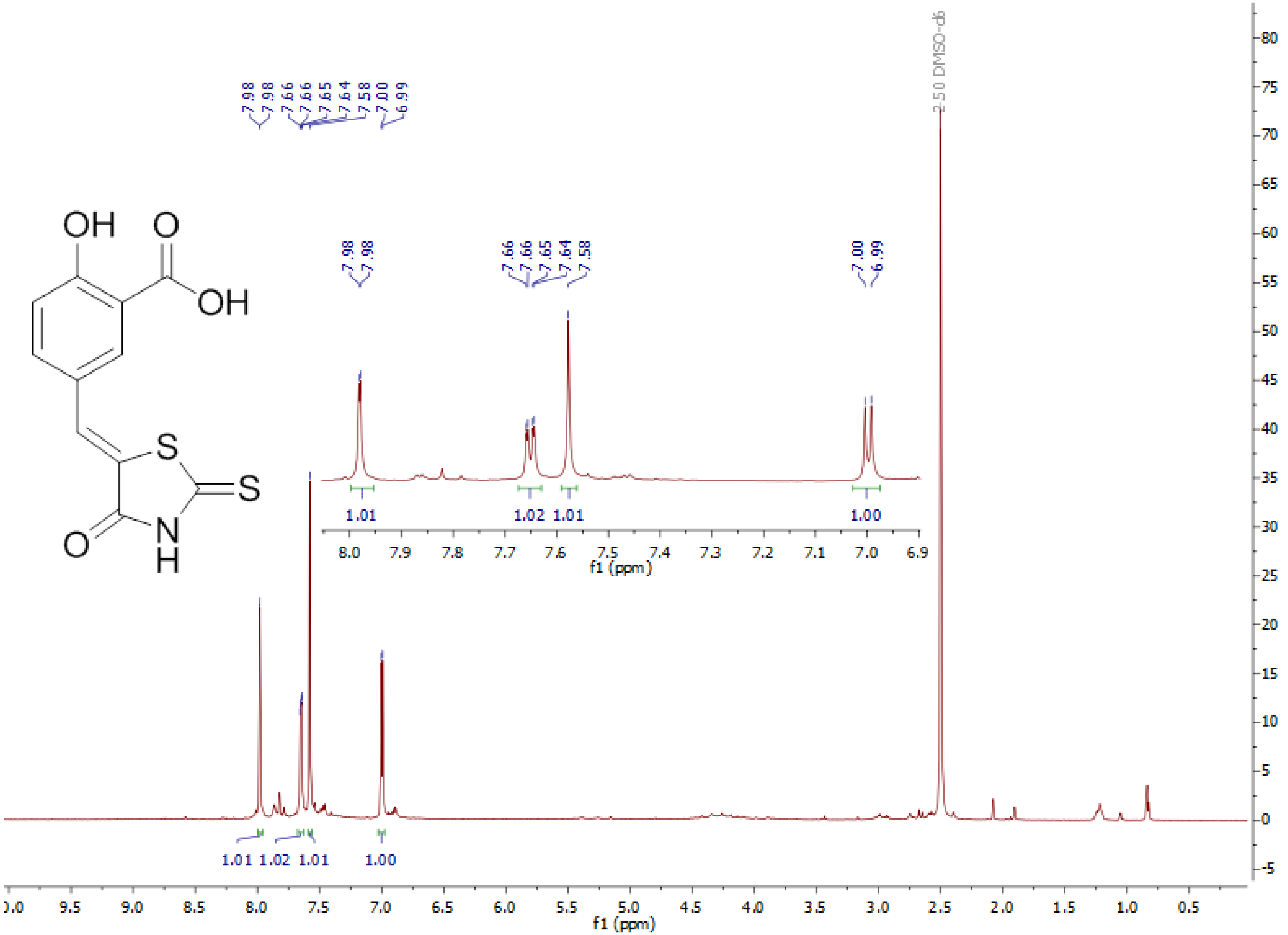

7.98
7.98
7.66
7.66
7.65
7.64
7.58
7.00
6.99
2.50 DMSO-d6
1.01
1.02
1.01
1.00
f1 (ppm)

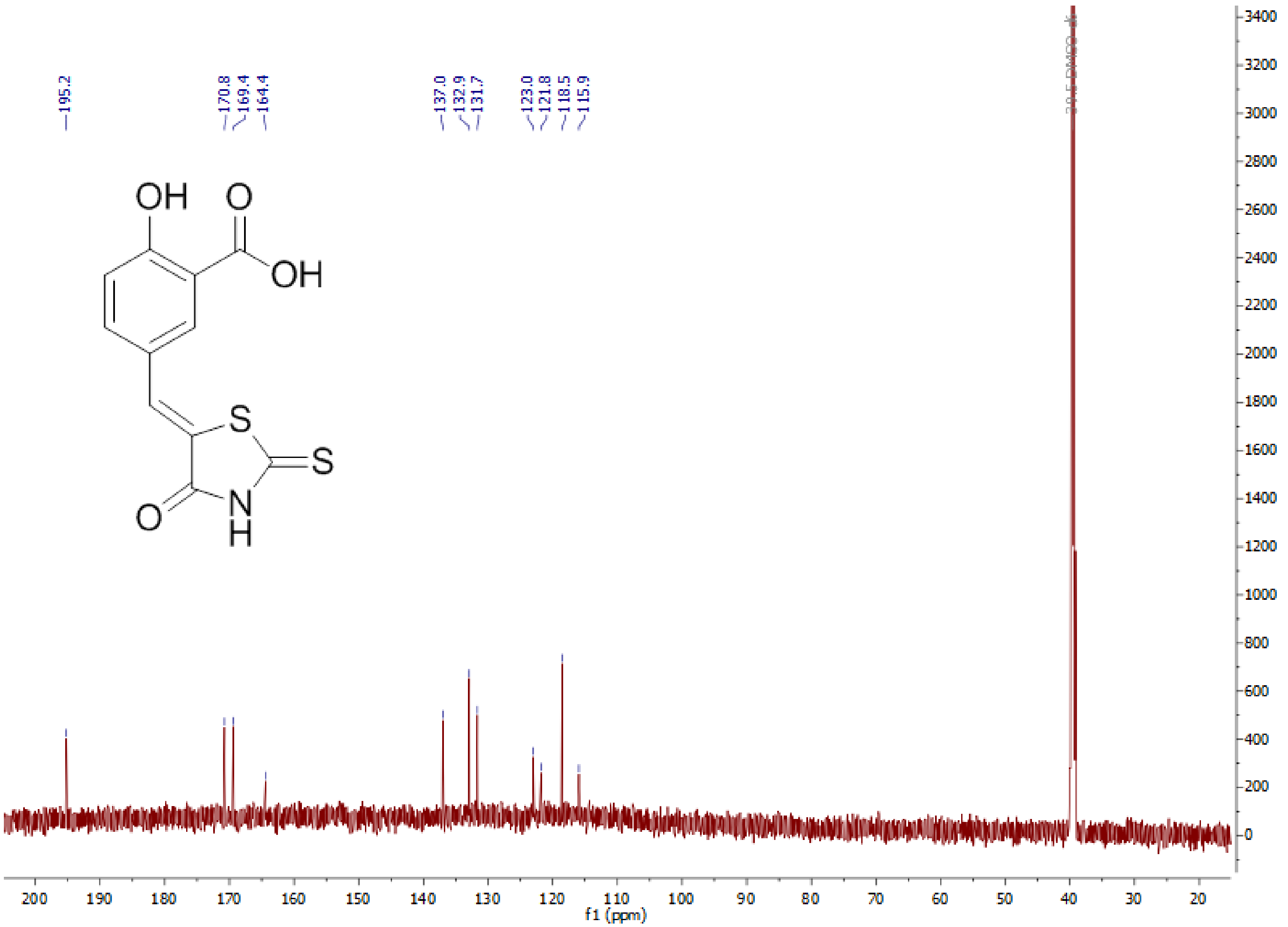
195.2
170.8
169.4
164.4
137.0
132.9
131.7
123.0
121.8
118.5
115.9
OH
O
OH
S
S
N
H
O
f1 (ppm)

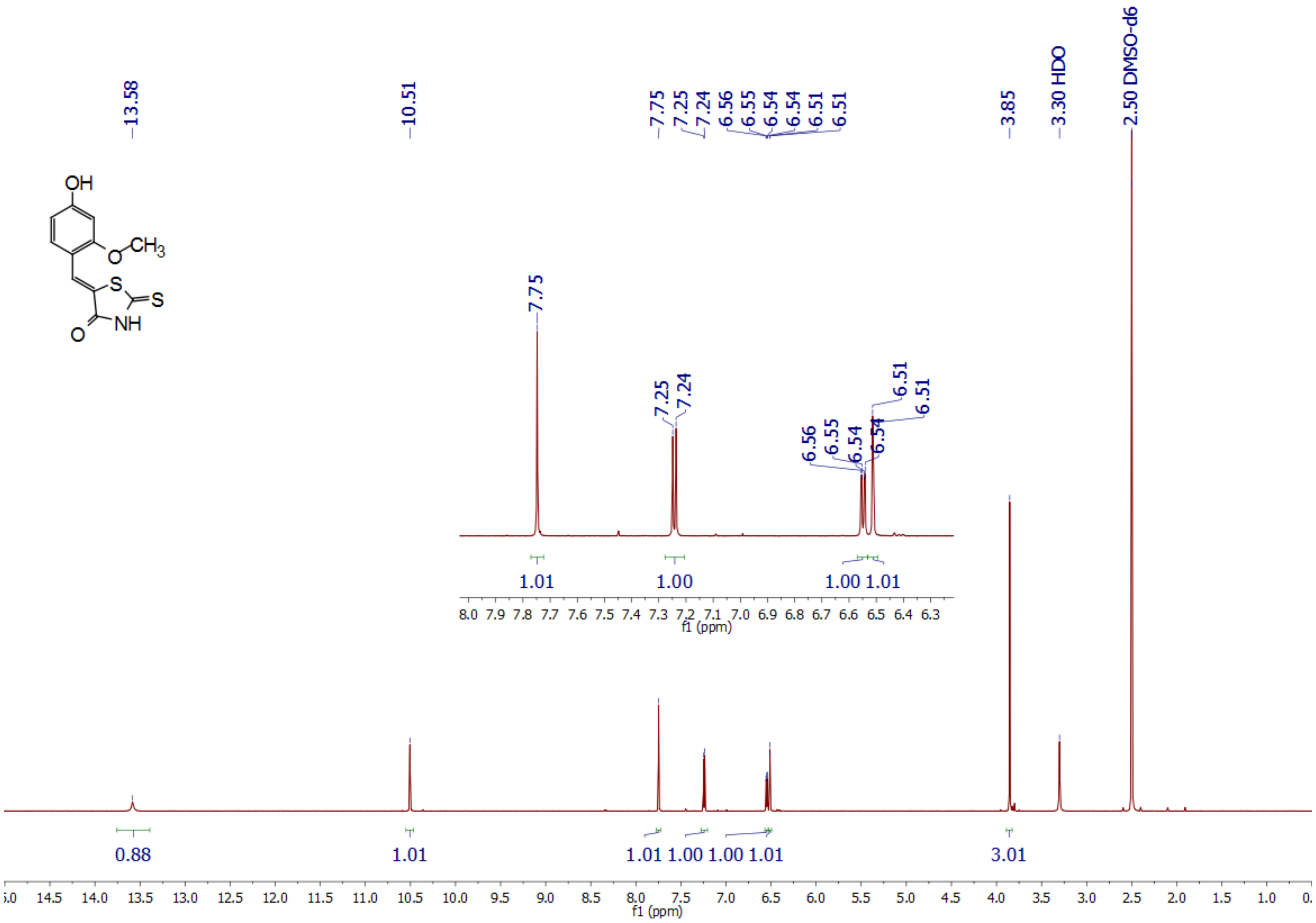
OH
O
CH3
S
S
NH
O
13.58
10.51
7.75
7.25
7.24
6.56
6.55
6.54
6.54
6.51
6.51
3.85
3.30 HDO
2.50 DMSO-d6
0.88
1.01
1.01 1.00 1.00 1.01
3.01
f1 (ppm)

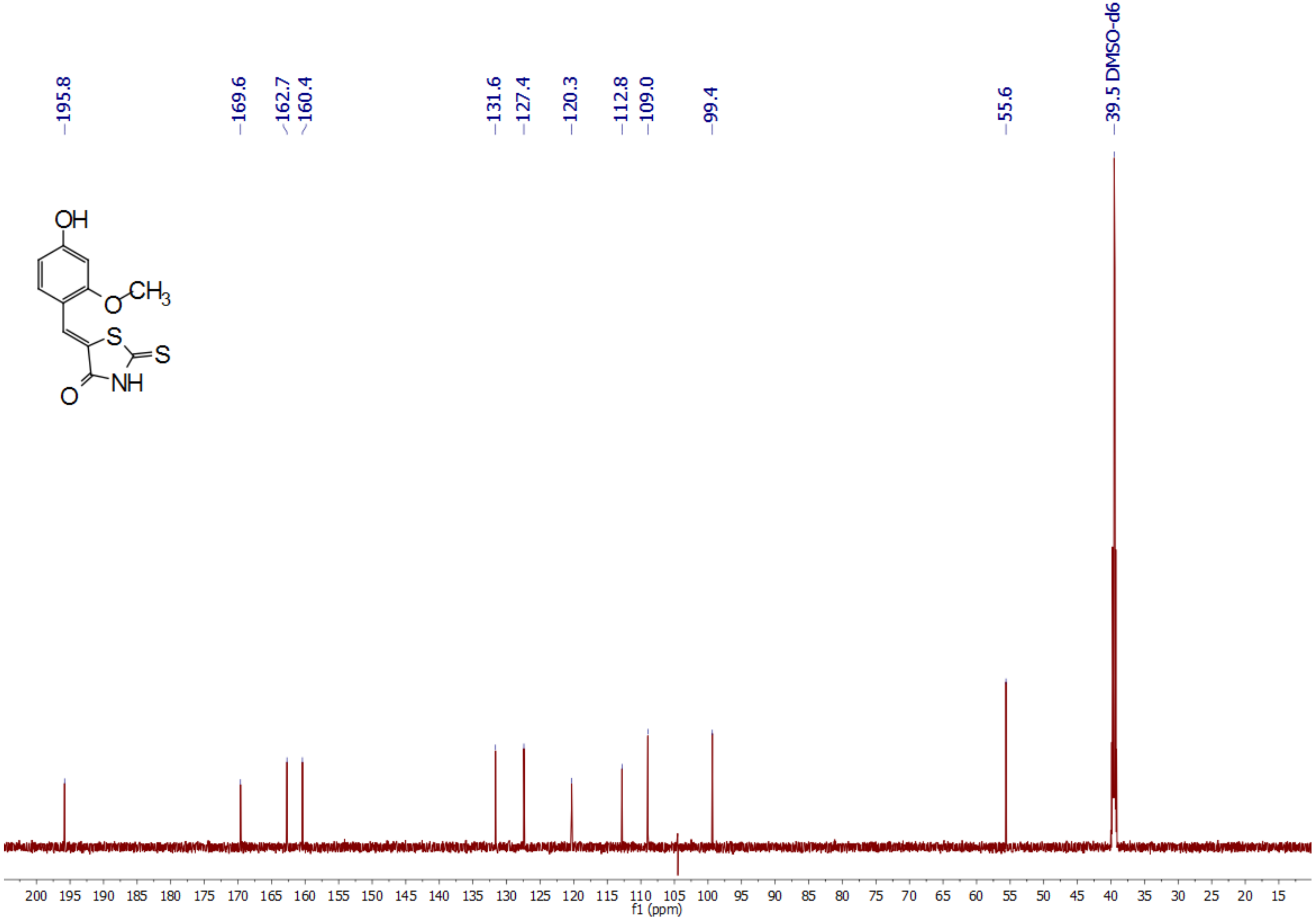
195.8
169.6
162.7
160.4
131.6
127.4
120.3
112.8
109.0
99.4
55.6
39.5 DMSO-d6
OH
CH3
O
S
S
NH
O
200 195 190 185 180 175 170 165 160 155 150 145 140 135 130 125 120 115 110 105 100 95 90 85 80 75 70 65 60 55 50 45 40 35 30 25 20 15
f1 (ppm)

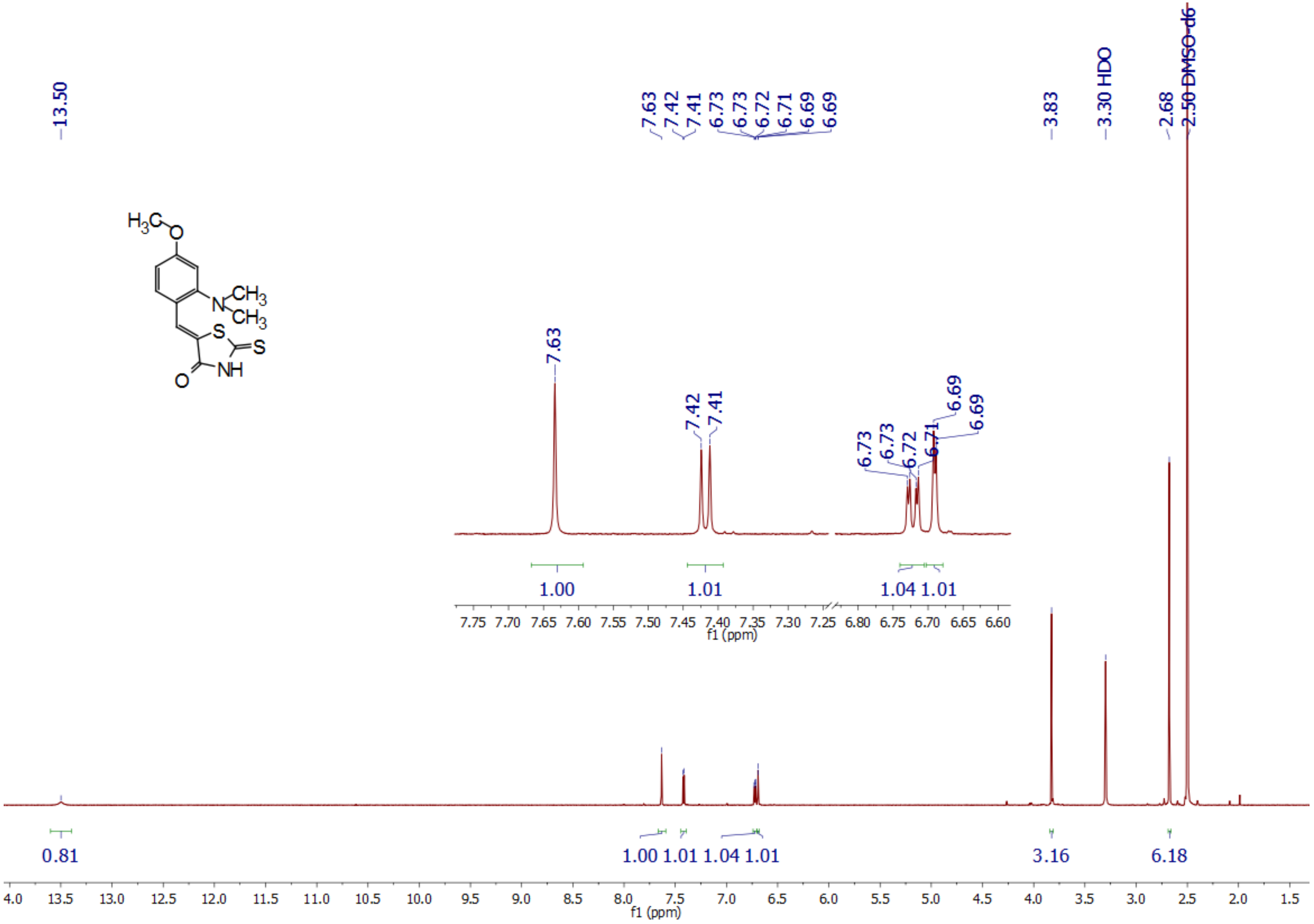
13.50
7.63
7.42
7.41
6.73
6.73
6.72
6.71
6.69
6.69
3.83
3.30 HDO
2.68
2.50 DMSO-d6
1.00
1.01
1.04 1.01
0.81
1.00 1.01 1.04 1.01
3.16
6.18
f1 (ppm)
H3C
O
N
CH3
CH3
S
S
NH
O

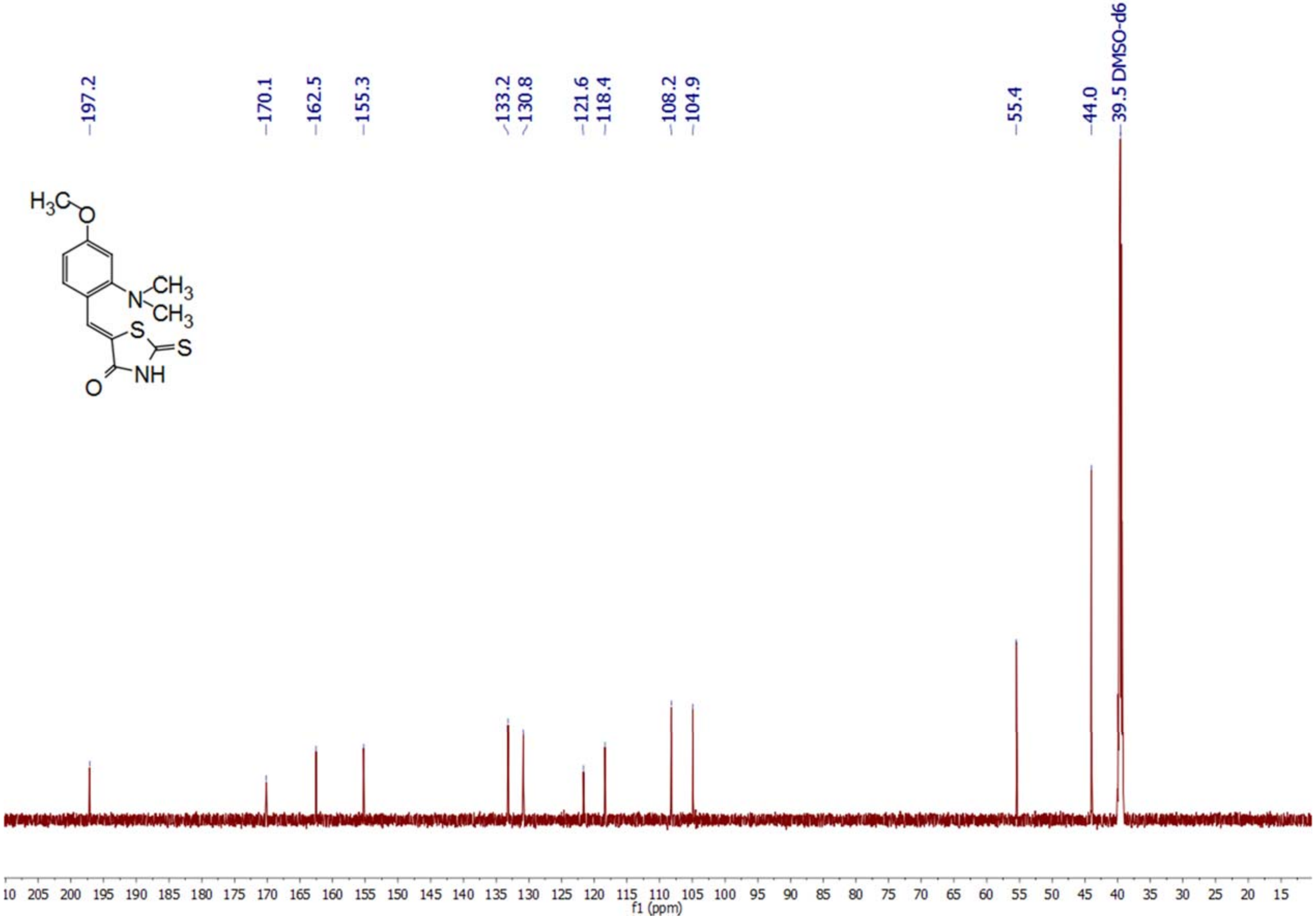

197.2
170.1
162.5
155.3
133.2
130.8
121.6
118.4
108.2
104.9
55.4
44.0
39.5 DMSO-d6
f1 (ppm)

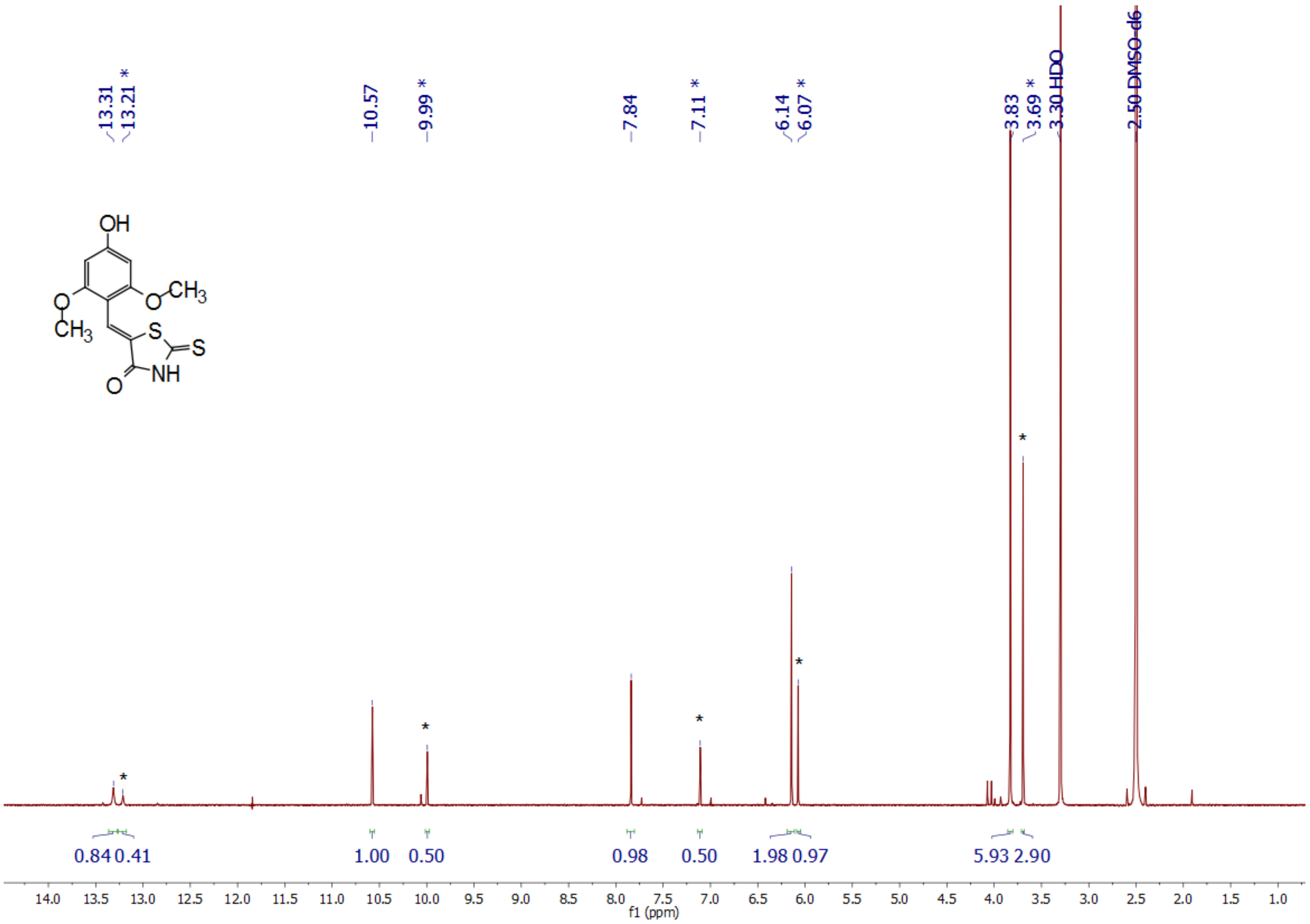

13.31
13.21 *
10.57
9.99 *
7.84
7.11 *
6.14
6.07 *
3.83
3.69 *
3.30 HDO
2.50 DMSO-d6
OH
O
CH3
S
S
NH
O
0.84
0.41
1.00
0.50
0.98
0.50
1.98
0.97
5.93
2.90
14.0 13.5 13.0 12.5 12.0 11.5 11.0 10.5 10.0 9.5 9.0 8.5 8.0 7.5 7.0 6.5 6.0 5.5 5.0 4.5 4.0 3.5 3.0 2.5 2.0 1.5 1.0
f1 (ppm)

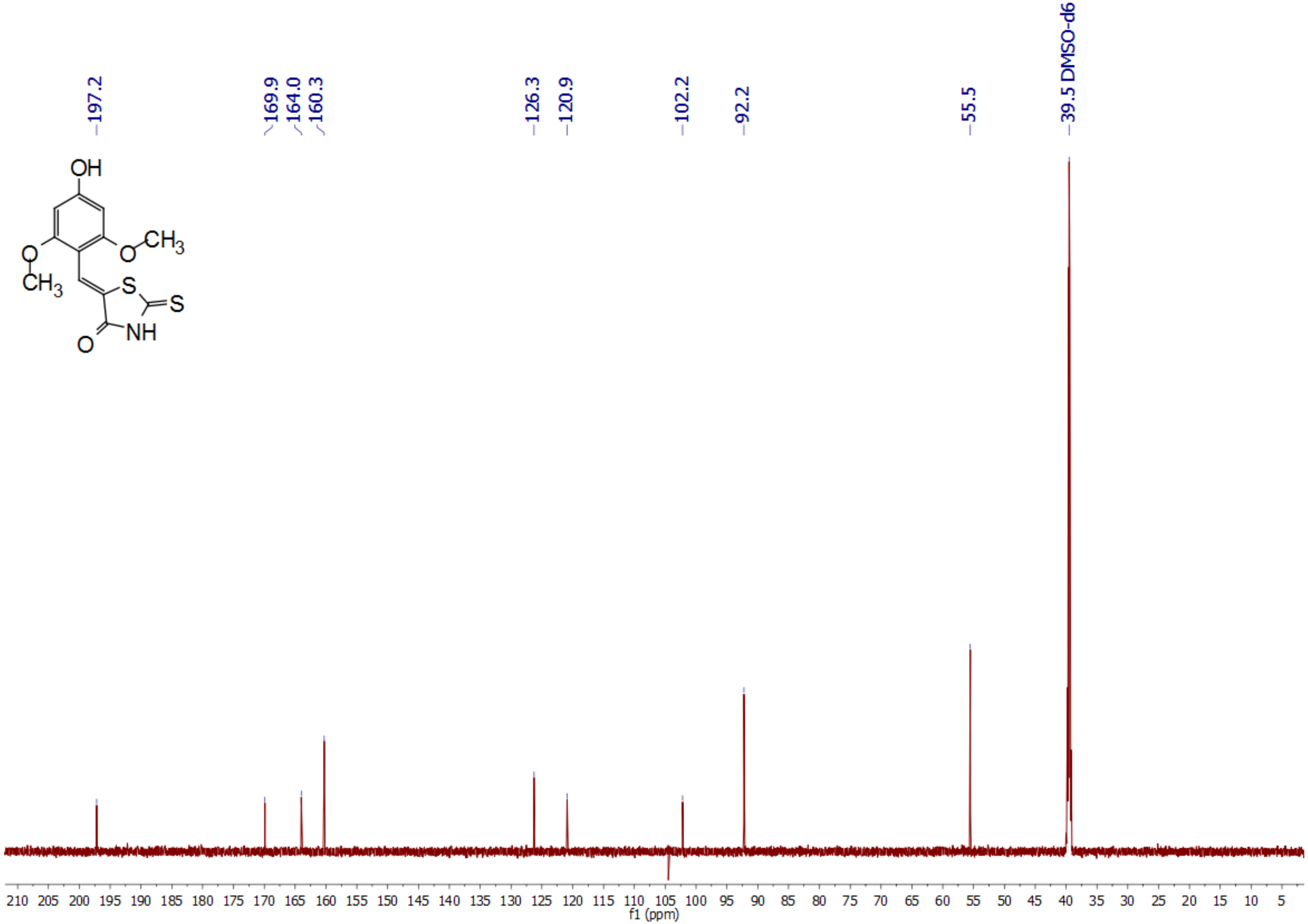
197.2
169.9
164.0
160.3
126.3
120.9
102.2
92.2
55.5
39.5 DMSO-d6
OH
CH3
O
S
S
NH
210 205 200 195 190 185 180 175 170 165 160 155 150 145 140 135 130 125 120 115 110 105 100 95 90 85 80 75 70 65 60 55 50 45 40 35 30 25 20 15 10 5
f1 (ppm)

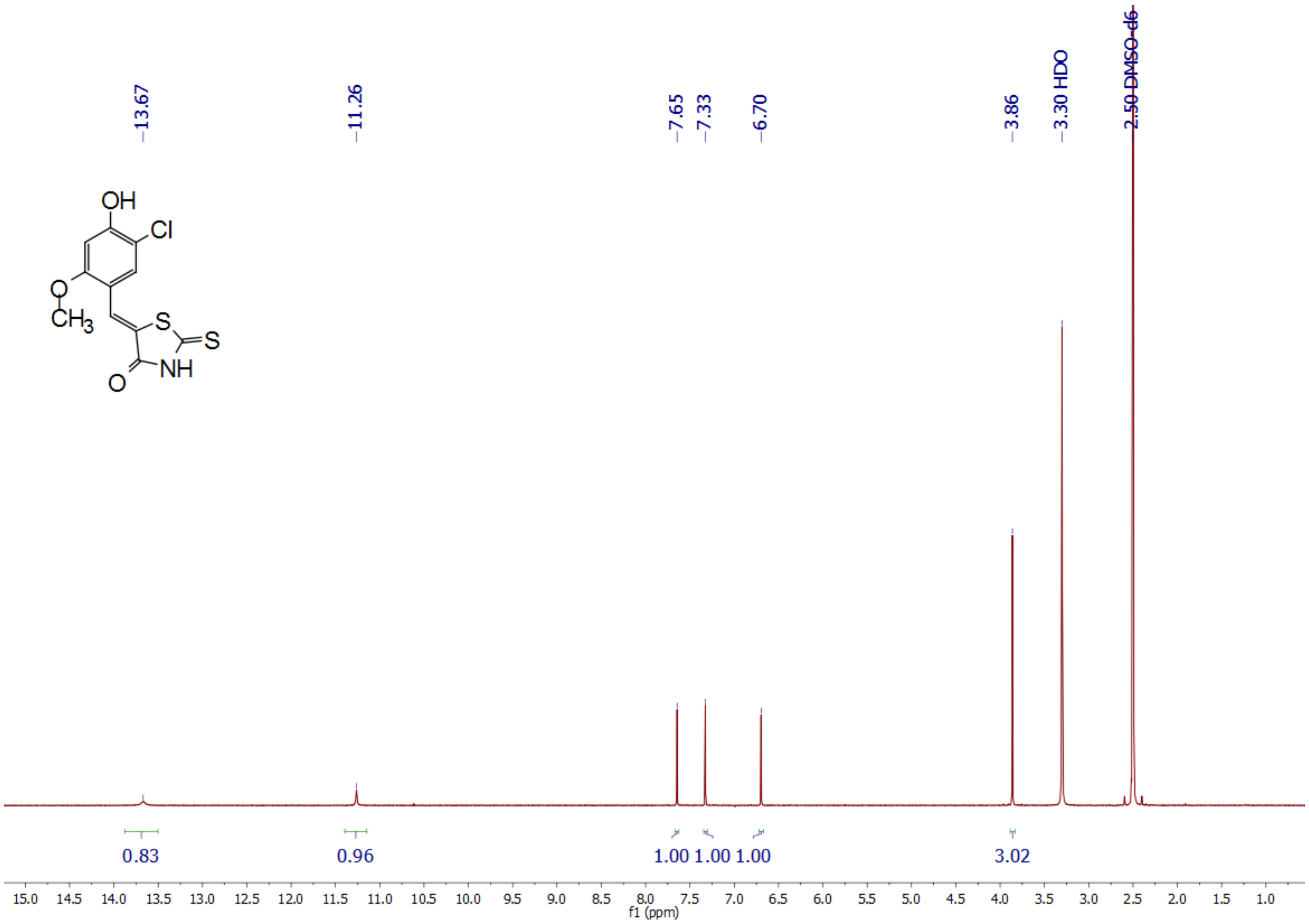

13.67
11.26
7.65
7.33
6.70
3.86
3.30 HDO
2.50 DMSO-d6
OH
Cl
O
CH3
S
S
NH
O
0.83
0.96
1.00
1.00
1.00
3.02
15.0
14.5
14.0
13.5
13.0
12.5
12.0
11.5
11.0
10.5
10.0
9.5
9.0
8.5
8.0
7.5
7.0
6.5
6.0
5.5
5.0
4.5
4.0
3.5
3.0
2.5
2.0
1.5
1.0
f1 (ppm)

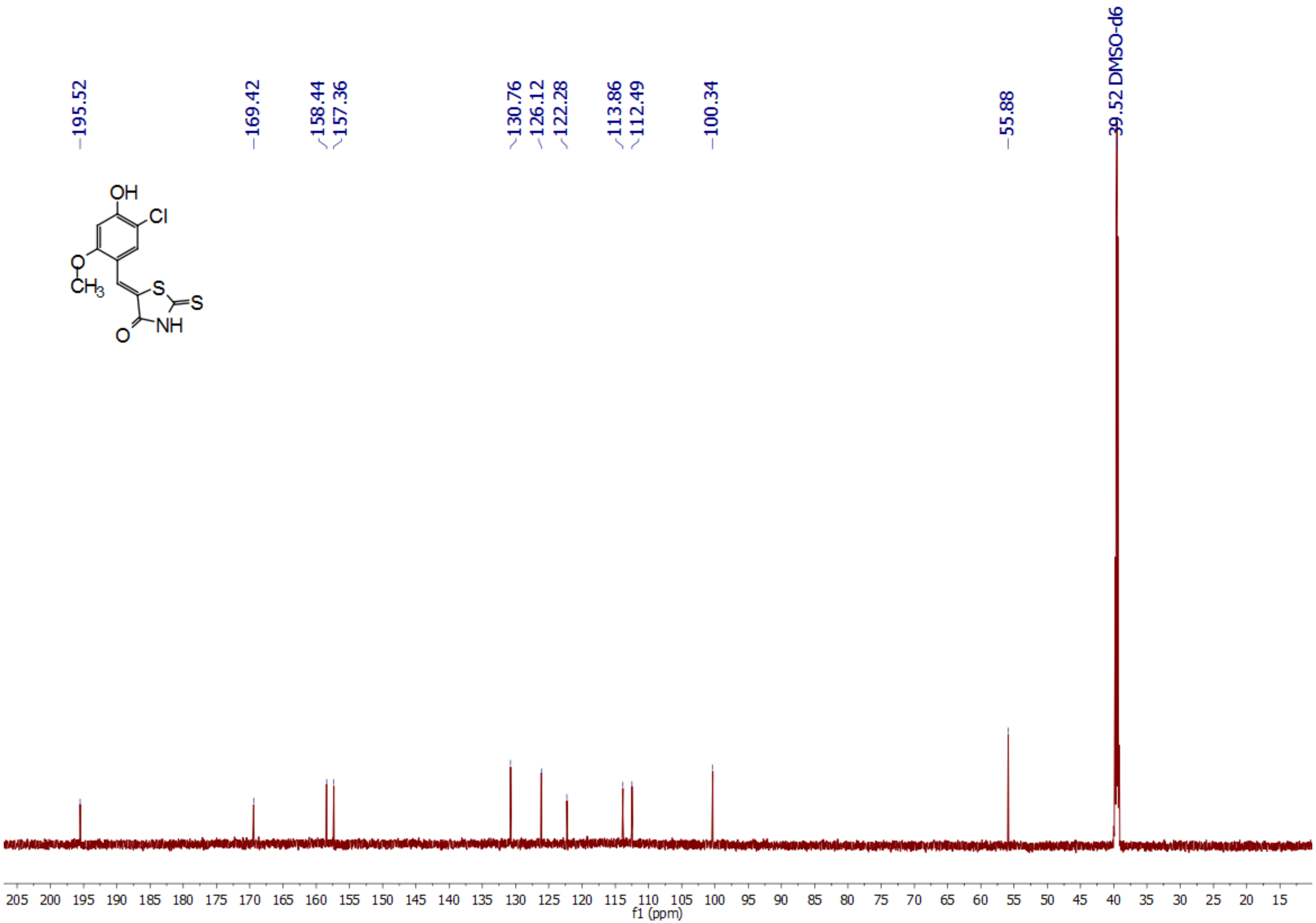
195.52
169.42
158.44
157.36
130.76
126.12
122.28
113.86
112.49
100.34
55.88
39.52 DMSO-d6
OH
Cl
O
CH3
S
S
NH
O
205 200 195 190 185 180 175 170 165 160 155 150 145 140 135 130 125 120 115 110 105 100 95 90 85 80 75 70 65 60 55 50 45 40 35 30 25 20 15
f1 (ppm)

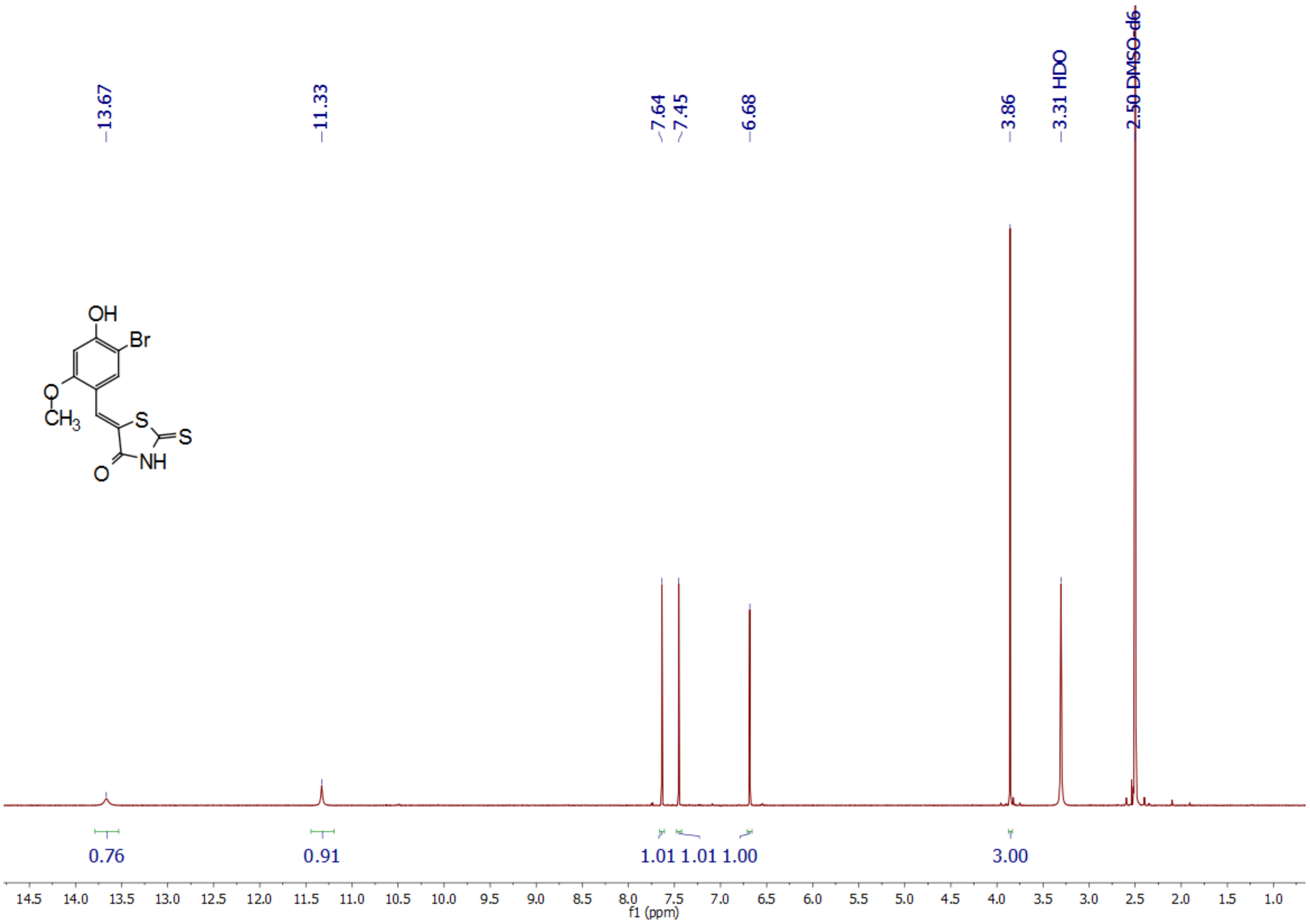
OH
Br
O
CH3
S
S
NH
O
13.67
11.33
7.64
7.45
6.68
3.86
3.31 HDO
2.50 DMSO-d6
0.76
0.91
1.01 1.01 1.00
3.00
14.5 14.0 13.5 13.0 12.5 12.0 11.5 11.0 10.5 10.0 9.5 9.0 8.5 8.0 7.5 7.0 6.5 6.0 5.5 5.0 4.5 4.0 3.5 3.0 2.5 2.0 1.5 1.0
f1 (ppm)

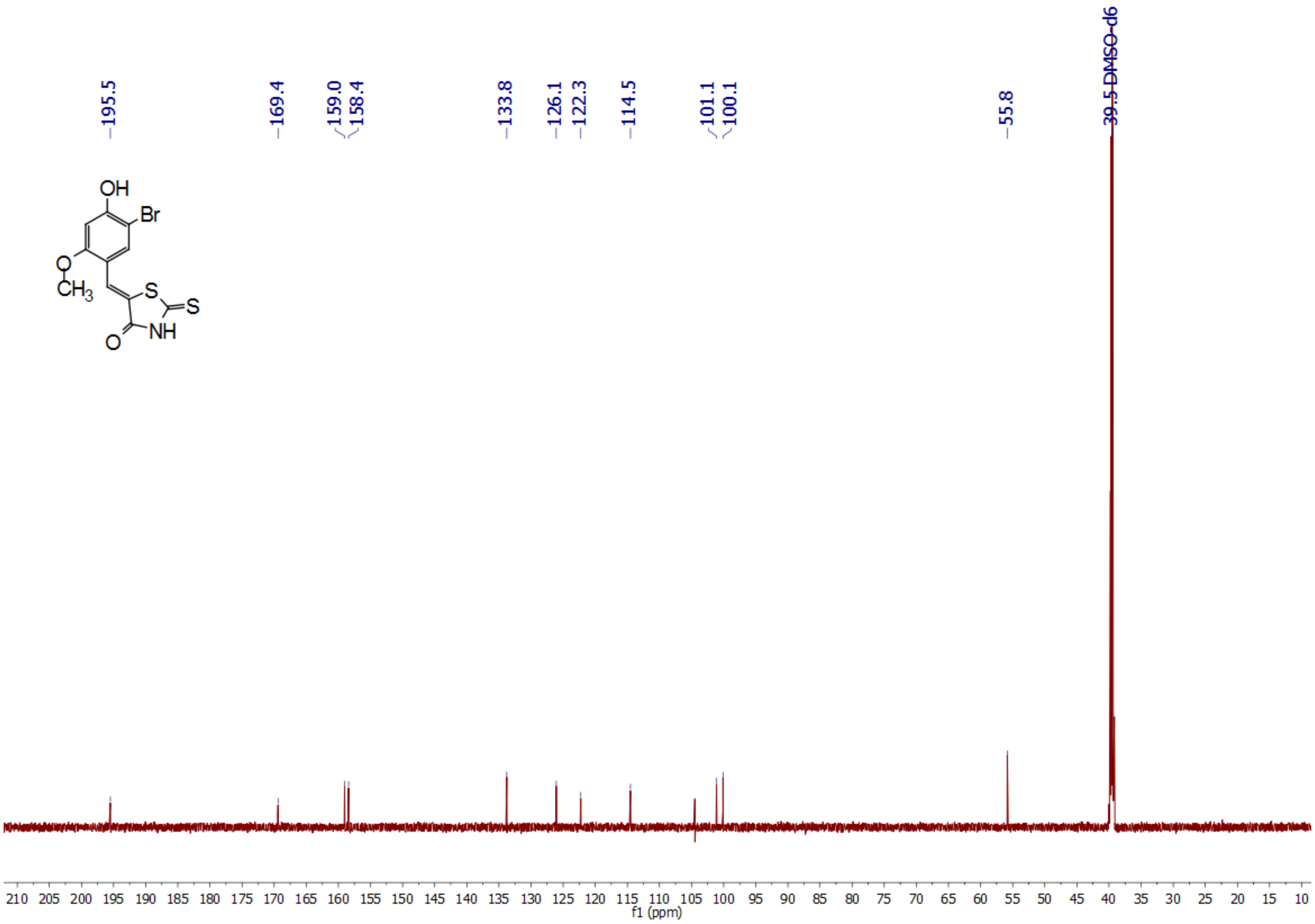

195.5
169.4
159.0
158.4
133.8
126.1
122.3
114.5
101.1
100.1
55.8
39.5 DMSO-d6
OH
Br
O
CH3
S
S
NH
O
210 205 200 195 190 185 180 175 170 165 160 155 150 145 140 135 130 125 120 115 110 105 100 95 90 85 80 75 70 65 60 55 50 45 40 35 30 25 20 15 10
f1 (ppm)

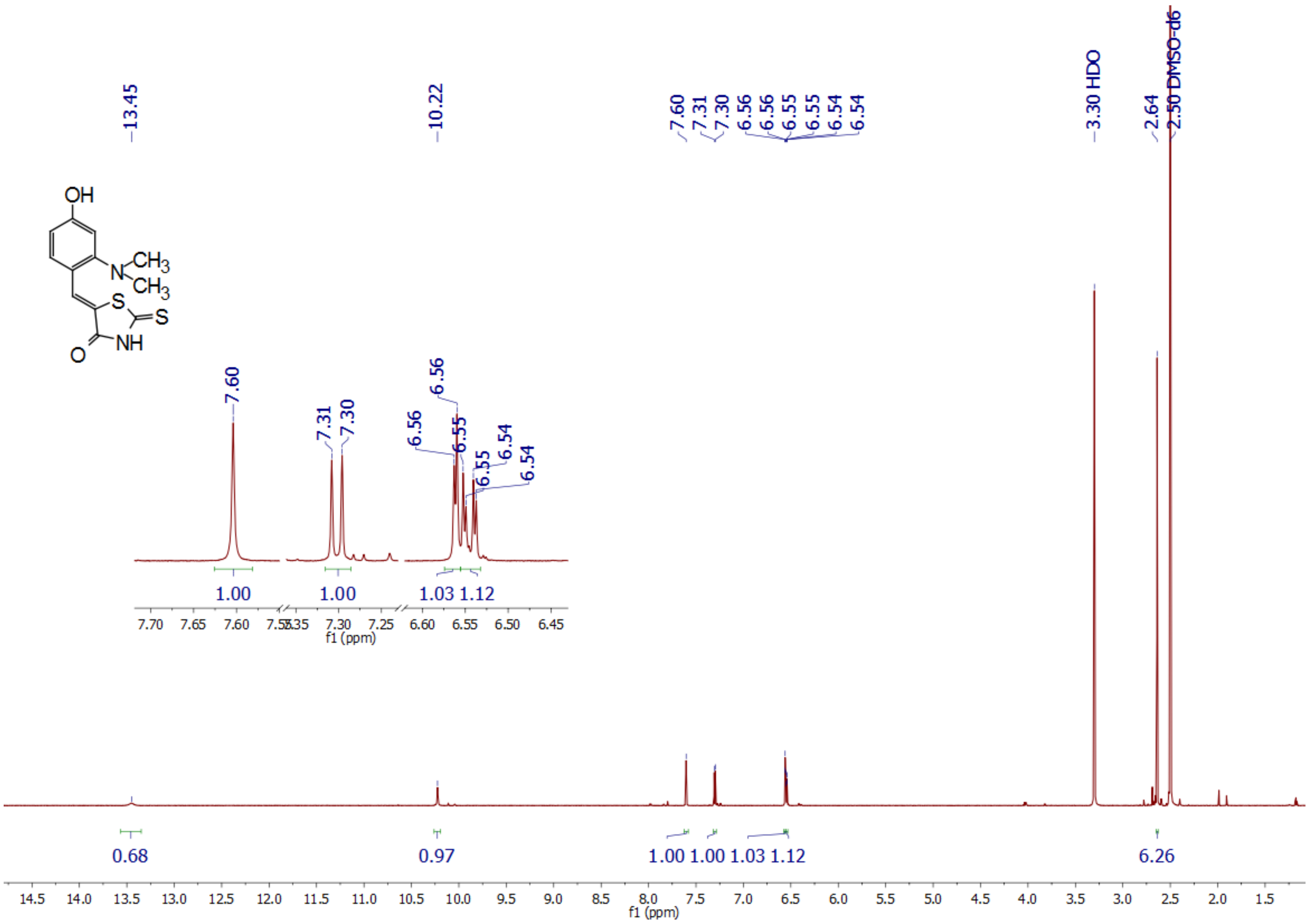

13.45
10.22
7.60
7.31
7.30
6.56
6.56
6.55
6.55
6.54
6.54
3.30 HDO
2.64
2.50 DMSO-d6
OH
N
CH3
CH3
S
S
NH
O
1.00
1.00
1.03 1.12
f1 (ppm)
0.68
0.97
1.00 1.00 1.03 1.12
6.26
f1 (ppm)

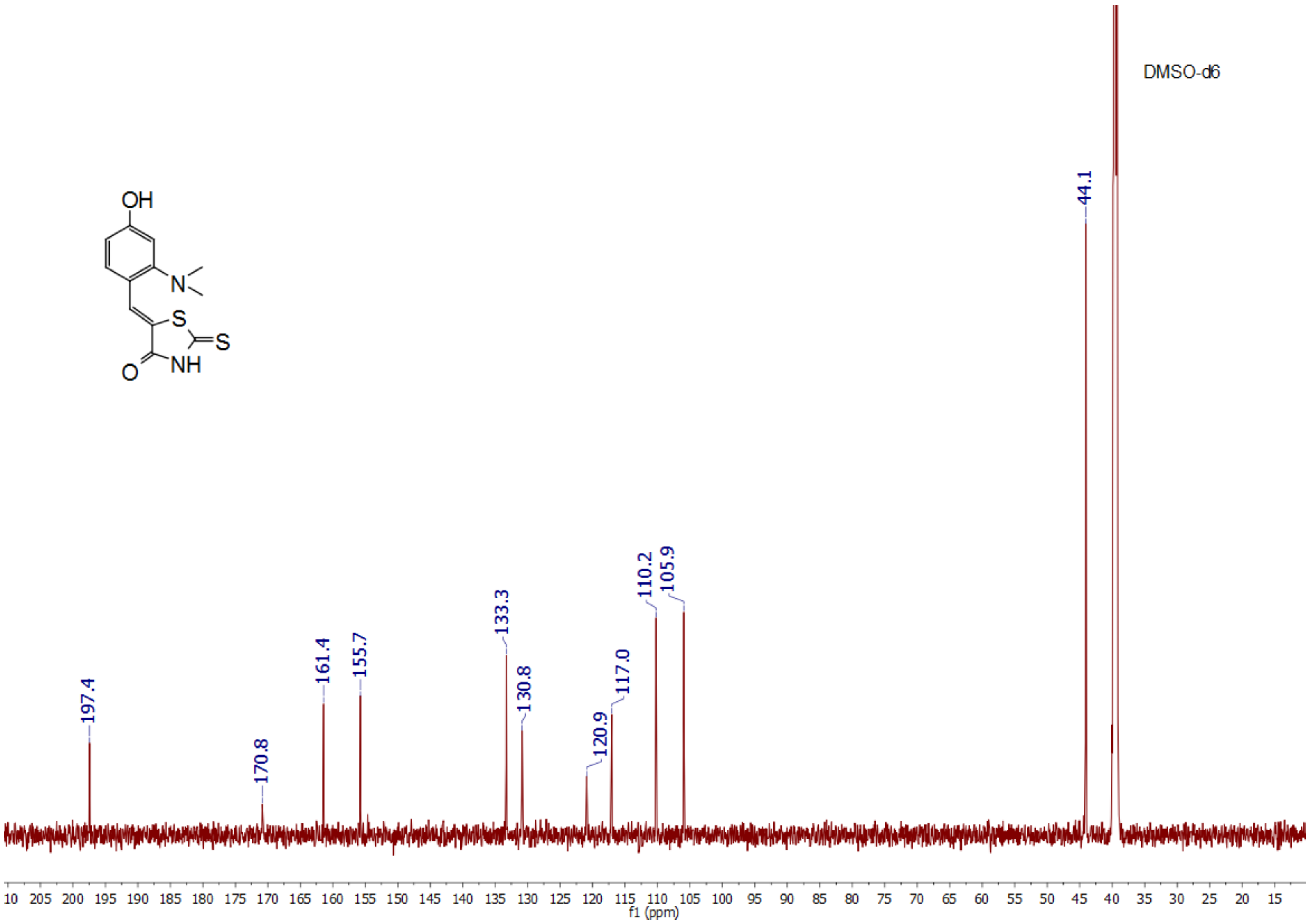
OH
N
S
S
NH
O
DMSO-d6
197.4
170.8
161.4
155.7
133.3
130.8
120.9
117.0
110.2
105.9
44.1
10 205 200 195 190 185 180 175 170 165 160 155 150 145 140 135 130 125 120 115 110 105 100 95 90 85 80 75 70 65 60 55 50 45 40 35 30 25 20 15
f1 (ppm)

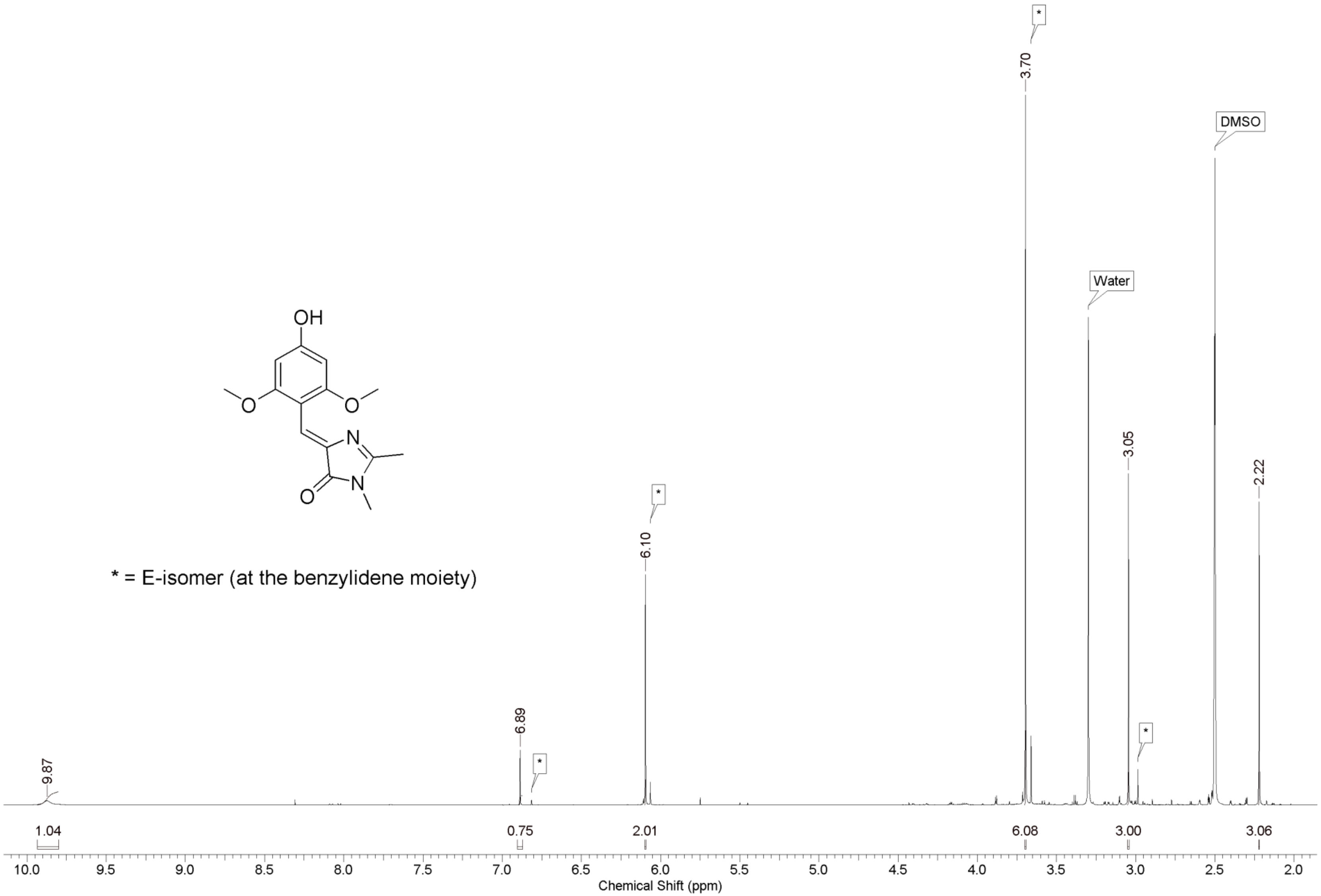

OH
O
O
N
N
O
* = E-isomer (at the benzylidene moiety)
9.87
6.89
6.10
3.70
3.05
2.22
*
Water
DMSO
1.04
0.75
2.01
6.08
3.00
3.06
10.0
9.5
9.0
8.5
8.0
7.5
7.0
6.5
6.0
5.5
5.0
4.5
4.0
3.5
3.0
2.5
2.0
Chemical Shift (ppm)

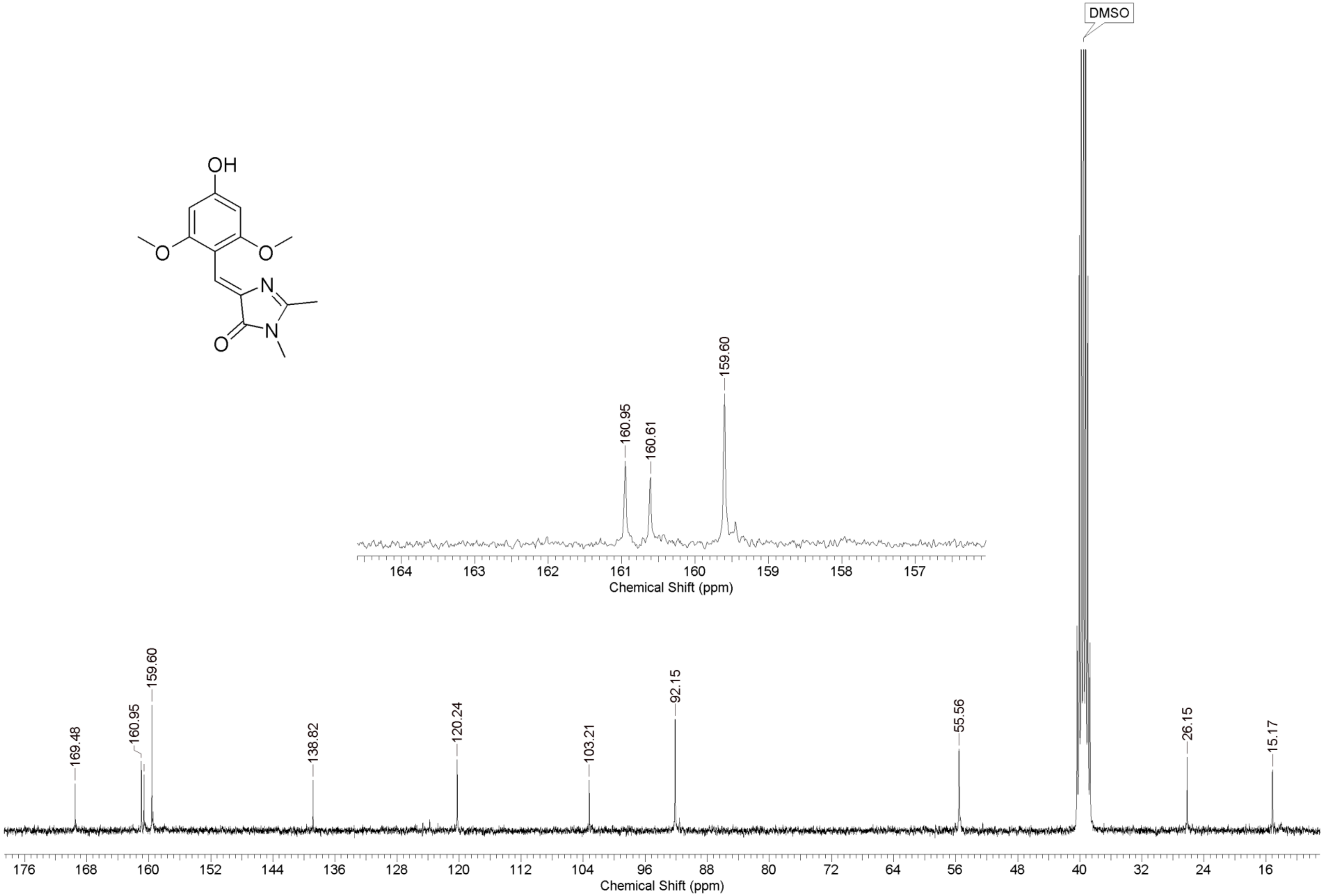
OH
O
O
N
O
N
DMSO
160.95
160.61
159.60
164 163 162 161 160 159 158 157
Chemical Shift (ppm)
169.48
160.95
159.60
138.82
120.24
103.21
92.15
55.56
26.15
15.17
176 168 160 152 144 136 128 120 112 104 96 88 80 72 64 56 48 40 32 24 16
Chemical Shift (ppm)

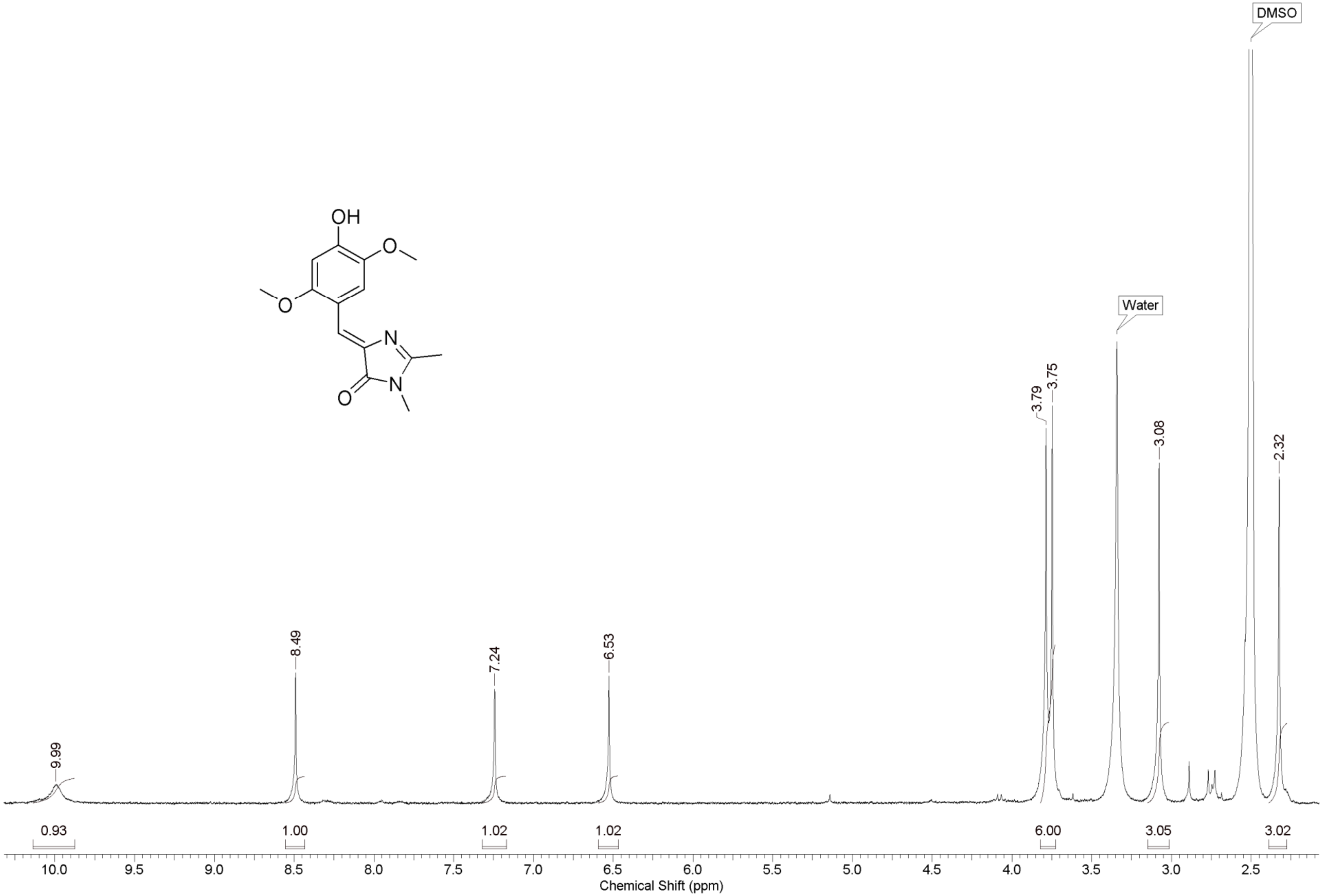
OH
O
O
N
N
O
DMSO
Water
9.99
8.49
7.24
6.53
3.79
3.75
3.08
2.32
0.93
1.00
1.02
1.02
6.00
3.05
3.02
10.0
9.5
9.0
8.5
8.0
7.5
7.0
6.5
6.0
5.5
5.0
4.5
4.0
3.5
3.0
2.5
Chemical Shift (ppm)

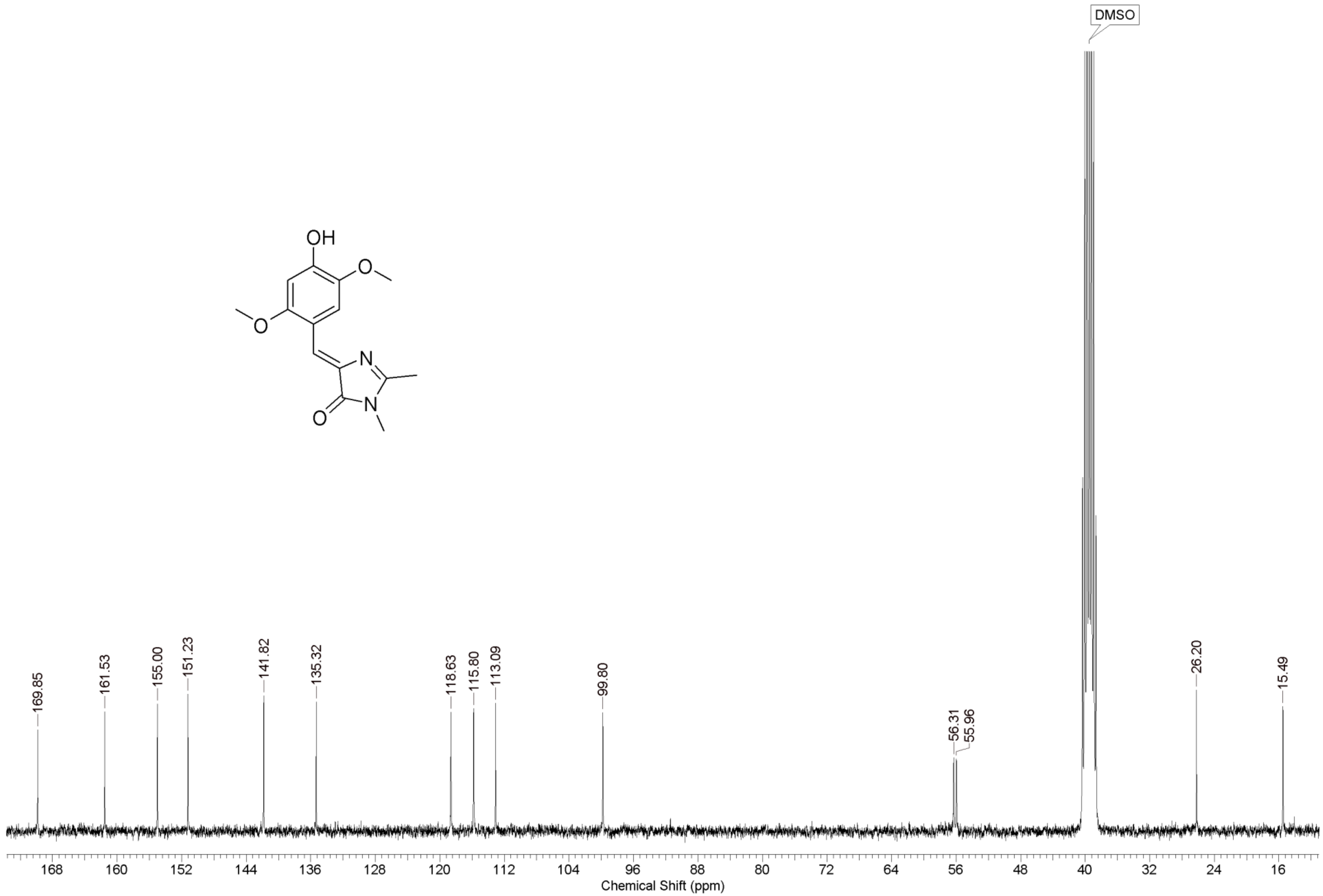

DMSO
OH
O
N
O
N
169.85
161.53
155.00
151.23
141.82
135.32
118.63
115.80
113.09
99.80
56.31
55.96
26.20
15.49
168
160
152
144
136
128
120
112
104
96
88
80
72
64
56
48
40
32
24
16
Chemical Shift (ppm)

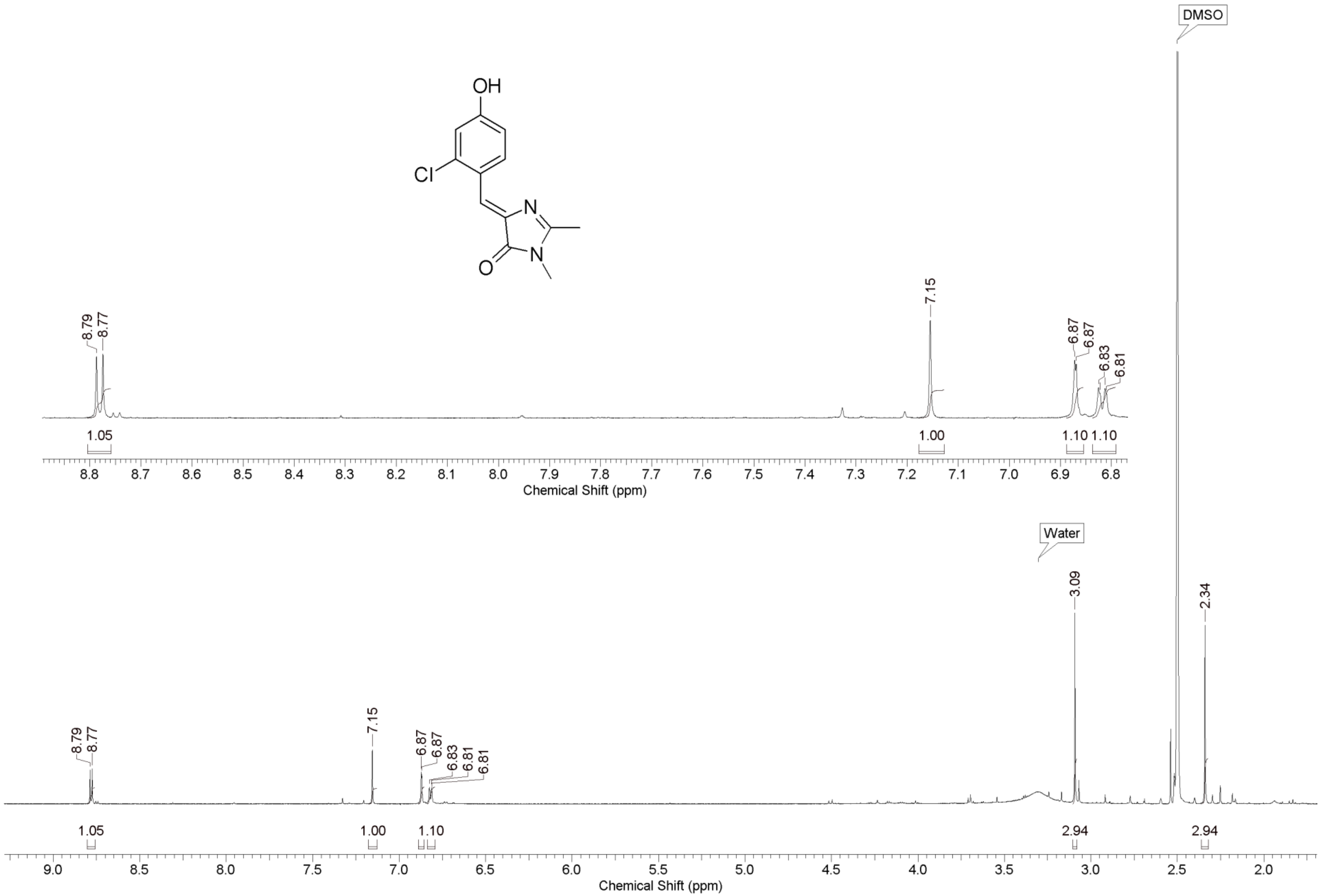

OH
Cl
N
N
O
DMSO
Water
8.79
8.77
7.15
6.87
6.87
6.83
6.81
6.81
3.09
2.34
1.05
1.00
1.10
1.10
2.94
2.94
Chemical Shift (ppm)
Chemical Shift (ppm)

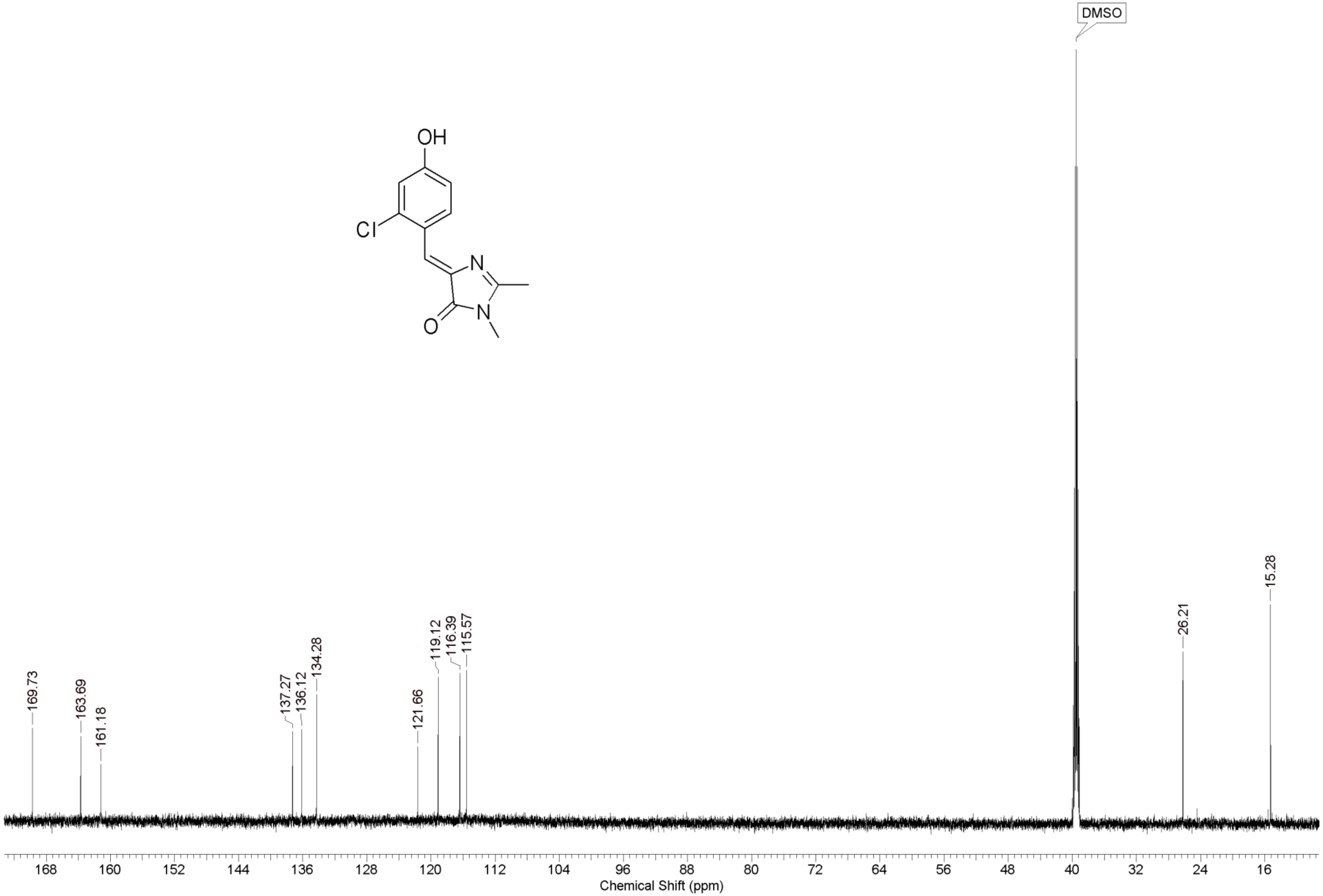

DMSO
OH
Cl
N
N
O
169.73
163.69
161.18
137.27
136.12
134.28
121.66
119.12
116.39
115.57
26.21
15.28
168
160
152
144
136
128
120
112
104
96
88
80
72
64
56
48
40
32
24
16
Chemical Shift (ppm)

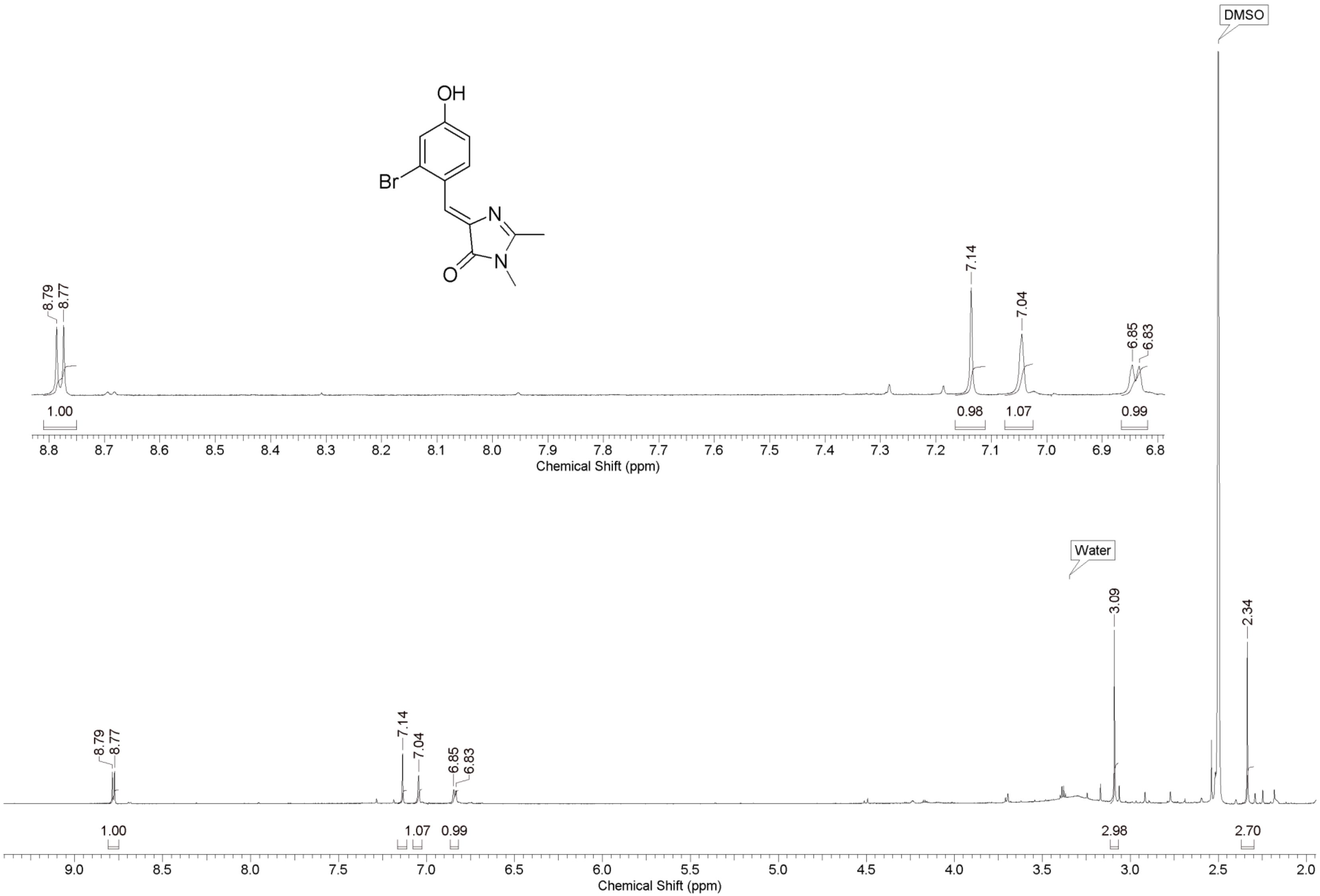
OH
Br
N
N
O
DMSO
8.79
8.77
7.14
7.04
6.85
6.83
1.00
0.98
1.07
0.99
8.8 8.7 8.6 8.5 8.4 8.3 8.2 8.1 8.0 7.9 7.8 7.7 7.6 7.5 7.4 7.3 7.2 7.1 7.0 6.9 6.8
Chemical Shift (ppm)
Water
3.09
2.34
8.79
8.77
7.14
7.04
6.85
6.83
1.00
1.07
0.99
2.98
2.70
9.0 8.5 8.0 7.5 7.0 6.5 6.0 5.5 5.0 4.5 4.0 3.5 3.0 2.5 2.0
Chemical Shift (ppm)

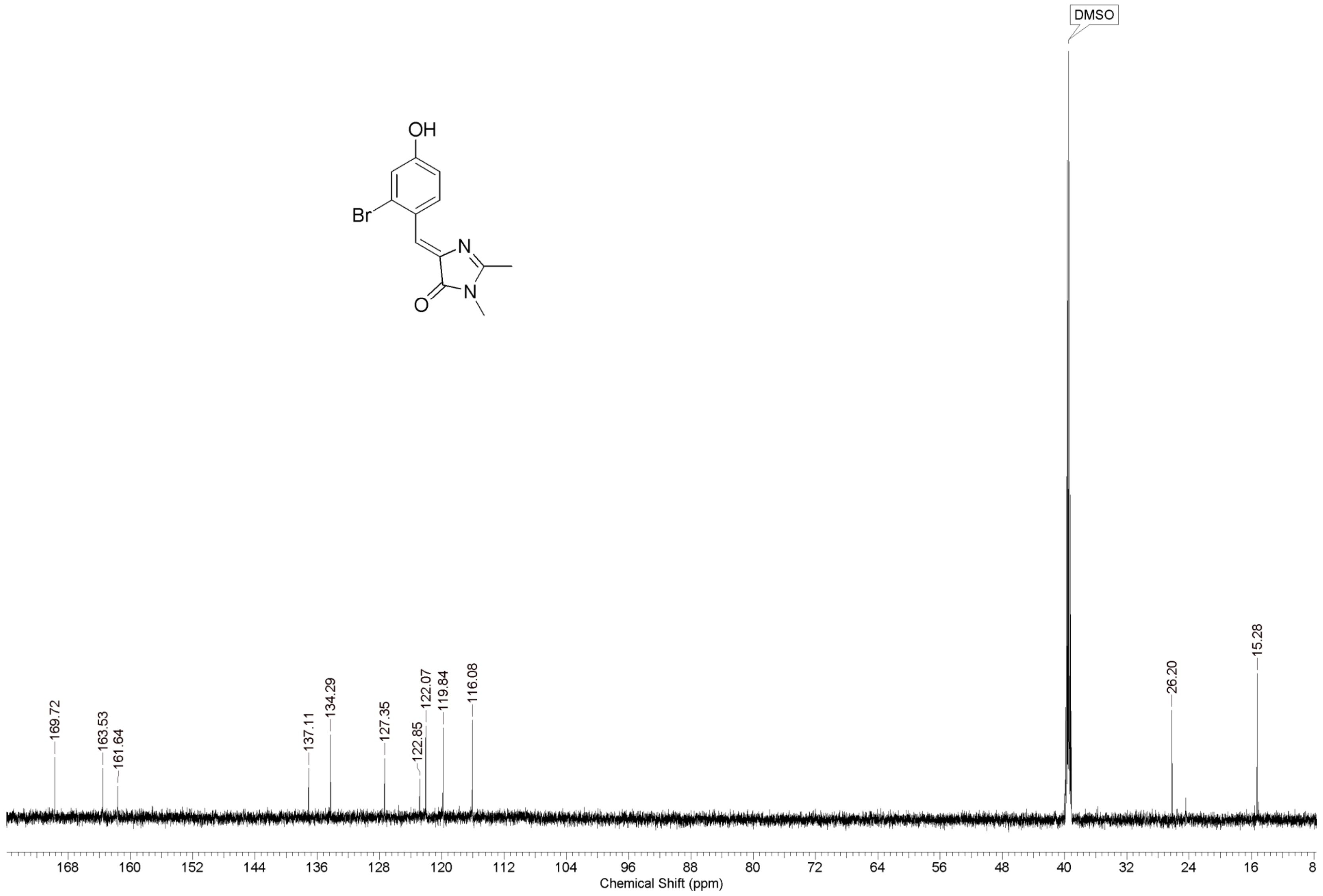
OH
Br
N
N
O
DMSO
169.72
163.53
161.64
137.11
134.29
127.35
122.85
122.07
119.84
116.08
26.20
15.28
168
160
152
144
136
128
120
112
104
96
88
80
72
64
56
48
40
32
24
16
8
Chemical Shift (ppm)

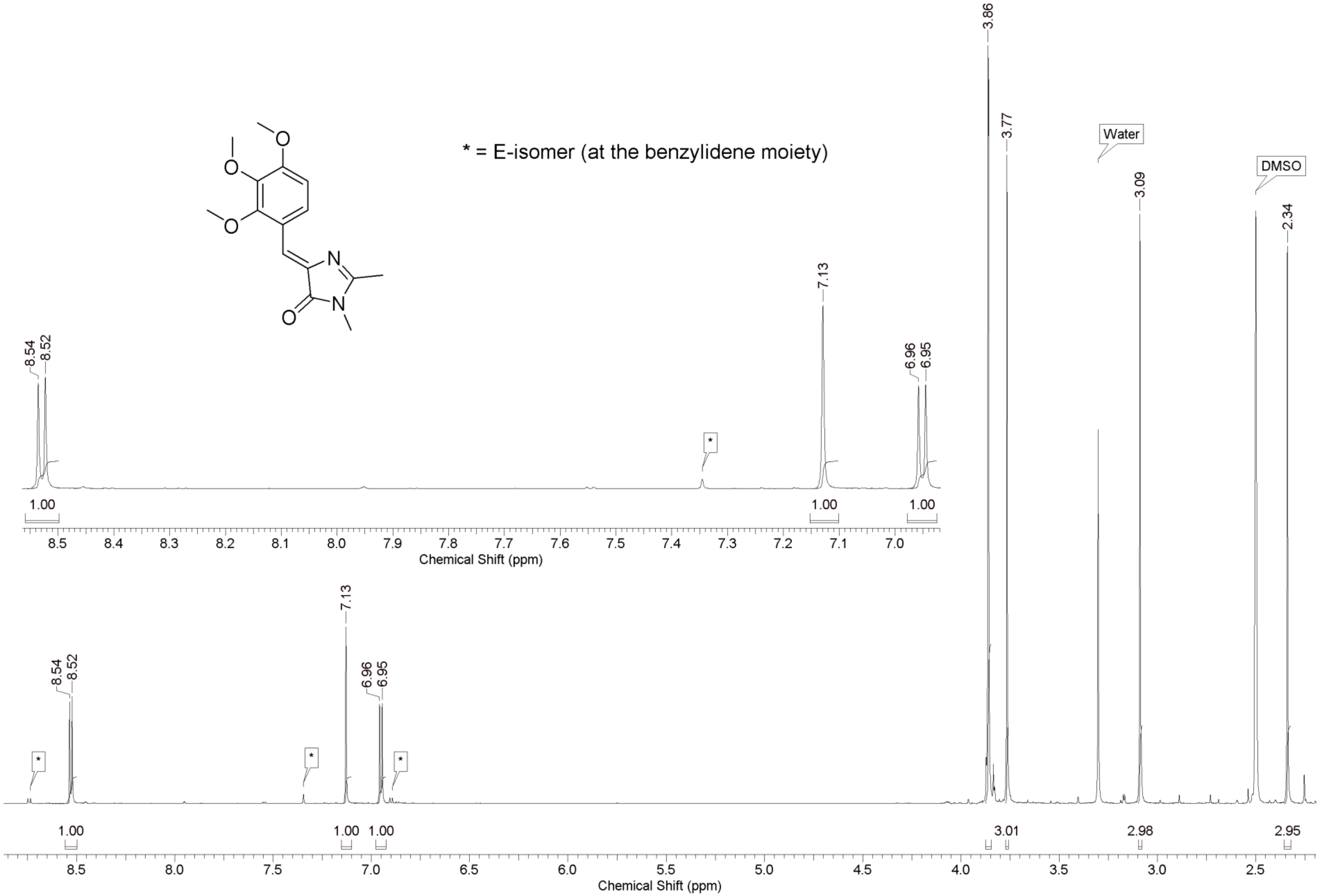
* = E-isomer (at the benzylidene moiety)
8.54
8.52
7.13
6.96
6.95
3.86
3.77
3.09
2.34
Water
DMSO
*
1.00
1.00
1.00
3.01
2.98
2.95
Chemical Shift (ppm)
Chemical Shift (ppm)

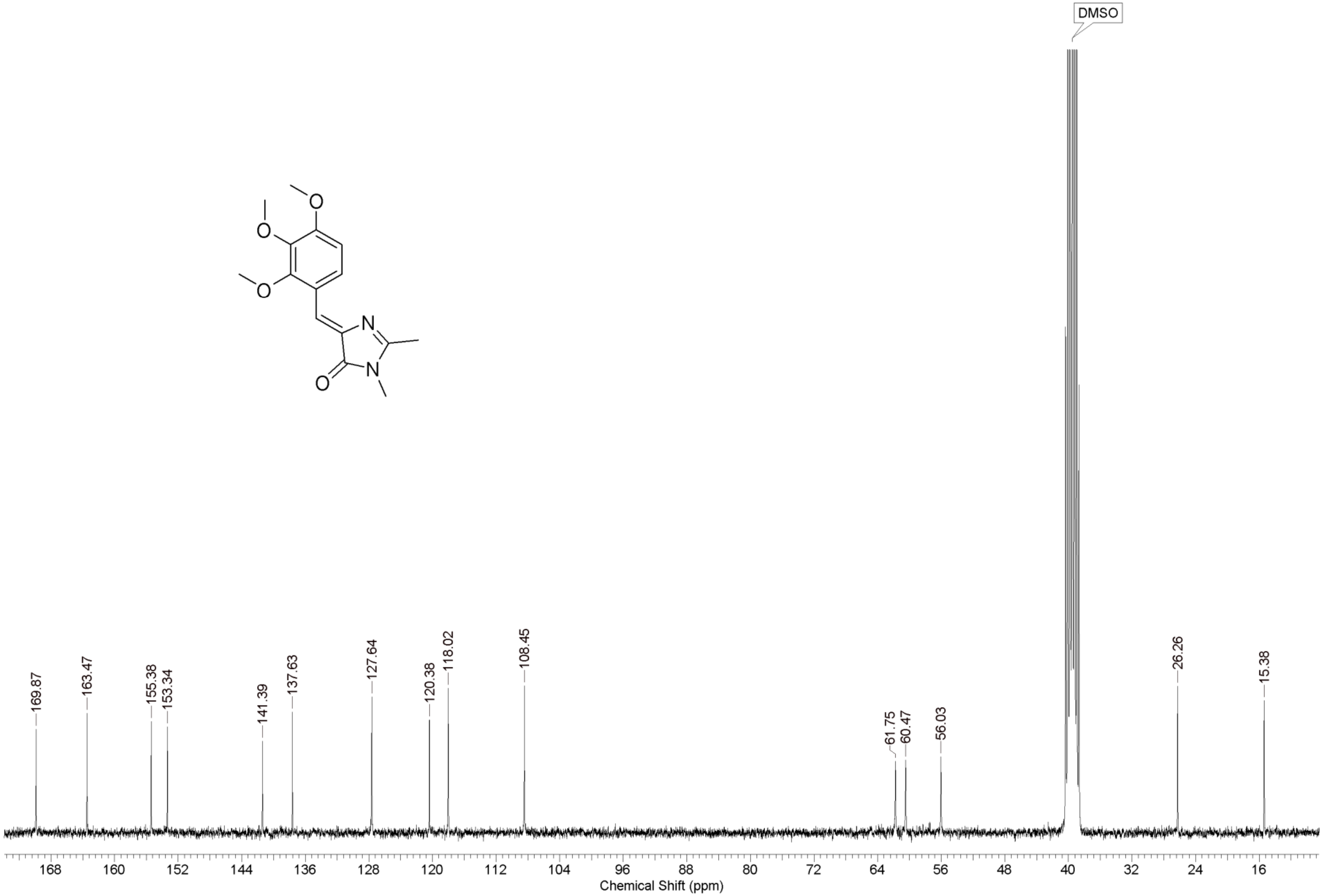
DMSO
169.87
163.47
155.38
153.34
141.39
137.63
127.64
120.38
118.02
108.45
61.75
60.47
56.03
26.26
15.38
168
160
152
144
136
128
120
112
104
96
88
80
72
64
56
48
40
32
24
16
Chemical Shift (ppm)

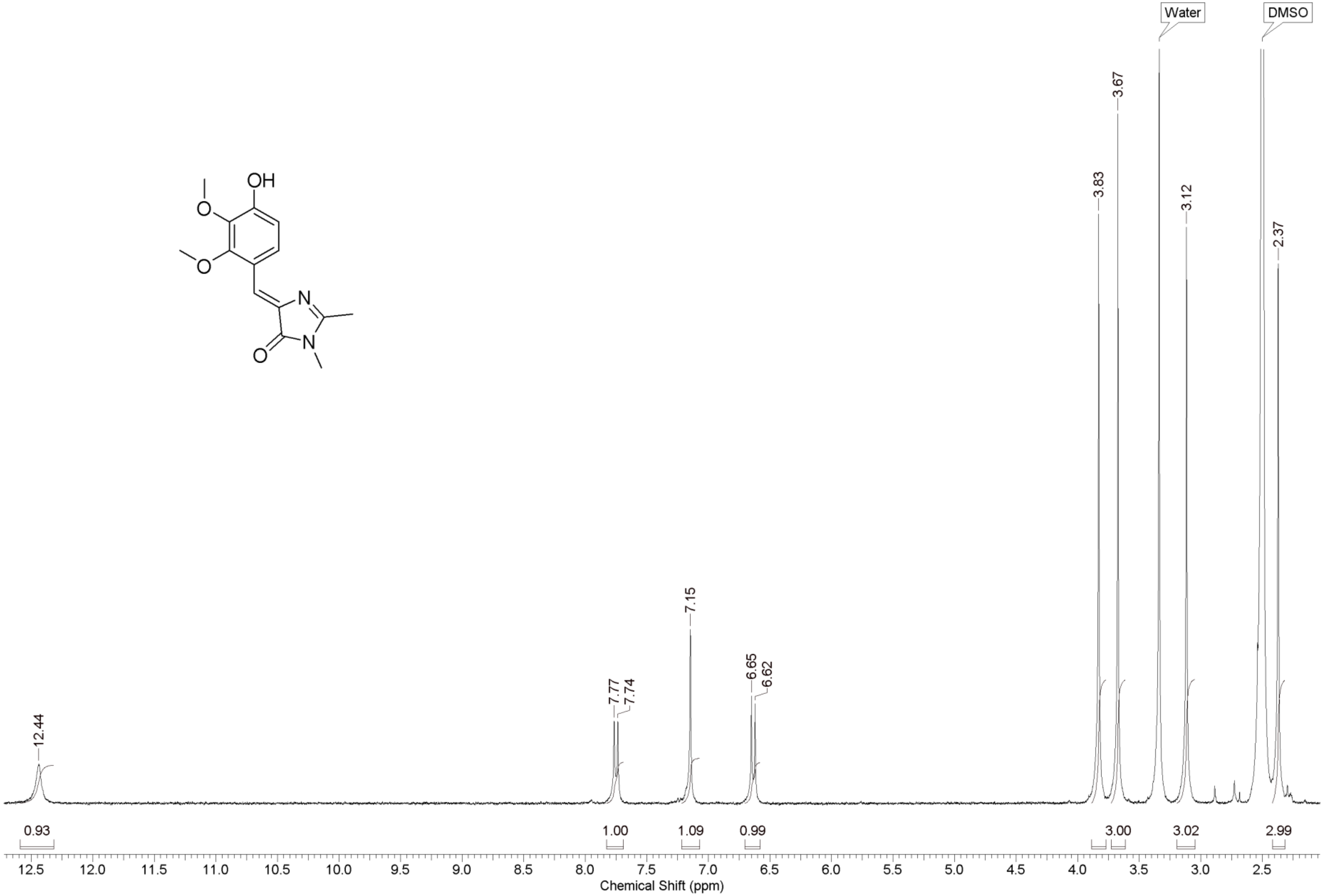
OH
O
O
N
O
N
Water
DMSO
3.67
3.83
3.12
2.37
7.15
7.77
7.74
6.65
6.62
12.44
0.93
1.00
1.09
0.99
3.00
3.02
2.99
12.5 12.0 11.5 11.0 10.5 10.0 9.5 9.0 8.5 8.0 7.5 7.0 6.5 6.0 5.5 5.0 4.5 4.0 3.5 3.0 2.5
Chemical Shift (ppm)

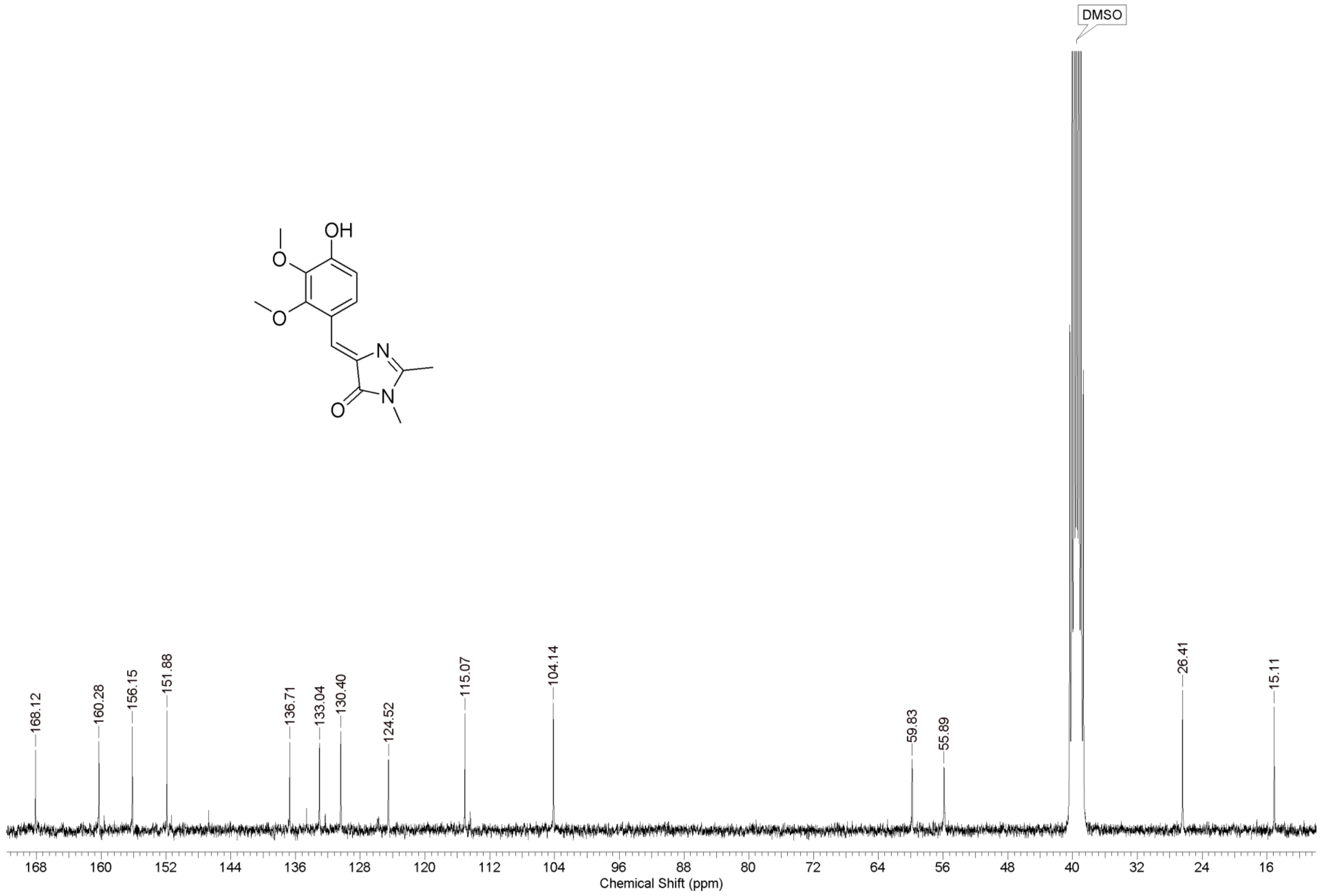

OH
O
O
N
N
O
DMSO
168.12
160.28
156.15
151.88
136.71
133.04
130.40
124.52
115.07
104.14
59.83
55.89
26.41
15.11
168
160
152
144
136
128
120
112
104
96
88
80
72
64
56
48
40
32
24
16
Chemical Shift (ppm)

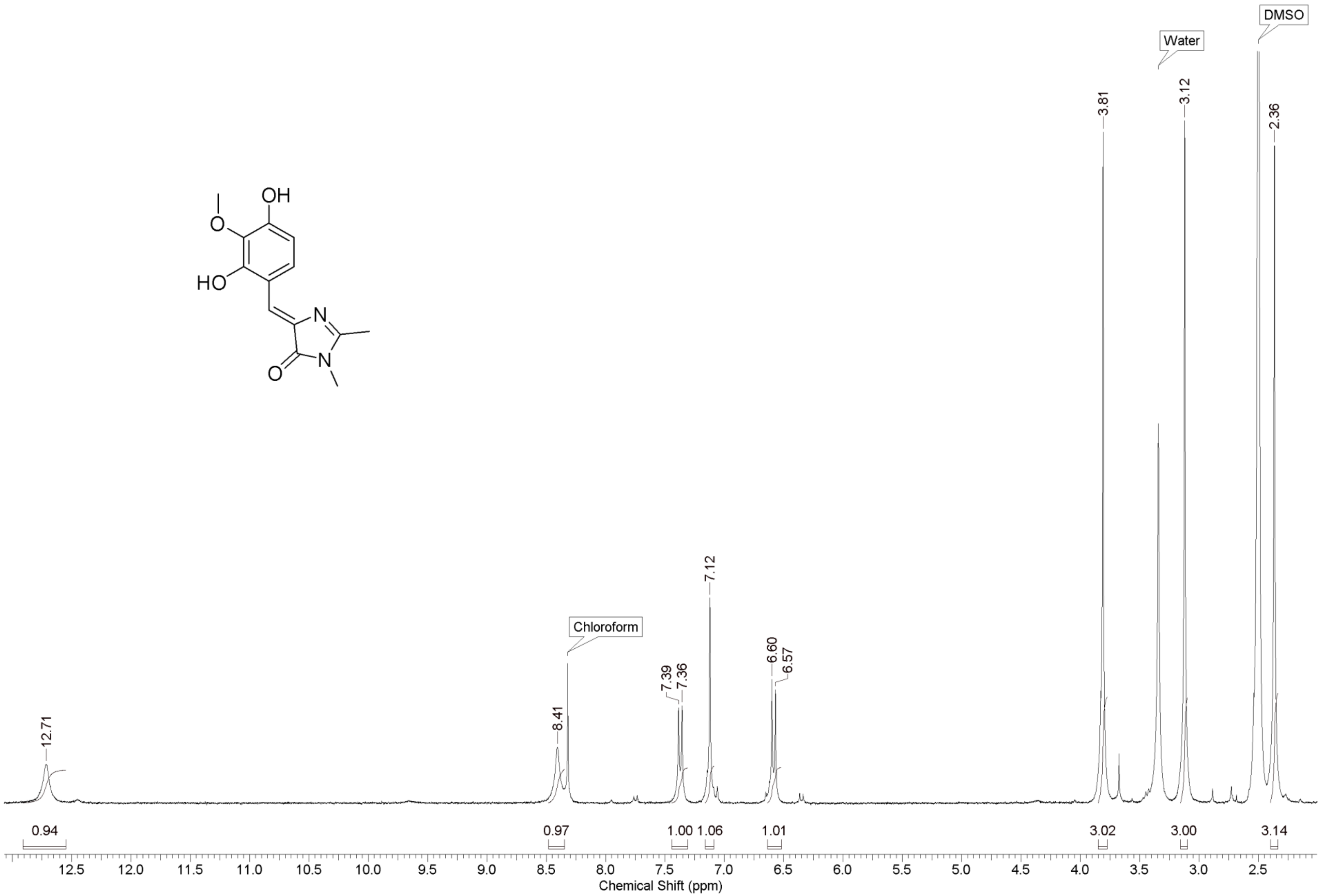
OH
O
HO
N
N
O
Water
DMSO
Chloroform
12.71
8.41
7.39
7.36
7.12
6.60
6.57
3.81
3.12
2.36
0.94
0.97
1.00
1.06
1.01
3.02
3.00
3.14
12.5
12.0
11.5
11.0
10.5
10.0
9.5
9.0
8.5
8.0
7.5
7.0
6.5
6.0
5.5
5.0
4.5
4.0
3.5
3.0
2.5
Chemical Shift (ppm)

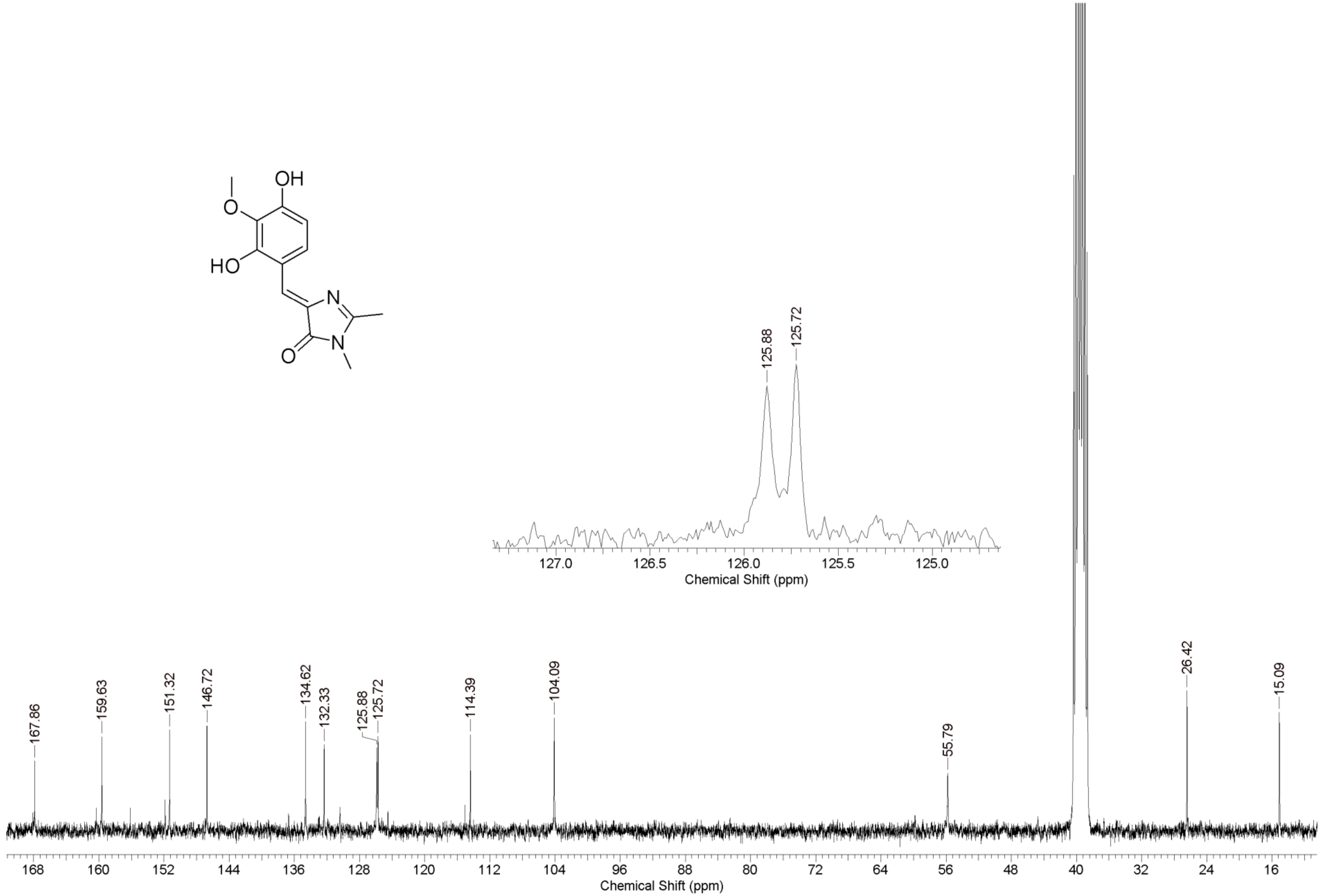
OH
O
HO
N
N
O
125.88
125.72
127.0
126.5
126.0
125.5
125.0
Chemical Shift (ppm)
167.86
159.63
151.32
146.72
134.62
132.33
125.88
125.72
114.39
104.09
55.79
26.42
15.09
168
160
152
144
136
128
120
112
104
96
88
80
72
64
56
48
40
32
24
16
Chemical Shift (ppm)

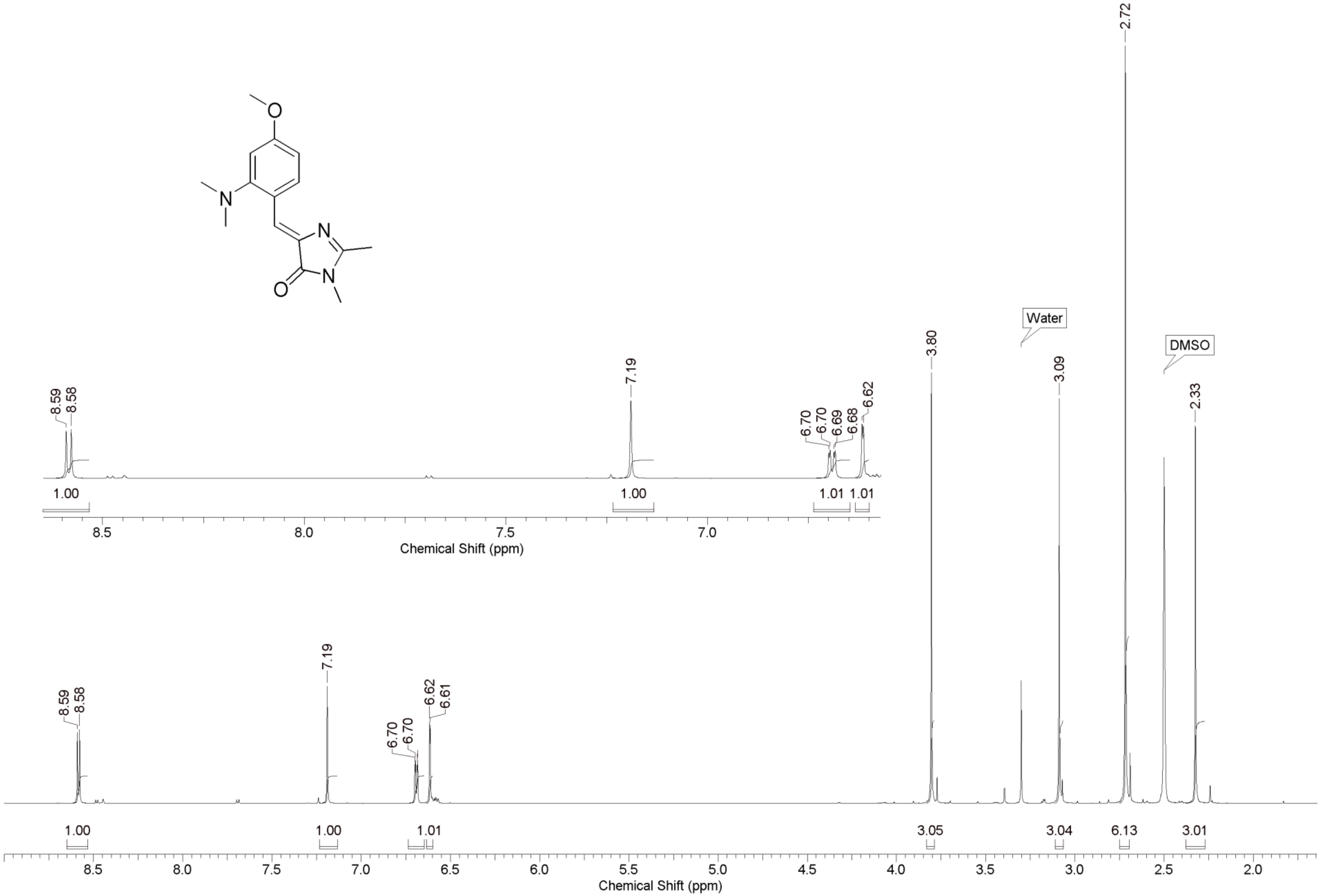
O
N
N
N
O
Water
DMSO
8.59
8.58
7.19
6.70
6.70
6.69
6.68
6.62
6.61
3.80
3.09
2.72
2.33
1.00
1.00
1.01
1.01
3.05
3.04
6.13
3.01
Chemical Shift (ppm)

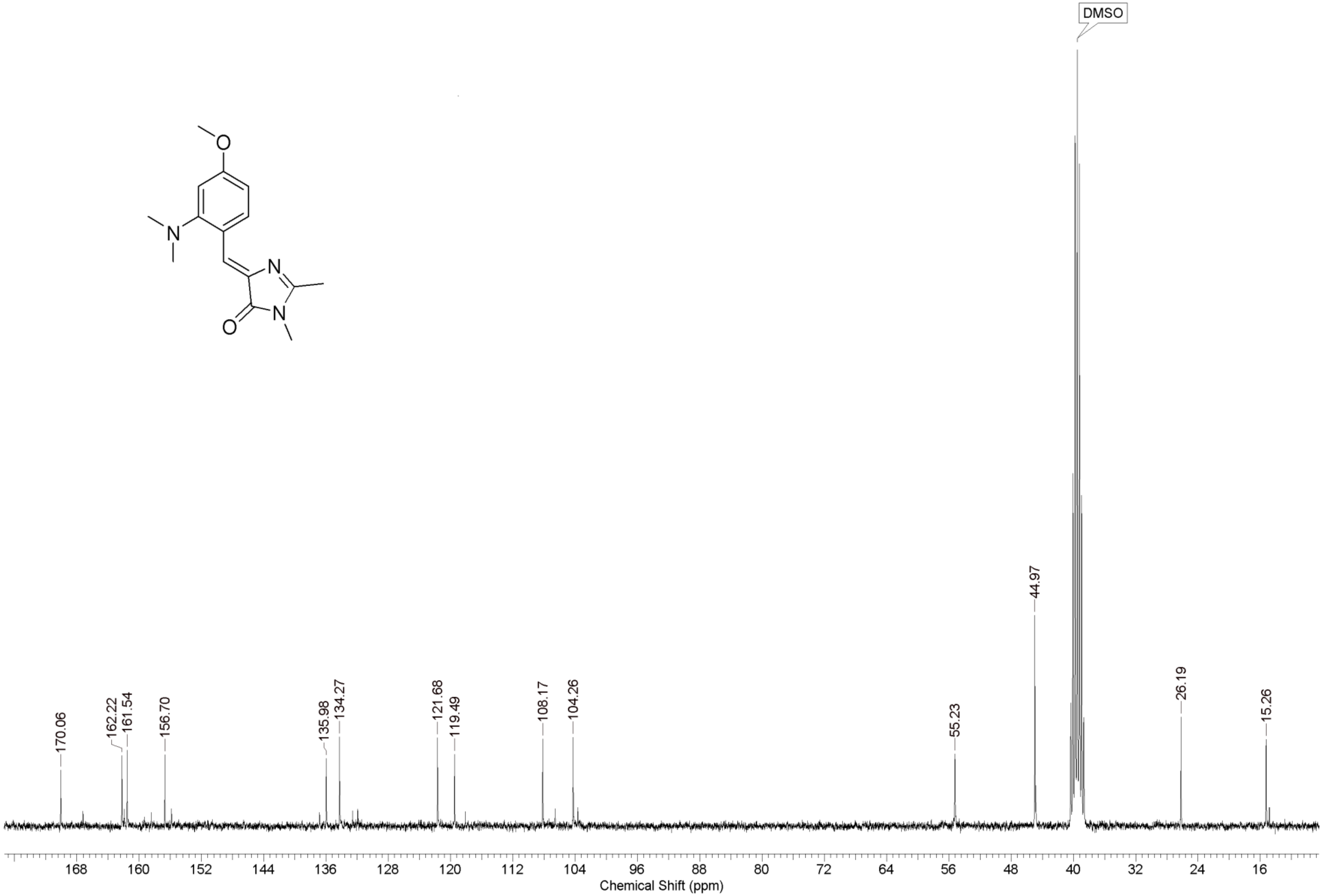
DMSO
170.06
162.22
161.54
156.70
135.98
134.27
121.68
119.49
108.17
104.26
55.23
44.97
26.19
15.26
168
160
152
144
136
128
120
112
104
96
88
80
72
64
56
48
40
32
24
16
Chemical Shift (ppm)

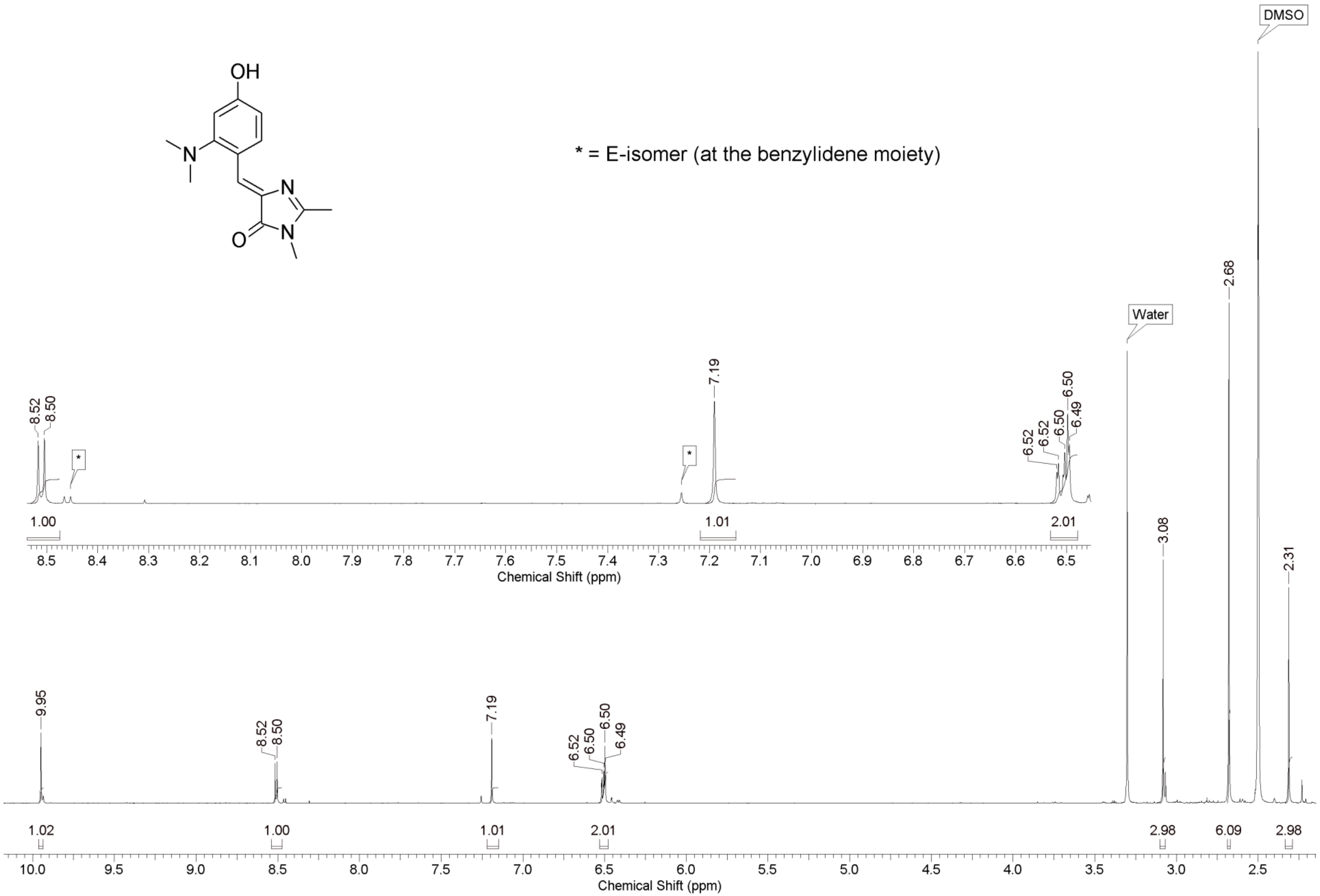

OH
N
N
O
N
* = E-isomer (at the benzylidene moiety)
DMSO
Water
2.68
3.08
2.31
8.52
8.50
7.19
6.52
6.52
6.50
6.50
6.49
1.00
1.01
2.01
Chemical Shift (ppm)
9.95
1.02
2.98
6.09
2.98
Chemical Shift (ppm)

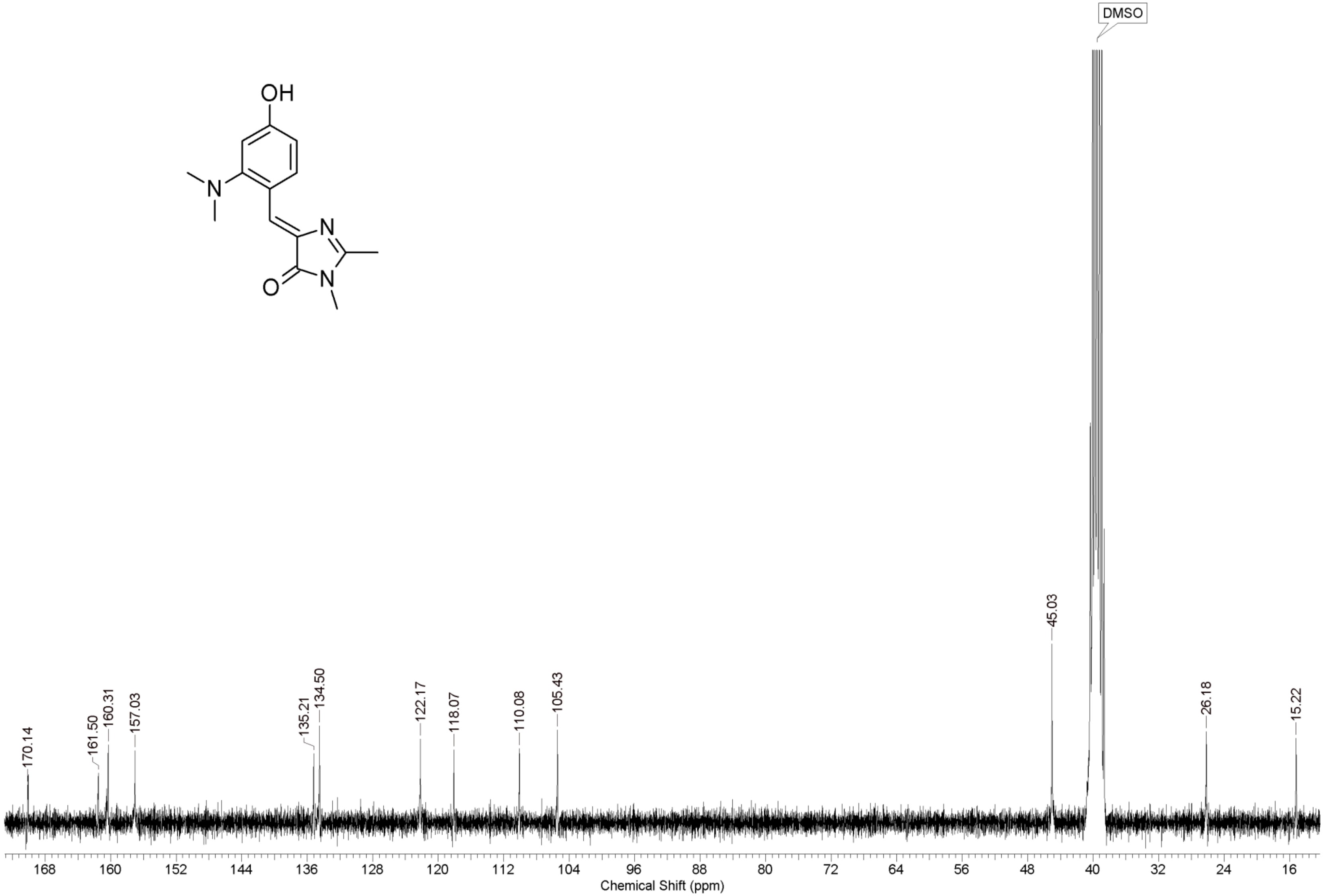
OH
N
N
N
O
DMSO
170.14
161.50
160.31
157.03
135.21
134.50
122.17
118.07
110.08
105.43
45.03
26.18
15.22
168 160 152 144 136 128 120 112 104 96 88 80 72 64 56 48 40 32 24 16
Chemical Shift (ppm)

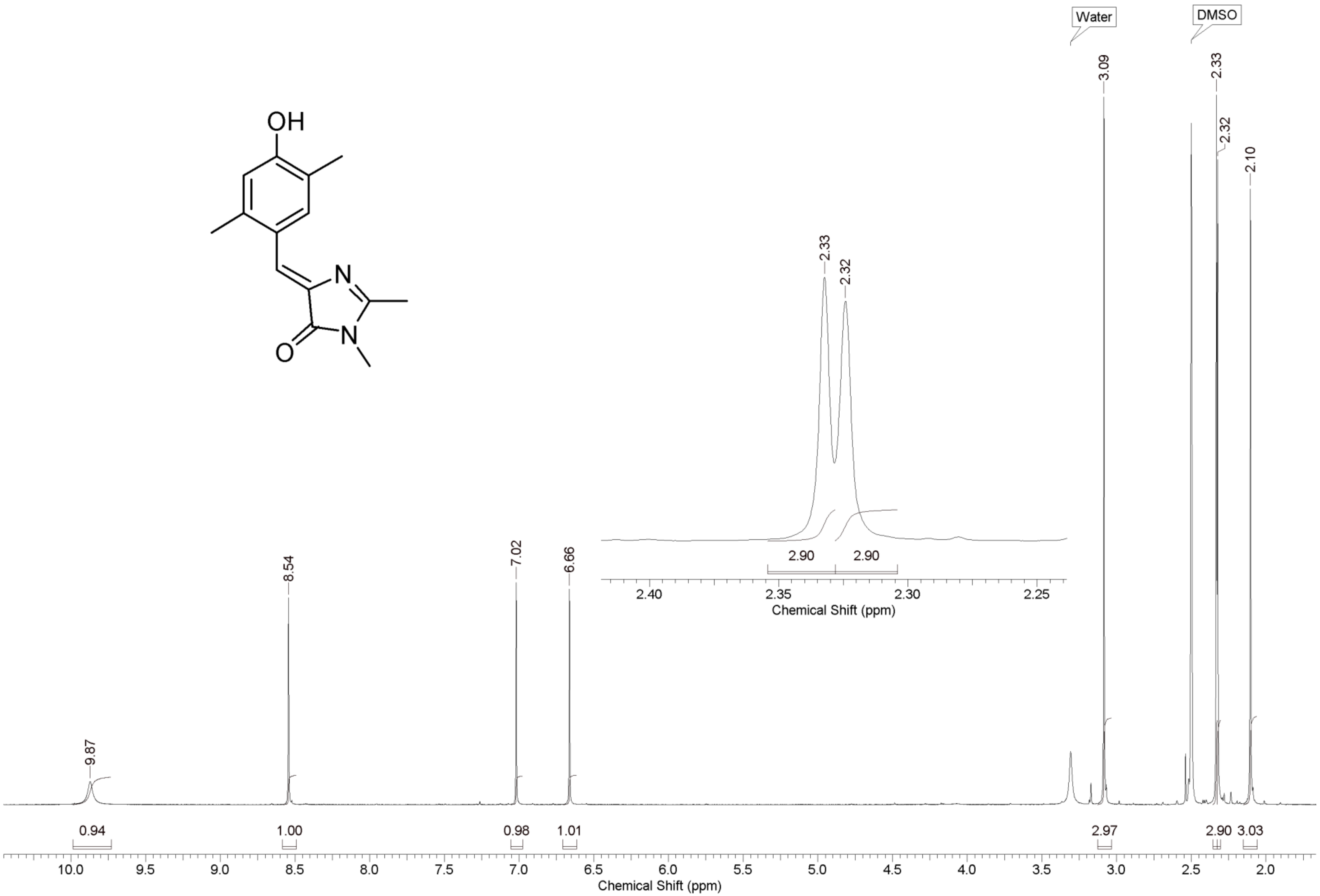
OH
N
N
O
Water
DMSO
3.09
2.33
2.32
2.10
9.87
8.54
7.02
6.66
0.94
1.00
0.98
1.01
2.97
2.90
3.03
10.0 9.5 9.0 8.5 8.0 7.5 7.0 6.5 6.0 5.5 5.0 4.5 4.0 3.5 3.0 2.5 2.0
Chemical Shift (ppm)
2.33
2.32
2.90
2.90
2.40 2.35 2.30 2.25
Chemical Shift (ppm)

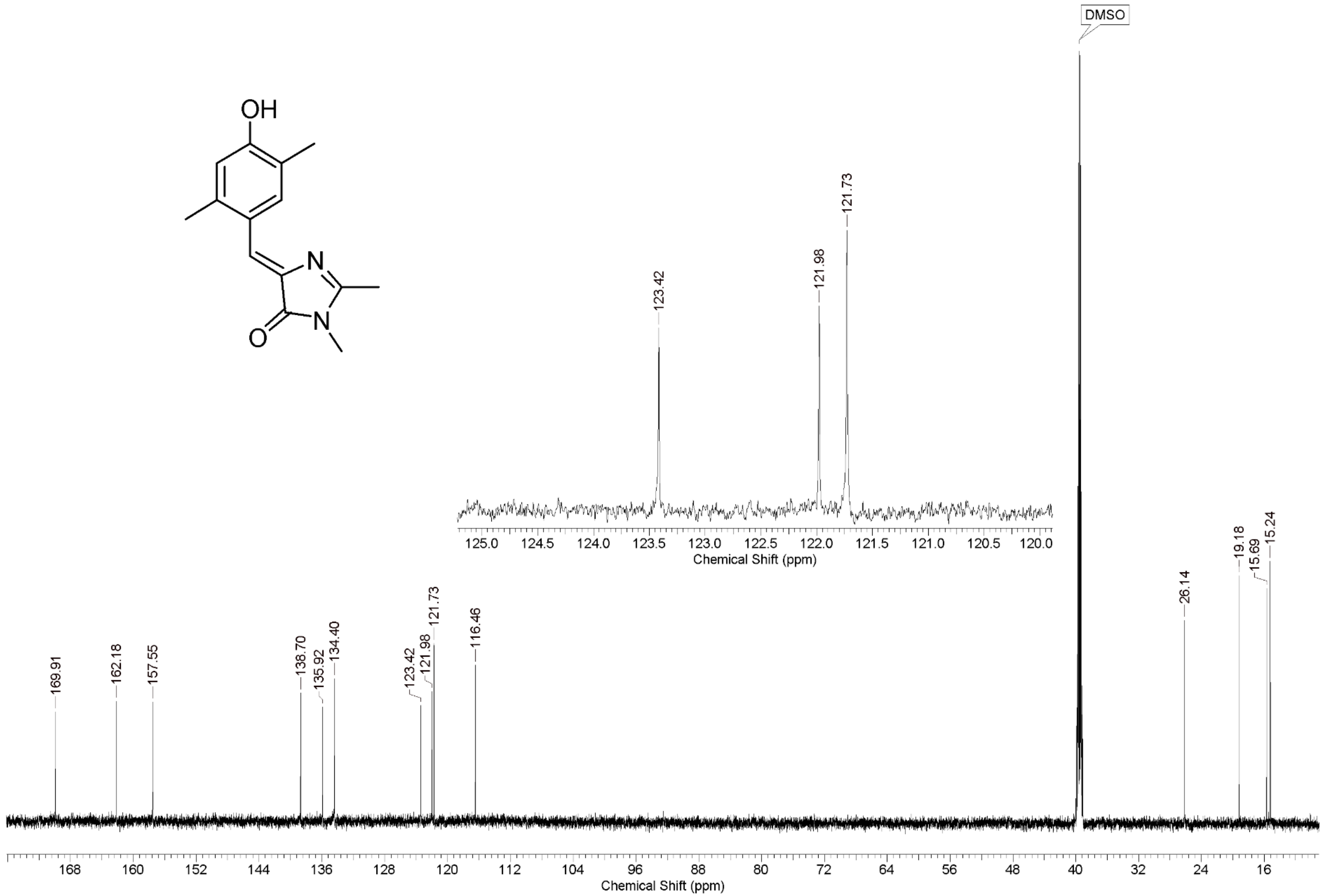
OH
N
N
O
DMSO
123.42
121.98
121.73
125.0 124.5 124.0 123.5 123.0 122.5 122.0 121.5 121.0 120.5 120.0
Chemical Shift (ppm)
169.91
162.18
157.55
138.70
135.92
134.40
123.42
121.98
121.73
116.46
26.14
19.18
15.69
15.24
168 160 152 144 136 128 120 112 104 96 88 80 72 64 56 48 40 32 24 16
Chemical Shift (ppm)

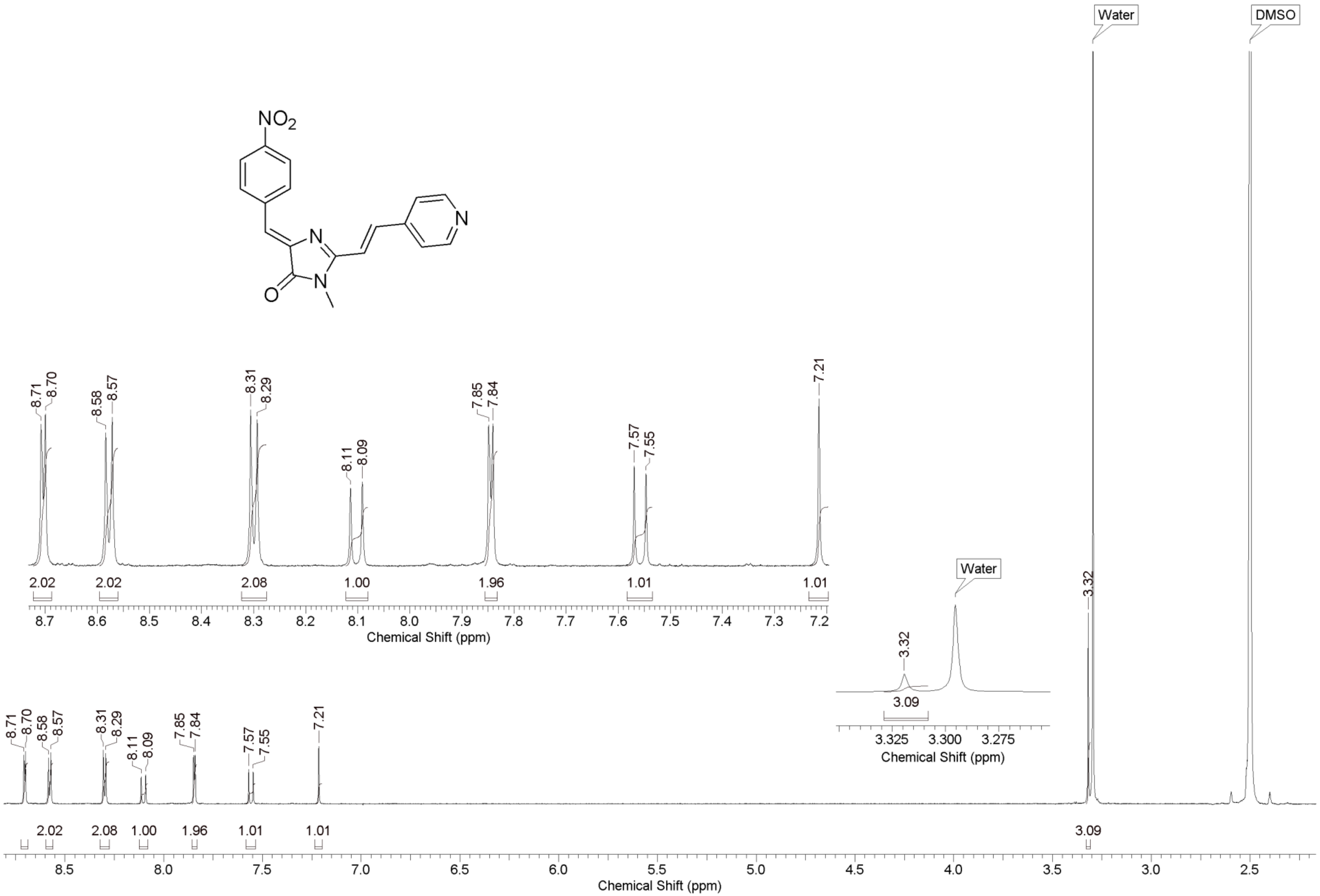

Water
DMSO
$NO_2$
N
N
N
O
8.71 8.70 8.58 8.57 8.31 8.29 8.11 8.09 7.85 7.84 7.57 7.55 7.21
2.02 2.02 2.08 1.00 1.96 1.01 1.01
Chemical Shift (ppm)
Water
3.32
3.09
3.325 3.300 3.275
Chemical Shift (ppm)
3.32
2.02 2.08 1.00 1.96 1.01 1.01
3.09
Chemical Shift (ppm)

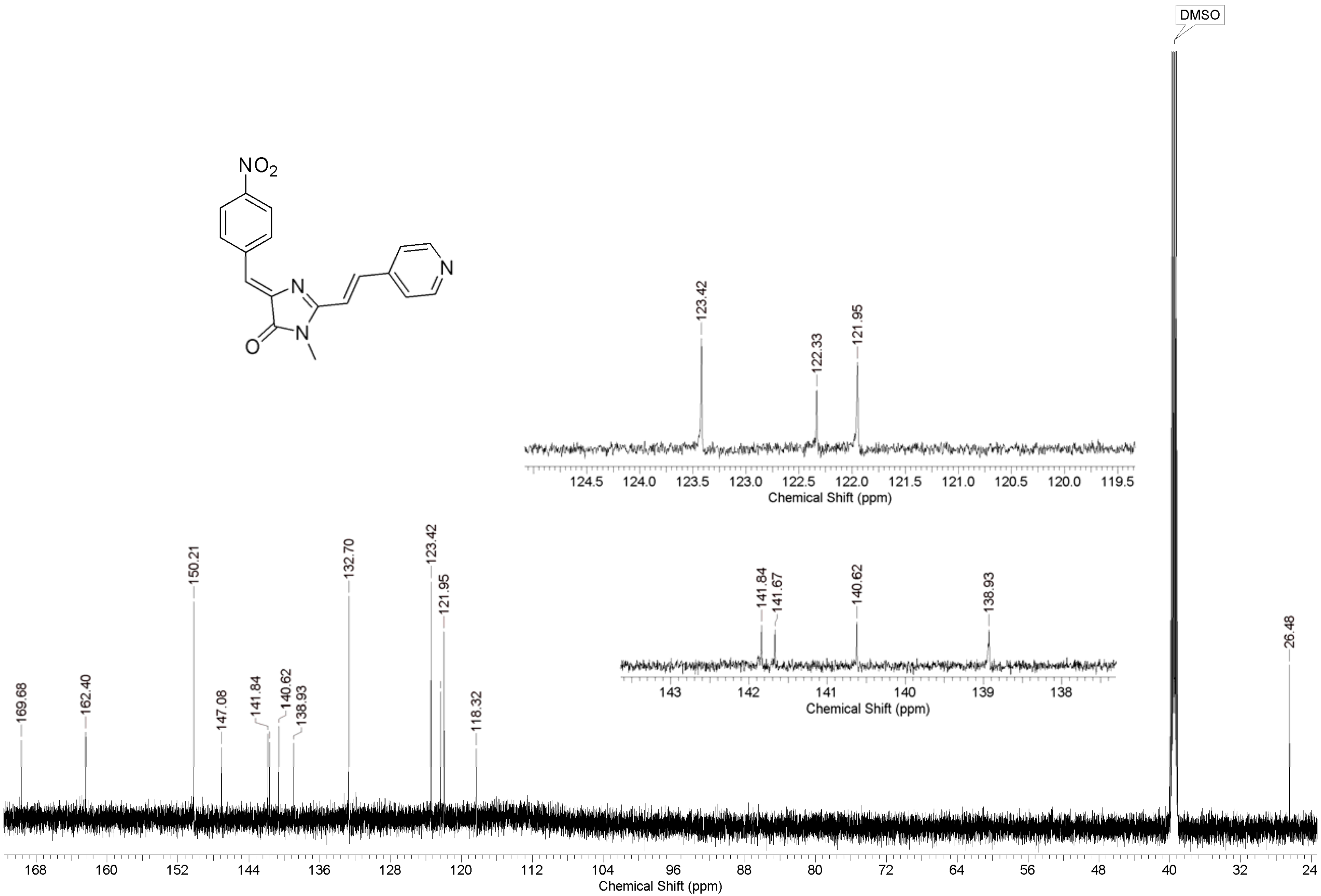

NO2
N
N
O
N
DMSO
123.42
122.33
121.95
124.5 124.0 123.5 123.0 122.5 122.0 121.5 121.0 120.5 120.0 119.5
Chemical Shift (ppm)
141.84
141.67
140.62
138.93
143 142 141 140 139 138
Chemical Shift (ppm)
169.68
162.40
150.21
147.08
141.84
140.62
138.93
132.70
123.42
121.95
118.32
26.48
168 160 152 144 136 128 120 112 104 96 88 80 72 64 56 48 40 32 24
Chemical Shift (ppm)

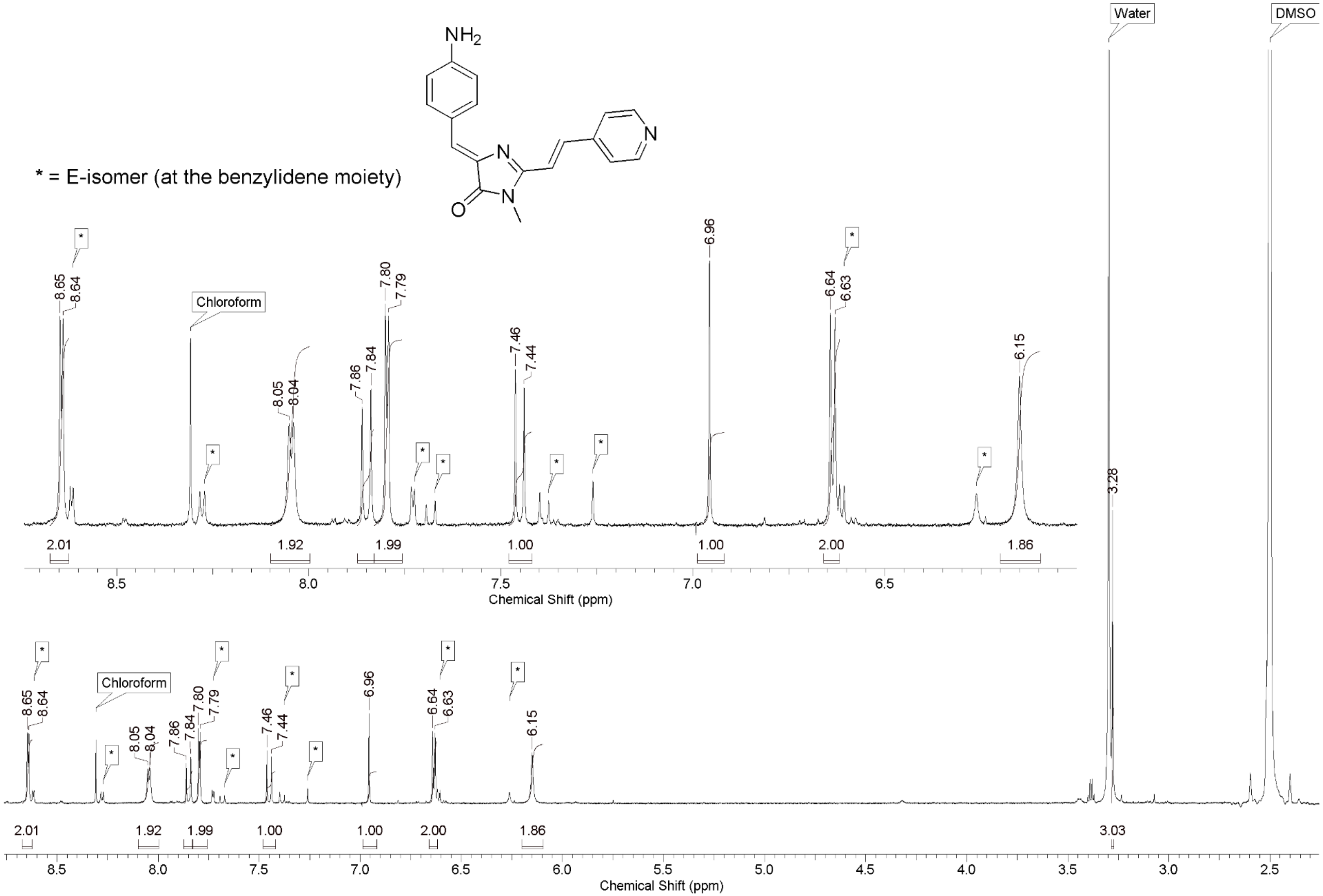
* = E-isomer (at the benzylidene moiety)
NH2
N
N
O
Water
DMSO
Chloroform
8.65
8.64
8.05
8.04
7.86
7.84
7.80
7.79
7.46
7.44
6.96
6.64
6.63
6.15
3.28
2.01
1.92
1.99
1.00
1.00
2.00
1.86
3.03
Chemical Shift (ppm)

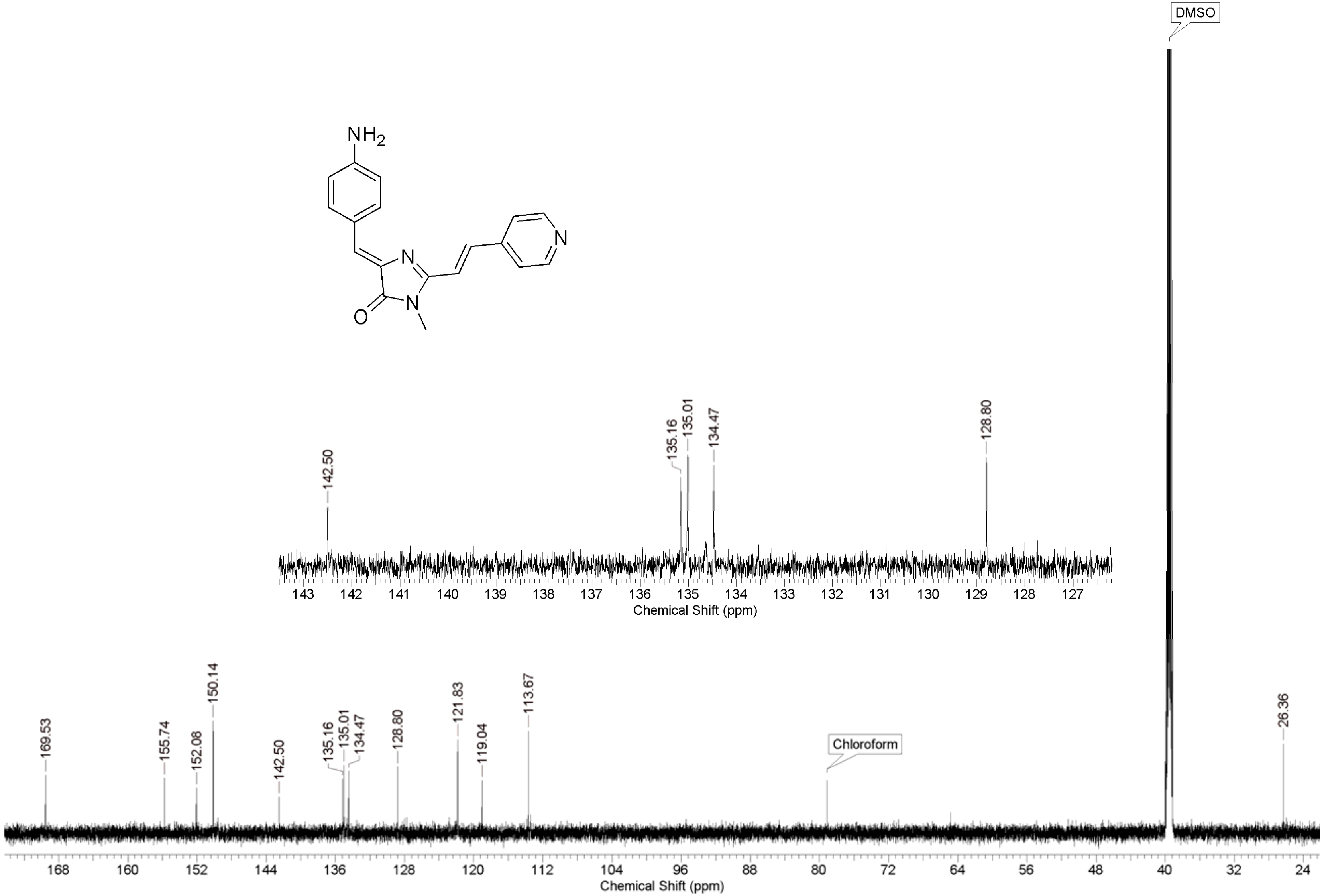
NH2
N
N
N
O
DMSO
142.50
135.16
135.01
134.47
128.80
143 142 141 140 139 138 137 136 135 134 133 132 131 130 129 128 127
Chemical Shift (ppm)
169.53
155.74
152.08
150.14
142.50
135.16
135.01
134.47
128.80
121.83
119.04
113.67
Chloroform
26.36
168 160 152 144 136 128 120 112 104 96 88 80 72 64 56 48 40 32 24
Chemical Shift (ppm)

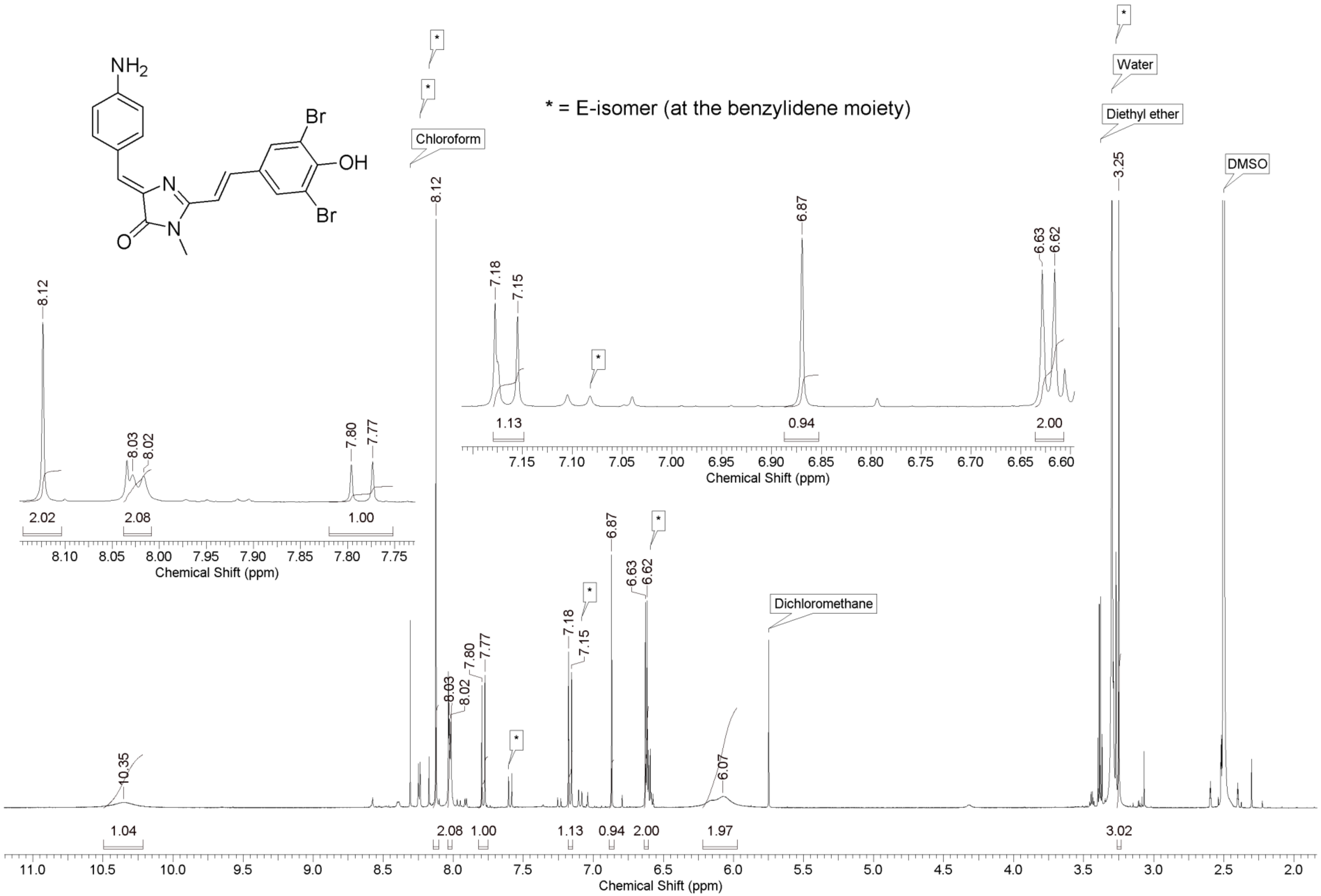
NH2
Br
OH
Br
N
O
N
* = E-isomer (at the benzylidene moiety)
Chloroform
Water
Diethyl ether
DMSO
Dichloromethane
Chemical Shift (ppm)

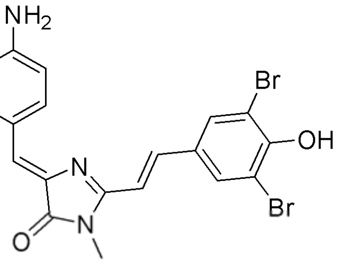
NH2
Br
OH
Br
N
O
N

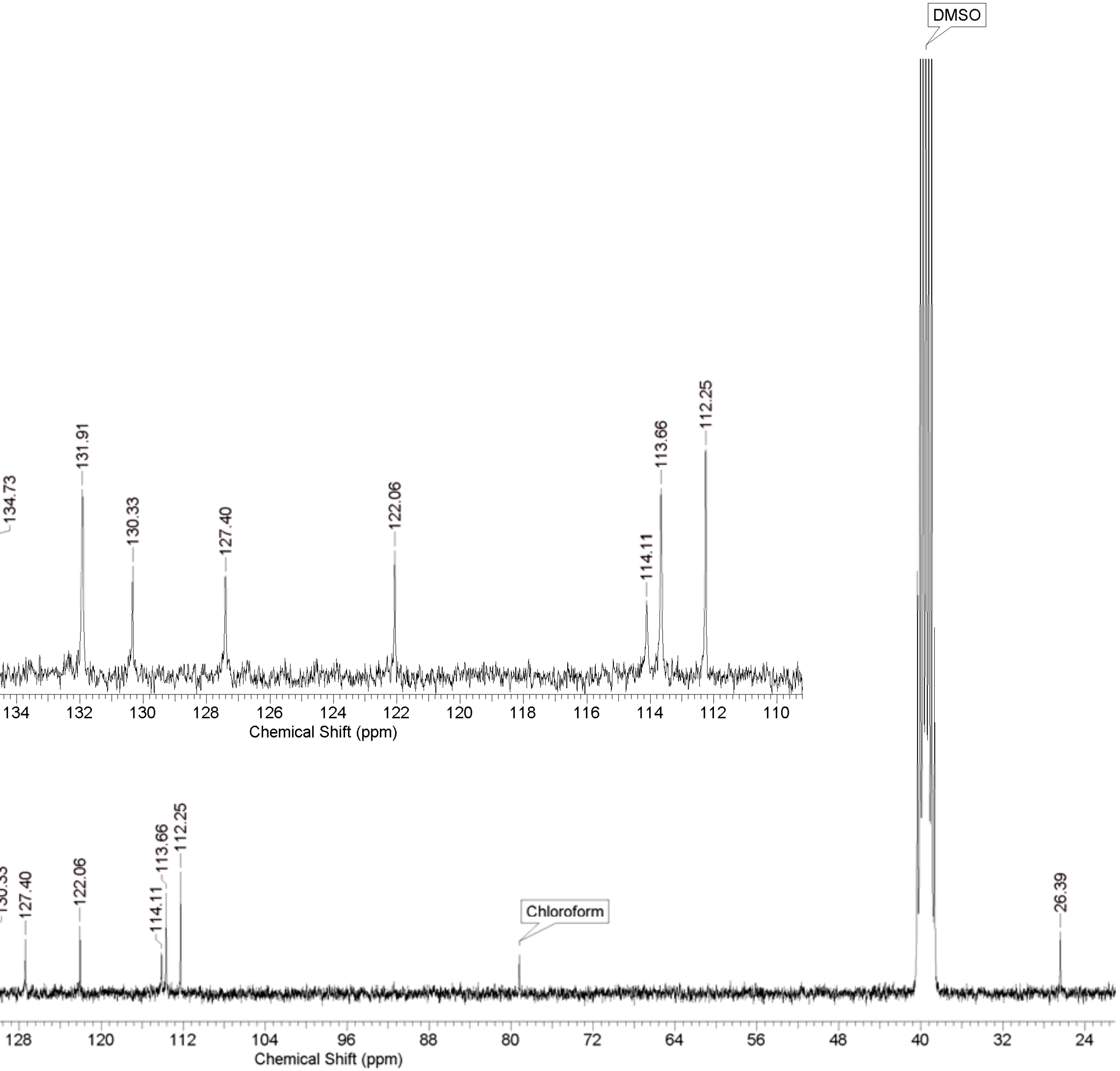
DMSO
135.49
134.80
134.73
131.91
130.33
127.40
122.06
114.11
113.66
112.25
Chemical Shift (ppm)
169.77
156.50
151.74
Chloroform
26.39
Chemical Shift (ppm)

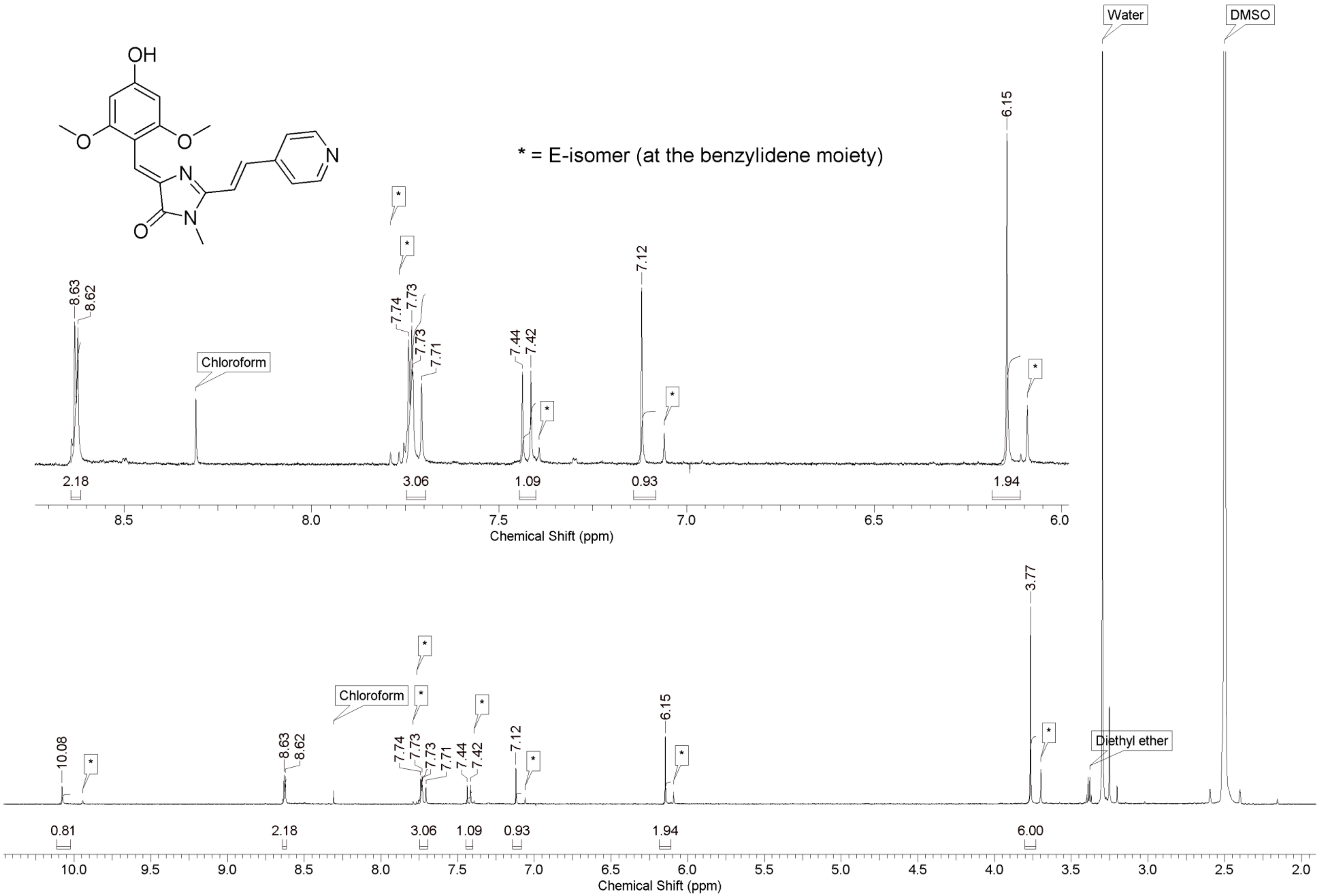
OH
O
N
O
N
N
* = E-isomer (at the benzylidene moiety)
Water
DMSO
Chloroform
Diethyl ether
8.63
8.62
7.74
7.73
7.73
7.71
7.44
7.42
7.12
6.15
10.08
3.77
2.18
3.06
1.09
0.93
1.94
0.81
6.00
Chemical Shift (ppm)
Chemical Shift (ppm)

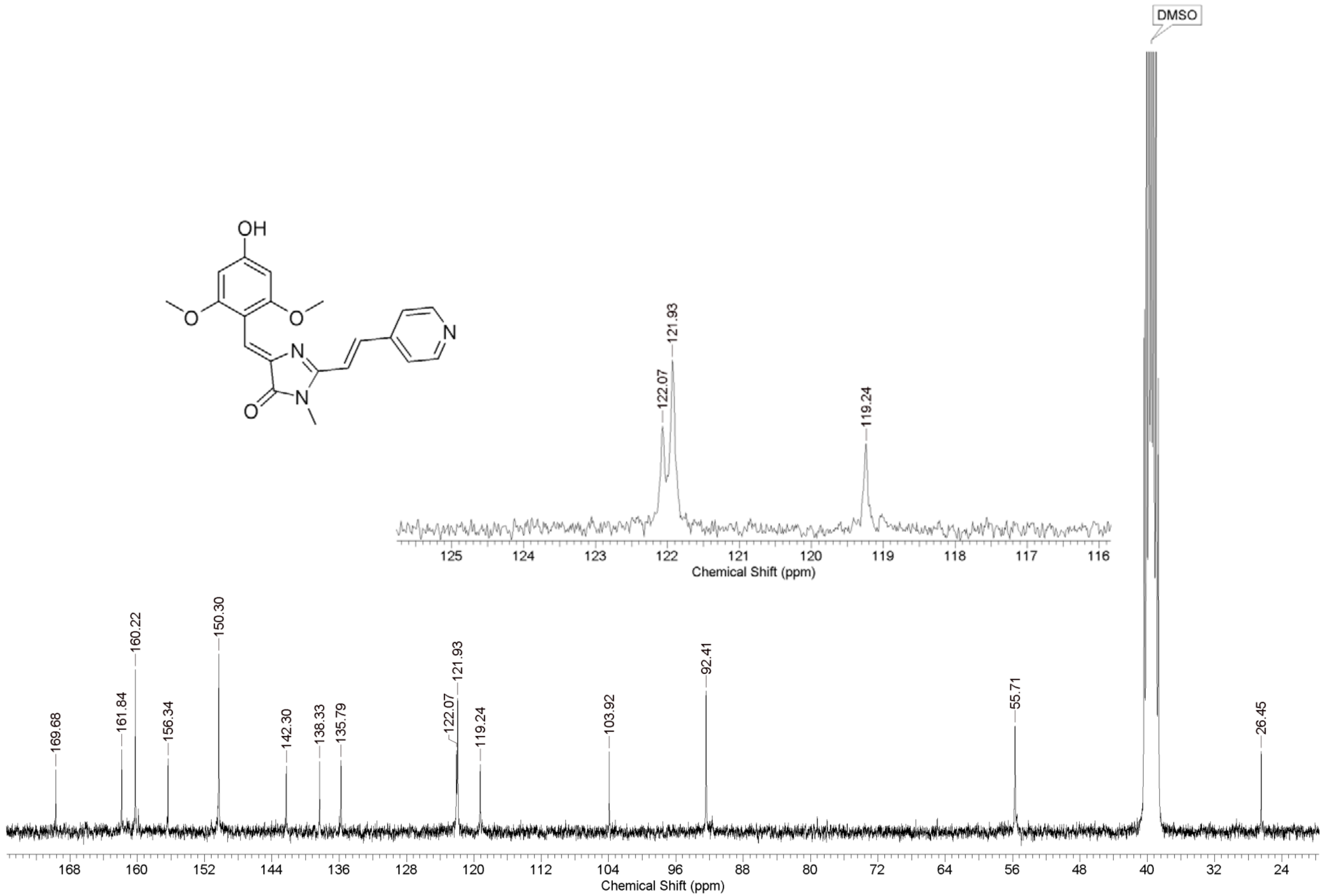
OH
O
N
N
O
N
O
DMSO
122.07
121.93
119.24
125
124
123
122
121
120
119
118
117
116
Chemical Shift (ppm)
169.68
161.84
160.22
156.34
150.30
142.30
138.33
135.79
122.07
121.93
119.24
103.92
92.41
55.71
26.45
168
160
152
144
136
128
120
112
104
96
88
80
72
64
56
48
40
32
24
Chemical Shift (ppm)

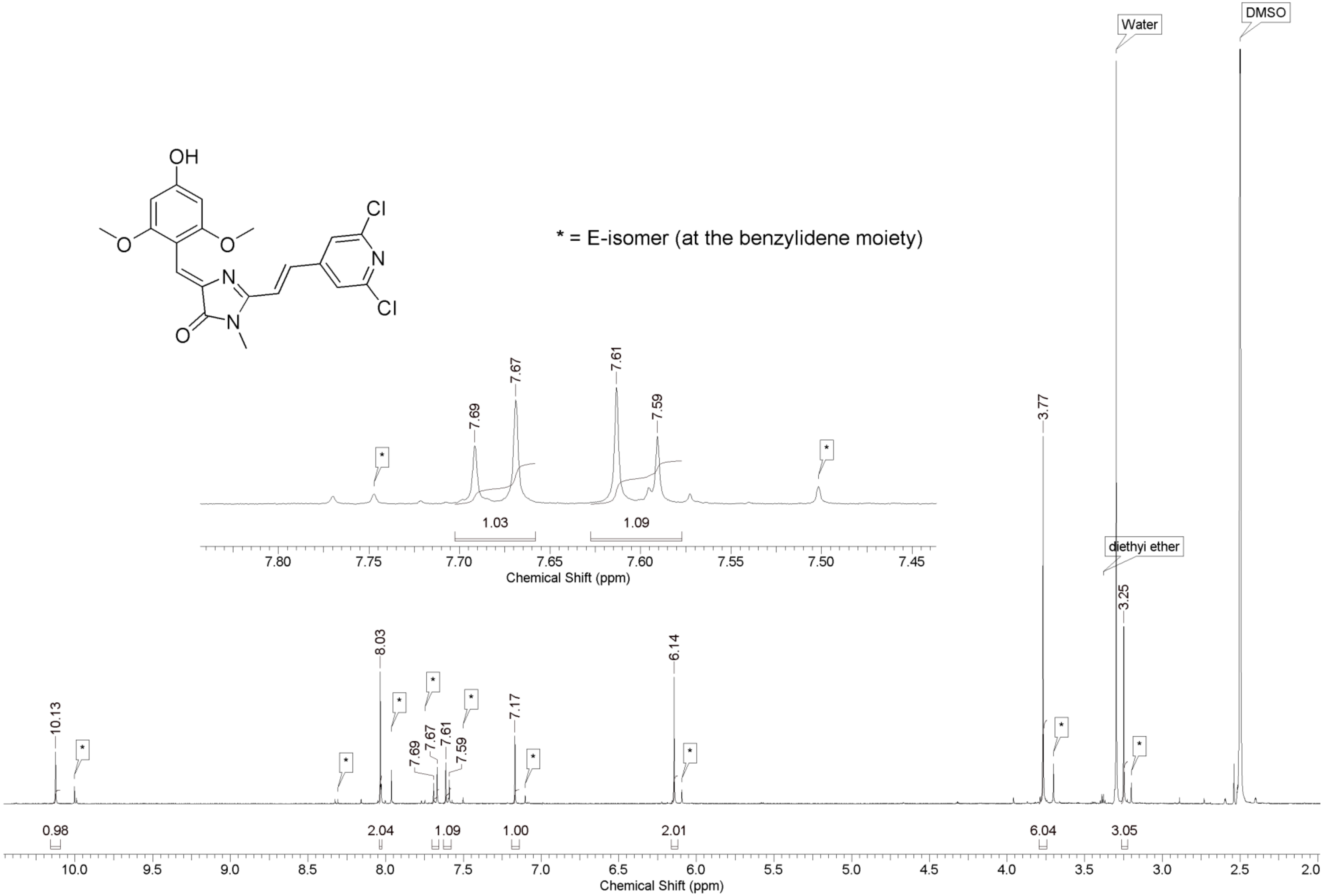

* = E-isomer (at the benzylidene moiety)
OH
O
N
Cl
Cl
Water
DMSO
diethyl ether
10.13
8.03
7.69
7.67
7.61
7.59
7.17
6.14
3.77
3.25
0.98
2.04
1.09
1.00
2.01
6.04
3.05
1.03
Chemical Shift (ppm)

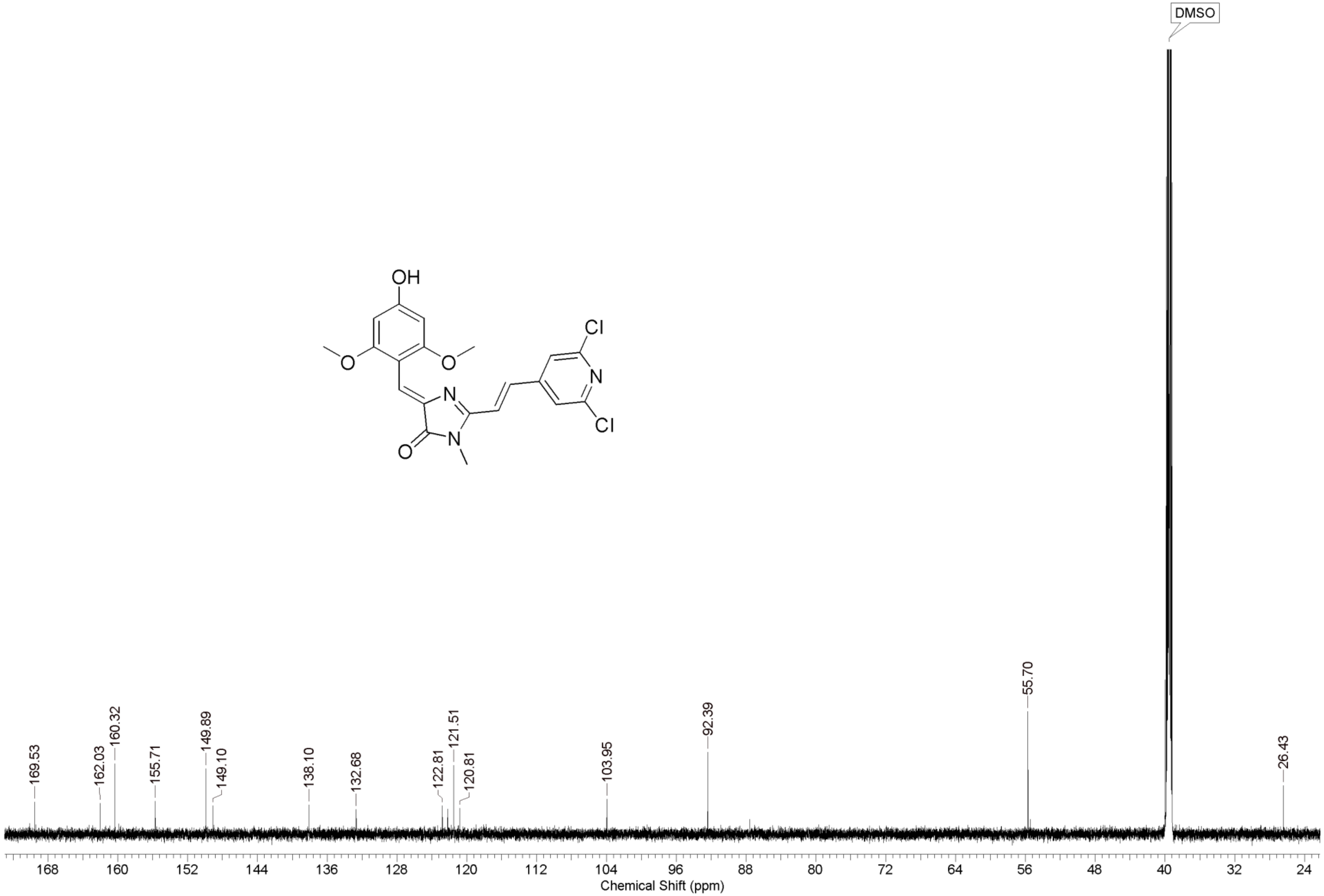
OH
O
O
N
N
O
Cl
N
Cl
DMSO
169.53
162.03
160.32
155.71
149.89
149.10
138.10
132.68
122.81
121.51
120.81
103.95
92.39
55.70
26.43
168
160
152
144
136
128
120
112
104
96
88
80
72
64
56
48
40
32
24
Chemical Shift (ppm)

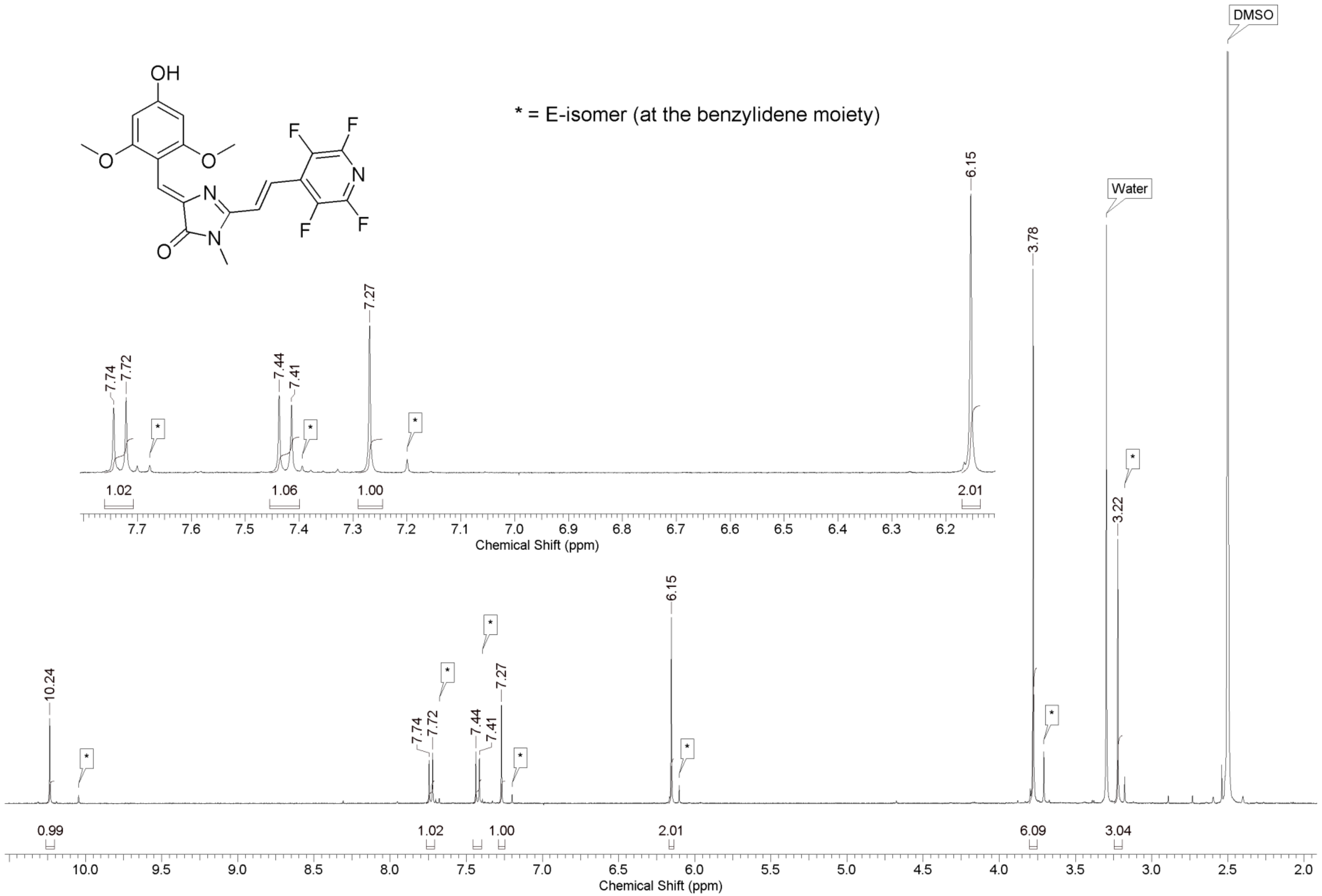
OH
O
N
F
F
N
F
F
O
N
* = E-isomer (at the benzylidene moiety)
DMSO
Water
10.24
7.74
7.72
7.44
7.41
7.27
6.15
3.78
3.22
0.99
1.02
1.06
1.00
2.01
6.09
3.04
Chemical Shift (ppm)

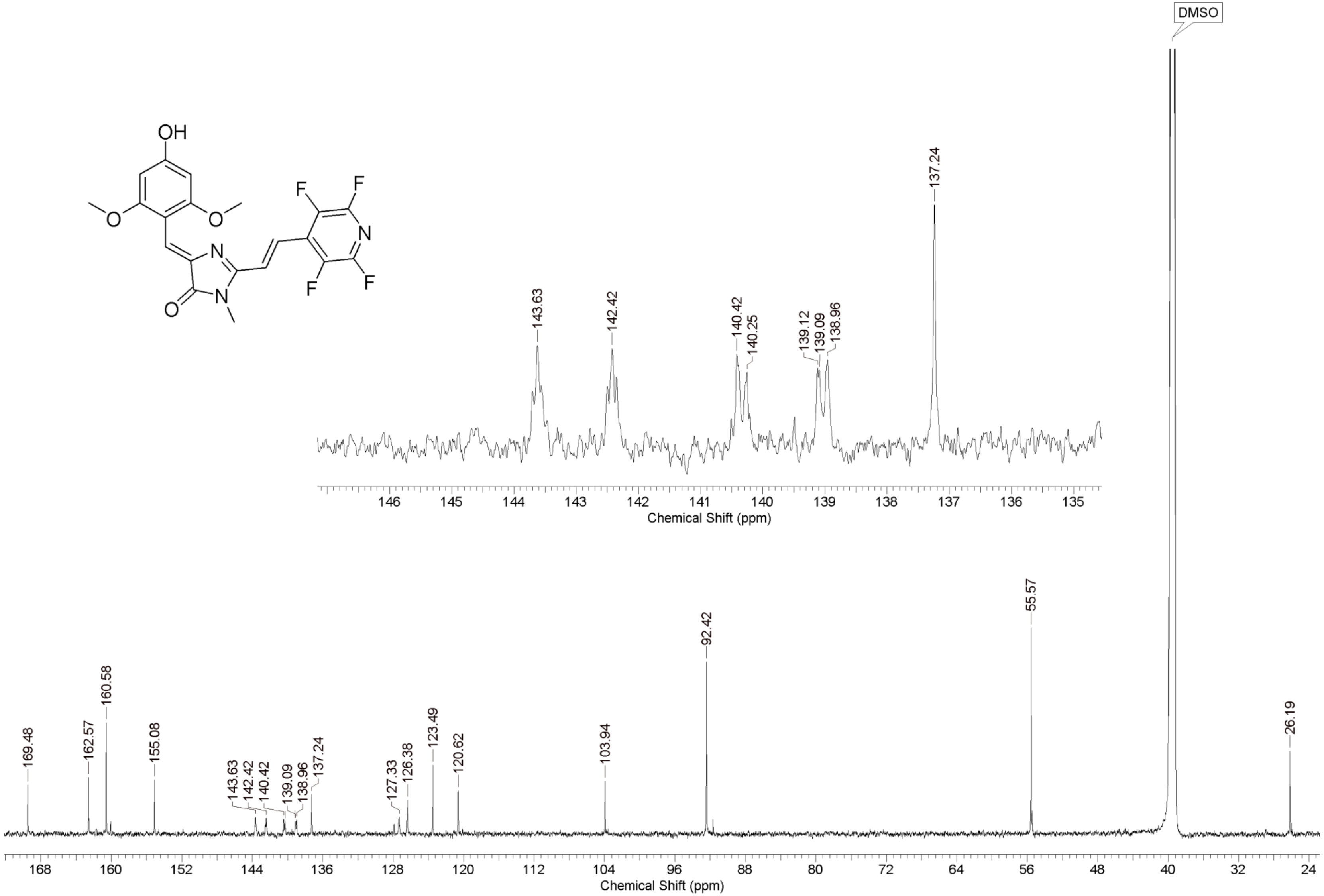
OH
O
O
N
N
O
F
F
N
F
F
DMSO
143.63
142.42
140.42
140.25
139.12
139.09
138.96
137.24
146 145 144 143 142 141 140 139 138 137 136 135
Chemical Shift (ppm)
169.48
162.57
160.58
155.08
143.63
142.42
140.42
139.09
138.96
137.24
127.33
126.38
123.49
120.62
103.94
92.42
55.57
26.19
168 160 152 144 136 128 120 112 104 96 88 80 72 64 56 48 40 32 24
Chemical Shift (ppm)

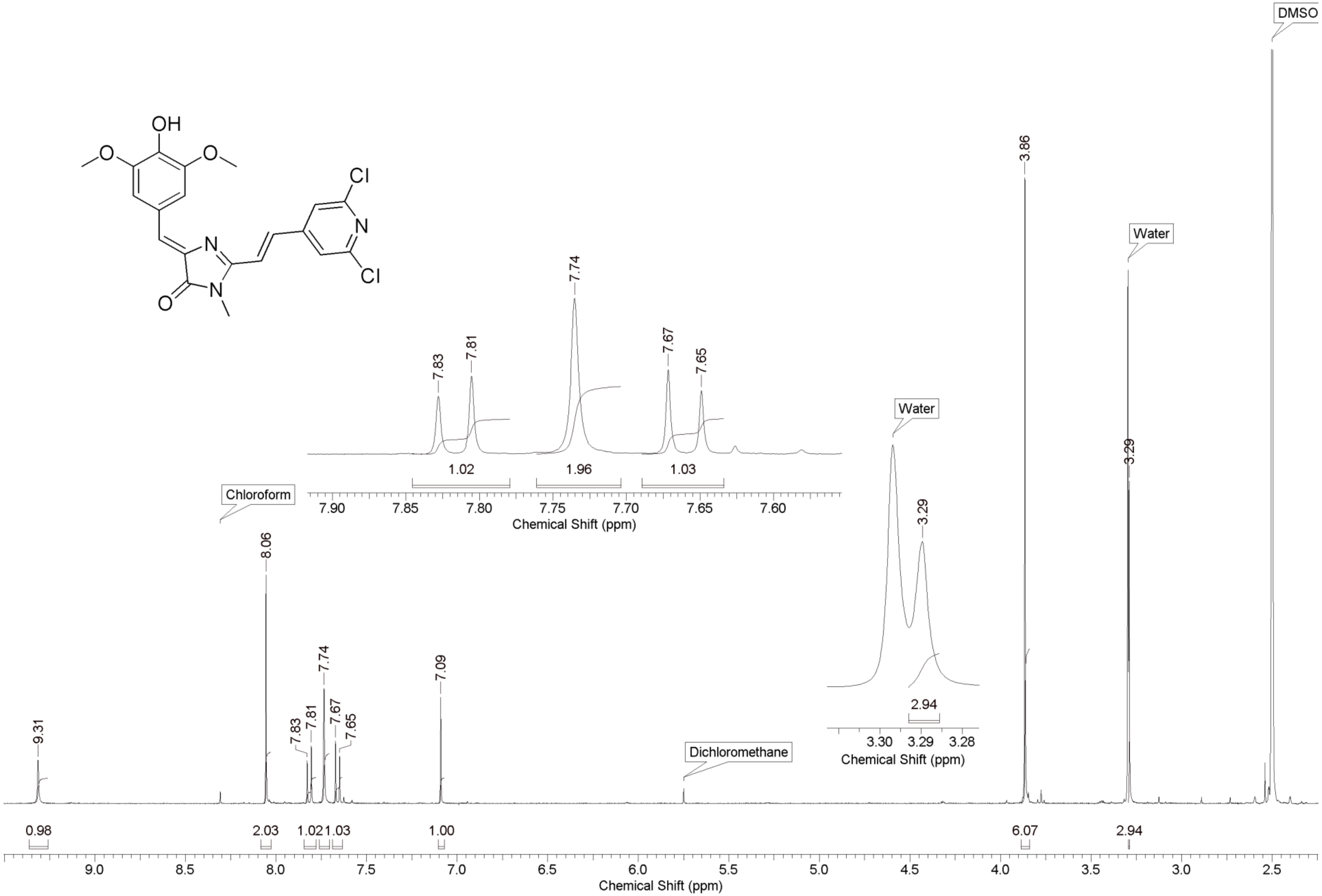
OH
O
O
Cl
N
Cl
N
N
O
DMSO
3.86
Water
7.74
7.83
7.81
7.67
7.65
1.02
1.96
1.03
7.90 7.85 7.80 7.75 7.70 7.65 7.60
Chemical Shift (ppm)
Water
3.29
3.29
2.94
3.30 3.29 3.28
Chemical Shift (ppm)
Chloroform
8.06
7.09
9.31
Dichloromethane
0.98
2.03
1.02
1.03
1.00
6.07
2.94
9.0 8.5 8.0 7.5 7.0 6.5 6.0 5.5 5.0 4.5 4.0 3.5 3.0 2.5
Chemical Shift (ppm)

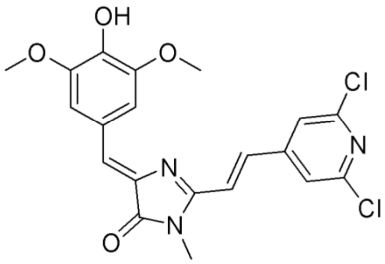

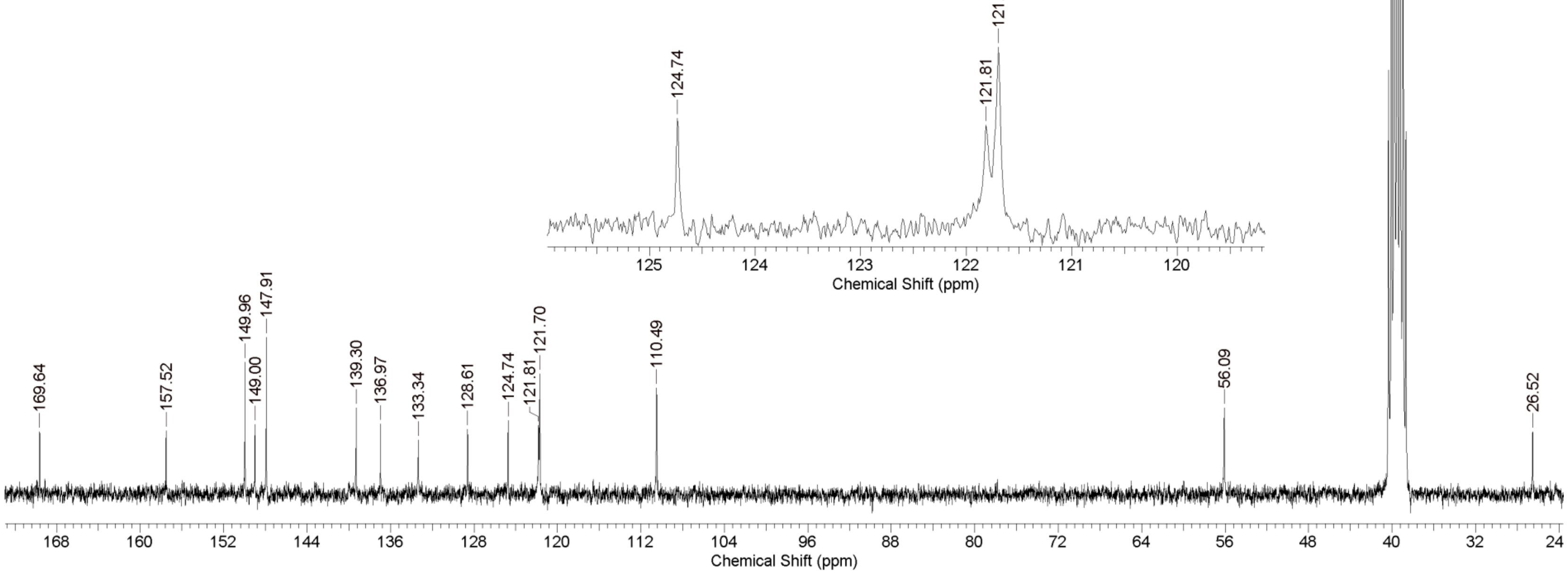

DMSO
169.64
157.52
149.96
149.00
147.91
139.30
136.97
133.34
128.61
124.74
121.81
121.70
110.49
56.09
26.52
168
160
152
144
136
128
120
112
104
96
88
80
72
64
56
48
40
32
24
Chemical Shift (ppm)
124.74
121.81
121.70
125
124
123
122
121
120
Chemical Shift (ppm)

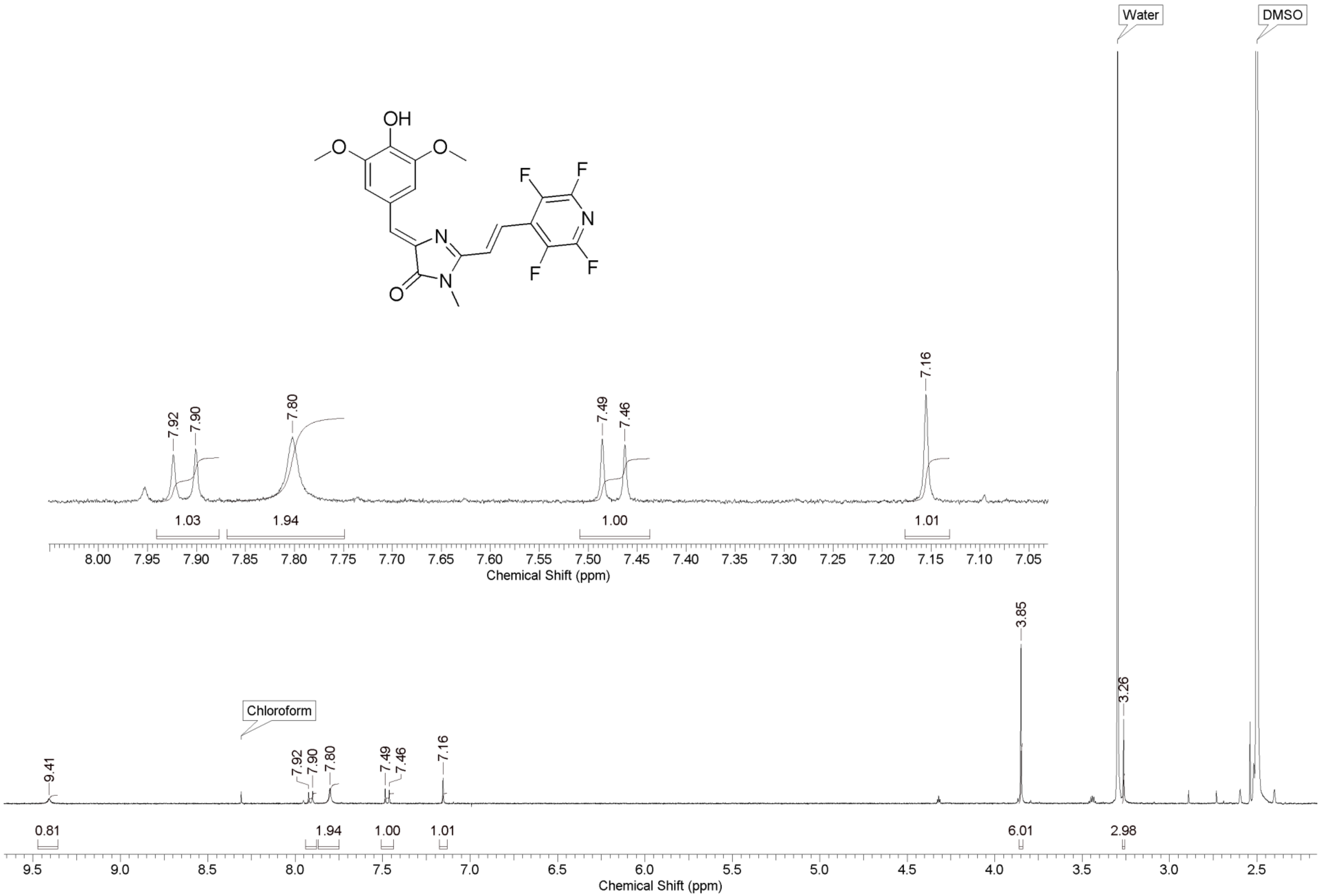

OH
O
O
F
F
N
F
F
N
N
O
Water
DMSO
7.92
7.90
7.80
7.49
7.46
7.16
1.03
1.94
1.00
1.01
8.00 7.95 7.90 7.85 7.80 7.75 7.70 7.65 7.60 7.55 7.50 7.45 7.40 7.35 7.30 7.25 7.20 7.15 7.10 7.05
Chemical Shift (ppm)
3.85
3.26
Chloroform
9.41
0.81
1.94
1.00
1.01
6.01
2.98
9.5 9.0 8.5 8.0 7.5 7.0 6.5 6.0 5.5 5.0 4.5 4.0 3.5 3.0 2.5
Chemical Shift (ppm)

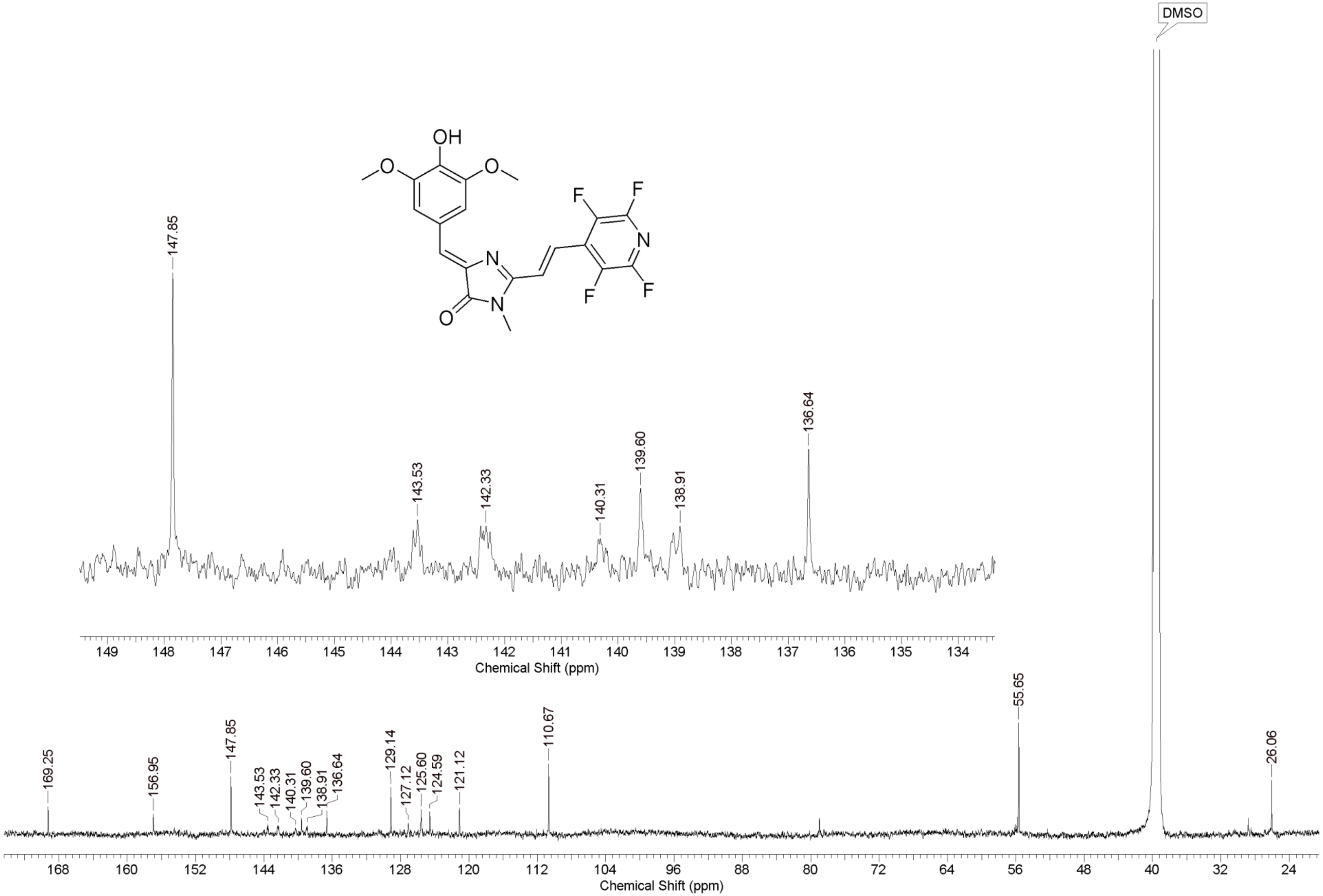
DMSO
OH
O
O
F
F
N
F
F
N
N
O
147.85
143.53
142.33
140.31
139.60
138.91
136.64
149 148 147 146 145 144 143 142 141 140 139 138 137 136 135 134
Chemical Shift (ppm)
169.25
156.95
147.85
143.53
142.33
140.31
139.60
138.91
136.64
129.14
127.12
125.60
124.59
121.12
110.67
55.65
26.06
168 160 152 144 136 128 120 112 104 96 88 80 72 64 56 48 40 32 24
Chemical Shift (ppm)

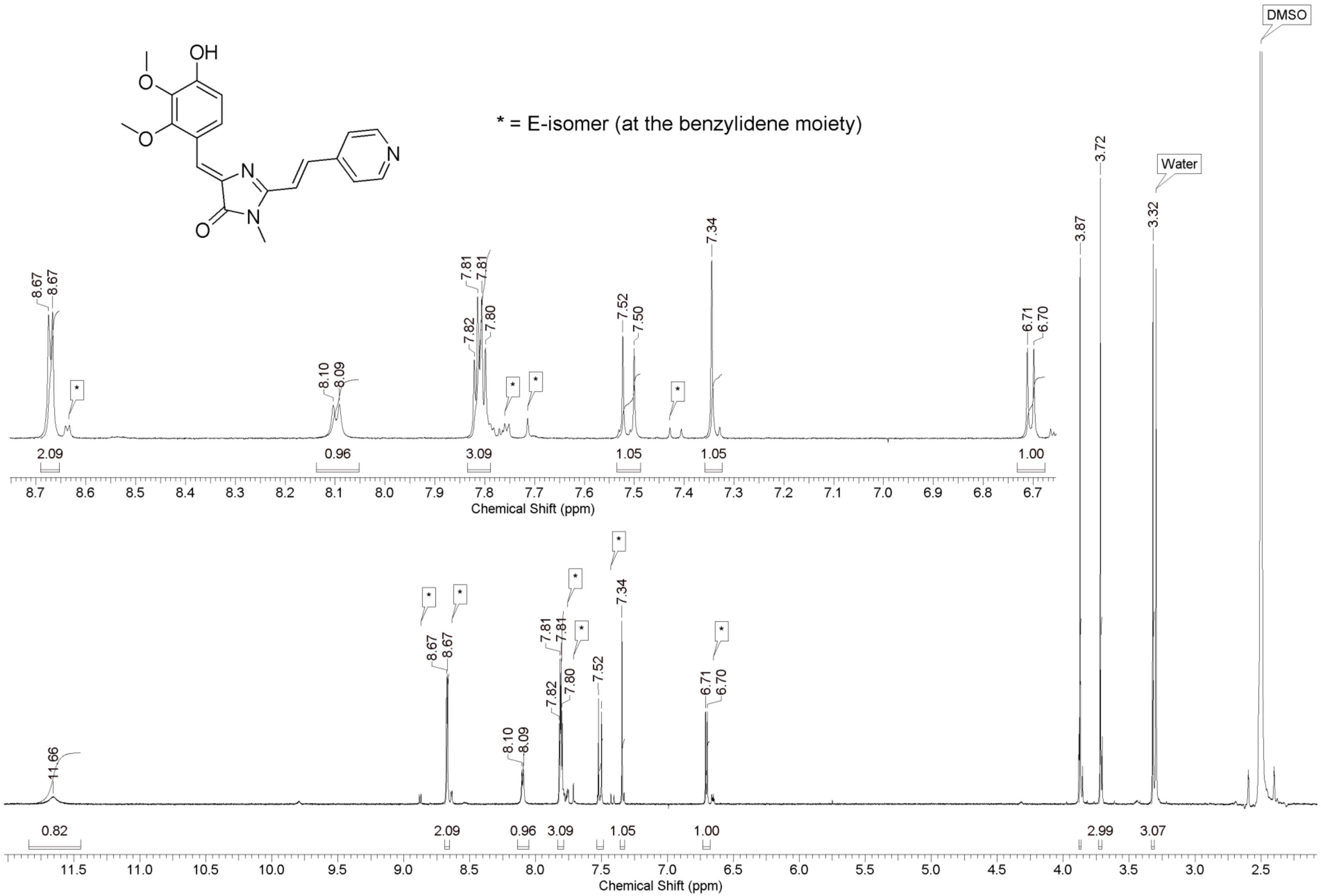
* = E-isomer (at the benzylidene moiety)
DMSO
Water
Chemical Shift (ppm)
Chemical Shift (ppm)

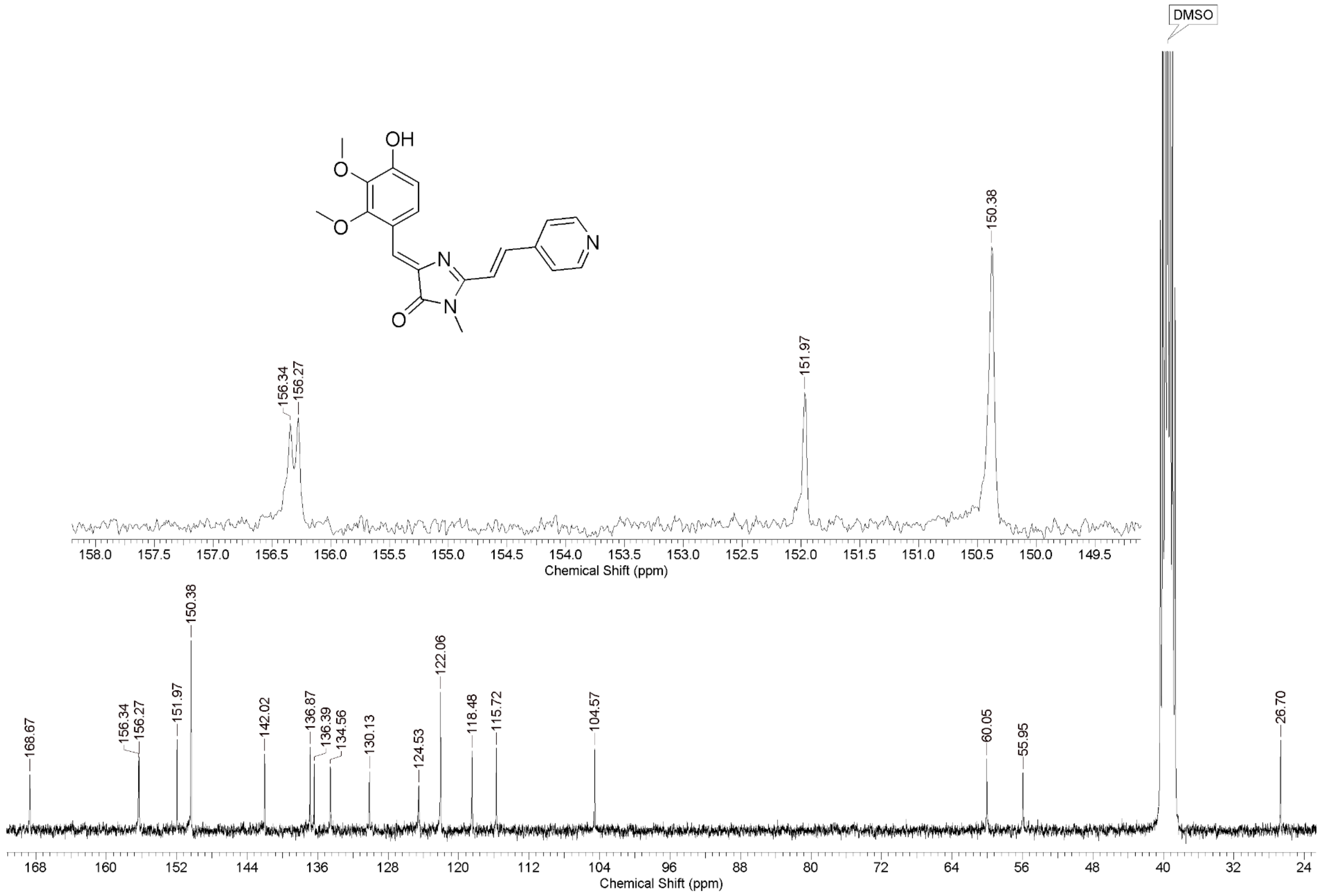

DMSO
OH
O
O
N
N
N
O
156.34
156.27
151.97
150.38
Chemical Shift (ppm)
168.67
142.02
136.87
136.39
134.56
130.13
124.53
122.06
118.48
115.72
104.57
60.05
55.95
26.70
Chemical Shift (ppm)

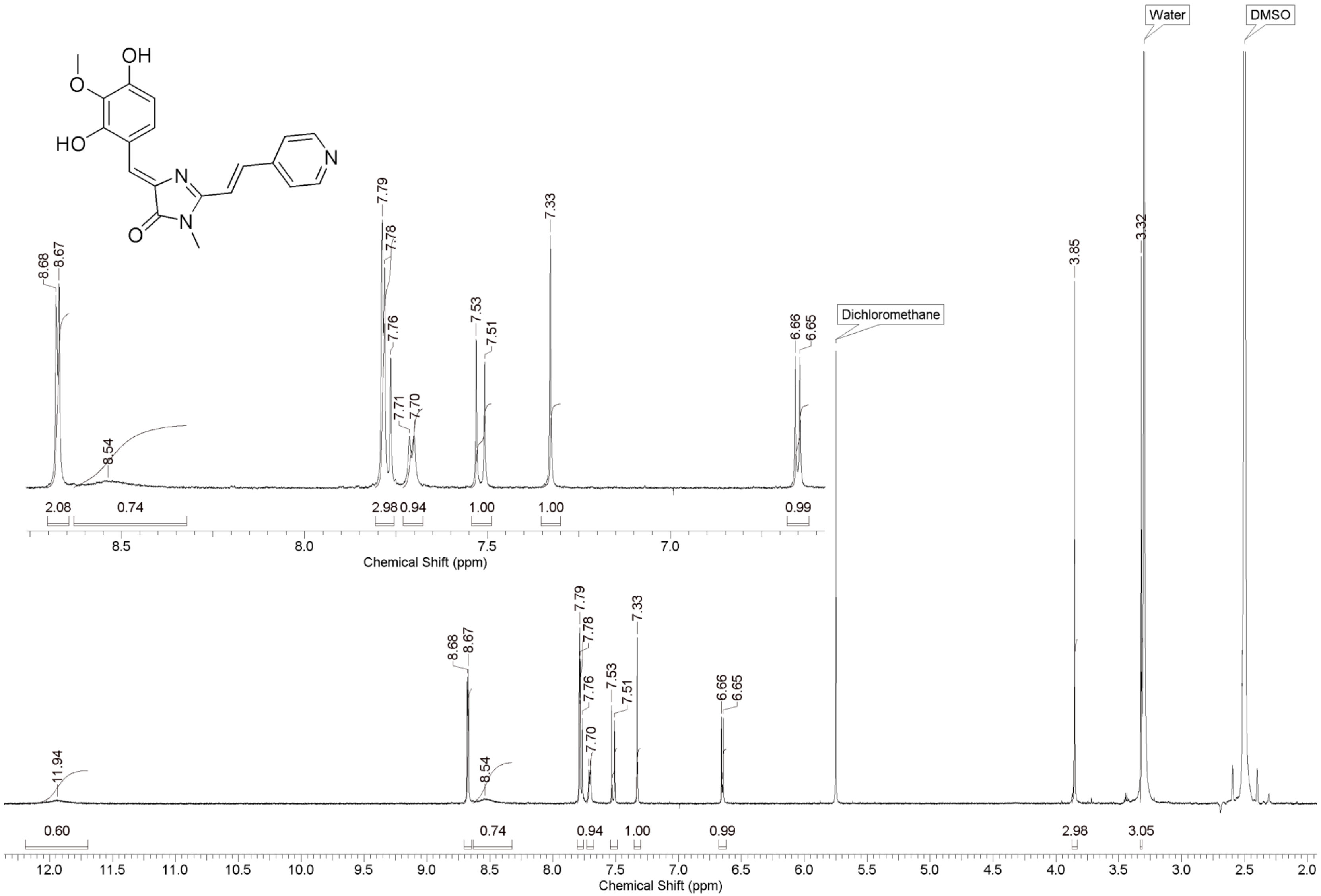
OH
O
HO
N
N
N
O
Water
DMSO
Dichloromethane
8.68
8.67
8.54
7.79
7.78
7.76
7.71
7.70
7.53
7.51
7.33
6.66
6.65
2.08
0.74
2.98
0.94
1.00
1.00
0.99
Chemical Shift (ppm)
11.94
3.85
3.32
0.60
0.74
0.94
1.00
0.99
2.98
3.05
Chemical Shift (ppm)

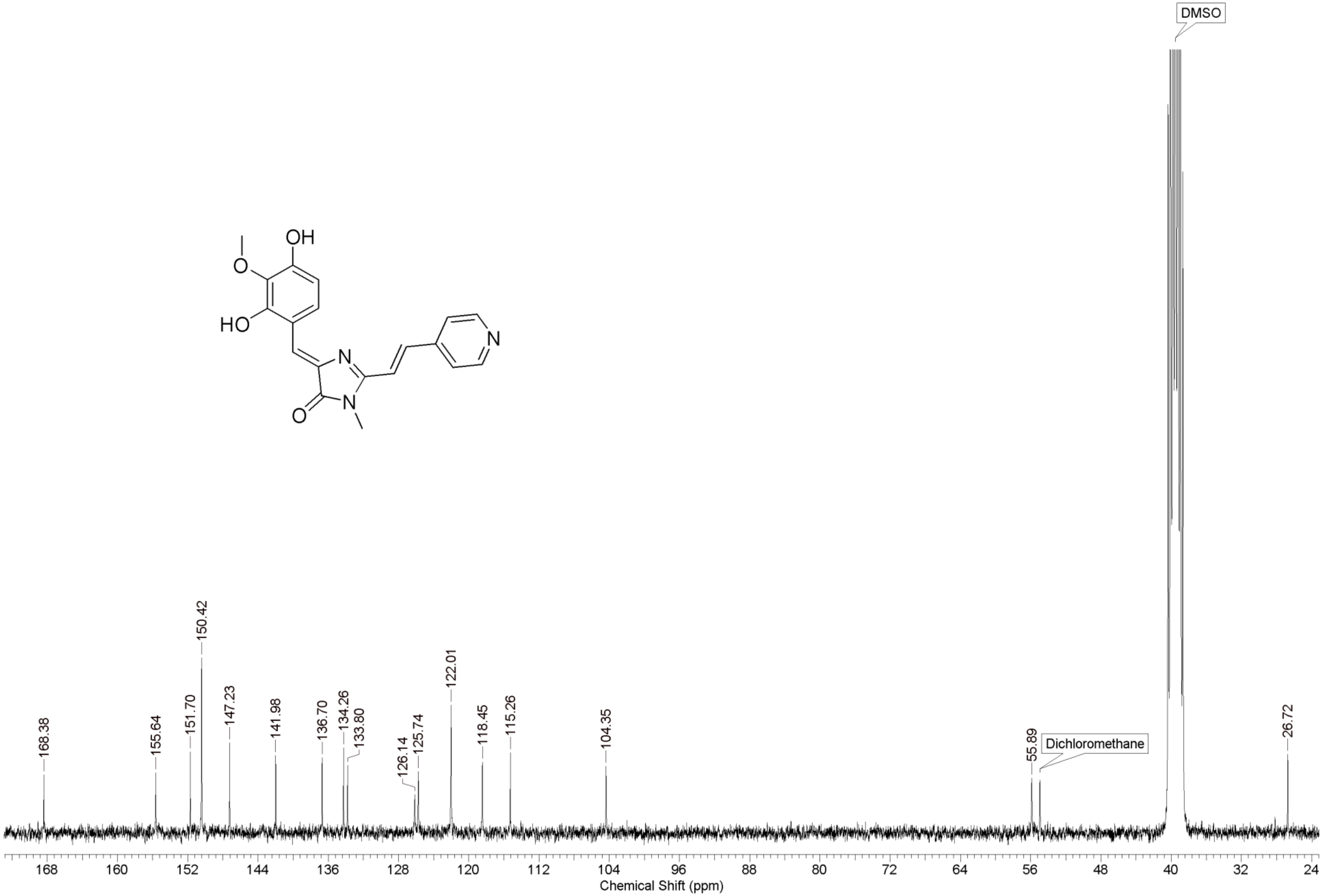
OH
O
HO
N
N
O
N
DMSO
Dichloromethane
168.38
155.64
151.70
150.42
147.23
141.98
136.70
134.26
133.80
126.14
125.74
122.01
118.45
115.26
104.35
55.89
26.72
168
160
152
144
136
128
120
112
104
96
88
80
72
64
56
48
40
32
24
Chemical Shift (ppm)

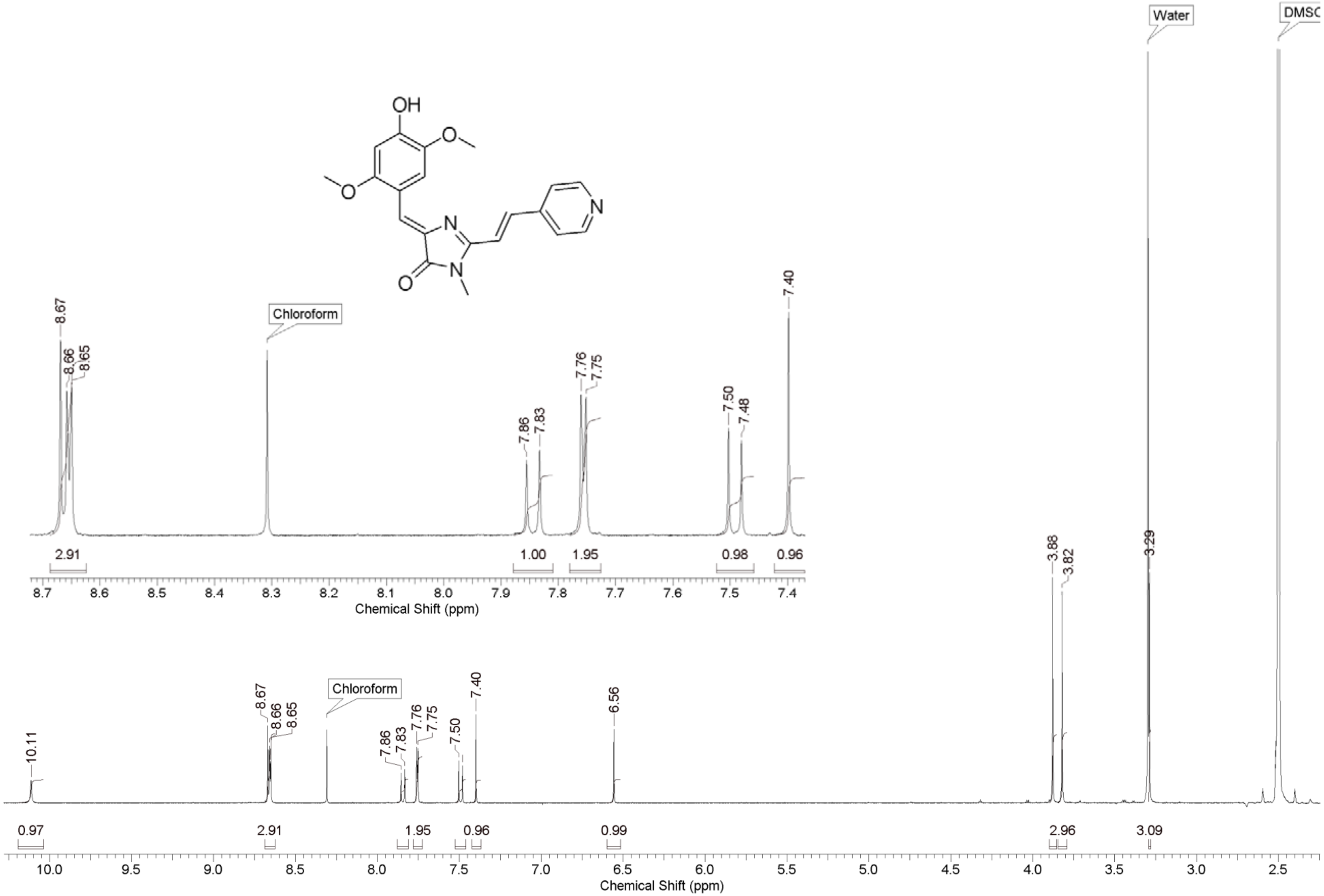
OH
O
O
N
N
O
N
Water
DMSO
Chloroform
10.11
8.67
8.66
8.65
7.86
7.83
7.76
7.75
7.50
7.48
7.40
6.56
3.88
3.82
3.29
0.97
2.91
1.00
1.95
0.98
0.96
0.99
2.96
3.09
Chemical Shift (ppm)

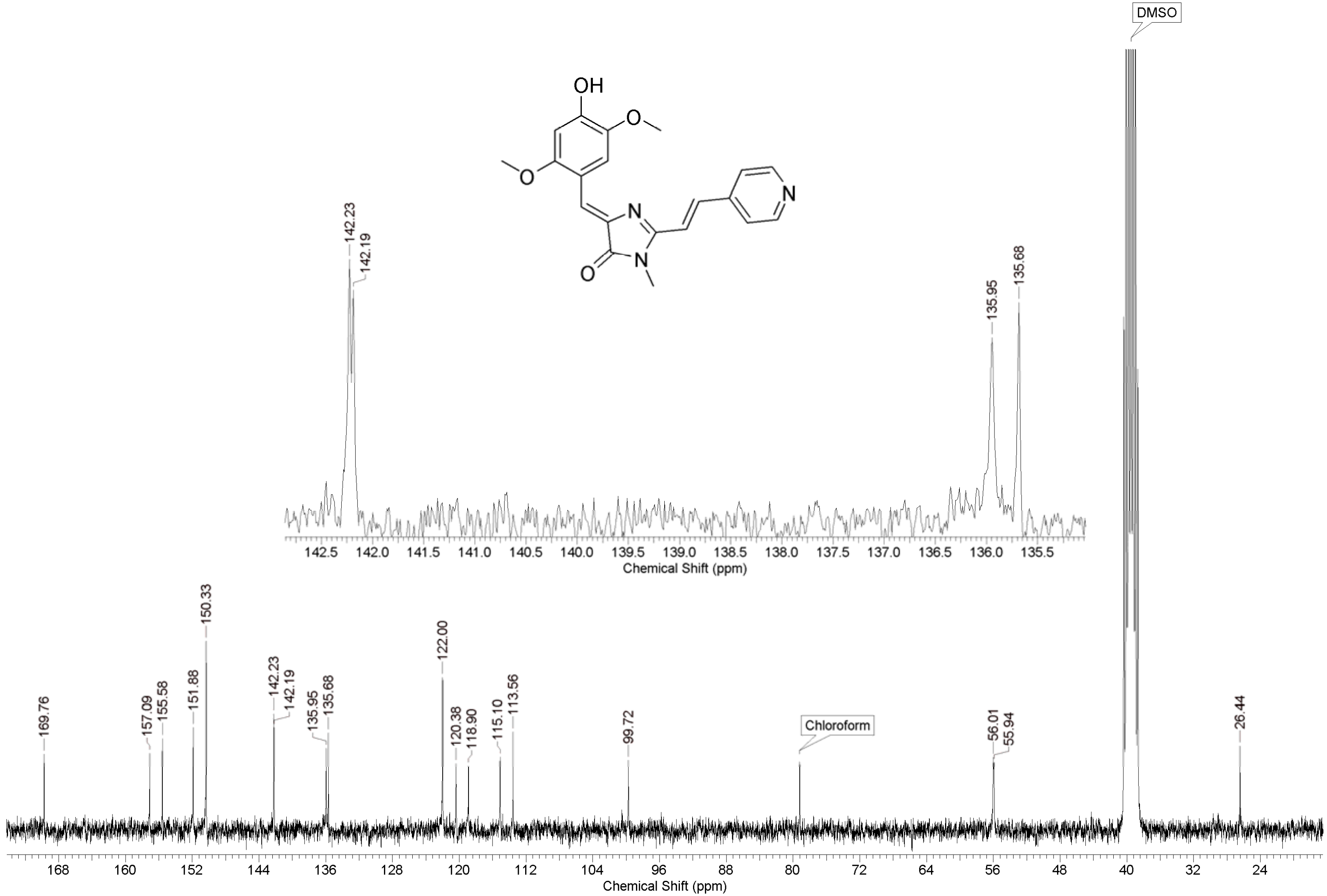
OH
O
N
N
O
N
DMSO
142.23
142.19
135.95
135.68
142.5 142.0 141.5 141.0 140.5 140.0 139.5 139.0 138.5 138.0 137.5 137.0 136.5 136.0 135.5
Chemical Shift (ppm)
169.76
157.09
155.58
151.88
150.33
142.23
142.19
135.95
135.68
122.00
120.38
118.90
115.10
113.56
99.72
Chloroform
56.01
55.94
26.44
168 160 152 144 136 128 120 112 104 96 88 80 72 64 56 48 40 32 24
Chemical Shift (ppm)

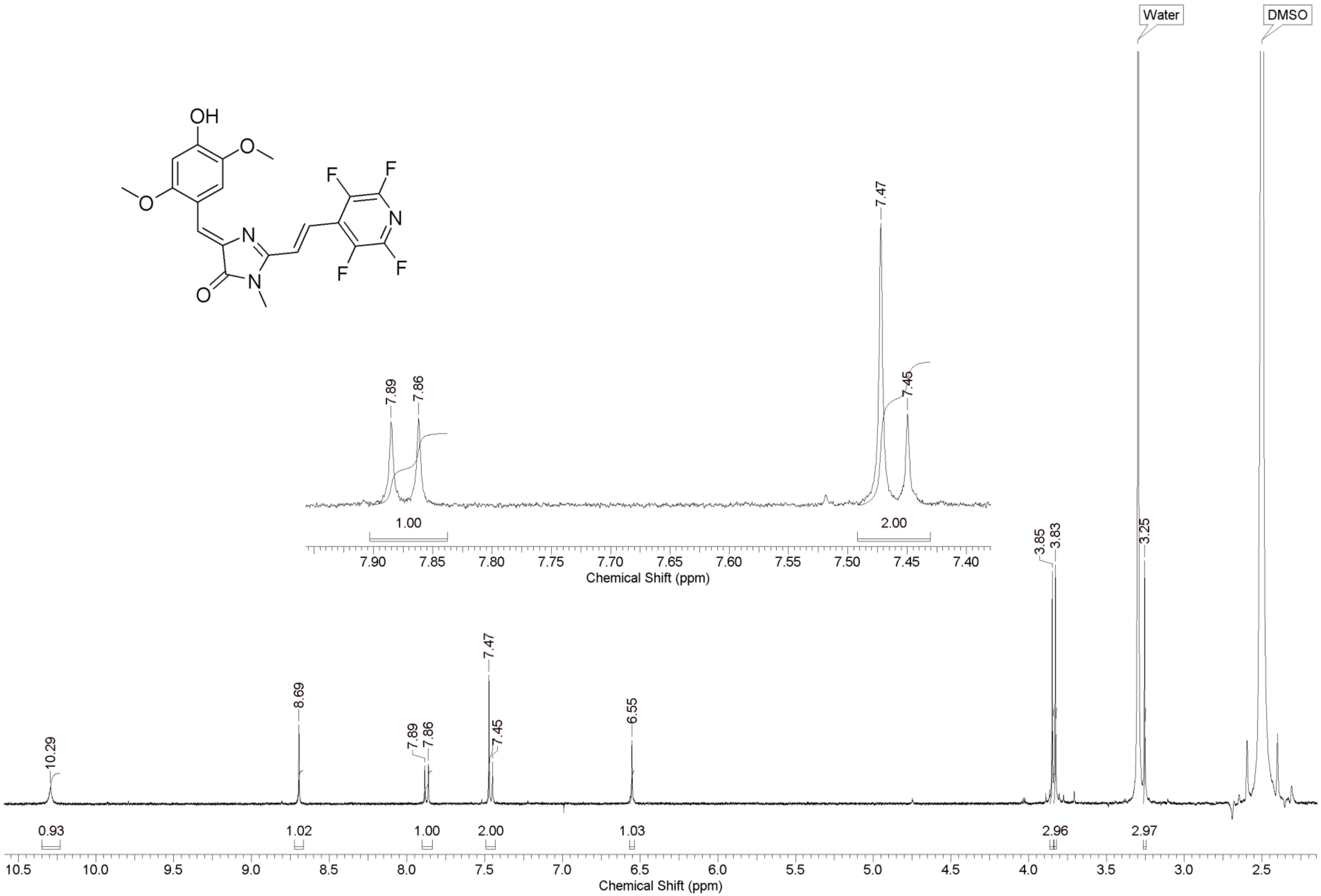
OH
F
F
N
F
F
N
N
O
Water
DMSO
7.89
7.86
7.47
7.45
1.00
2.00
7.90 7.85 7.80 7.75 7.70 7.65 7.60 7.55 7.50 7.45 7.40
Chemical Shift (ppm)
10.29
8.69
7.89
7.86
7.47
7.45
6.55
3.85
3.83
3.25
0.93
1.02
1.00
2.00
1.03
2.96
2.97
10.5 10.0 9.5 9.0 8.5 8.0 7.5 7.0 6.5 6.0 5.5 5.0 4.5 4.0 3.5 3.0 2.5
Chemical Shift (ppm)

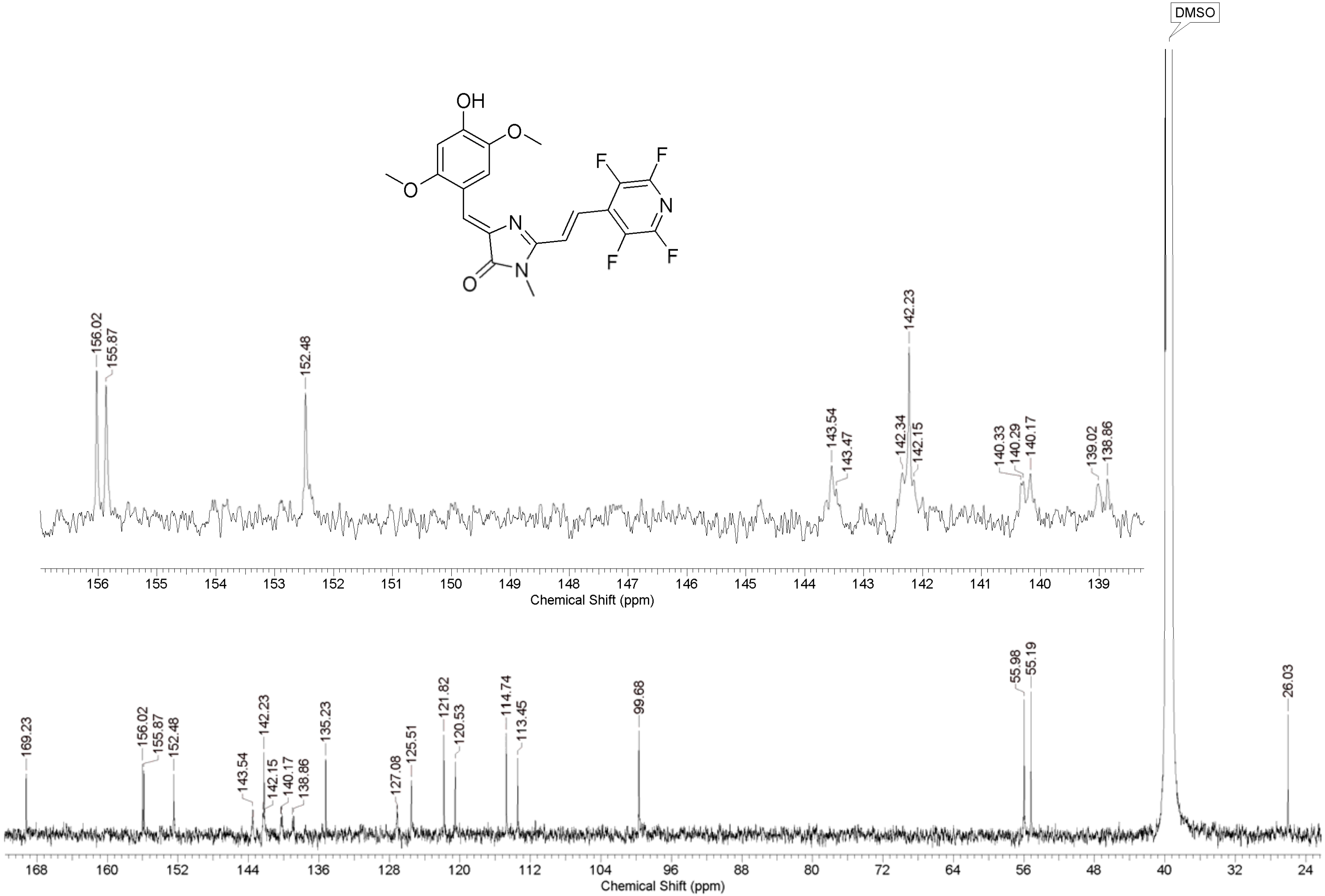

OH
O
O
N
N
O
F
F
N
F
F
DMSO
156.02
155.87
152.48
143.54
143.47
142.34
142.23
142.15
140.33
140.29
140.17
139.02
138.86
Chemical Shift (ppm)
169.23
156.02
155.87
152.48
143.54
142.23
142.15
140.17
138.86
135.23
127.08
125.51
121.82
120.53
114.74
113.45
99.68
55.98
55.19
26.03
Chemical Shift (ppm)

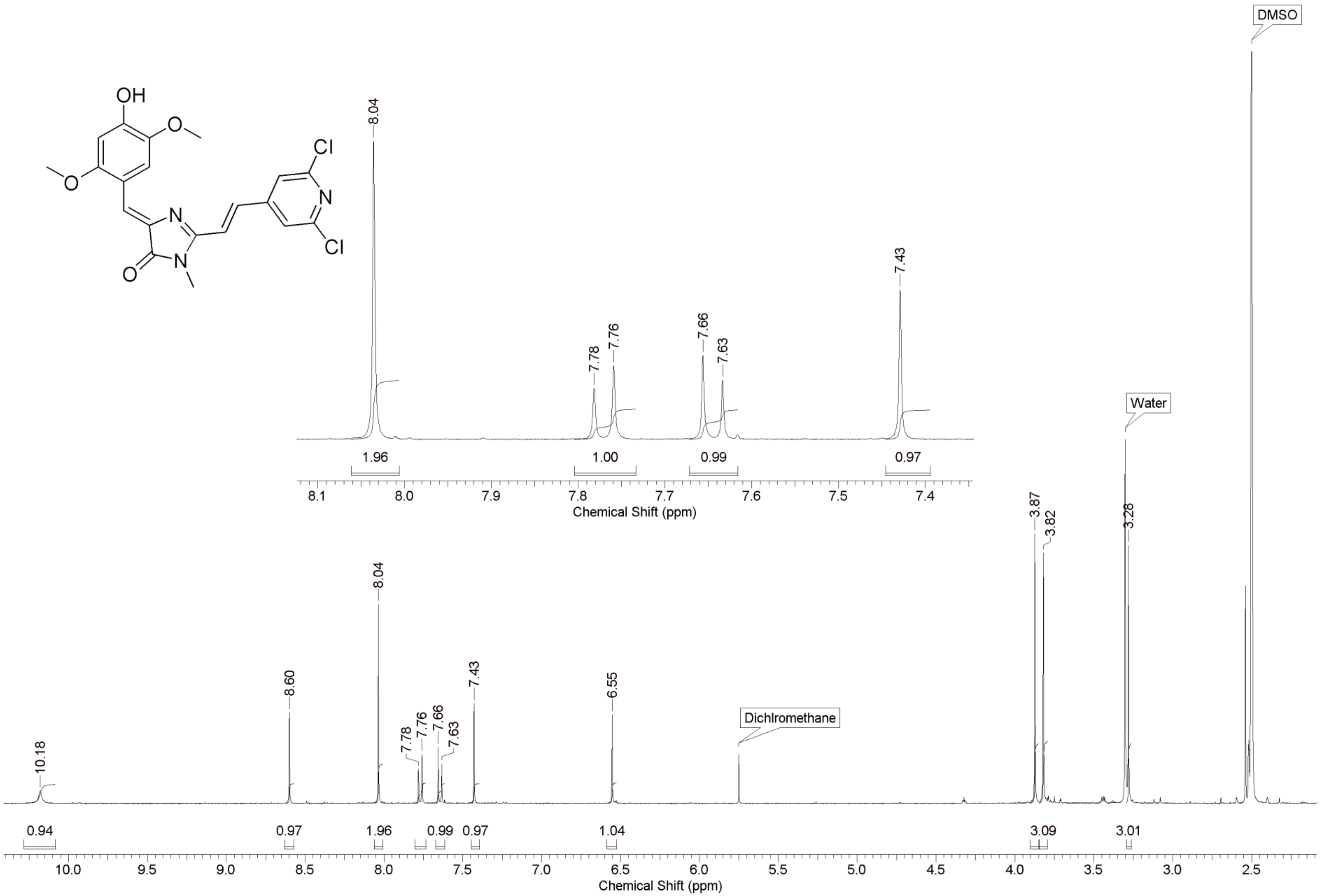
OH
Cl
N
Cl
O
N
N
8.04
7.78
7.76
7.66
7.63
7.43
1.96
1.00
0.99
0.97
Chemical Shift (ppm)
DMSO
Water
3.87
3.82
3.28
10.18
8.60
6.55
Dichlromethane
0.94
0.97
1.04
3.09
3.01
Chemical Shift (ppm)

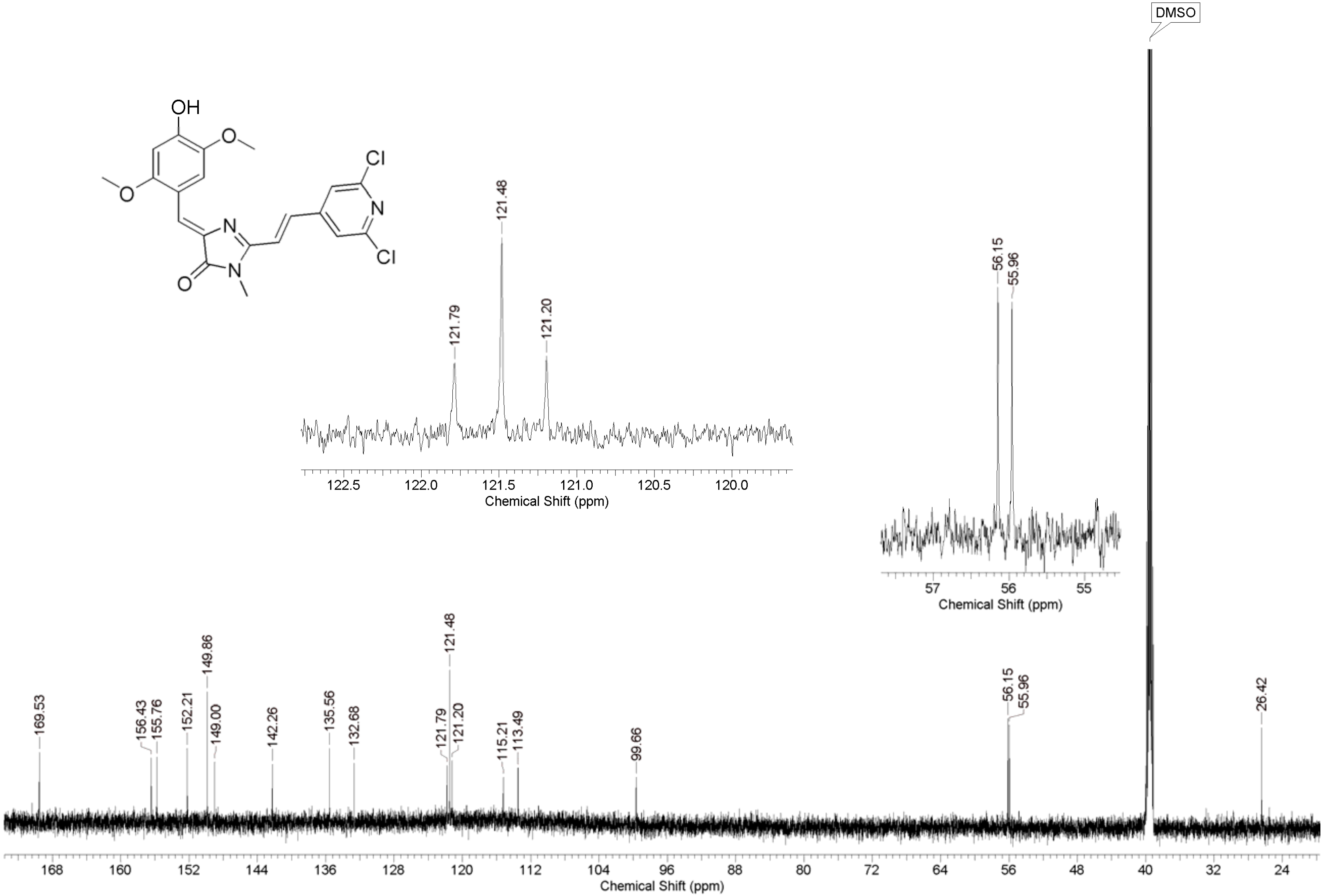
OH
O
O
N
N
O
Cl
N
Cl
121.48
121.79
121.20
122.5
122.0
121.5
121.0
120.5
120.0
Chemical Shift (ppm)
56.15
55.96
57
56
55
Chemical Shift (ppm)
DMSO
169.53
156.43
155.76
152.21
149.86
149.00
142.26
135.56
132.68
121.79
121.48
121.20
115.21
113.49
99.66
56.15
55.96
26.42
168
160
152
144
136
128
120
112
104
96
88
80
72
64
56
48
40
32
24
Chemical Shift (ppm)

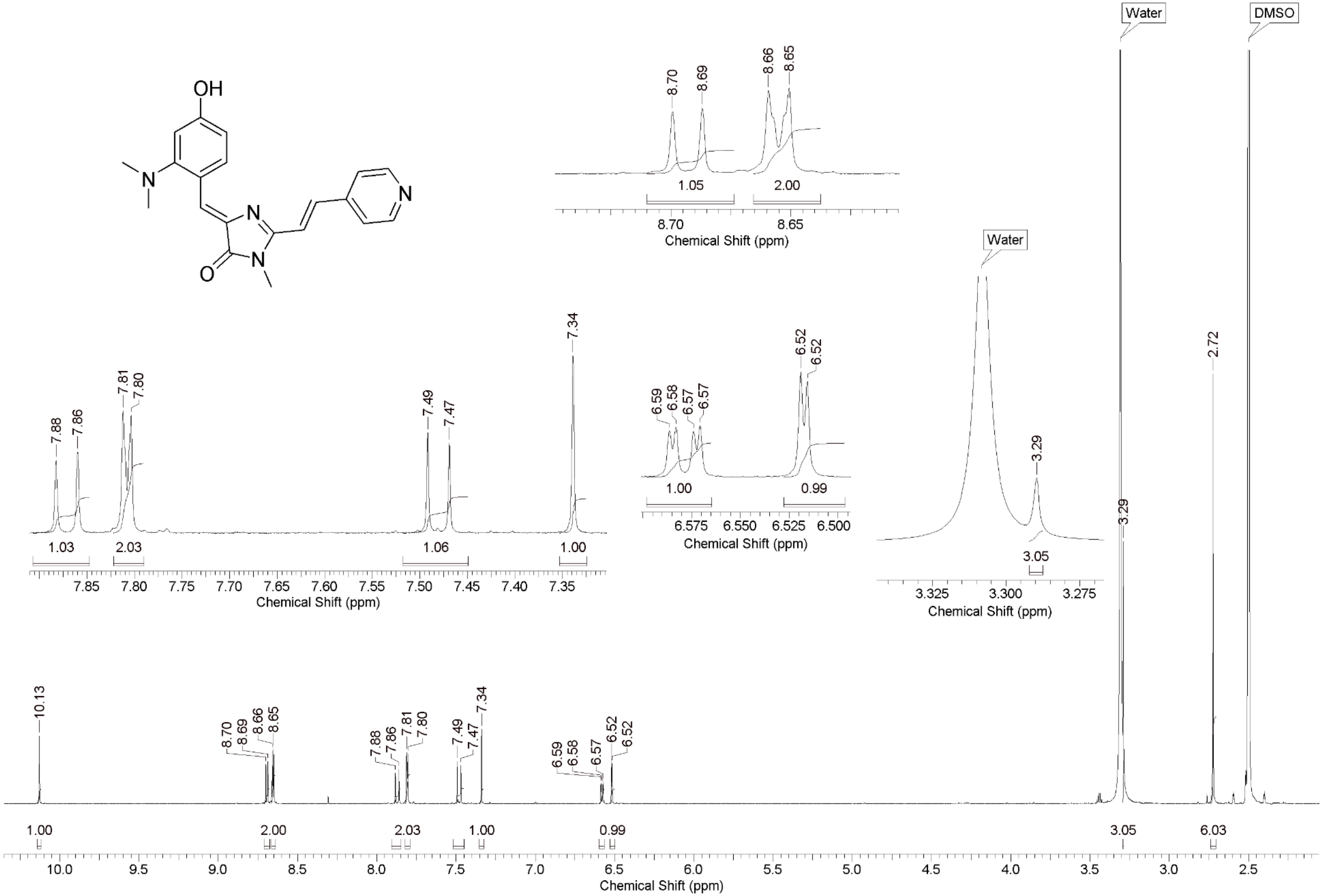
OH
N
N
N
N
O
Water
DMSO
8.70
8.69
8.66
8.65
1.05
2.00
Chemical Shift (ppm)
7.88
7.86
7.81
7.80
7.49
7.47
7.34
1.03
2.03
1.06
1.00
6.59
6.58
6.57
6.57
6.52
6.52
1.00
0.99
3.29
3.05
2.72
10.13
1.00
2.00
2.03
1.00
0.99
3.05
6.03

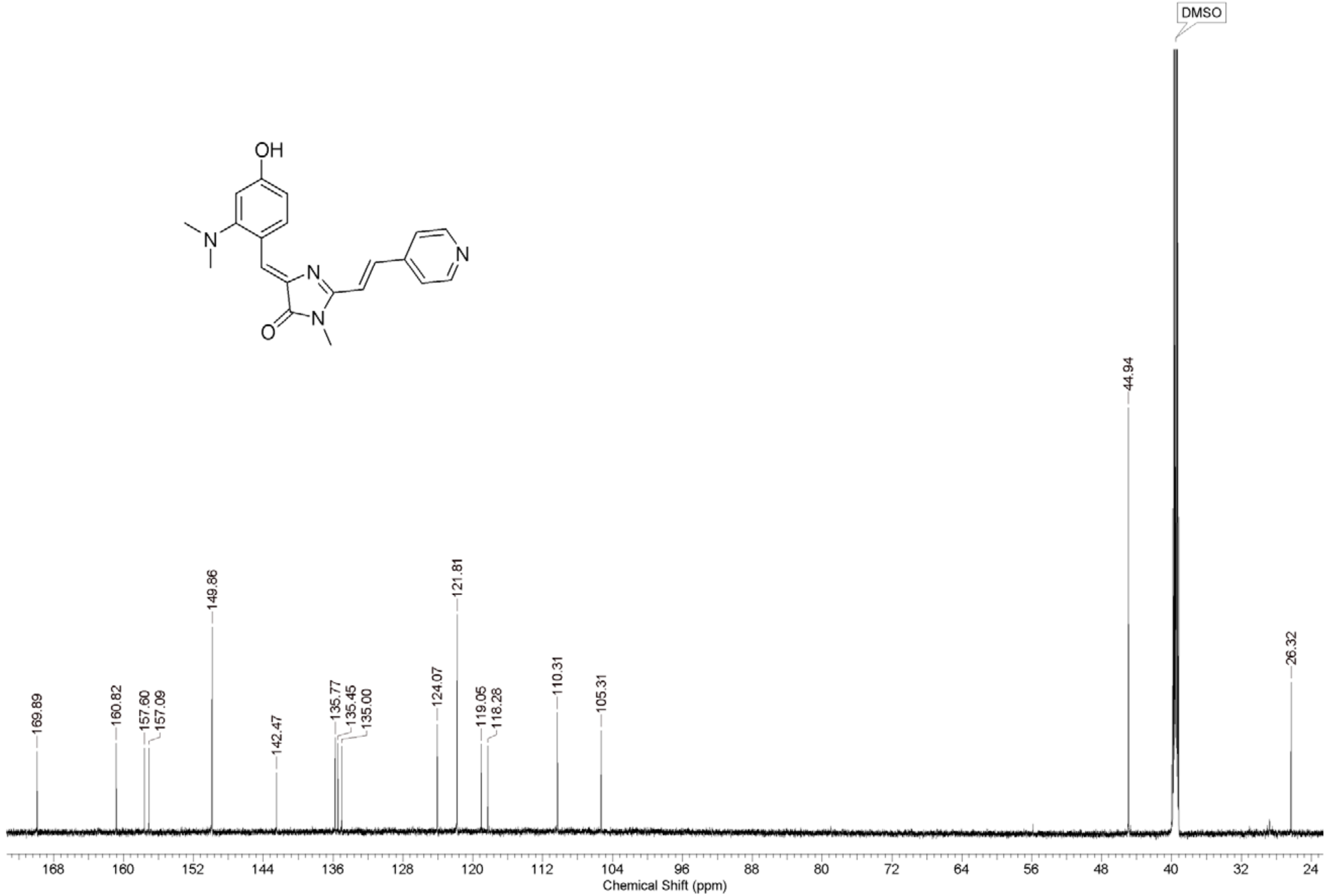
OH
N
N
N
N
O
DMSO
44.94
26.32
169.89
160.82
157.60
157.09
149.86
142.47
135.77
135.45
135.00
124.07
121.81
119.05
118.28
110.31
105.31
168 160 152 144 136 128 120 112 104 96 88 80 72 64 56 48 40 32 24
Chemical Shift (ppm)

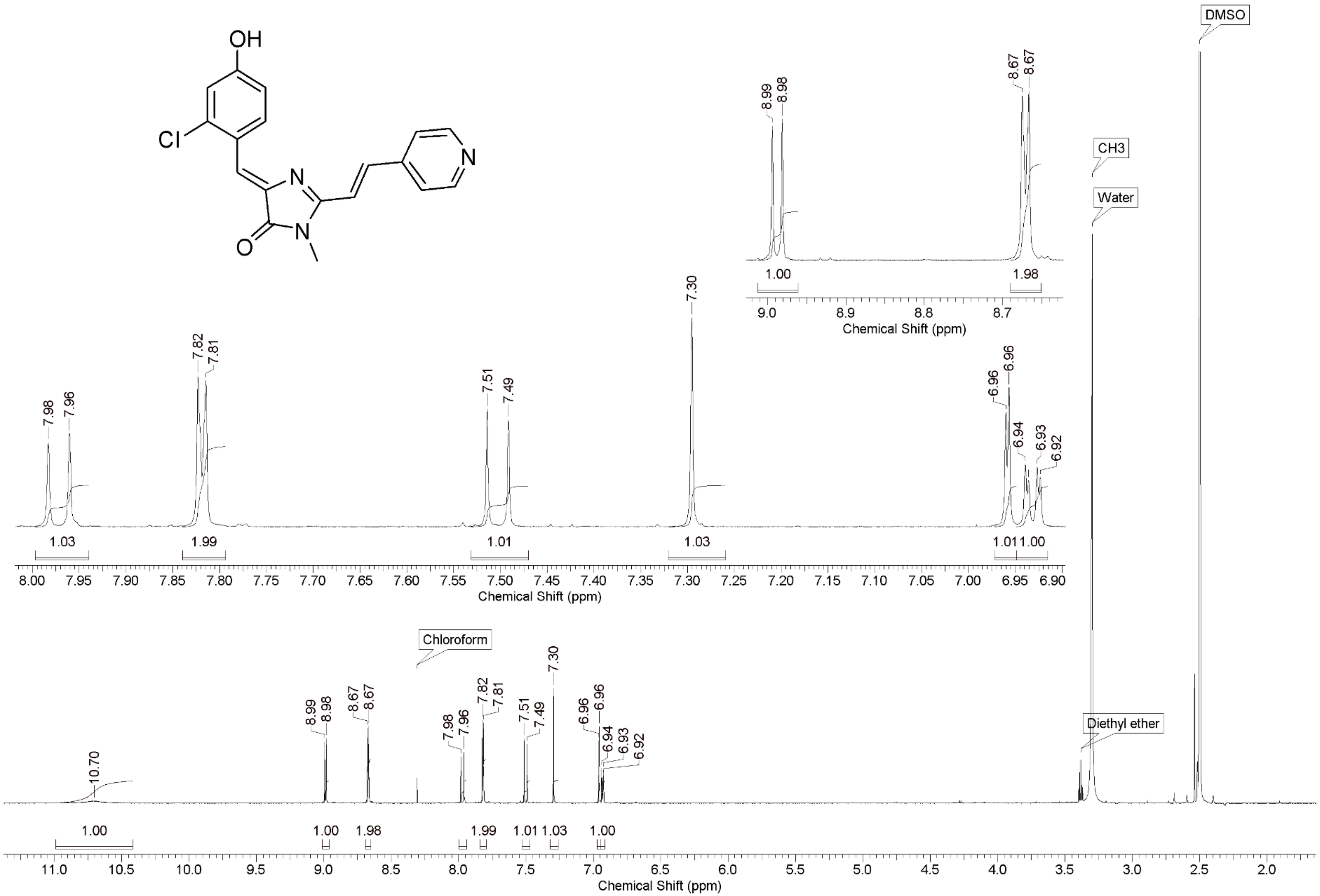
OH
Cl
N
N
O
N
DMSO
CH3
Water
Diethyl ether
Chloroform
8.99
8.98
8.67
8.67
1.00
1.98
9.0
8.9
8.8
8.7
Chemical Shift (ppm)
7.98
7.96
7.82
7.81
7.51
7.49
7.30
6.96
6.96
6.94
6.93
6.92
1.03
1.99
1.01
1.03
1.01
1.00
8.00 7.95 7.90 7.85 7.80 7.75 7.70 7.65 7.60 7.55 7.50 7.45 7.40 7.35 7.30 7.25 7.20 7.15 7.10 7.05 7.00 6.95 6.90
Chemical Shift (ppm)
10.70
1.00
1.00
1.98
1.99
1.01
1.03
1.00
11.0 10.5 10.0 9.5 9.0 8.5 8.0 7.5 7.0 6.5 6.0 5.5 5.0 4.5 4.0 3.5 3.0 2.5 2.0
Chemical Shift (ppm)

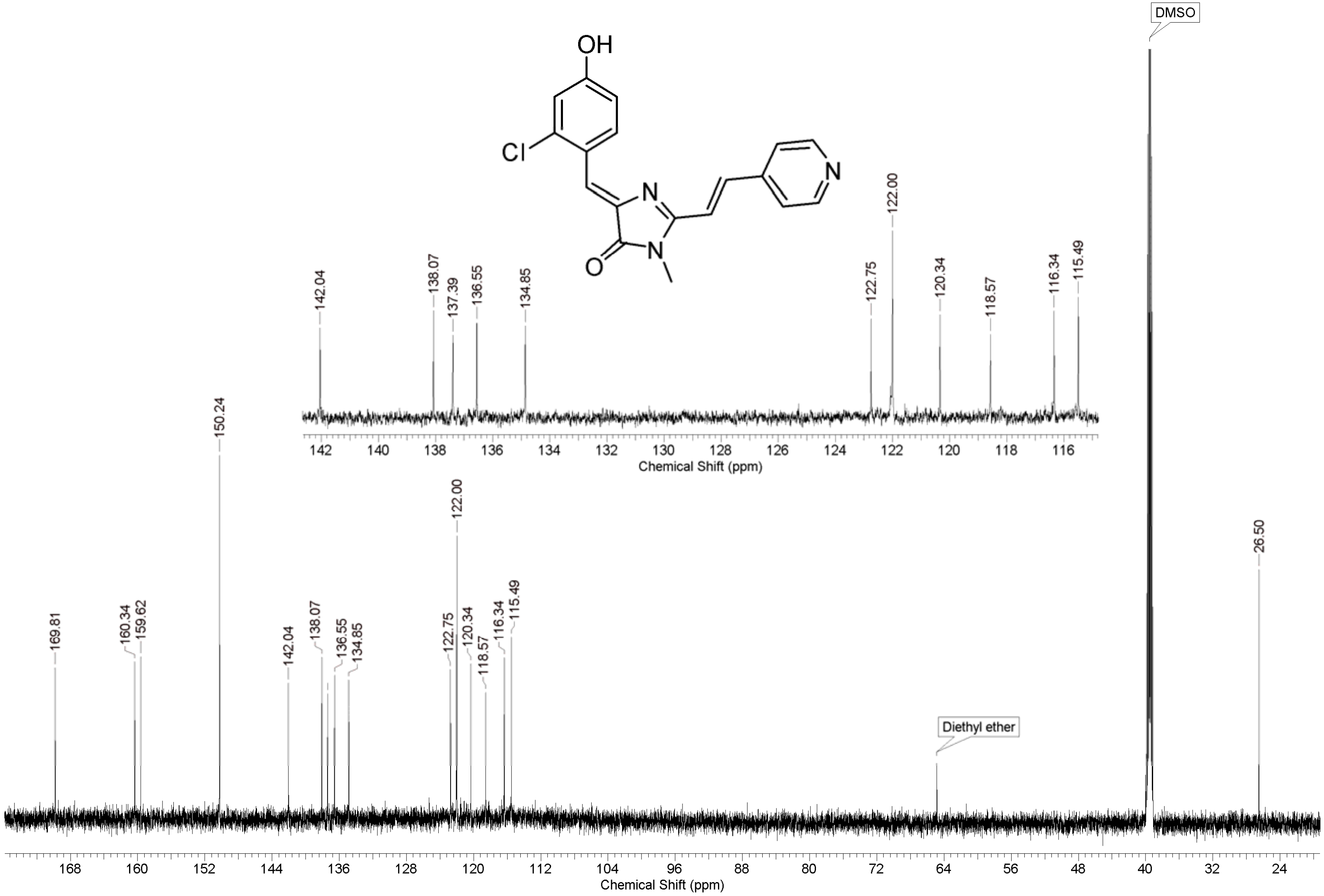
OH
Cl
N
N
O
DMSO
Diethyl ether
142.04
138.07
137.39
136.55
134.85
122.75
122.00
120.34
118.57
116.34
115.49
142 140 138 136 134 132 130 128 126 124 122 120 118 116
Chemical Shift (ppm)
169.81
160.34
159.62
150.24
142.04
138.07
136.55
134.85
122.75
122.00
120.34
118.57
116.34
115.49
26.50
168 160 152 144 136 128 120 112 104 96 88 80 72 64 56 48 40 32 24
Chemical Shift (ppm)

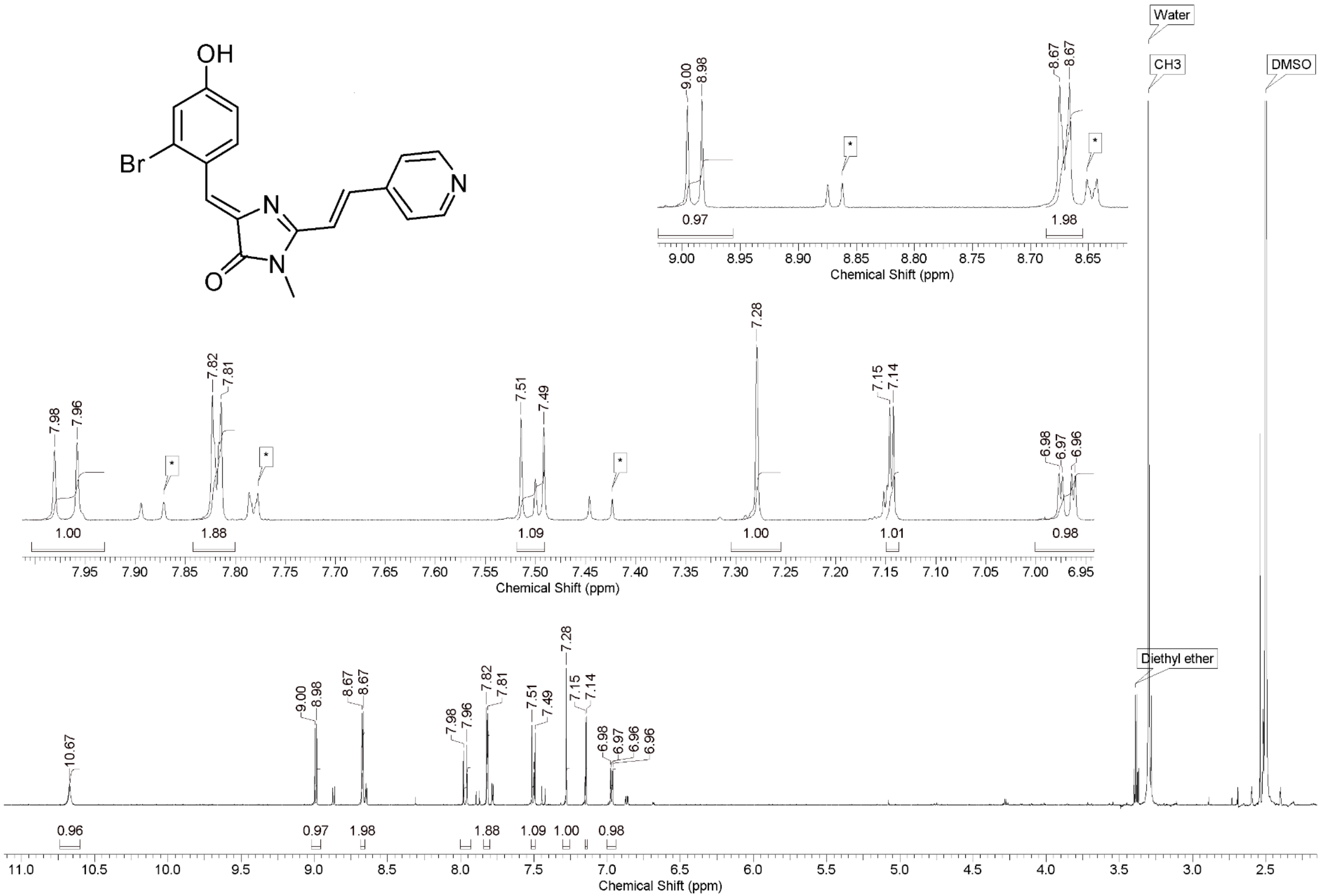
OH
Br
N
N
O
N
Water
CH3
DMSO
Diethyl ether
Chemical Shift (ppm)
Chemical Shift (ppm)
Chemical Shift (ppm)

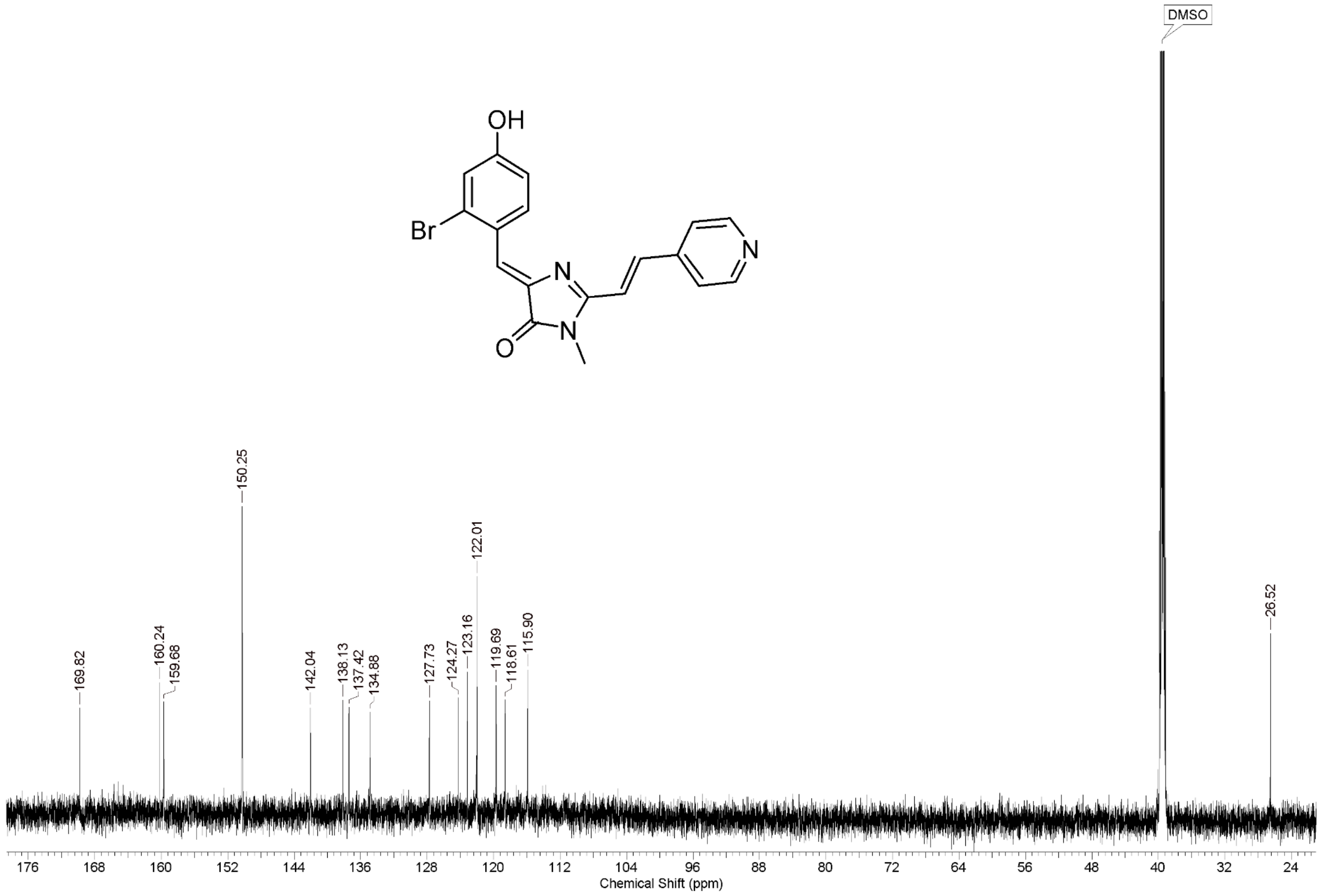
OH
Br
N
N
N
O
DMSO
169.82
160.24
159.68
150.25
142.04
138.13
137.42
134.88
127.73
124.27
123.16
122.01
119.69
118.61
115.90
26.52
176
168
160
152
144
136
128
120
112
104
96
88
80
72
64
56
48
40
32
24
Chemical Shift (ppm)

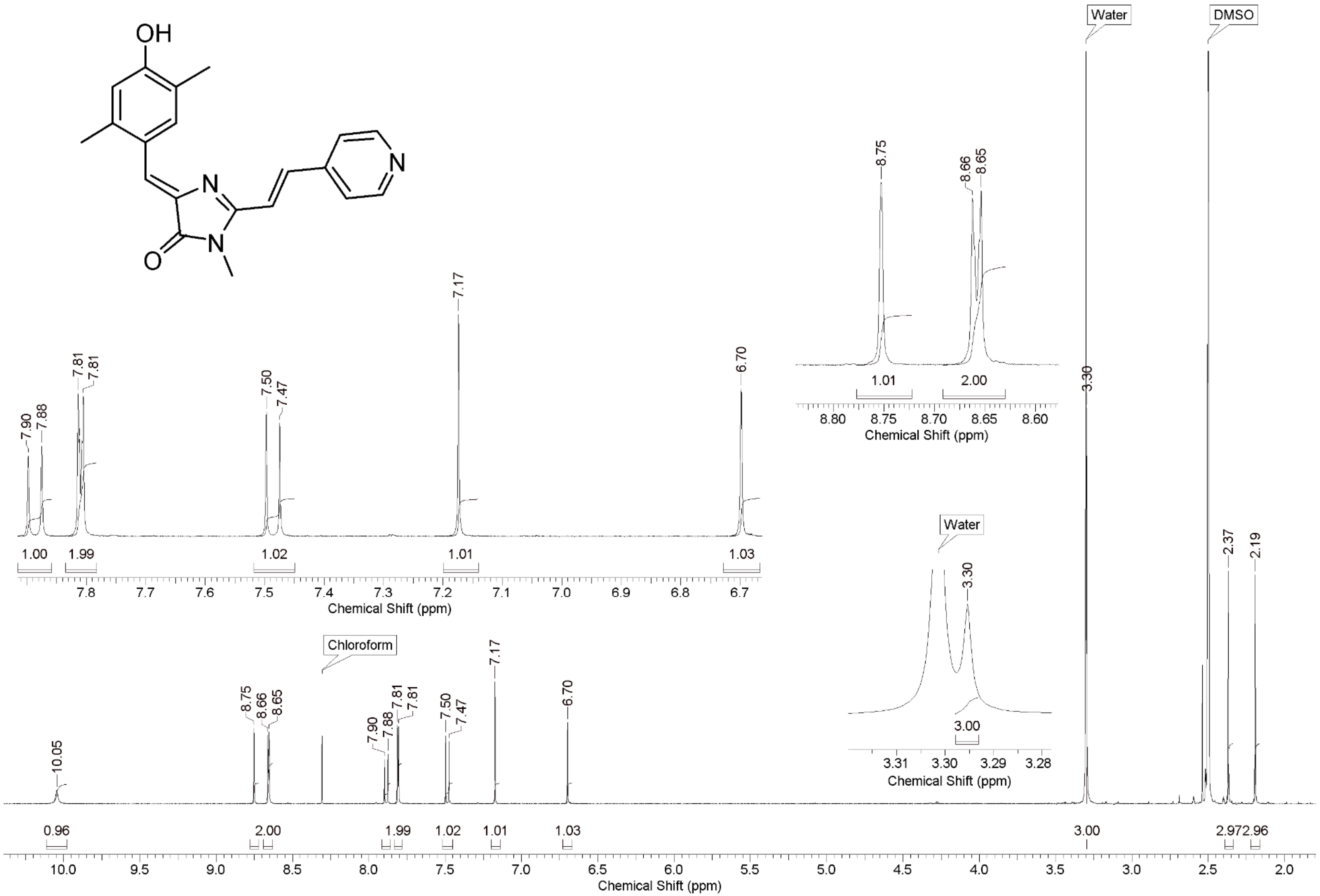
OH
N
N
O
N
Water
DMSO
Chloroform
10.05
8.75
8.66
8.65
7.90
7.88
7.81
7.81
7.50
7.47
7.17
6.70
3.30
2.37
2.19
0.96
2.00
1.99
1.02
1.01
1.03
3.00
2.97
2.96
1.00
Chemical Shift (ppm)

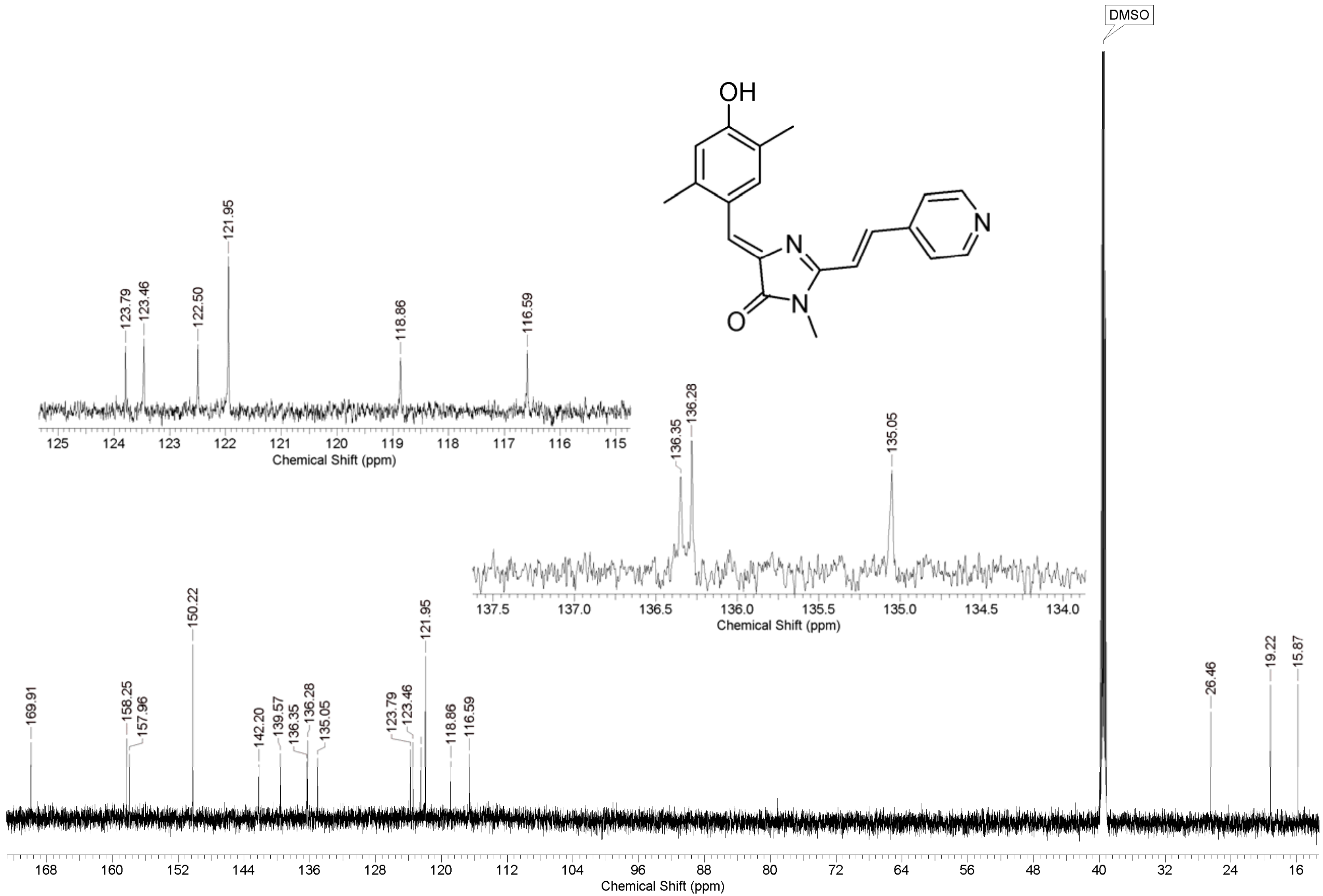
DMSO
OH
N
N
N
O
169.91
158.25
157.96
150.22
142.20
139.57
136.35
136.28
135.05
123.79
123.46
121.95
118.86
116.59
26.46
19.22
15.87
123.79
123.46
122.50
121.95
118.86
116.59
136.35
136.28
135.05
Chemical Shift (ppm)
